# Slow Electromechanical Interference Links Division Geometry, Differentiation Bias, and Senescence-like Arrest

**DOI:** 10.1101/2025.10.28.684574

**Authors:** Alfonso De Miguel Bueno

## Abstract

This work develops a biophysical theory in which a bioelectric field *V* (*x, t*) and a cortical stress field *σ*(*x, t*) are weakly and reciprocally coupled via an overdamped electromechanical coupler. We show that interference between two fast latent modes produces a measurable slow beat *f*_slow_ that acts as a tissue-level clock. By sampling the dynamics at “neutral moments”—recurring instants of phase symmetry—we derive a reduced even circle map in which healthy homeostasis corresponds to locking within a specific 2/21 Arnold tongue.

We then introduce a coarse-grained dual-field algebra that collapses the continuum description into three effective blocks (Γ, *a, b*) capturing net electromechanical gain and dissipation. In this algebraic picture, ionic and rheological perturbations are represented as smooth deformations of the parametrization space (Γ, *a, b*), while the clock variables (Ω, *f*_slow_, *K*_2_) provide experimentally accessible coordinates on those deformations. This construction offers a concrete bridge between molecular-scale regulation and tissue-level mechanics, connecting subcellular control to the emergent seconds–minutes slow clock that constrains division geometry.

Evaluating the membrane-potential profile at neutral moments defines a neutral charge-asymmetry observable Δ*Q*_*n*_ that quantifies left–right voltage imbalance at the division axis and links the slow-phase map to directly measurable bioelectric patterns. The same neutral-map construction admits a rotational interpretation in terms of circle maps and slow precession of the locked orbit: small detunings *δ* from the ideal 2/21 plateau generate a hierarchy of time scales and predict a scaling law *T*_dev_ ~ 1/(*f*_slow_ |*δ*|) relating the fast electromechanical beat to developmental timing.

The theory yields four falsifiable predictions. (P1) Homeostatic epithelia exhibit a narrow shared slow-band peak in voltage and stress with high coherence. (P2) The effective forcing and coupling (Ω, *K*_2_), derived from physical parameters, reside within the 2/21 tongue while avoiding broad low-order resonances. (P3) A weak, frequency-specific drive at *f*_slow_ (phase-targeted entrainment) selectively increases coherence and reduces spindle-angle dispersion in unlocked states, providing a physical basis for bioelectric modulation of regenerative dynamics. (P4) Across conditions with comparable *f*_slow_, the number of neutral compensation cycles required to complete a phenotypic transition scales inversely with the detuning |*δ*|, linking slow precession of the neutral map to macroscopic developmental time.

Ultimately, this framework treats cancer-like instability and senescence-like arrest not as independent pathologies, but as opposite failures of navigation in a single underlying electromechanical cycle, from persistent unlocking to rigid oversynchronization.

**Highlights:**

- Proposes an overdamped double-oscillator model of tissue electromechanics in which two fast latent modes generate a slow beat (*f*_slow_).
- Links the slow beat to an even circle map via neutral moments, identifying a specific 2/21 Arnold tongue that governs stable spindle orientation.
- Introduces a quantitative dual-field algebra in which ionic (pump-like) and rheological (stiffness-like) perturbations act as deformations of coarse-grained blocks (Γ, *a, b*), predicting matched shifts in Ω and *f*_slow_.
- Defines a neutral charge-asymmetry observable Δ*Q*_*n*_ at neutral moments, recasting the locking scenarios (P1–P3) as constraints on left–right membrane-potential imbalance that can be computed from existing bioelectric models.
- Relates the slow beat to capture kinetics *T*_lock_, linking fast carrier interference to mitotic (minute-scale) timing through progressive synchronization.
- Derives a scaling law (P4) in which the number of neutral compensation cycles required for a phenotypic transition scales inversely with the phase detuning |*δ*|, naturally generating a hierarchy of time scales from minutes to days.
- Models “proliferative unlocking” and “senescent overlock” as opposite dynamical failures of the dual-field clock (drift vs. rigidity), providing a unified view of cancer instability and aging.
- Validates a protocol for phase-targeted entrainment, predicting that a weak drive at *f*_slow_ selectively recovers coherence and reduces spindle-angle dispersion in unlocked states.

## 1 Introduction

Biological tissues exhibit coupled electrical and mechanical dynamics. Bioelectric potentials shape patterning, growth, and large-scale morphogenesis, while cortical tension and poroelastic stresses govern force transmission, division geometry, and tissue-scale mechanics [3, 5, 6, 8]. Controlled bioelectric cues can actively reshape tissues *in vivo* and *in vitro*, and multicellular responses to electrophysiological perturbations can be captured quantitatively by bioelectrical models [3, 4, 5]. Here we treat these two sectors explicitly as fields: a bioelectric field *V* (**x**, *t*) and a cortical stress field *σ*(**x**, *t*) that are weakly and *reciprocally* coupled.

In the fast band, the observable dynamics arise from *two weakly coupled first-order latent modes* that generate two damped carriers in the 1–3 Hz range. Their slight detuning produces a measurable slow beat *f*_slow_ = |Ω_+_ − Ω_−_|/(2*π*) at the mesoscale set by the effective thickness *𝓁* (i.e., *k*^∗^ = *π/𝓁*). This two-mode coupling succinctly accounts for the apparent second-order features in the measured time series–namely, voltage-imaging traces *V* (*t*) and traction/AFM-based stress reconstructions *σ*(*t*) in epithelial monolayers.

Sampling the dynamics at the *neutral moment* –the recurring near-symmetric crossing between *V* and *σ* where the relative phase *ψ* passes through 0 (mod *π*) with positive slope–yields an *even* circle map (mod *π*) for the slow phase. In this reduced description, the forcing Ω and the even-harmonic coupling *K*_2_ are defined directly from data: to first order,

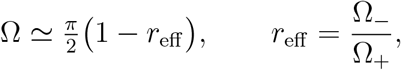

with *r*_eff_ obtained from the fast-band carriers, while *K*_2_ is estimated by neutral-moment regression.

Within this framework we propose that the slow electromechanical phase does not just exist spectrally, but coordinates two biological scales that are usually treated separately. On one side, it constrains spindle geometry and cortical anchoring during mitosis, limiting spindle-angle dispersion and chromatin torsional load at division checkpoints. On the other side, it provides a dynamical compatibility window for helical coordination. We focus on a narrow high-order lock, the 2/21 tongue, used here as a concrete target whose center is Ω_2/21_ = *π* · 2/21. We do not assume a direct forcing of molecular helical pitch by a tissue-scale cadence; any apparent alignment remains to be established empirically. The analysis concerns tissue-level observables (*f*_slow_, Λ, and (Ω, *K*_2_)), with cross-scale links treated operationally via the cycle-resolved neutral-reset analysis (App. E).

A central observable is the cross-spectral coherence Λ ≡ *C*_*V σ*_(*f*_slow_) between *V* and *σ* at *f*_slow_. High Λ indicates that the bioelectric and mechanical sectors share a common slow cadence and remain phase-locked across neutral moments, whereas low Λ indicates slow-cadence drift between sectors. Together with the neutral-moment map parameters (Ω, *K*_2_), Λ provides a concrete test of whether tissue sits within a specific high-order Arnold tongue (notably 2/21) and whether departures from that window correlate with pathological states.

To ground these phase dynamics in a measurable bioelectric quantity, we introduce the neutral charge-asymmetry observable Δ*Q*_*n*_, obtained by sampling the membrane-potential profile across the division axis at neutral moments with an antisymmetric weight. This quantity directly probes the antisymmetric electrical mode and links the abstract locking geometry to specific patterns of left–right voltage imbalance at the coupler.

Physiological tissue is not expected to sit forever in a perfectly locked, stationary regime. Instead, App. E formalizes a *cycle-resolved* analysis in which healthy tissue repeatedly *re-enters* the 2/21 band on each slow cycle. Operationally, we identify each neutral moment, compute local slow-band coherence around it, and track the sliding estimates (Ω, *K*_2_) over many cycles. Failure to re-enter takes two characteristic forms. First, *persistent unlocking* : the system drifts out of 2/21, Λ collapses, and spindle orientation becomes erratic. Second, *aging-like overlock* : the system remains trapped near the neutral state, Λ stays high but rigid, and adaptability is lost. In this reading, cancer-like genomic instability and aging-like loss of plasticity can be viewed as opposite dynamical arrests of the same electromechanical cycle.

Beyond mitotic timing, the same neutral-map construction admits a rotational outlook in terms of circle maps and slow precession of the locked orbit (App. L.2). Small detunings *δ* from the ideal 2/21 plateau generate a second slow scale: the precessive phase slip of the neutral orbit in function space. In a reduced description, this precession controls how rapidly neutral compensation cycles accumulate a phenotypic bias, leading to a scaling law

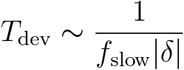

that links the fast electromechanical beat to developmental timing and to the number of neutral cycles required for a macroscopic phenotypic transition.

This framework addresses three persistent gaps in the current understanding of tissue morphogenesis, at the interface between molecular biology, bioelectricity, and mechanical physics. First, a *dimensional gap*: how high-dimensional molecular and cellular noise (arising from many ionic and cytoskeletal degrees of freedom) collapses into robust, lowdimensional transitions in cell state and tissue geometry. Second, a *temporal gap*: how fast bioelectric and mechanical dynamics on millisecond–second scales can instruct developmental processes unfolding over hours or days without invoking ad hoc accumulation counters. Third, a *mechanistic gap* between continuous physical fields and discrete genetic programs: how field-level constraints are converted into fate decisions and division patterns.

In our formulation, the neutral phase map acts as a low-dimensional attractor organizing high-dimensional electromechanical fluctuations, slow precession of the locked orbit provides a geometric origin for developmental time scales, and the neutral charge asymmetry Δ*Q*_*n*_ links the locking geometry to measurable left–right voltage imbalance at the division axis.

The theory yields three preregisterable predictions (P1–P3) with direct experimental readouts, together with a fourth scaling prediction (P4) that can be confronted with time-resolved division and differentiation data. P1 states that homeostatic epithelia exhibit two fast-band carriers together with a narrow shared slow-band peak and a coherence maximum Λ = *C*_*Vσ*_(*f*_slow_). P2 predicts that the forcing/coupling pair (Ω, *K*_2_), obtained from the fast-band carriers (for Ω) and neutral-moment stroboscopy (for *K*_2_), lies inside the predicted 2/21 Arnold tongue while avoiding broad low-order tongues. P3 posits that a weak, frequency-specific electrical or mechanical drive at *f*_slow_ should increase Λ and reduce spindle-angle dispersion relative to an energy-matched off-resonant control, providing a physical basis for bioelectric modulation of regenerative dynamics. P4 predicts that, across conditions with comparable *f*_slow_, the number of neutral compensation cycles required to complete a phenotypic transition scales inversely with the phase detuning |*δ*|, effectively relating *T*_dev_*f*_slow_ to 1/ |*δ*| .

App. G gives a preregistered assay for P3 and an *in silico* prevalidation in which a synthetic unlocked state is driven weakly at *f*_slow_, showing selective recovery of slow-phase locking (higher Λ) and reduced angular noise in the neutral map, while off-band controls at *f*_slow_(1± 0.10) fail to reproduce both effects together. Small persistent biases are also considered: a tonic offset breaks the *π*-symmetry of the neutral map, mixes parities, elevates even harmonics and (*f*_0_, *f*_0_ → 2*f*_0_) bicoherence, and produces a non-monotonic power versus coherence response (App. H).

Within P3 we also preregister three mechanistic subtests–**P3a** (phase-locked neutral crossings), **P3b** (transition sharpness vs. slow-band power), and **P3c** (bias-induced parity mixing and rectified pumping)–which share the same weak-drive paradigm and serve as falsifiers of linear, phase-preserving alternatives (see App. H and App. G).

Beyond the slow-phase map, App. K develops a complementary functional and algebraic formulation of the dual-field clock. In that framework, the coarse-grained parameters (Ω, *K*_2_, *f*_slow_) arise from three operator blocks (Γ, *a, b*) inherited from the overdamped electromechanical coupler, and small Na^+^/K^+^-like and pH/stiffness-like perturbations are represented as controlled deformations in this dual-field algebra. This provides an analytic explanation for the near-colinearity of the susceptibilities of Ω and *f*_slow_ and offers a compact language to interpret ionic and pharmacological interventions as trajectories in a constrained parameter space.

All analysis code, the preregistered workflow, and the cycle-resolved diagnostics are provided in App. E and App. G; the main text is self-contained.

## 2 Background and gaps the model addresses

Existing accounts of tissue coordination tend to split into two largely separate traditions. One focuses on molecular and statistical detail, treating voltage gradients, gene-regulatory programmes and double-helix structure as self-contained, with mechanics appearing only as boundary conditions. The other works at the scale of tissues and organs, modelling collective mechanics, division geometry or cortical tension without an explicit dual electrical– mechanical dynamics or a concrete link back to molecular carriers. Both perspectives lack a dynamical, directly falsifiable mechanism that can coordinate cell-cycle events and division geometry without relying on manual integer tuning.

Here we introduce a dual-field framework that is designed to address this gap. We treat the bioelectric field *V* (**x**, *t*) and the cortical-stress field *σ*(**x**, *t*) as weakly and reciprocally coupled. In the fast band, this interaction can be reduced to two first-order latent modes with slightly detuned damped carriers Ω_*±*_; their interference generates an emergent slow beat

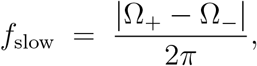

which we propose as a tissue-scale clock for division timing and spindle geometry. Crucially, *f*_slow_ is not introduced as a free parameter but is determined by the electrical and mechanical latencies of the two coupled sectors.

To turn this slow cadence into a predictive structure, we sample the fast electrical phase at recurrent near-symmetric crossings between *V* and *σ* (the *neutral moments*). Operationally, these are the instants at which the analytic-phase difference *ψ*(*t*) = arg *Z*_*V*_ (*t*) − arg *Z*_*σ*_(*t*) crosses 0 (mod *π*) with positive slope. Strobing at neutral moments yields an *even* circle map, defined modulo *π*, for the reduced phase variable *θ*_*n*_,

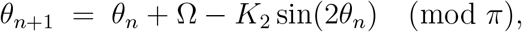

whose rotation number *ρ*_*π*_ and Arnold-tongue structure are entirely controlled by two parameters (Ω, *K*_2_). The effective forcing Ω is set by the ratio of the two carriers, *r*_eff_ = Ω_−_/Ω_+_, via 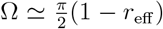, and the even-harmonic coupling *K*_2_ is estimated by neutral-moment regression on the observed strobe. In this setting, the high-order lock *ρ*_*π*_ = 2/21 used throughout the paper is not imposed as an ad hoc quantization rule; it *emerges* from the even map once the carriers have been measured.

Because the map is even (mod *π*), the same construction naturally distinguishes between symmetric (latency-matched) and desynchronized (latency-mismatched) regimes. Neutral-moment strobing enforces even parity and reveals a specific locking pattern; full-cycle strobing (odd parity) is predicted to degrade that structure in a characteristic way. This built-in parity control provides an internal falsification test: if the proposed mechanism is correct, mode locking and curvature of the circle map should be visible under neutral-moment sampling and should change in a parity-sensitive manner when sampling is shifted away from neutral moments.

Finally, all quantities entering the reduced slow-phase map are traceable to measurable tissue properties. Membrane time constants, cortical stiffness and viscosity, diffusivities (*D*_*V*_, *D*_*σ*_) and weak electromechanical gains (*γ, β*_*R*_, *β*_*I*_) constrain the latent carriers and their ratio *r*_eff_, and thus Ω. The coupling *K*_2_ is obtained from data by neutral-moment regression. This parameter provenance enables preregistered tests of specific locking scenarios, rather than post hoc tuning of integers or free phases.

### 2.1 Relation to prior bioelectric models

Classical bioelectric frameworks emphasize voltage gradients, ionic currents and patterning cues, often without an explicitly reciprocal mechanical field. In many such models, mechanics appears only as a static substrate, and no slow interference cadence is identified. In contrast, the present dual-field reduction

i. treats *V* and *σ* as weakly but reciprocally coupled degrees of freedom,
ii. predicts two narrow-band carriers in the fast band together with a slow interference beat *f*_slow_ that is, in principle, directly measurable rather than inferred,
iii. yields an *even* circle map at neutral moments, providing a built-in parity switch (even vs. odd strobing) that acts as an internal falsification control,
iv. computes the effective forcing Ω and the even coupling *K*_2_ from directly measurable electrical and mechanical properties of the tissue, with no ad hoc integer quantization, and
v. defines a functional order parameter Λ = *C*_*V σ*_(*f*_slow_) as the cross-spectral coherence between *V* and *σ* at the slow beat, providing a compact way to test whether different observables (slow-band voltage power, slow-band stress power, division geometry) are governed by the same underlying electromechanical locking.
vi. introduces a neutral charge-asymmetry observable Δ*Q*_*n*_ by sampling the membrane potential at neutral moments, providing a direct way to reinterpret locking, proliferative unlocking and senescent overlock as distinct patterns of left–right voltage imbalance along the division axis.

In this sense, our model is intended to complement existing bioelectric accounts of growth control, morphogenesis and tissue-scale coordination, and to align with recent work showing that bioelectric control of tissue shape and collective responses to electrophysiological perturbations can be quantified and tested in a predictive manner [3, 7, 4, 5].

## 3 Physical identity of the fields and oscillatory reduction

### 3.1 Latent-mode representation and slow-beat emergence

In the fast band we represent the coupled bioelectric and mechanical dynamics by two weakly coupled first-order latent modes. In complex-amplitude form,

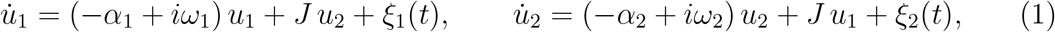

where *α*_*k*_ *>* 0 are decay rates, *ω*_*k*_ *>* 0 are carrier frequencies, *J* is a small complex coupling, and *ξ*_*k*_(*t*) collect fast noise and residual driving. The measured channels are real projections of these modes,

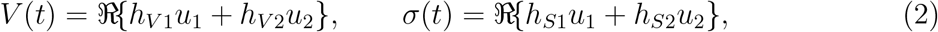

with complex coefficients *h*_(*·*)*k*_ encoding the mixing between latent modes and observed fields. Apparent second-order behaviour of the fast band is captured here by the two-mode coupling itself: no extra “hidden” oscillator is introduced beyond Eqs. (1)–(4).

Writing 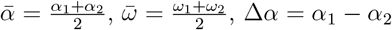 and Δ*ω* = *ω*_1_ − *ω*_2_, the four eigenvalues of the 4 × 4 real system can be written as

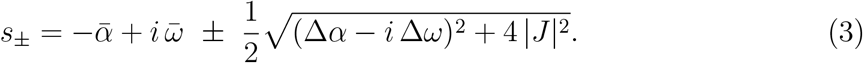

Generically this yields *two* damped carriers with observable frequencies Ω_*±*_ = ℑ *s*_*±*_. In the symmetric case Δ*α* = 0,

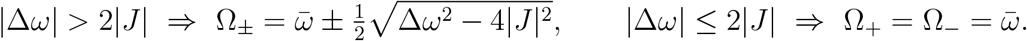

The emergent slow beat is then

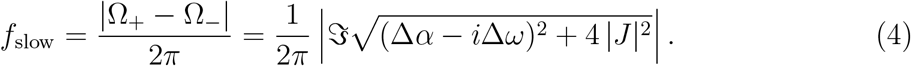

Thus, two weakly coupled first-order modes are sufficient to account for the fast-band structure. When |Δ*ω*| *>* 2|*J*| the spectrum exhibits two damped carriers Ω_*±*_ with a slow beat *f*_slow_ = |Ω_+_ − Ω_−_ |/(2*π*); under locking (|Δ*ω*| ≤2 |*J*|), the split collapses to a single carrier and *f*_slow_ → 0. In this sense, the “effective second order” of the observed response is generated by the two-mode coupling rather than by an ad hoc second-order term.

In what follows we refer to the regime with two distinct carriers and ongoing relative precession between the two sectors as the *antisymmetric, latency-mismatched* regime. When the carriers merge into a single frequency and the electric and mechanical sectors stop precessing against each other, we speak of the *symmetric, latency-matched* regime.

#### Observed phases and projection structure

If the projections are cross-dominant, |*h*_*V* 1_| ≫ |*h*_*V* 2_| and |*h*_*S*2_| ≫ |*h*_*S*1_|, each channel follows, in the antisymmetric latency-mismatched regime, the phase of the latent mode that dominates its projection. Under locking, approximately odd/even combinations *h*_*V* 1_ ≈ − *h*_*V* 2_ and *h*_*S*1_ ≈ *h*_*S*2_ produce phase opposition in *V* and phase alignment in *σ* for the locked solution, matching the phenomenology we seek to explain at the tissue level.

In the antisymmetric, latency-mismatched regime, dominance between the two sectors alternates over each slow cycle: during one half-beat the electrical projection is closer to the fast carrier, whereas during the other half-beat the mechanical projection leads. This periodic exchange of roles preserves the slow beat while preventing either sector from becoming permanently dominant. In the symmetric, latency-matched regime one sector effectively freezes the relative phase and the alternation collapses, a point that will be important below when we discuss proliferative versus senescent regimes.

#### Role of nonlinearity (P3)

Nonlinearity does not create the carriers; it mixes and rectifies their interference under a weak tonic bias *ε*_*b*_. This raises even harmonics, enhances the (*f*_+_, *f*_+_ → 2*f*_+_) bicoherence and transmits the cadence *f*_slow_ into the slow map. In other words, nonlinearity is needed to *read out* the beat, not to generate it.

#### Falsifiable check for two damped carriers

The presence of two complex poles can be tested directly. A brief, spatially uniform impulse followed by traction or stress readout should exhibit a short-lived oscillatory ring-down in the carrier band if the spectrum contains two damped modes. Fitting the impulse response with a single-carrier versus a two-carrier model discriminates whether a split consistent with Eqs. (3)–(4) is present. In the antisymmetric, latency-mismatched regime (|Δ*ω*| *>* 2|*J*|), Hilbert-phase drift between *V* and *σ* provides an independent estimate of *f*_slow_ and its collapse when |Δ*ω*| ≤ 2|*J*|.

### 3.2 Phase-map reduction and parity structure

In the antisymmetric, latency-mismatched regime the two damped normal modes at *k*^∗^ = *π/𝓁* alternate in dominance. We define the *neutral moment* as the recurrent instant when the instantaneous phase difference between *V* and *σ* is 0 (mod *π*) with positive slope,

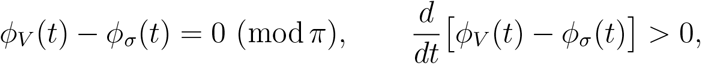

so that the signals are locally mirror-symmetric even if amplitudes remain unbalanced. Operationally, this marks a transient balance between inward (electrical) and outward (mechanical) tendencies and resets the slow cadence.

Taking the neutral moments as a Poincaré section identifies *θ* ≡ *θ* +*π*. In this *π*-quotient the leading nonlinearity is even, and the reduced slow-phase map is

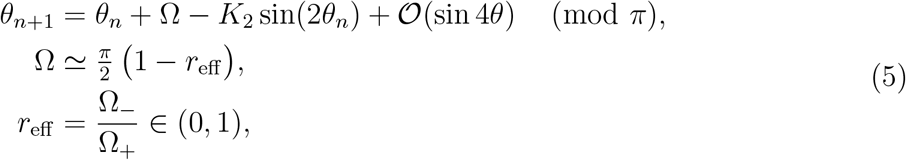

where *r*_eff_ is the carrier-frequency ratio obtained from Eqs. (1)–(4). The forcing Ω is therefore fixed by the measured carriers, and the even-harmonic coupling *K*_2_ is estimated by neutral-moment regression on the observed strobe.

#### Type of nonlinearity and empirical checks

Because neutral-moment strobing identifies *θ* with *θ* + *π*, the reduced map Eq. (5) must be parity-even. The leading nonlinear correction is therefore quadratic and appears as −*K*_2_sin(2*θ*), with higher even orders 𝒪(sin 4*θ*). Odd harmonics (e.g. sin *θ*) are symmetry-forbidden in the unbiased regime and only emerge when parity is broken by a sustained bias (Sec. H.4). In the Supplementary Material we report two compact empirical checks supporting this classification: the even/odd spectral ratio *P* (2*f*_+_)*/P* (3*f*_+_) and the normalized bicoherence at (*f*_+_, *f*_+_ →2*f*_+_) with phase-randomization *p*-values.

#### Bias and parity mixing

A small tonic bias (electrical DC offset or mechanical prestress) does not remove slow alternation; it perturbs the reduced phase map by mixing even and odd harmonics. To leading order,

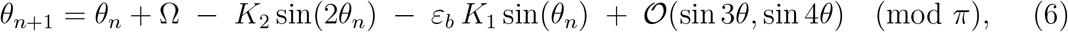

where *ε*_*b*_ measures the tonic offset and *K*_1_ is the odd-parity projection at neutral moments. In the 2*π* lift of the section the odd term is 2*π*-periodic; Eq. (31) (Sec. H.4) gives the explicit form used in the rectification analysis. The odd correction skews dwell times around neutral moments and *rectifies* the phase-gated exchange, increasing slow-band power and reducing adaptability as the operating point approaches tongue boundaries. Here Ω and *r*_eff_ are obtained directly from the fast band, and *K*_2_ is estimated by neutral-moment regression, with no ad hoc integer fitting.

Within the *π*-circle convention, a *p/q* phase lock is centered at Ω ≃ *π p/q*. For 2/21,

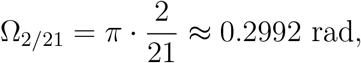

so the 2/21 plateau we focus on is a high-order Arnold tongue of the even (mod *π*) circle map in the standard sense of driven phase locking [1, 2].

When the lagged/quadrature component locks to the in-phase component (symmetric, latency-matched regime; the two sectors stop precessing against each other and the two carriers merge into a single synchronized oscillation), the *π* identification is lost. The relevant section becomes 2*π*-periodic and the map reverts to odd parity,

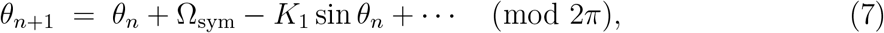

with constant forcing Ω_sym_ defined by the locked fast carrier. In the biological interpretation developed below, the antisymmetric, latency-mismatched regime underlies normal proliferative cycling, whereas progressive bias and eventual symmetry lead to “overlock” and senescence-like arrest.

## 4 Even circle map and rotation number

After neutral-moment reduction, the slow phase dynamics of the coupled bioelectric– mechanical system are captured by the even (mod *π*) circle map

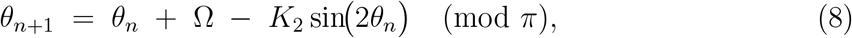

where Ω is the effective forcing set by the carrier-frequency ratio *r*_eff_ = Ω_−_/Ω_+_ and *K*_2_ is the even-harmonic coupling obtained by neutral-moment regression (Sec. 3). Eq. (8) describes how the slow phase *θ*_*n*_ advances from one neutral moment to the next.

Let {Θ_*n*_} be a lift of Eq. (8) to ℝ, so that Θ_*n*+1_ − Θ_*n*_ = Ω − *K*_2_ sin(2Θ_*n*_). The associated rotation number on the *π*-quotient is

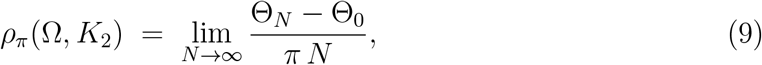

whenever the limit exists (which it does for circle maps with lifts of bounded variation). Intuitively, *ρ*_*π*_ measures the average phase advance (in units of *π*) per neutral moment; in our biological interpretation it controls the average angular offset of spindle orientation and the effective helical pitch of the integrated dynamics.

At fixed *K*_2_, the function Ω ↦ *ρ*_*π*_(Ω, *K*_2_) is a nondecreasing “devil’s staircase” [2, 10]: it is constant on mode-locked plateaus at rationals *ρ*_*π*_ = *p/q* and strictly increases only on a Cantor set of Ω values. Each rational *p/q* defines an Arnold tongue in the (Ω, *K*_2_)-plane, within which the map is locked with average rotation *ρ*_*π*_ = *p/q*. For *K*_2_ ≪ 1, the width of the tongue associated with *p/q* scales generically like 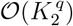, so higher-order locks become progressively narrower but can still be robust if *K*_2_ is not vanishingly small. In the uncoupled limit *K*_2_ = 0 one recovers the trivial relation *ρ*_*π*_ = Ω*/π*.

Within this even (mod *π*) convention, the lock of interest in this work is *ρ*_*π*_ = 2/21. It corresponds to a narrow high-order tongue centred at Ω ≃ *π* · 2/21 ≈ 0.2992 rad, and it implies that the slow phase advances by 2*π/*21 per neutral moment. In subsequent sections we show that this 2/21 locking regime organizes both spindle-orientation statistics and P1– P3 timing, and we outline how its presence can be tested numerically and experimentally.

## 5 Helical quantization without fitted integers

In the even (mod *π*) convention, a neutral-moment step advances the slow phase on average by *ρ*_*π*_*π* radians. For a rational lock *ρ*_*π*_ = *p/q, p* neutral cycles correspond to a net phase advance of *π*, and *q* neutral cycles produce an advance of *pπ*. Defining a helical unit as *p* neutral-moment increments that yield a mechanical half-turn (*π*), the number of units per full 2*π* turn is

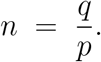

The value *n* ≈ 10.5 corresponds numerically to the rational *ρ*_*π*_ = 2/21.

In the present framework, the 2/21 plateau is an emergent locking regime of the even map, determined by the carrier ratio and the coupling (Ω, *K*_2_) inferred from tissue measurements. The coincidence with the B-DNA pitch is obtained without introducing fitted integers or explicit slow forcing at the molecular scale. We regard this as a cross-scale numerical correspondence; no direct dynamical coupling between the tissue beat and individual base pairs is assumed.

We ask whether physiologically plausible parameters can place the system inside the 2/21 tongue while avoiding low-order locks.

### Observation (Compatibility of physiological strips with 2/21 locking)

Let independently measured parameters induce a strip

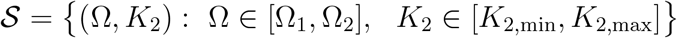

in the (Ω, *K*_2_) plane via Eqs. (1)–(4) and (8). For ranges consistent with homeostatic epithelia, numerical continuation shows that 𝒮 can intersect the 2/21 tongue of the even map while remaining separated from lower-order tongues with *q* ≤ 6. If, for a given tissue, the empirically derived strip 𝒮 does not intersect the 2/21 tongue, the 2/21 scenario is not supported under those conditions.

In practical terms, once (Ω_1_, Ω_2_) and (*K*_2,min_, *K*_2,max_) have been constrained from data, either 𝒮 intersects the 2/21 tongue or it does not; no additional integer fitting is available to enforce that lock.

## 6 An experimental coherence observable

Let *S*_*V*_ (*f*) and *S*_*σ*_(*f*) denote the power spectral densities of *V* and *σ*, and let *S*_*V σ*_(*f*) be their cross-spectrum. We define the electromechanical coherence at the slow cadence as

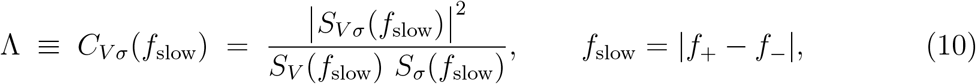

with *f*_slow_ given by Eqs. (3)–(4). The quantity Λ ∈ [0, 1] is a directly measurable order parameter that quantifies how strongly the electrical field *V* and the cortical stress *σ* are phase-locked at the slow beat.

Practically, *V* can be recorded with voltage-sensitive dyes or fast optical electrophysiology, and *σ* can be estimated from traction force microscopy or AFM-based stress reconstruction in epithelial monolayers. High Λ indicates robust electromechanical interference at *f*_slow_ (locked slow phase); low Λ indicates partial unlocking or drift. The definition of Λ does not rely on the circle-map description and can therefore be used as an independent test of the proposed locking scenario.

## 7 Mechanistic link to division and differentiation

Within the dual-field framework, the position of (Ω, *K*_2_) relative to the relevant Arnold tongue (here, the 2/21 tongue of the even map) controls slow-phase locking. When (Ω, *K*_2_) lies inside the tongue, the slow phase is locked, the cross-spectral coherence Λ at *f*_slow_ is high and stable, and spindle-angle dispersion is predicted to be low. When (Ω, *K*_2_) approaches tongue boundaries or moves outside, locking becomes fragile or is lost: Λ decreases, and division-plane statistics broaden.

We interpret proliferative and differentiated states as operating at different locations in this control plane. Differentiated, homeostatic epithelia correspond to effective parameters (Ω, *K*_2_) within a locked region, with high Λ and low spindle error. Proliferative states are expected to hover near tongue edges, where small parameter shifts can induce partial unlocking. Two distinct failure modes follow from the same slow-phase mechanism: a regime of *proliferative unlocking*, with persistent drift of (Ω, *K*_2_) away from the tongue and low Λ, and a regime of *overlock*, in which locking remains strong but adaptability is reduced.

### 7.1 Testable pathology signatures

#### Aging (overlock / drift)

*Prediction*. In tissues from aged donors or long-term cultures, neutral-moment strobing should yield higher effective *K*_2_ and slow drift in *r*_eff_, with Λ remaining high but changing only slowly over time. Under full-cycle (odd-parity) strobing, the curvature structure associated with even harmonics is expected to be only weakly disrupted. *Refutation*. No systematic differences in (Ω, *K*_2_, Λ) relative to young controls.

#### Proliferative unlocking

*Prediction*. In hyperproliferative monolayers with impaired checkpoint control, (Ω, *K*_2_) should display a broad distribution of Ω values displaced from *π* · 2/21, Λ at *f*_slow_ should be low, and spindle misorientation should increase. Under parity switching (even to odd), only limited additional change is expected, because the even-harmonic organization is already degraded. *Refutation*. Persistently high Λ with (Ω, *K*_2_) remaining inside the 2/21 tongue despite unchecked proliferation.

These two signatures correspond, in parameter space, to different displacements of (Ω, *K*_2_) away from the interior of the 2/21 tongue, together with predictable changes in Λ and division-plane statistics.

#### Parameter-grounded view

Independent of any specific gene-expression program, aging-related changes in tissue mechanics and electrophysiology are expected to move (Ω, *K*_2_). For example, cortical or ECM stiffening and increased viscous loss (*κ*↑, *η*↑) act through *b* = *κ* + *D*_*σ*_*k*^∗2^ to lower Ω and strengthen even-harmonic feedback (larger *K*_2_). Electrical remodeling (changes in gap-junctional coupling *D*_*V*_ or membrane time constant *τ*_*m*_) can bias Ω in either direction. The net displacement is empirical and, in principle, quantifiable through (Ω, *K*_2_, Λ) under neutral-moment stroboscopy.

### 7.2 Phase-guided modulation (phase-targeted entrainment)

If a partial phase-locking capacity is present, the same slow-phase structure suggests a simple control strategy: low-amplitude entrainment at *f*_slow_ aimed at moving (Ω, *K*_2_) back towards the interior of the locking tongue. In this view, (Ω, *K*_2_, Λ) estimated in vivo provide both a baseline and a target region in the control plane.

A practical protocol would proceed in three steps: (i) estimate (Ω, *K*_2_, Λ) from voltage imaging and traction/stress readouts using neutral-moment stroboscopy; (ii) apply a weak electrical or mechanical drive at *f*_slow_, with detuning chosen to lie well inside the inferred tongue; and (iii) test for an increase in Λ and a reduction in spindle misorientation relative to matched off-resonant controls. The weak-drive Popper test (Methods) rules out purely linear alternatives and justifies on-band, low-amplitude stimulation.

### 7.3 Regenerative bioelectrical stimulation and dynamical specificity

Low-frequency electrical stimulation is used empirically in regenerative settings (dental pulp, bone, peripheral nerves) with carrier frequencies in the spanning the sub-Hz to tens of Hz range. Reported protocols typically arise from screening and differ across tissues, with no shared mechanism for why a given frequency is effective in one context and not in another.

In the dual-field framework the relevant quantity is not a universal carrier frequency but the tissue-specific slow cadence *f*_slow_ emerging from electromechanical interference. Successful stimulation is expected when the effective drive envelope lies inside the interior of the locking tongue for the target tissue. In particular, a fixed-frequency protocol may be pro-regenerative in one preparation and ineffective in another if the corresponding (Ω, *K*_2_) place the two tissues in different regions of the control plane.

A testable prediction follows. Outcomes of regenerative protocols should correlate more strongly with individualized matching of the slow envelope to *f*_slow_ (with (Ω, *K*_2_) inside the interior of the relevant tongue) than with fixed carrier frequency alone. In crossover designs using the same hardware, moving the envelope from off-band to on-band, while holding total delivered charge and carrier constant, is predicted to improve efficacy if the locking mechanism is operative.

## 8 Falsifiable experimental targets

### P1 Spectral coherence (slow band, even parity)

Homeostatic monolayers are expected to exhibit a coincident slow-band peak in *S*_*V*_ and *S*_*σ*_ at *f*_slow_ = |*f*_+_ − *f*_−_|, together with elevated cross-coherence Λ ≡ *C*_*V σ*_(*f*_slow_) *>* 0.6 estimated as in Methods (band-limited Welch, neutral-moment strobing; 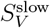 and 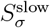 as defined there). Failure to detect a slow-band peak at *f*_slow_, or observing Λ *<* 0.2, falsifies the dual-field locking picture for that tissue and set of conditions, or indicates that the sample is in a non-homeostatic regime (unlocking or overlock).

In the latency-based refinement, homeostatic preparations that satisfy P1 are also expected to show latency parity consistent with even synchrony, with the electric and mechanical relaxation times lying on the same side of *T*_slow_ and of comparable magnitude (latency-parity indices *S*_lag_ *>* 0 and *D*_lag_ ≈ 1 in the notation of the Appendices). Persistent *S*_lag_ *<* 0 together with low slow-band coherence would instead signal an odd-desynchronized regime, i.e. effective unlocking or overlock of the dual-field oscillator.

### P2 Mode selection in (Ω, *K*_*2*_) without integer fitting

Independently measured physical parameters are used to compute the two carriers and their beat (Eqs. (1)– (4)), and the slow forcing via the even map (Eq. (8)), yielding a point (Ω, *K*_2_) in the control plane. The model predicts that (Ω, *K*_2_) falls inside the 2/21 Arnold tongue and avoids lower-order tongues, provided that the latency-based coupling proxy *C*_proxy_ (App. F.3) is not collapsed toward zero (near-quadrature mismatch), so that even-harmonic coupling at neutral moments remains effective.

### P2 (Quantitative)

P2 holds if (Ω, *K*_2_) lies within the 2/21 tongue and the distance in Ω to the nearest tongue centre with *q* ≤ 6 exceeds 3 *σ*_Ω_. In addition, a Monte Carlo under physiological priors places (Ω, *K*_2_) inside 2/21 with probability ≥ 0.30. This assumes negligible low-order overlap for *K*_2_ ≪ 1; if local overlaps occur, the exclusion margin is computed with respect to the nearest overlapping boundary instead of the tongue centre. A systematic displacement of (Ω, *K*_2_) outside 2/21, robust locking to a lower-order tongue, or a consistently collapsed *C*_proxy_ ≈ 0 falsifies the proposed selection mechanism.

### P3 Phase-targeted entrainment (weak, frequency specific)

A weak external electrical or mechanical drive at *f*_slow_ is predicted to increase Λ and decrease spindle-angle dispersion (proxy for spindle-orientation noise), whereas an off-resonant control of matched energy and duty cycle but ~10% detuning should not produce both effects simultaneously. Operationally, preregister the contrasts

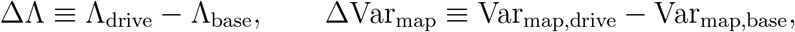

with the “weakness” constraint |Δ*f*_*±*_| *<* 3%. Absence of a selective on-band improvement in both ΔΛ and ΔVar_map_ relative to off-band controls falsifies the phase-targeted entrainment mechanism.

App. G provides the preregistration and the in-silico prevalidation.

### P3 (Refined capture-time prediction)

In the latency-based, first-order formulation the capture time is further constrained by the inter-channel phase at the slow cadence. Let *τ*_e_, *τ*_m_ be the electric/mechanical relaxation times, and *C*_proxy_ the latency-based coupling proxy defined in App. F.3. Near a tongue boundary the slow locking time obeys the scaling

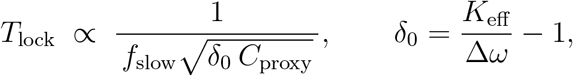

so that on-band weak drive should shorten *T*_lock_ primarily by increasing *C*_proxy_ (improved latency parity and coincidence), whereas an equal-energy off-band drive should not. Empirically, pro-meta and metaphase dwell times are expected to regress against 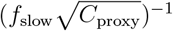 with approximately unit slope on log-log axes; systematic deviations or loss of on-band superiority relative to off-band controls falsify the refined capture picture.

## 9 Materials and methods

### 9.1 Electrical and mechanical parameters

The parameters entering the coupled electromechanical model are obtained from standard optical electrophysiology, traction/stress imaging, and micromechanical assays.

The electrical diffusivity *D*_*V*_ and membrane time constant *τ*_*m*_ are estimated from voltagesensitive dyes / fast optical electrophysiology by fitting a diffusive RC response: spatial decay of *V* constrains *D*_*V*_, and step responses constrain *τ*_*m*_ (after correcting for dye kinetics). The in-plane elastic modulus *κ*, viscous coefficient *η*, and stress diffusivity *D*_*σ*_ are obtained from AFM indentation, traction force microscopy (TFM), and poroelastic/viscoelastic relaxation fits. In this overdamped framework, *κ* captures effective cortical stiffness in the carrier band, *η* quantifies viscous loss in the ~1–3 Hz band, and *D*_*σ*_ characterizes stress redistribution.

The cross-coupling coefficients (*β*_*R*_, *β*_*I*_, *γ*) are inferred from linear-response gains and phase lags around the mechanical carrier band: *β*_*R*_ and *β*_*I*_ describe the in-phase and quadrature components of the stress-to-voltage gain, and *γ* the voltage-to-displacement gain. Together with *D*_*V*_, *τ*_*m*_, *κ, η, D*_*σ*_, these parameters determine two latent carrier branches *ω*_*±*_(*k*) of an overdamped electromechanical coupler. The mesoscale ratio

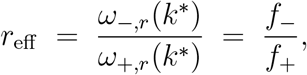

and the slow interference cadence Ω_slow_ = *ω*_+,*r*_(*k*^∗^) − *ω*_−,*r*_(*k*^∗^) are evaluated at the morphological wavenumber *k*^∗^ = *π/𝓁* set by cortical shell thickness *𝓁*.

### 9.2 Candidate systems and parameter acquisition

Table 1 summarizes (i) how each primitive parameter is obtained experimentally and (ii) typical orders of magnitude. These values are indicative placeholders: system-specific values (e.g. MDCK vs. HaCaT vs. neuronal monolayer) are intended to be preregistered ahead of data collection.

**Table 1:** Parameter provenance for computing (Ω, *K*_2_), *r*_eff_, and *f*_slow_ in the overdamped, two-latent-mode framework. Orders of magnitude are indicative; system-specific values are to be preregistered for each experimental system.

| Parameter | Units | Typical order | Acquisition (method / note) |
| --- | --- | --- | --- |
| $D_V$ | $\text{m}^2/\text{s}$ | $10^{-10}$–$10^{-8}$ | Optical electrophysiology; spatial decay of $V$; fit diffusive RC model. |
| $\tau_m$ | s | 0.1–2 | Membrane charging from voltage steps; corrected for dye kinetics. |
| $\kappa$ | N/m | $10^{-4}$–$10^{-2}$ | Effective cortical stiffness from small-amplitude TFM/AFM fits. |
| $\eta$ | N·s/m | $10^{-4}$–$10^{-2}$ | Viscous loss from frequency sweeps in the 1–3 Hz band. |
| $D_\sigma$ | $\text{m}^2/\text{s}$ | $10^{-12}$–$10^{-10}$ | Stress diffusion inferred from relaxation of traction patterns. |
| $\gamma$ | m/V | small | Voltage-to-displacement gain of $u$ per volt (report peak gain near $f_\pm$). If only stress drive is available, report in $\text{N m}^{-2}\text{V}^{-1}$. |
| $\beta_R, \beta_I$ | V per (stress unit) | small | Stress-to-voltage cross-gain; in-phase ($\beta_R$) and quadrature ($\beta_I$) via lock-in / phase-lag measurements. |
| $\ell$ | m | $10^{-5}$–$10^{-3}$ | Effective cortical/shell thickness from confocal optical sectioning. |

### 9.3 Spectral estimation and coherence

Paired time series *V* (*t*) and *σ*(*t*) are acquired simultaneously at sampling rate ≥100 Hz for durations ≥ 600 s in confluent monolayers. After detrending, Welch spectra are computed with 10 s Hanning windows, 50 % overlap, and averaged to estimate *S*_*V*_ (*f*), *S*_*σ*_(*f*), and the cross-spectrum *S*_*V σ*_(*f*). The electromechanical coherence

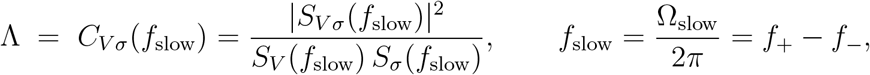

is then evaluated at the slow interference cadence.

For prediction P1, slow-band spectra are explicitly isolated. For each signal *X* ∈ {*V, σ* }, a narrowband analytic signal *X*_slow_(*t*) is constructed by demodulating around *f*_slow_ with a symmetric band-pass (typically ±0.05–0.08 Hz about *f*_slow_, or a multitaper equivalent). 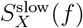 denotes the power spectrum of *X*_slow_(*t*), which is used to assess whether both channels exhibit a coincident slow-band peak. Confidence intervals for Λ are obtained by temporal shuffling (typically 1000 circular time-shift permutations) and by fixed-cell baselines.

### 9.4 Rotation number and stroboscopic regression

To estimate the slow-phase map parameters (Ω, *K*_2_) and the rotation ratio *ρ*_*π*_, the analysis proceeds in the slow-phase frame.

First, analytic signals *Z*_*X*_(*t*) = *X*(*t*)+*i* ℋ [*X*(*t*)] are formed for *V* and *σ*, where ℋ denotes the Hilbert transform. The instantaneous phase difference *ψ*(*t*) = arg *Z*_*V*_ (*t*) − arg *Z*_*σ*_(*t*) tracks the lag between electrical and mechanical sectors.

Neutral moments {*τ*_*n*_} are defined as the recurrent instants where the two sectors align modulo *π*:

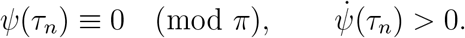

Sampling at these neutral moments enforces the even (mod *π*) Poincaré section described in the main text. Let *ϕ*_+_(*t*) be the instantaneous phase of the + carrier branch (the higher-frequency carrier), and set *θ*_*n*_ = *ϕ*_+_(*τ*_*n*_). The slow-phase increments are Δ*θ*_*n*_ = *θ*_*n*+1_ − *θ*_*n*_.

Regressing Δ*θ*_*n*_ against sin(2*θ*_*n*_) yields estimates of Ω (intercept) and *K*_2_ (slope), and the inferred rotation number *ρ*_*π*_ follows from the extracted Ω via 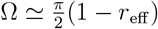.

### 9.5 Statistical model for the coupling *K*_2_

Let Δ*θ*_*n*_ = *θ*_*n*+1_ − *θ*_*n*_, unwrapped to ℝ. The stroboscopic slow-phase map is modeled as

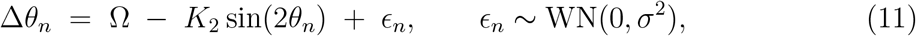

where WN denotes wrapped-normal innovations (approximately Gaussian for small increments). Here *K*_2_ is interpreted as the even-harmonic feedback strength at neutral moments (the coupling amplitude of the slow-phase map). Parameters (Ω, *K*_2_) are estimated by constrained ordinary least squares with *K*_2_ ≥ 0, and Newey–West robust standard errors are reported. Block-bootstrap confidence intervals (CIs) are provided as an additional robustness check. Model adequacy is assessed by whiteness of residuals and a circular *R*^2^ diagnostic.

### 9.6 Biological King plot: common-curvature test (pre-experimental)

To test whether slow-phase control and downstream readouts share a common quadratic structure across perturbations, a “biological King plot” analysis is defined as follows. Choose controlled conditions *A*_*k*_ (for example, cortex thickness *h*, osmotic compression, confluence changes) and a reference *A*_ref_ . For each *A*_*k*_, compute Ω(*A*_*k*_) via neutral-moment stroboscopy (even parity) and extract the slow-band observables

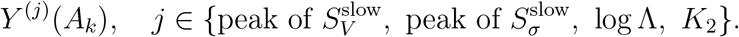

To account for morphology, apply a rescaling 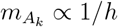 and define

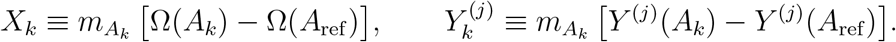

Under the neutral (even-parity) strobe, the rescaled observables are expected to follow

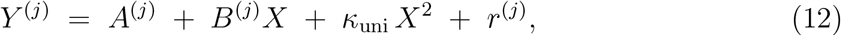

with approximately the same quadratic curvature *κ*_uni_ across all *j*. This “common curvature” is interpreted as evidence that voltage slow-band power, cortical-stress slow-band power, log Λ, and *K*_2_ are co-driven by a single electromechanical interference mode at *f*_slow_. Switching to the full-cycle strobe (odd parity) provides an internal control: it introduces an additional odd component, disrupts the shared quadratic curvature across readouts, and increases the alternation index (App. A).

### 9.7 Experimental candidates and realism

Homeostatic epithelial monolayers (e.g. MDCK, HaCaT) and dense neuronal cultures provide paired *V* (*t*) and *σ*(*t*), controllable *𝓁*, and standard AFM / TFM / optical readouts. These systems allow direct estimation of (*D*_*V*_, *τ*_*m*_, *κ, η, D*_*σ*_, *β*_*R*_, *β*_*I*_, *γ*), construction of the slow-phase map (Ω, *K*_2_), and measurement of Λ. Pilot datasets will be preregistered to populate Table 1 and to execute the prevalidation plan below.

#### Algorithm 1 Cycle-resolved workflow (overdamped, two-latent-mode framework)

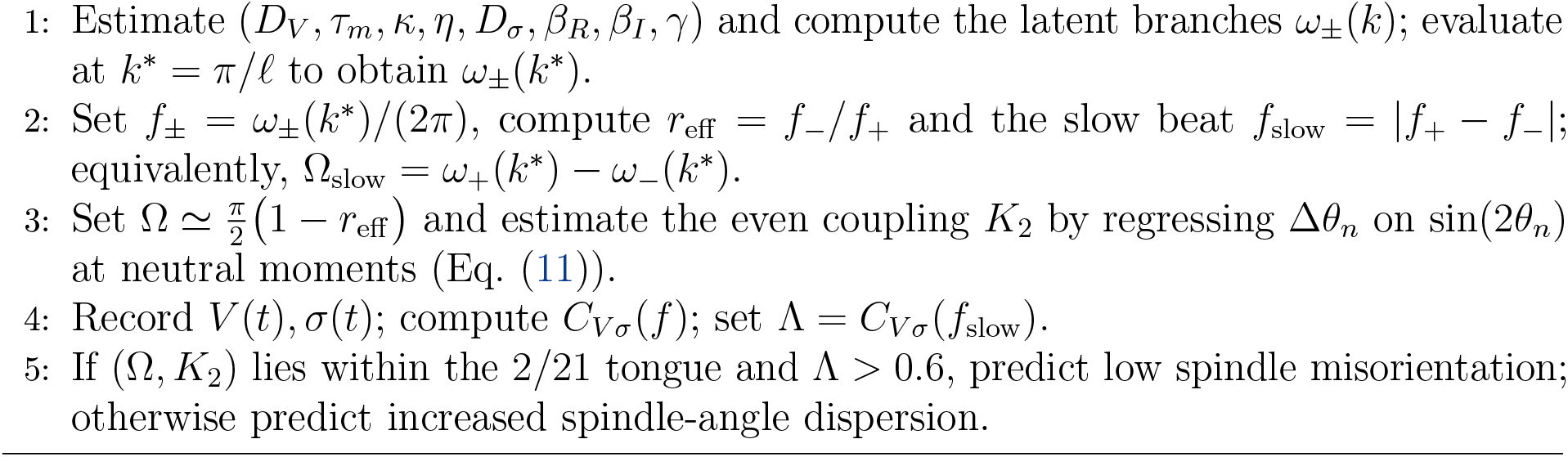

#### Prevalidation plan (addressing P2 robustness)

Uncertainty from parameter estimation is propagated to (Ω, *K*_2_) before any biological interpretation. Profile likelihoods and Monte Carlo sampling are used to propagate uncertainty in (*γ, a, b, k*^∗^), where 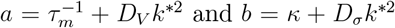. The resulting sampling distribution of (Ω, *K*_2_) is summarized by two preregistered conditions: *(C1) P*((Ω, *K*_2_) ∈ tongue(2/21)) ≥ 0.3 (95% CI), and *(C2)* the safety margin to the nearest lower-order tongue (1/2, 1/3, 2/5) exceeds 3*σ*_Ω_, where *σ*_Ω_ is the robust SE for Ω from neutral-moment regression. In parallel, Λ is estimated with temporal shuffling (*N* ≥ 1000 permutations) and a fixed-cell baseline; Λ *>* 0.6 (95% CI) is preregistered as the operational “high-locking” threshold.

### 9.8 Preregistered reanalysis on public datasets (Level-1 verification)

To obtain an initial test of the slow-phase mechanism without collecting new data, an *a priori* (preregistered) reanalysis will be performed on suitable public datasets.

#### Dataset types and inclusion criteria

Candidate datasets include: (i) confluent epithelial monolayers (MDCK/HaCaT/Caco-2), (ii) dense neuronal cultures as a positive dynamical control, and (iii) organoid monolayers or early spheroids. Inclusion criteria:

- **Sampling and duration:** frame rate ≥ 80 Hz (to resolve carrier peaks in the 1 −− 3 Hz band) and total duration ≥ 300 s.
- **Observables:** (A) paired electrical/mechanical channels (preferred): a voltage reporter (GEVI or fast dye) and traction/stress (TFM/AFM or micropillar arrays); or (B) a voltage channel plus a mechanical proxy (bright-field kymographs, opticalflow divergence, speckle-based strain) that can be collapsed to a scalar *σ*(*t*); or (C) a mechanical channel plus a slower electrical proxy with documented SNR.
- **Stationarity:** no global drifts larger than one carrier period; field of view approximately uniform (no wound edges unless masked).

#### Reconstruction of observables

From raw movies, *V* (*t*) is taken as the field-averaged GEVI/dye signal (after photobleach correction and detrending). *σ*(*t*) is obtained as either the spatial mean traction (TFM) / principal component of the traction field, or a calibrated mechanical proxy (e.g. optical-flow divergence). Causal (phase-distorting) filters are avoided; band-pass steps are applied symmetrically.

#### Analysis pipeline (fixed, preregistered)

1. **Carrier identification:** estimate *S*_*V*_ (*f*) and *S*_*σ*_(*f*) (Welch; 10 s windows; 50 % overlap). Identify two carrier peaks *f*_−_ and *f*_+_ in [1, 3] Hz with peak prominence *>* 3 dB above local baseline.
2. **Neutral-moment stroboscopy:** form analytic signals *Z*_*X*_ and the phase lag *ψ*(*t*) = arg *Z*_*V*_ − arg *Z*_*σ*_. Define neutral moments 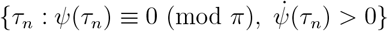.
3. **Even-map regression:** sample *θ*_*n*_ = *ϕ*_+_(*τ*_*n*_) (phase of the + carrier) and regress Δ*θ*_*n*_ = *θ*_*n*+1_ − *θ*_*n*_ on sin(2*θ*_*n*_) to estimate 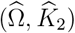 with Newey–West standard errors and block-bootstrap CIs (Eq. (11)).
4. **Slow-band coherence:** demodulate around *f*_slow_ = *f*_+_ − *f*_−_ and compute Λ = *C*_*V σ*_(*f*_slow_), using 1000-fold temporal shuffling for confidence intervals.
5. **Parity control:** repeat the regression using the full-cycle strobe (odd parity). Report the alternation index and loss of common quadratic curvature as in App. A.

#### Level-1 pass/fail criteria for homeostatic monolayers consistent with the dual– field locking picture

- (E1) two resolved carrier peaks *f*_*±*_ with *f*_slow_ = *f*_+_ − *f*_−_ above noise;
- (E2) Λ *>* 0.6 at *f*_slow_ (95% CI) under neutral strobing;
- (E3) 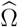 lies within ±0.1 rad of Ω_2/21_ = *π* · 2/21, and switching parity (even→odd) disrupts common curvature / increases alternation index.

*Because most public datasets lack independent estimates of* (*D*_*V*_, *τ*_*m*_, *κ, η*, …), *criterion (E3) is interpreted as a reduced form of P2:* 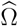 *is obtained from r*_eff_ = *f*_−_*/f*_+_ *rather than from full parameter provenance*.

- (E4) 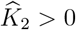 with significant slope (two-tailed *p <* 0.05).

#### Contra-evidence in the same datasets

- (C1) no resolvable *f*_slow_ or Λ *<* 0.2;
- (C2) 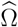 displaced *>* 0.2 rad from Ω_2/21_ with tight CI, or non-significant *K*_2_ regression;
- (C3) parity switching fails to alter alternation / common-curvature metrics.

#### Candidate repositories and data modalities

Suitable material is expected in (i) traction-microscopy movies (micropillar arrays, TFM), (ii) AFM-based viscoelastic sweeps on confluent epithelia, and (iii) GEVI/dye voltage recordings of confluent monolayers. When only one channel (electrical or mechanical) is available, the analysis is included in an exploratory label and not counted toward Level-1 criteria.

#### Negative and adversarial controls

Fixed-cell preparations (no dynamics), sparse low-density cultures lacking collective mechanics, and small-molecule decoupling of the cortex (e.g. actomyosin inhibition) are expected to suppress *K*_2_ and lower Λ. Off-strobe windows (misaligned stroboscopic sampling) serve as additional negative controls.

#### Reporting

For each dataset a one-page report will be generated, including QC plots, *S*_*V*_ (*f*) and *S*_*σ*_(*f*), slow-band spectra, (Ω, *K*_2_, Λ) with confidence intervals, parity metrics, and pass/fail flags for (E1)–(E4). This reporting structure is preregistered to avoid hindsight tuning.

## 10 In-silico validation: continuum sandbox and cell-based models

### Level A: continuum sandbox (overdamped PDE electromechanical coupler)

Eqs. (16)–(18) are integrated on a 2D periodic domain of linear size *L* using pseudo-spectral or finite-difference time stepping.

Parameters (*D*_*V*_, *τ*_*m*_, *κ, η, D*_*σ*_, *β*_*R*_, *β*_*I*_, *γ*) are drawn from physiologically plausible priors; no ad hoc integer constraints are imposed. Observables are the spatial means *V* (*t*) = ⟨*V* (**x**, *t*)⟩ and *σ*(*t*) = ⟨*σ*(**x**, *t*)⟩, or equivalently the Fourier mode at *k*^∗^ = *π/𝓁* (the mesoscale set by cortical thickness *𝓁*). The sandbox is required to reproduce: (i) two carrier peaks *f*_*±*_ in the fast band generated by an *overdamped* dual-mode coupler, (ii) a slow interference band at *f*_slow_ = |*f*_+_ − *f*_−_|, (iii) neutral-moment stroboscopy (even parity) that yields a well-defined slow-phase map and estimates 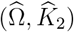, and (iv) Monte Carlo sampling over priors that places (Ω,*K*_2_) near the 2/21 Arnold tongue while avoiding lower-order tongues for a non-negligible fraction of draws. Weak external drives (electrical or mechanical) at *f*_slow_ are then applied to test phase-targeted entrainment (P3) by checking whether, relative to matched off-band controls, an on-band drive produces ΔΛ *>* 0 together with a drop in neutral-map angular variance Var_map_.

#### Level B: cell-based model (vertex / Cellular Potts / agent-based)

A confluent epithelial sheet is simulated with explicit cells and a diffusive bioelectric layer. Each cell *i* carries a transmembrane potential *V*_*i*_ with gap-junctional coupling; the mechanical sheet is represented as an *overdamped* (first-order) vertex / Cellular Potts / agent-based layer whose active cortical tension is modulated by *V* . Readouts are processed through exactly the same analysis pipeline as in experiments: estimation of *S*_*V*_ (*f*) and *S*_*σ*_(*f*), detection of the two fast-band carriers *f*_*±*_ and the slow beat *f*_slow_ = |*f*_+_ − *f*_−_|, neutral-moment stroboscopy (even parity) to obtain (Ω, *K*_2_), and slow-band electromechanical coherence Λ at *f*_slow_.

#### Phase-targeted entrainment tests and falsification

To assess P3, a weak periodic drive is applied at *f*_slow_: (i) an electrical perturbation *V* ← *V* + *A*_*V*_ sin(2*πf*_slow_*t*) (globally or within a subdomain), and/or (ii) a mechanical/traction drive *A*_*σ*_ sin(2*πf*_slow_*t*). Controls use off-resonant drives at *f*_slow_(1 ± 0.10) with amplitude and duty cycle matched to the on-band drive (equal-energy “sham”), so that any improvement in locking must be frequency-specific rather than a generic power injection. The primary endpoints are (a) ΔΛ *>* 0 specifically under on-band drive, and (b) a selective reduction in Var_map_ (interpreted as reduced spindle-angle dispersion) under on-band drive but not under the off-band controls. Secondary endpoints include an increase in the even-harmonic coupling estimate *K*_2_. Falsification criteria are: absence of a resolvable *f*_slow_ band, persistently low coherence (Λ *<* 0.2), non-significant even-map regression (ill-defined 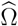 or 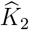), or lack of on-band specificity relative to off-resonant controls. App. G implements and prevalidates this logic in silico.

### 10.1 Reference implementation (concept and usage)

#### Purpose

Listing M.1 is a minimal, CPU-light reference implementation of the dual-field sandbox. It operationalizes P1 and P2 (shared slow-band cadence, placement of (Ω, *K*_2_) in the 2/21 tongue) and reproduces the basic Case D “recovery by on-band entrainment” scenario.

App. G adds a dedicated script (Listing M.4) that implements the full preregistered P3 assay, including off-band controls and Var_map_.

#### What it does

In each run the code:

1. generates two *latent carriers* in the fast band with small noise and slow drift (either by integrating the full overdamped PDE coupler or by using a lightweight two-carrier signal generator with the same structure);
2. computes Welch spectra *S*_*V*_ (*f*), *S*_*σ*_(*f*) and the magnitude-squared coherence Λ(*f*);
3. identifies the carrier pair (*f*_−_, *f*_+_) and the slow interference cadence *f*_slow_ =| *f*_+_ − *f*_−_| ;
4. performs neutral-moment stroboscopy (even parity) and regresses the even slow-phase map to estimate (Ω, *K*_2_);
5. optionally applies a weak, narrow-band drive during a defined time window (phase-targeted entrainment test) and reports ΔΛ;
6. saves diagnostic figures (space–time *V* (*x, t*), spectra with *f*_*±*_ and *f*_slow_, and timeresolved Λ(*t*)) and prints summary diagnostics.

#### How it is used here

For the illustrative regimes in Cases A–D, the same pipeline is run on a lightweight signal generator (two carriers, controlled noise, slow drift) to make the logic transparent. The parameter tweaks defining each regime are summarized in Table 2 and listed explicitly in App. M. Listing M.1 is the pseudo-spectral reference implementation of this workflow.

**Table 2:**
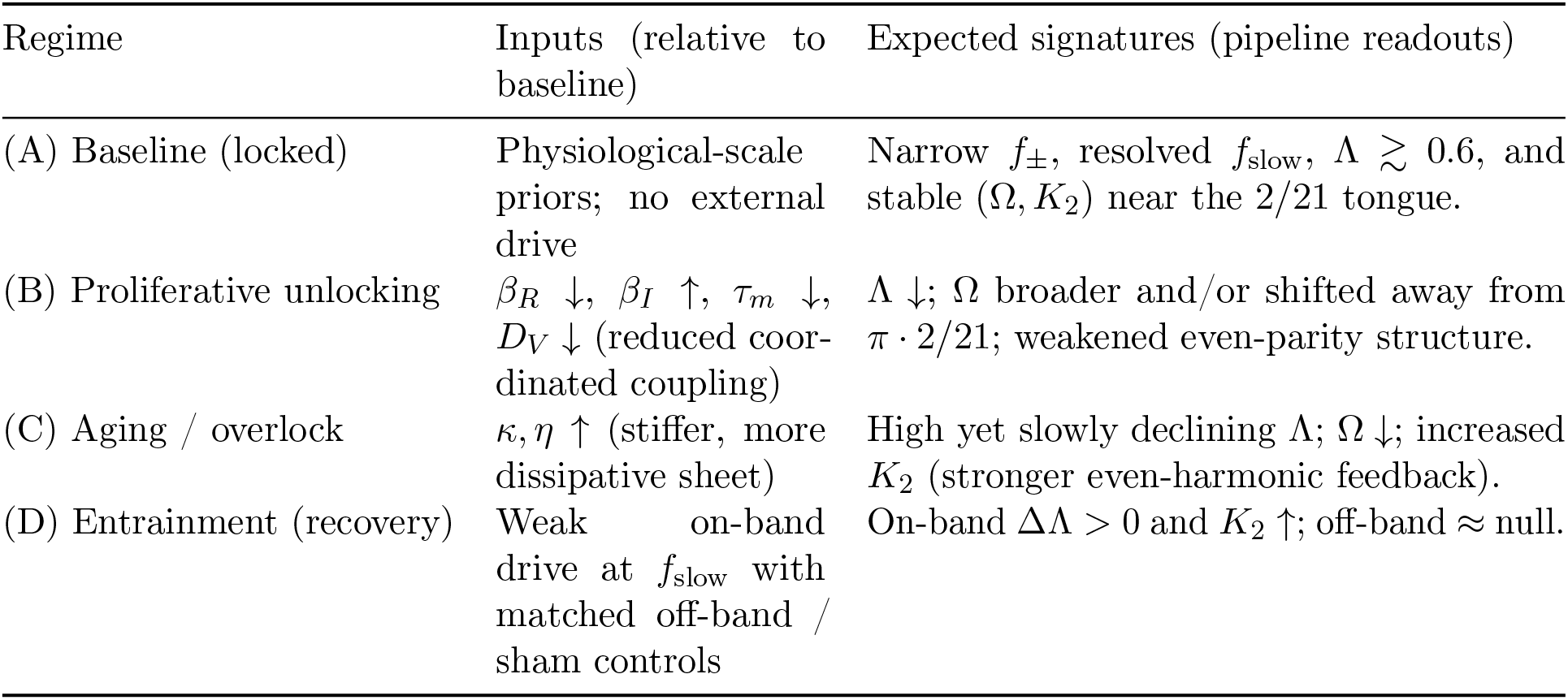
Checklist of in-silico regimes (Cases A–D). “Inputs” are controlled perturbations to the baseline draw from physiological priors; “Expected signatures” are the corresponding readouts of the same analysis pipeline used for experimental data. Full code, including parameter dictionaries and drive protocol, is provided in App. M.

| Regime | Inputs (relative to baseline) | Expected signatures (pipeline readouts) |
| --- | --- | --- |
| (A) Baseline (locked) | Physiological-scale priors; no external drive | Narrow $f_{\pm}$, resolved $f_{\text{slow}}$, $\Lambda \gtrsim 0.6$, and stable $(\Omega, K_2)$ near the 2/21 tongue. |
| (B) Proliferative unlocking | $\beta_R \downarrow$, $\beta_I \uparrow$, $\tau_m \downarrow$, $D_V \downarrow$ (reduced coordinated coupling) | $\Lambda \downarrow$; $\Omega$ broader and/or shifted away from $\pi \cdot 2/21$; weakened even-parity structure. |
| (C) Aging / overlock | $\kappa, \eta \uparrow$ (stiffer, more dissipative sheet) | High yet slowly declining $\Lambda$; $\Omega \downarrow$; increased $K_2$ (stronger even-harmonic feedback). |
| (D) Entrainment (recovery) | Weak on-band drive at $f_{\text{slow}}$ with matched off-band / sham controls | On-band $\Delta\Lambda > 0$ and $K_2 \uparrow$; off-band ≈ null. |

#### How to run and modify

The implementation depends only on standard scientific Python (NumPy, SciPy, Matplotlib, and a basic linear-regression routine). Running the main script produces: (i) a figure with three panels (space–time signal, spectra with *f*_*±*_ and *f*_slow_, and Λ(*t*)); (ii) numerical estimates of *f*_−_, *f*_+_, *f*_slow_ and Λ(*f*_slow_); (iii) 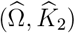 and, if an on-band drive was applied, ΔΛ comparing driven vs. undriven windows. Stable, high Λ corresponds to locking (Case A); low, bursty Λ corresponds to unlocking (Case B/B’); a gently declining Λ(*t*) corresponds to overlock with drift (Case C); and an on-band increase in Λ under weak drive corresponds to partial recovery by phase-targeted entrainment (Case D).

#### Scope

The sandbox is designed to validate the dynamical mechanism captured by P1 and P2, and to generate preregisterable targets for P3. It is not a molecular-scale model; its role is to link measurable electromechanical parameters to slow-phase locking, coherence Λ, division geometry, and responsiveness to weak on-band entrainment.

The explicit computational prevalidation of P3, including frequency-specific rescue of Λ and suppression of Var_map_ under on-band (but not off-band) drive, is provided in App. G; see also Figs. G.2 and G.1.

## 11 Reproducible in-silico regimes: implementation and outputs

### Case A – Baseline (locked)

#### Setup

Carrier frequencies in the ~1–2 Hz band (e.g. *f*_−_ ≈1.6 Hz, *f*_+_ ≈2.0 Hz). Both *V* (*t*) and *σ*(*t*) contain these carriers with small noise and minimal detuning. No external drive is applied.

#### Expected outcome

Narrow peaks at *f*_−_ and *f*_+_; a slow interference band at *f*_slow_ = |*f*_+_ − *f*_−_| in the sub-Hz range; Λ ≳ 0.6 at *f*_slow_; and stroboscopic estimates 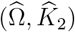 lying near the center of the 2/21 tongue (i.e. close to Ω_2/21_ = *π* · 2/21 with nonzero *K*_2_).

#### Diagnostics (illustrative numbers)

- Carrier estimate: *f*_−_ ∈ [1.55, 1.70] Hz, *f*_+_ ∈ [1.95, 2.05] Hz; *f*_slow_ ≈ 0.37–0.40 Hz.
- Λ(*f*_slow_) ≥ 0.60; median Λ(*t*) typically 0.5–0.8 with only small window-induced ripples.
- Neutral-moment stroboscopy: thousands of strobes in ~ 600 s; even-map regression with circular 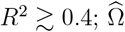 within ±0.2 rad of *π* · (2/21).

**Figure 11.1:**
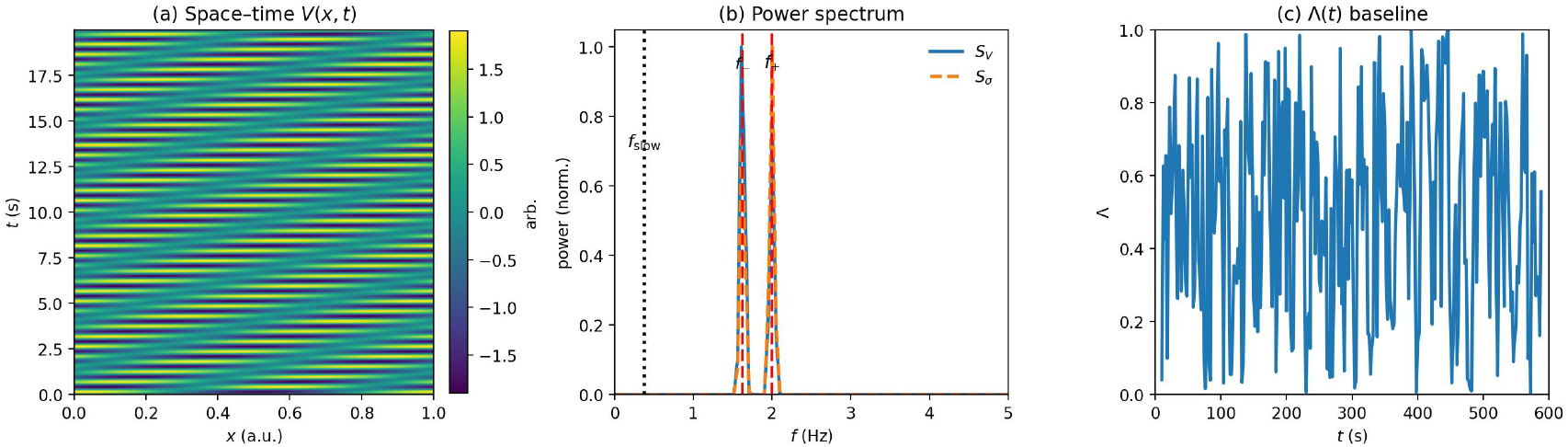
Case A (baseline/locked). The slow band at *f*_slow_ is present and Λ(*t*) remains high; 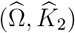 fall near the 2/21 tongue. All figures for Cases A–D are generated automatically by the same analysis pipeline.

#### Quick internal checks

- **Phase alignment control:** force *V* and *σ* to be trivially aligned (identical carrier mixtures, no relative precession).
Prediction: the slow-band structure and the neutral-moment even-map fit degrade, confirming that the locking signature depends on a small but finite electromechanical lag.
- **Mild detuning robustness:** shift *f*_*±*_ by ±0.02 Hz. Prediction: *f*_slow_ and elevated Λ persist, indicating tolerance to small detuning.

**Figure 11.2:**
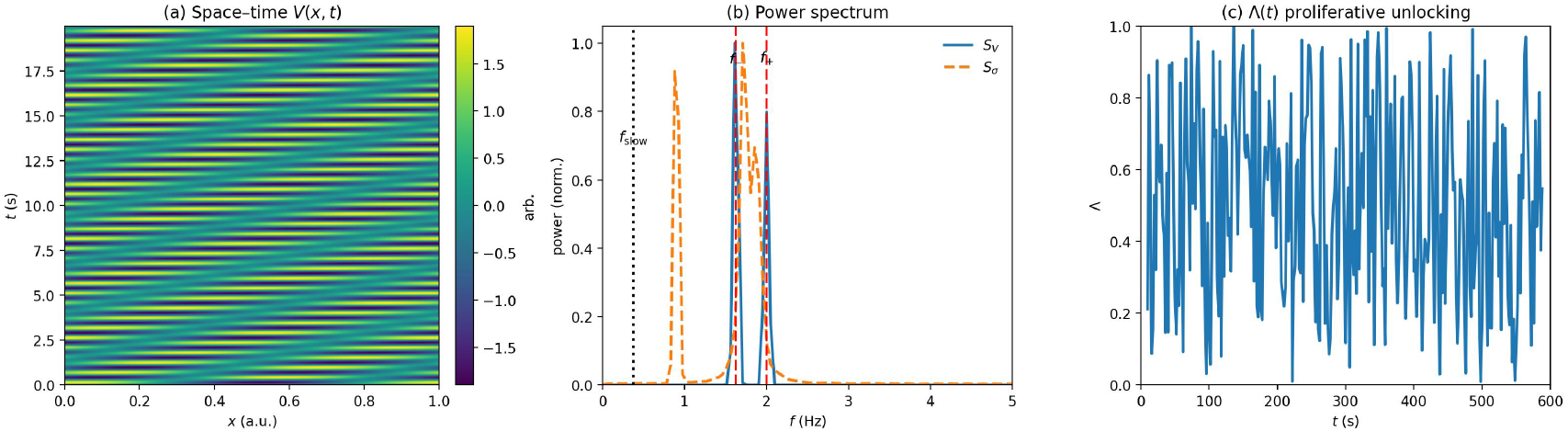
Case B (proliferative unlocking, moderate). Left to right: (a) toy space–time *V* (*x, t*); (b) power spectra with broadened *S*_*σ*_ (orange); (c) Λ(*t*) showing a lower, burstier baseline compared with Case A, including transient drops.

### Case B – Proliferative unlocking (moderate)

We emulate proliferative unlocking by selectively degrading coordinated electromechanical coupling in the mechanical surrogate *σ*(*t*): we weaken the in-phase component, introduce slow drift in the *σ* phase, add modest detuning only in *σ*, and increase its noise.

#### Implementation (summary; details in App. M)

- Keep *V* (*t*) carriers near the baseline pair (~1.6 Hz, ~2.0 Hz).
- Reduce the in-phase carrier weights in *σ*(*t*).
- Impose a slow phase drift and mild frequency detuning in *σ* only.
- Add moderate noise to *σ*.
- Track time-resolved coherence Λ(*t*) at *f*_slow_ using a fixed window (e.g. 20 s) and a ±0.05 Hz band.

#### Expected outcome

*S*_*σ*_(*f*) broadens around *f*_*±*_ relative to *S*_*V*_ (*f*); *f*_slow_ remains detectable; and Λ(*t*) shows a lower and more bursty baseline than in Case A, typically ~ 0.4–0.6 with transient drops below 0.3. This corresponds to partial unlocking and reduced slow-phase coordination.

### Case B’ – Strong proliferative unlocking (sensitivity)

To stress-test the signature of unlocking, we push decoupling harder by further weakening in-phase alignment, adding stronger detuning, and injecting an off-band mechanical component.

#### Implementation (summary)

- Strongly suppress the in-phase carrier contribution in *σ*.
- Impose larger carrier-frequency detuning in *σ*.
- Add a weak off-band tone (e.g. ~ 0.9 Hz) to *σ*.
- Increase *σ*-channel noise.
- Recompute Λ(*t*) around *f*_slow_ with a longer window (e.g. 30 s) and a narrower band (±0.01 Hz).

*Result. S*_*σ*_(*f*) broadens substantially and develops an off-band shoulder around the injected ~0.9 Hz component.

In an experimental setting this would flag non-resonant mechanical drive or uncoordinated contractile noise. Λ(*t*) shows a much lower baseline and bursty excursions, with median values often in the ~ 0.2–0.4 range.

This captures the expected trend **proliferative unlocking** ⇒ **coherence loss** (P1).

**Figure 11.3:**
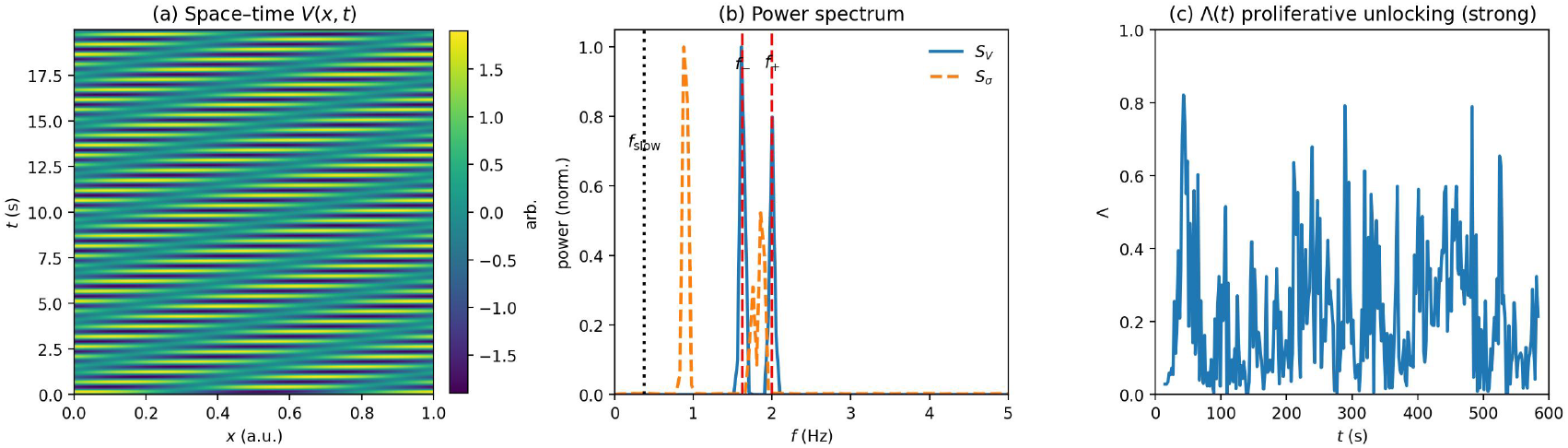
Case B’ (strong unlocking). Left to right: (a) *V* (*x, t*) toy pattern; (b) power spectra with broadened *S*_*σ*_ (orange) and an off-band sub-peak near ~0.9 Hz; (c) time-resolved coherence Λ(*t*) with a reduced baseline and bursty profile.

### Case C – Aging / overlock

We emulate aging / overlock by increasing effective stiffness and viscosity: *κ* ↑, *η* ↑.

This shifts carriers slightly lower in frequency, strengthens even-harmonic feedback, and reduces adaptability while maintaining high coherence.

#### Implementation (summary)

- Shift both carrier frequencies slightly lower (e.g. *f*_−_ ≈1.5 Hz, *f*_+_ ≈1.85 Hz).
- Impose a slow amplitude decay in *σ*(*t*) and a small, quasi-static phase bias between *V* and *σ*.
- Keep noise relatively low.
- Track Λ(*t*) with a sliding window and plot a smoothed trend.

#### Expected outcome

A lower carrier pair in the 1–2 Hz band; Λ(*t*) remains high but displays a gentle downward drift over time (reduced adaptability); and the neutral-moment regression returns a modestly increased *K*_2_, consistent with stronger even-harmonic feedback.

**Figure 11.4:**
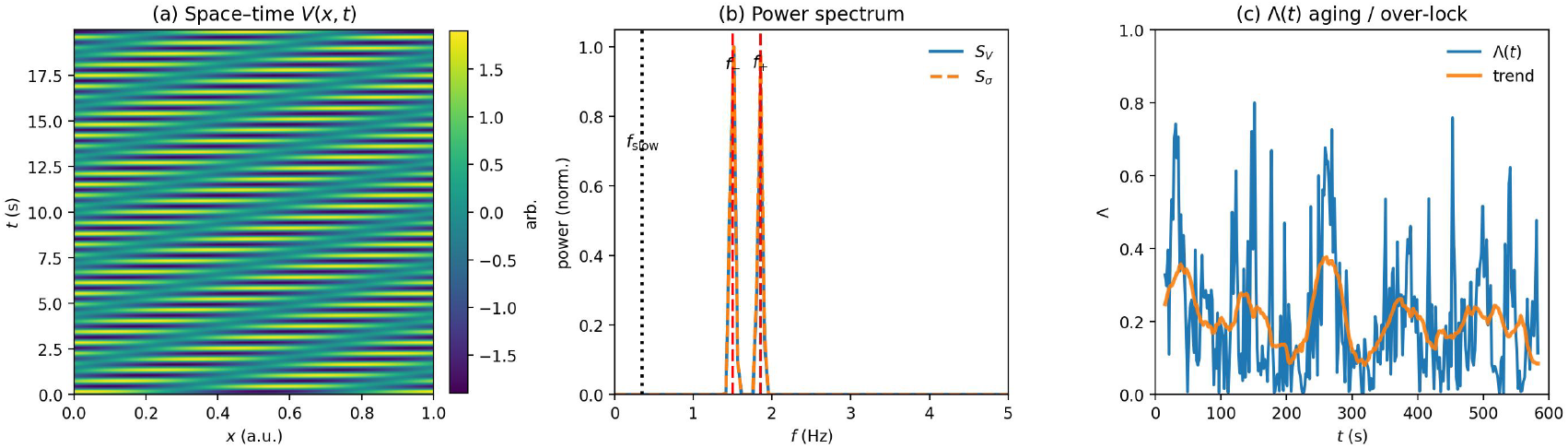
Case C (aging / overlock). Left to right: (a) *V* (*x, t*) toy pattern; (b) power spectra showing the lowered carrier pair; (c) time-resolved coherence Λ(*t*) with a slowly declining trend over the run.

### Case D – Recovery by on-band entrainment

Starting from an unlocking configuration (Case B), we apply a weak global drive at the slow cadence *f*_slow_ during a limited time window to test phase-targeted entrainment (P3). Case D demonstrates qualitatively that an on-band drive can transiently restore coordination between *V* and *σ*.

App. G formalizes this by adding symmetric off-band controls (±10% detuning), quantifying ΔΛ and Var_map_, and spelling out preregistered falsification criteria.

#### Implementation (summary)

- Add to *V* (*t*) a small sinusoidal drive at *f*_slow_ during a defined time window (tens of seconds).
- Keep the drive amplitude low enough that the carrier peaks *f*_*±*_ shift by no more than ~ 1–3%.
- Repeat with an *off-band* control drive at frequency *f*_slow_ · (1 + 0.10), with amplitude and duty cycle matched to the on-band drive.
- Define 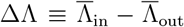, where “in” refers to the drive window and “out” to the surrounding baseline.

#### Result

In Case D, the on-band drive produces an increase in Λ(*t*) confined to the driven window, yielding ΔΛ *>* 0, while an off-band control with matched amplitude and duty cycle produces ΔΛ ≈ 0. Carrier frequencies *f*_*±*_ remain essentially unchanged, indicating that the drive is weak and spectrally specific rather than a gross perturbation.

App. G extends this test and shows that, in a proliferative-like unlocked state with high Var_map_, a weak on-band drive at *f*_slow_ simultaneously increases Λ and reduces Var_map_, whereas off-band drives of identical amplitude fail to reproduce both effects together. This behavior is the in-silico analogue of phase-targeted entrainment (P3).

**Figure 11.5:**
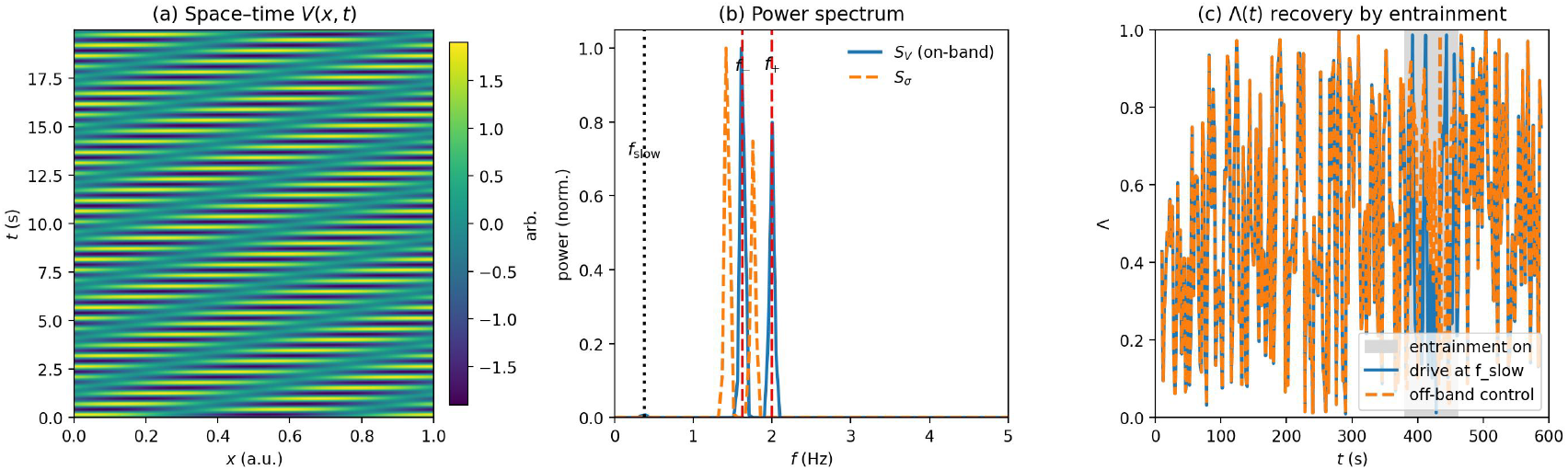
Case D (recovery by on-band entrainment). Left to right: (a) *V* (*x, t*) toy pattern; (b) power spectra (carrier peaks unchanged, consistent with a weak drive); (c) time-resolved coherence Λ(*t*): the on-band drive (solid) increases Λ inside the shaded window, whereas the off-band control (dashed) shows no effect despite matched amplitude and duty cycle.

#### Scope and extensions

These in-silico regimes are designed to validate the *dynamic* mechanism summarized by P1–P3: slow-phase locking with high Λ, proliferative unlocking with reduced Λ and bursty coherence, aging-like overlock with rigid high Λ but reduced adaptability, and recovery by weak on-band entrainment with frequency-specific rescue of Λ and suppression of Var_map_. In baseline draws from physiological-scale priors, Λ(*f*_slow_) *>* 0.6 occurs for a non-negligible fraction; on-band drives produce selective improvements that off-band drives (detuned by ~10%) do not.

Extension to 2D/3D sheets introduces anisotropic carriers and potentially multiple slow interference modes. Neutral-moment locking remains applicable provided local homogeneity at the mesoscale *k*^∗^ = *π/𝓁*. Weak broadband noise can assist coherence (stochastic resonance) but amplitudes must remain below thresholds that shift *f*_*±*_ or cause damage in vitro.

All implementation details (parameter dictionaries for Cases A–D, drive schedules, plotting commands) are consolidated in App. M, and the full preregistered P3 prevalidation (including off-band controls and Var_map_) is given in App. G.

## 12 Results: analytical and numerical evidence

Before turning to width bounds and numerical scans, we summarize how the reduced even circle map arises from the overdamped electromechanical model. Starting from the continuum coupler (Eqs. (16)–(18)), we project onto two latent carriers with complex eigenfrequencies Ω_*±*_ (Eqs. (1)–(4)), whose real parts define the fast-band pair *f*_*±*_ and hence the slow beat *f*_slow_ = |Ω_+_ − Ω_−_| /(2*π*). In the time domain, we then identify neutral crossings {*τ*_*n*_} where the electrical and mechanical phases momentarily align modulo *π* with positive slope (App. E). Sampling the fast phase at these times, *θ*_*n*_ = *ϕ*_+_(*τ*_*n*_), yields the neutral-moment stroboscopic map

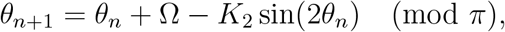

with forcing Ω fixed by the carrier ratio *r*_eff_ = *f*_−_*/f*_+_ and even-harmonic coupling *K*_2_ estimated from regression at neutral moments. The analytical structure of this even (mod *π*) map, and its numerical Arnold tongue diagram in the (Ω, *K*_2_) plane, are developed in App. J.

### 12.1 Analytical width bounds

To characterize slow-phase locking we consider the even circle map

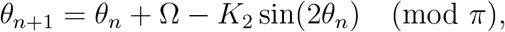

sampled at the neutral moment. For *K*_2_ ≪ 1, the Arnold tongue of a rational lock *p/q* has a width in Ω that scales as

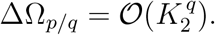

Low-order ratios (small *q*, such as 1/2, 1/3, 2/5) therefore occupy parametrically broader regions of the (Ω, *K*_2_) plane, whereas higher-order ratios (large *q*) carve out much narrower slivers. This scaling suffices to rank low versus high order and to motivate P2: within physiological variability of (Ω, *K*_2_), it should be feasible to avoid persistent capture by *q* ≤ 6 tongues while still intersecting the narrow 2/21 tongue, centred at Ω_2/21_ = *π* · 2/21≃ 0.30 rad.

Importantly, this construction means that the 2/21 plateau is not chosen by hand. Once the two fast carriers *f*_*±*_ have been measured, the effective forcing Ω is fixed (up to uncertainty) by their ratio 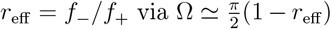. Together with an independent estimate of the even coupling *K*_2_ from neutral-moment regression, this defines a narrow physiological strip in the (Ω, *K*_2_) plane. Because low-order tongues *p/q* with *q* ≤ 6 are parametrically wide whereas higher-order tongues are much narrower, requiring that this strip avoid the broad low-order plateaus while still permitting slow-phase capture leaves very little freedom: within the measured range of (Ω, *K*_2_) the first accessible high-order plateau of the even map is precisely the 2/21 tongue. In this sense the integer ratio 2/21 is not tuned as an extra parameter; it emerges from the carrier frequencies and the tongue-width hierarchy encoded in the even map.

### 12.2 Sensitivity and low-order exclusion for P2

In the overdamped dual-field picture, the slow forcing follows the carrier-frequency ratio:

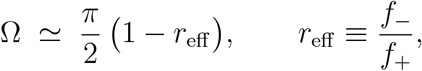

so small changes obey

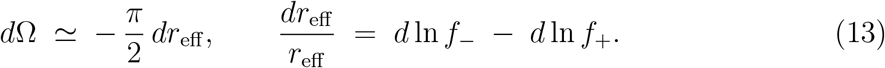

For weak electromechanical coupling the carrier split is small. Writing 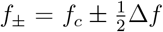 with 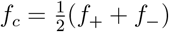a local mean,

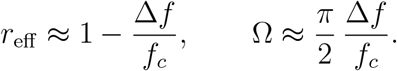

At a coarse-grained level the split scales like

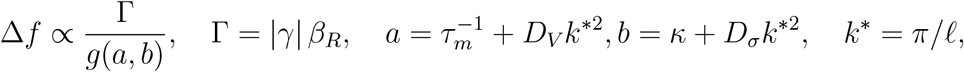

where *g*(*a, b*) *>* 0 lump the electrical and mechanical dissipative blocks. The relative sensitivity of Ω can then be organized as

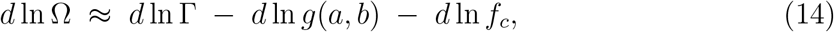

which depends only on coupling strength and dissipation, without invoking areal inertia. Operationally, we (i) estimate *σ*_Ω_ from neutral-moment regression using Newey-West robust errors, (ii) compute the distance in Ω to the nearest low-order centre {1/2, 1/3, 2/5}, and (iii) require simultaneously that the safety margin ΔΩ_low_ ≥ 3 *σ*_Ω_ and that Monte Carlo over physiological priors places (Ω, *K*_2_) inside the 2/21 tongue with probability at least 0.3.

(Ω, *K*_2_) should sit in or near the 2/21 strip at a distance from broad low-order tongues that is large compared with its own uncertainty. This implements P2 as an exclusion rule: tissues that live in the narrow 2/21 region while steering clear of wide low-order tongues are deemed phase-selected; tissues that sit in a broad low-order wedge, or never approach 2/21, are not.

Numerically, Monte Carlo sampling over physiological priors for the parameter vector

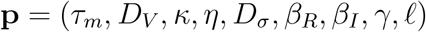

populates a compact band in the (Ω, *K*_2_) plane that overlaps the 2/21 tongue while remaining separated from the centres of the *q* ≤ 6 tongues by several *σ*_Ω_ (App. J). This acts as a coarse-grained phase diagram: typical parameter draws visit the 2/21 strip but do not realize persistent capture in low-order wedges, reinforcing that the 2/21 plateau is a robust outcome of physiological parameter variability rather than a fine-tuned choice.

**Figure 12.1:**
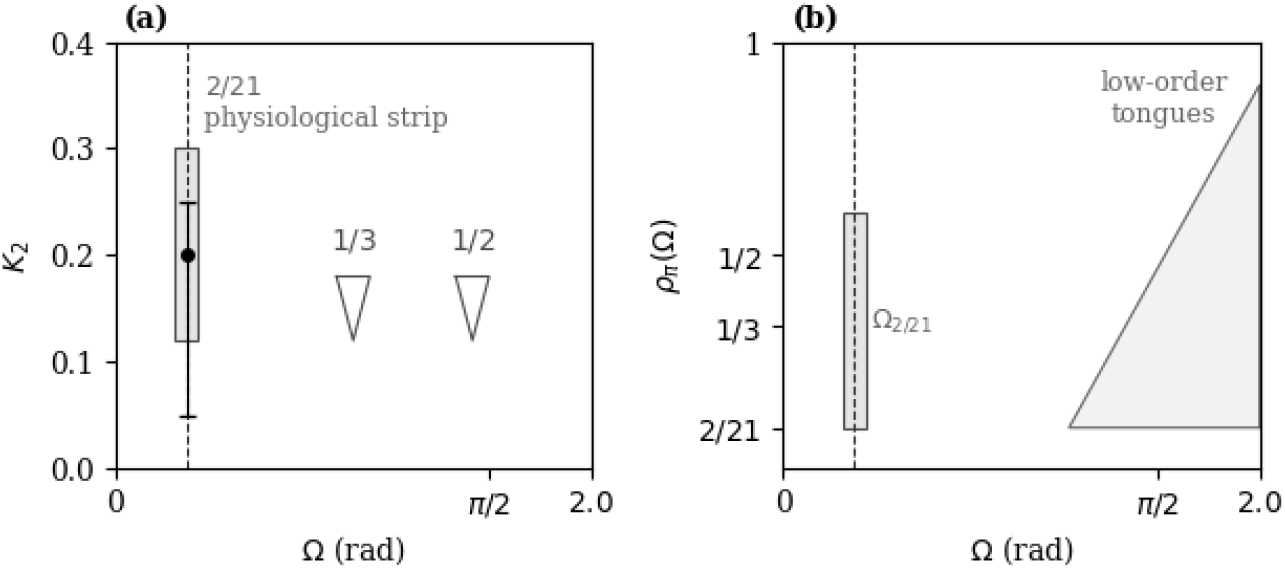
Electromechanical locking landscape in the even (mod *π*) convention. (a) Schematic (Ω, *K*_2_) plane for the even circle map *θ*_*n*+1_ = *θ*_*n*_ + Ω − *K*_2_ sin 2*θ*_*n*_ (mod *π*) sampled at the neutral moment. The shaded vertical band marks the strip around Ω_2/21_ = *π* · 2/21≃ 0.30 rad, interpreted here as the high-order 2/21 tongue. Farther to the right, broad low-order tongues (e.g. 1/3, 1/2) open near Ω ≃ *π/*2 and occupy a larger region. (b) Stylized devil’s staircase view of the locked ratio *ρ*_*π*_(Ω) versus Ω in the same even convention, highlighting the narrow 2/21 plateau and broad low-order plateaus.

### 12.3 Phase-locking geometry in the even convention

Figure 12.Figure 1 summarizes the slow-phase geometry. In panel (a) the tissue is placed on the (Ω, *K*_2_) plane of the even map under neutral-moment stroboscopy. Ω is the effective slow forcing set by the beat between the two latent carriers; *K*_2_ is the even-harmonic feedback strength estimated from data. The vertical band around Ω_2/21_ represents the 2/21 tongue; farther to the right schematic wedges denote broad low-order tongues. P2 asks whether homeostatic epithelia systematically fall inside the narrow 2/21 strip while avoiding the wide low-order regions.

Panel (b) recasts the same structure as a staircase *ρ*_*π*_(Ω), with *ρ*_*π*_ the effective winding number (locked ratio between slow-phase advance and the neutral strobe). The 2/21 plateau is narrow in Ω compared with generic low-order plateaus such as 1/3 or 1/2. In this picture, proliferative unlocking corresponds to drift away from the 2/21 plateau, and aging/overlock corresponds to remaining trapped in a broad low-order wedge with reduced adaptability.

While Fig. 12.Figure 1 is schematic, the same structure arises from direct numerical analysis of the even (mod *π*) phase map. The existence of a narrow 2/21 locking tongue is not imposed by construction but emerges from the reduced equations used for the electromechanical analysis. To verify this numerically, we scanned the (Ω, *K*_2_) plane and estimated the rotation number *ρ* = *f*_1_*/f*_2_ from long integrations of the coupled-phase model. A zoom around the 2/21 tongue and a transverse cut at Ω = 2/21 are shown in App. J, Figs. J.1 and J.2. These calculations confirm that the parameter range used for the electromechanical map lies in the transition region between the 2/21 plateau and the onset of near-1:1 locking, rather than deep inside a low-order tongue.

### 12.4 Slow-band spectra and coherence (P1)

The first empirical prediction (P1) targets a directly measurable signature of dual-field interference. In frequency space, the model predicts an electromechanical slow cadence *f*_slow_ that is shared by the bioelectric sector *V* (**x**, *t*) and the mechanical sector *σ*(**x**, *t*).

**Figure 12.2:**
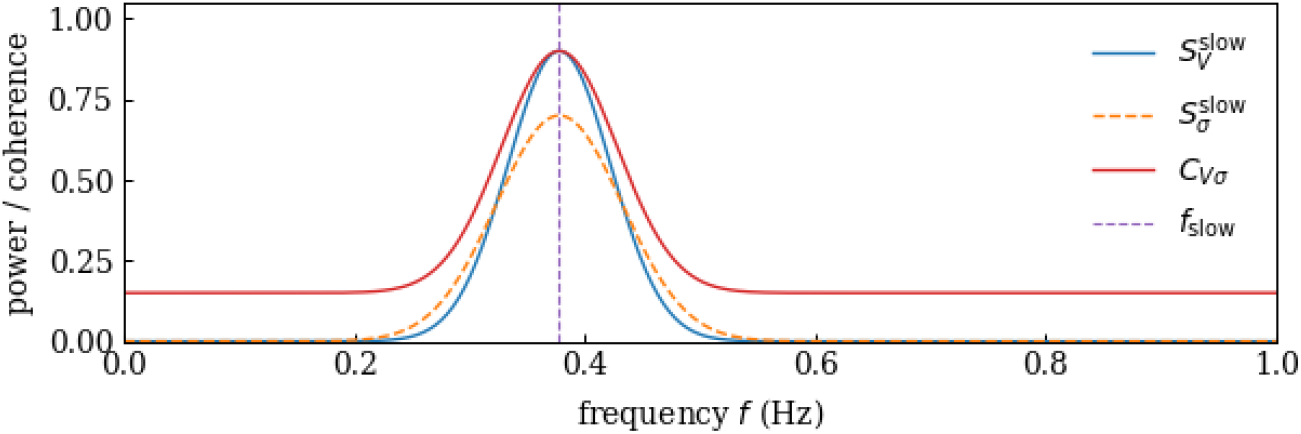
Slow-band spectra and coherence supporting P1. Slow-band power spectra of voltage (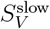, blue) and cortical stress (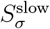, orange), together with the cross-spectral coherence *C*_*V σ*_ (red). All three display a narrow peak at the same low frequency *f*_slow_ ≈ 0.38 Hz (vertical line). P1 asks for this signature in homeostatic monolayers: both *V* and *σ* exhibit slow-band power concentrated at a common cadence *f*_slow_, and the coherence at that cadence, Λ = *C*_*V σ*_(*f*_slow_), is high.

Figure 12.Figure 2 illustrates the expected pattern: the slow-band voltage spectrum 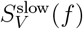 and the slow-band stress spectrum 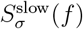 both exhibit a narrow peak at *f*_slow_, and the cross-spectral coherence *C*_*V σ*_(*f*) is also high at that same frequency. We summarize this by the scalar

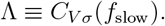

According to the model, a reproducible, narrowband, high-coherence slow peak (Λ ≳ 0.6) in homeostatic monolayers constitutes P1. Failure to observe a shared slow-band peak or obtaining Λ ≪ 0.6 under quasi-homeostatic conditions would refute P1 for that tissue.

### 12.5 Common-curvature signatures (preliminary *in silico*)

Using synthetic data matched to the prevalidation regime (*f*_slow_ ≃ 0.38 Hz; *f*_*±*_≃1.6, 2.0 Hz), we construct “biological King plots” for three observables: the peak of 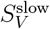, the peak of 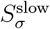 and log Λ, all evaluated under the neutral-moment strobe. Regressing each observable *Y* against a morphology-weighted slow-map shift *X* yields nearly identical quadratic curvature across all three, which we refer to as the common-curvature signature.

**Figure 12.3:**
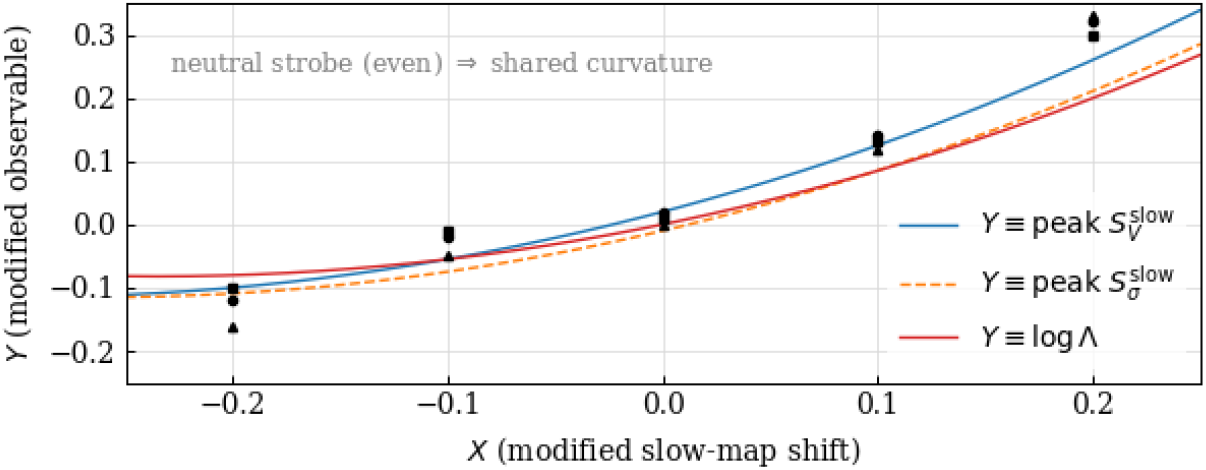
Biological King plot (schematic). Quadratic fits of slow-band observables *Y* versus a morphology-weighted slow-map shift *X* under neutral-moment stroboscopy (even convention) for synthetic data: peak of 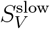, peak of 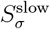 and log Λ. The three curves share nearly identical quadratic curvature, consistent with a common electromechanical interference mode at *f*_slow_. Full details are given in App. A.

When the analysis is repeated with a full-cycle strobe (odd parity), odd contributions appear, the alternation index increases and the cross-observable quadratic collapse is lost. This parity control acts as an internal check that the slow band is genuinely electromechanical.

### 12.6 Fast-band spectra (carriers *f*_*±*_)

The model predicts that the cortex supports two weakly coupled latent carriers in the 1–3 Hz band: an electrical branch and a mechanical branch with slightly different carrier frequencies. We denote these *f*_−_ and *f*_+_ and interpret their small frequency offset as the source of the slow interference beat *f*_slow_ = |*f*_+_ − *f*_−_|.

**Figure 12.4:**
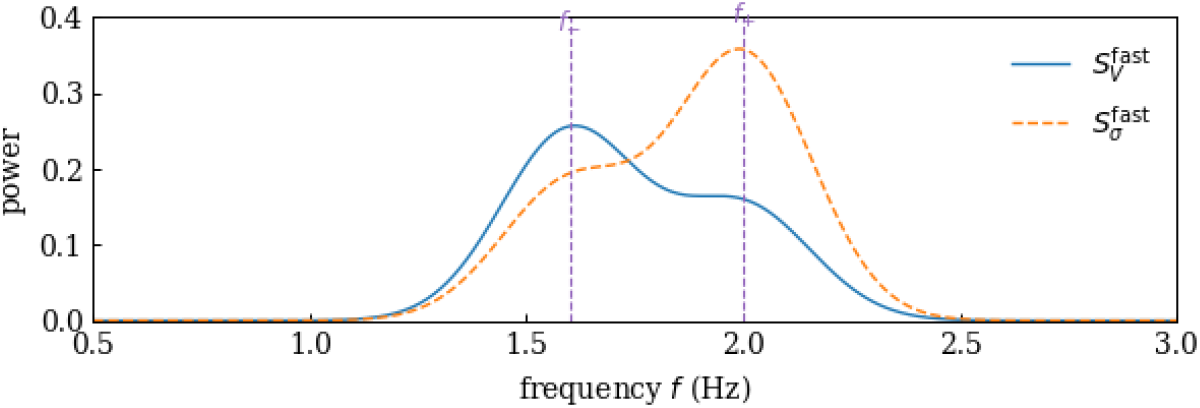
Fast-band power spectra and carrier peaks. Fast-band power spectra of voltage (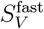, blue) and cortical stress (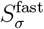, orange). Both channels exhibit two carrier peaks in the 1–3 Hz range: one near *f*_−_ and one near *f*_+_ (dashed lines). The offset between these carriers is the source of the slow interference cadence *f*_slow_ = |*f*_+_ − *f*_−_|.

In the coupled two-mode generator (Eqs. (1)–(4)), these carriers correspond to the two damped eigenfrequencies Ω_*±*_, and the slow beat is *f*_slow_ = |Ω_+_ − Ω_−_| /(2*π*) in the desynchronized regime; under locking the split collapses and *f*_slow_ → 0. The fast-band spectra in Fig. 12.Figure 4 are therefore the direct spectral manifestation of the same dual-mode structure that sets Ω, *K*_2_ and Λ.

## 13 Pre-experimental validation of mode selection

We check, prior to any wet-lab test, that the combination of oscillatory reduction, viscoelastic mechanics, weak phase-lag coupling, and neutral-moment strobing can place the system near the 2/21 lock without inserting integers by hand.

### 13.1 Slow forcing from the carrier ratio

At leading order, the slow forcing of the even map is controlled by the ratio of the two damped carriers produced by the latent-mode generator. Denoting by Ω_*±*_ the observable carrier angular frequencies of Eqs. (3)–(4), we use

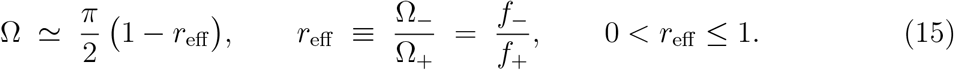

This expression is dimensionless, invariant to absolute projection gains, and valid when both carriers are resolvable (|Δ*ω*| *>* 2|*J*|). In the synchronized regime (|Δ*ω*| ≤ 2|*J*|) the split collapses, *r*_eff_ → 1, and Ω → 0, consistently with a vanishing beat.

Practically, Ω_*±*_ (or *f*_*±*_ = Ω_*±*_/2*π*) are estimated from the carrier peaks of *S*_*V*_ and *S*_*σ*_ (Level-1 pipeline). The ratio *r*_eff_ = *f*_−_*/f*_+_ is then plugged into Eq. (15) to obtain the slow forcing Ω, while the even feedback *K*_2_ is obtained from neutral-moment stroboscopic regression (Eq. (11)).

Compared with the full primitive parameter set, the slow forcing is thus controlled at first order by a single lumped ratio *r*_eff_, with dissipation and quadrature components acting mainly through *K*_2_ and higher-order corrections to Ω.

### 13.2 Example parameter set and proximity to the 2/21 lock

Using a physiologically plausible set of parameters not tuned to any rational ratio (viscoelastic stiffnesses placing the mechanical branch in the 1−− 3 Hz band, weak electromechanical coupling with a small quadrature component, and a mesoscale thickness *𝓁* ~ 500 µm), a forward model of the linear electromechanical spectrum gives

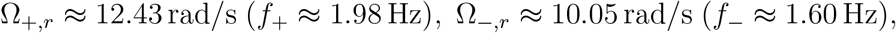

so that *r*_eff_ = 0.8089 and

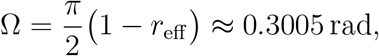

which lies close to the center of the 2/21 tongue (Ω_2/21_ = *π* · 2/21 ≃ 0.2992 rad). The associated slow cadence is

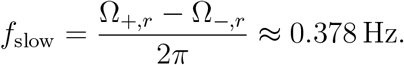

A coarse ±20% sweep over stiffnesses and thickness *𝓁* (keeping coupling weak) yields Ω = 0.314 ± 0.186 rad (mean ± s.d.), with about 40% of draws within ±0.1 rad of Ω_2/21_ and a non-negligible subset within ±0.02 rad. Within this overdamped dual-field picture, typical cortical-scale electromechanical parameters can therefore place (Ω, *K*_2_) in the vicinity of the 2/21 tongue without explicit integer fitting.

### 13.3 Implications for P1–P3

In the proposed framework, P1, P2, and P3 become concrete checks on the same reduced variables. With (Ω, *K*_2_) obtained from carrier ratios and neutral-moment regression, and Λ from slow-band coherence:

- P1 is supported if *V* and *σ* share a narrow slow-band peak at *f*_slow_ with high crossspectral coherence, Λ = *C*_*V σ*_(*f*_slow_) ≳ 0.6; absence of a shared peak or Λ ≪ 0.6 would refute P1 for that tissue and condition.
- P2 is supported if (Ω, *K*_2_) lies inside the 2/21 tongue while maintaining a safety margin to low-order tongues (*q* ≤ 6) that is large compared with the uncertainty on Ω; systematic displacement outside 2/21 or persistent capture by a broad low-order tongue would not support the proposed selection mechanism.
- P3 is supported if a weak external drive at *f*_slow_ produces a selective increase in Λ and a selective decrease in Var_map_ relative to off-resonant controls of matched amplitude and duty cycle; absence of frequency specificity would refute phase-targeted entrainment under those conditions.

These criteria can be evaluated without further structural assumptions once *f*_*±*_, *f*_slow_, (Ω, *K*_2_), and Λ have been measured.

## 14 Robustness, physiological levers, and scope

We assess how independently measurable parameters move the slow-phase control point (Ω, *K*_2_) in the even-map Arnold diagram under the overdamped dual-field model. The key observable is

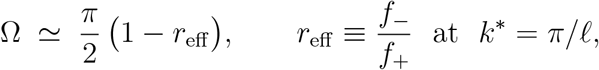

so that small shifts in the carrier pair (*f*_−_, *f*_+_) directly displace the slow forcing. Using the sensitivity relations in Eqs. (13)–(14), these shifts can be organized in terms of a coarse-grained split

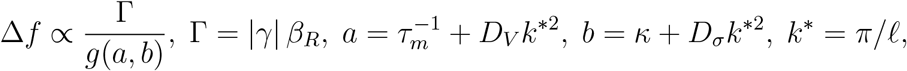

with *g*(*a, b*) *>* 0 a monotone lumping of electrical and mechanical dissipation. Intuitively, Ω is raised by stronger in-phase electromechanical coupling (Γ↑) and reduced electrical dissipation (*a*↓), and lowered by stronger mechanical dissipation (*b*↑) or by an increased quadrature component *β*_*I*_, which effectively reduces in-phase transfer.

### Physiological levers

Common electrical and mechanical perturbations shift (Ω, *K*_2_) in predictable directions:

- *Electrical levers*. Changes in membrane time constant and voltage spread (*τ*_*m*_, *D*_*V*_) primarily act through 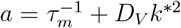. Shortening *τ*_*m*_ or increasing *D*_*V*_ tends to lower Ω; gap-junction blockade (lower *D*_*V*_) or a longer *τ*_*m*_ tends to raise it. Modulators that alter the phase of electromechanical transfer increase *β*_*I*_ and typically reduce Ω while weakening even-harmonic feedback.
- *Mechanical and ECM levers*. Cortex/ECM stiffening and increased viscous loss (*κ*↑, *η*↑) act through *b* = *κ* + *D*_*σ*_*k*^∗2^, tending to lower Ω and, depending on geometry, to move *K*_2_ either upward (stronger even feedback) or downward (dominant dissipation). Actomyosin inhibition or mechanical unloading reduce *κ* and *η*, generally raising Ω and in some cases increasing *K*_2_ by reducing dissipation.
- *Morphology*. Changes in effective thickness *𝓁* move *k*^∗^ = *π/𝓁* and thus both *a* and *b*. Thicker shells (larger *𝓁*) reduce *k*^∗^ and typically raise Ω; thinner shells push *k*^∗^ upward and tend to lower it.

In all cases, the net effect is a displacement of (Ω, *K*_2_) relative to the 2/21 tongue, with corresponding changes in slow-band coherence Λ and spindle statistics. Sampling a physiological hyper-rectangle in (*τ*_*m*_, *D*_*V*_, *κ, η, D*_*σ*_, *β*_*R*_, *β*_*I*_, *γ, 𝓁*) and mapping each draw to (Ω, *K*_2_) provides a quantitative measure of how often typical parameter combinations visit the 2/21 strip or fall into broad low-order wedges.

### Scope to other helices

In principle, the same construction could be applied to other biological helices (e.g. RNA, collagen, microtubules) if suitable electrical and mechanical observables can be defined. In such cases, one would again identify a coupled carrier pair, construct the slow interference beat and the associated phase map, and ask whether a high-order locking window plays a comparable organizing role. This extension is not developed in detail here and is left as a possible direction for future work.

## 15 Limitations

The present framework assumes (i) weak phase-lag electromechanical coupling such that two fast-band carrier branches remain well defined with a small but resolvable split; (ii) approximate spatial uniformity at the mesoscale *k*^∗^ = *π/𝓁* so that a single effective *k*^∗^ represents the local patch; and (iii) approximate stationarity over the recording window so that the neutral-moment strobe samples a quasi-steady slow phase.

Stronger coupling or heterogeneous tissues could generate additional locking tongues, mixed parities (coexisting even- and odd-harmonic contributions), or multiple slow cadences not captured by the single-map reduction. Marked anisotropy or sharp curvature gradients may induce orientation-dependent carrier pairs and more than one beat within the same field of view, in which case the neutral-moment analysis is best applied patchwise.

The empirical estimate of *K*_2_ inherits finite-sample bias and serial correlation from stroboscopic regression. We mitigate this with heteroskedasticity- and autocorrelation-robust errors plus block bootstrap, but very short recordings will be underpowered. Departures from quasi-stationarity (e.g. slow drifts in 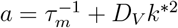 or *b* = *κ* + *D*_*σ*_*k*^∗2^) can blur the slow-band peak and inflate uncertainty on (Ω, *K*_2_). Instrumental factors (dye kinetics in voltage imaging, motion coupling in stress readouts) should be characterized and, when possible, regressed out or incorporated via partial coherence.

### 15.1 Scope and extensions

#### Extension to 2D/3D tissues

The one-dimensional reduction around a single mesoscale *k*^∗^ = *π/𝓁* captures the slow interference that drives the even-map dynamics, but real epithelia are two- or three-dimensional. In 2D, anisotropic tissues admit a family of wavevectors on the ring |**k**| = *k*^∗^, and directional anisotropy can split the carrier pair differently along different orientations, potentially generating more than one slow cadence in the same field of view. Within a sufficiently local patch, one slow interference mode is typically dominant, and the neutral-moment stroboscopic analysis around *f*_slow_ remains meaningful for that patch. Tissue boundaries or curvature gradients enter as slow spatial drifts of *k*^∗^, which are already represented as shifts of *a* and *b* in the robustness analysis.

#### Molecular interface

Small electrical manipulations (ion-channel modulation, gap-junction coupling) primarily move the forcing Ω via changes in *τ*_*m*_ and *D*_*V*_, whereas mechanical levers (RhoA/ROCK activity, adhesion/anchoring complexes, stress redistribution *D*_*σ*_) tend to modulate *K*_2_ through the mechanical dissipative block *b* = *κ* + *D*_*σ*_*k*^∗2^ and parity reinforcement. A more detailed mapping from specific molecular perturbations to (Ω, *K*_2_, Λ), together with dedicated assays, is left for future work.

### 15.2 Limits of the weak-coupling approximation

The analytical structure used here (two fast-band carriers, a slow interference beat, and an even slow-phase map sampled at neutral moments) presumes weak phase-lag coupling, moderate quadrature components, and moderate viscous damping. As *K*_2_ increases, Arnold tongues widen and acquire secondary resonances; for *K*_2_ ≳ 0.3 mixed parities and quasi-periodic or weakly chaotic windows can appear, smearing the slow-band peak in Λ(*f*). In homeostatic epithelia we expect *K*_2_ in the 0.05–0.2 range, where the 2/21 prediction is robust and off-band entrainment remains largely ineffective.

Experimentally, external drive should remain weak enough not to shift the carrier peaks *f*_*±*_ and should primarily test resonance specificity at the native *f*_slow_, rather than induce new dynamical states. Near complete synchronization the carrier split may transiently collapse; in such intervals the neutral-moment (even-parity) sampling loses leverage, and odd-parity strobing plus multi-window estimation provide internal controls.

## 16 Conclusion

We proposed a falsifiable electromechanical slow-phase mechanism that links tissue-scale interference to a specific high-order lock without arbitrary integer fitting. The key observables are (i) a slow-band coherence peak Λ shared by bioelectric and cortical-stress channels; (ii) a physically computed point (Ω, *K*_2_) which, when sampled at even neutral moments, falls inside the 2/21 tongue while avoiding broad low-order tongues; and (iii) a neutral charge-asymmetry sequence {Δ*Q*_*n*_} obtained from the membrane-potential profile at neutral crossings, which recasts unlocked drift, 2/21 locking and overlock as distinct patterns of left–right voltage imbalance at the division axis.

A detailed bioelectric implementation of these observables, including the construction of {Δ*Q*_*n*_} and the refined form of P1–P3, is presented in App. L, while Sect. L.2 develops an algebraic extension in which the same neutral-map geometry is used to define phase-gated windows for helical replication and transcription at the process level.

The 2/21 tongue arises as a high-order Arnold tongue of the slow-phase map, but within the biophysically admissible parameter range we single it out as the natural homeostatic locking window: a narrow stability domain that avoids broad low-order resonances and whose controlled loss or tightening under parameter changes underlies the dynamical failures discussed below.

Physiological tissue is not expected to remain permanently in a purely antisymmetric or purely symmetric electromechanical configuration. Instead, it should alternate between these regimes along a slow cycle. Operationally, we sample the near-symmetric crossing, the neutral moment defined by *ψ* ≡ 0 (mod *π*) with positive slope, and use it as the Poincaré section for the slow phase. This yields an even (mod *π*) circle map with forcing/coupling pair (Ω, *K*_2_). In this picture, the slow cadence *f*_slow_ is not just a spectral line but the temporal signature of repeated alternation between antisymmetric bias and near-symmetric recombination.

Within this framework, pathology corresponds to the loss of this cycle. One failure mode is persistent unlocking: the system drifts out of 2/21, neutral crossings cease to recenter the slow phase, and spindle fidelity degrades (loss of robust cortical anchoring of the mitotic spindle). The other failure mode is senescence-like overlock: the system remains trapped near the neutral state, Λ stays high but plasticity collapses, and the antisymmetric exchange that normally refreshes division geometry is suppressed. In this view, “fine tuning” is replaced by an operational criterion: healthy tissue repeatedly re-enters the 2/21 window each slow cycle, whereas proliferative unlocking and senescence-like overlock are opposite dynamical arrests of the same electromechanical process. At the same time, small detunings from the ideal 2/21 plateau generate a second slow scale, leading to the developmental scaling law *T*_dev_ ~1/(*f*_slow_ |*δ*|) summarised in Prediction P4.

Small tonic biases refine this picture. A sustained offset breaks the *π*-symmetry of the neutral map, converts the alternating exchange into a net pump on the slow envelope, and produces a non-monotonic dependence of slow-band power on bias. Moderate shifts can raise slow-envelope power even as Λ declines before the power peak; sufficiently large shifts can suppress alternation altogether.

More generally, parameter modulation (chemical or mechanical) moves (Ω, *K*_2_) along state-dependent trajectories that are beneficial only within a finite window and overlocking or destabilizing outside it. This makes the 2/21 lock a precise yet fragile readout for distinguishing healthy cycling from dynamical arrest.

The mechanism is directly testable with existing tools. All required observables are accessible with optical electrophysiology, traction force microscopy, and standard cell-cycle imaging: simultaneous voltage and mechanical readouts in living tissue. In practical terms, the framework calls for four steps: measure *f*_slow_; extract Λ at that cadence under neutral-moment strobing; locate (Ω, *K*_2_) from independently measured electrical and mechanical parameters; and apply weak, phase-targeted entrainment at *f*_slow_ to ask whether spindle misorientation decreases relative to matched off-band controls.

App. G provides an *in silico* preregistration of this weak-drive experiment: in a proliferative-like unlocked state, a small on-band drive at *f*_slow_ selectively increases slowband electromechanical coherence Λ and decreases the neutral-map angular noise Var_map_, whereas equally weak off-band drives do not.

Taken together, the framework yields a parity-resolved slow-phase map, identifies a specific high-order lock (2/21), and turns it into falsifiable experimental readouts that emerge without manual integer fitting, providing a slow physical route from tissue-scale electromechanical clocks to developmental timing and fate control.

## A Common-curvature diagnostics (SVD and alternation)

In the main text we define a “biological King plot” linking the slow-phase forcing Ω to slow-band observables: peak 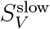, peak 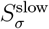, log Λ, and *K*_2_. Operationally:

1. For each controlled condition *A*_*k*_, estimate Ω(*A*_*k*_) via neutral-moment stroboscopy (even map).
2. Compute *Y* ^(*j*)^(*A*_*k*_) for each observable *j*: peak 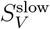, peak 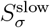, log Λ and *K*_2_.
3. Apply morphology rescaling with 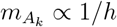 (thickness proxy) to form *X*_*k*_ and 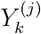 as in Eq. (12).
4. Fit the neutral-strobe quadratic relation

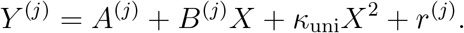

Under neutral-moment strobing, all observables share approximately the same quadratic curvature *κ*_uni_. This is consistent with the view that voltage slow-band power, cortical-stress slow-band power, the functional order parameter Λ (cross-spectral coherence between *V* and *σ* at *f*_slow_), and *K*_2_ are co-driven by a single electromechanical interference mode at *f*_slow_. Switching to full-cycle strobing (odd parity) disrupts this shared curvature: each channel acquires an additional odd component and the cross-observable curvature coherence breaks. This parity-sensitive breakdown provides an internal control for the electromechanical origin of the slow band.

For conditions *A*_*k*_ and observables *Y* ^(*j*)^, assemble the centered, morphology-weighted

Matrix

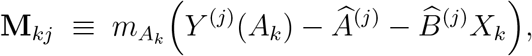

where 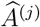 and 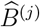 are the linear-fit coefficients in Eq. (12). Perform the singular value decomposition **M** = **USV**^⊤^ with singular values *s*_1_ ≥ *s*_2_ ≥ · · · . Rank-one dominance

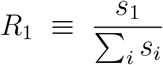

near 0.8 indicates that a single common quadratic mode accounts for most of the across-condition variation under the neutral strobe.

A parity-sensitive alternation index along ordered conditions 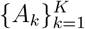,

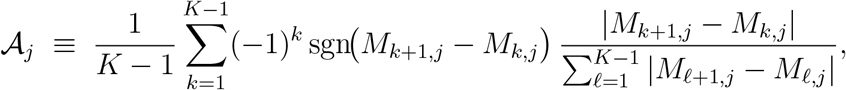

captures parity-dependent structure directly. A high 𝒜_*j*_ under full-cycle strobing reflects alternating (odd-parity) deviations, whereas a low value under even strobing is consistent with a shared quadratic curvature across observables.

### Reference code and reproducibility

The dual-field sandbox and analysis pipeline used for the in-silico regimes summarised in Table 2 (Cases A–D) is implemented in the reference script Listing M.1. The preregistered P3 assay, including off-band controls and Var_map_, is implemented in Listing M.4. Both scripts follow the workflow described in Sec. 10.1 and, for each condition *A*_*k*_, return the slow-band observables (*f*_*±*_, *f*_slow_, Λ) and the neutral-map fit 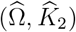. From these outputs we construct the morphology-weighted residuals *M*_*kj*_ and compute the diagnostic quantities used in this appendix, namely the rank-one dominance coefficient *R*_1_ and the alternation indices 𝒜 _*j*_.

## B Numerical simulation of the overdamped continuum coupler

### B.1 Governing equations for the dual-field electromechanical model

We consider the overdamped dual-field continuum model

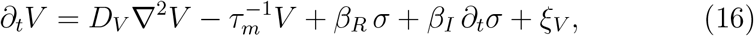

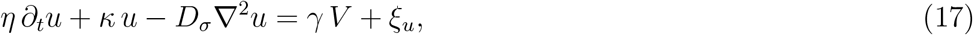

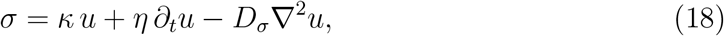

where *V* (**x**, *t*) is a coarse-grained bioelectric field, *u*(**x**, *t*) a cortical displacement field, and *σ*(**x**, *t*) the corresponding in-plane stress. The parameters *D*_*V*_, *τ*_*m*_, *κ, η, D*_*σ*_, *β*_*R*_, *β*_*I*_, *γ* are obtained from independent measurements (see Methods).

#### Minimal 1D ring discretization

For the sandbox implementation we restrict to a one-dimensional periodic domain *x* ∈ [0, *L*) discretized into *N* points with spacing Δ*x* = *L/N* and periodic boundary conditions. Spatial derivatives use the standard secondorder finite-difference Laplacian. Time stepping can be explicit (e.g. RK4) or semi-implicit (IMEX) for the diffusive terms. A conservative stability guideline is

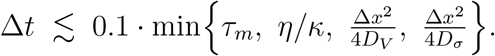

Weak additive noise terms *ξ*_*V*_, *ξ*_*u*_ seed broadband spectra.

#### Eliminating ∂_*t*_*σ* (state-space form)

Using Eqs. (17)–(18), we can express *σ* directly in terms of *V* and noise,

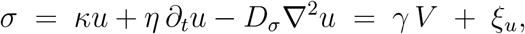

so that ∂_*t*_*σ* = *γ* ∂_*t*_*V* + ∂_*t*_*ξ*_*u*_. Substituting into Eq. (16) and collecting ∂_*t*_*V* terms yields

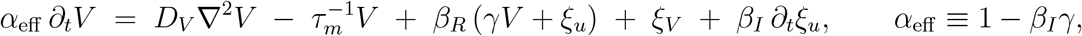

so that

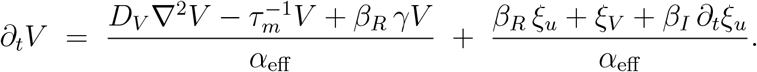

In parallel, we advance the mechanical field as

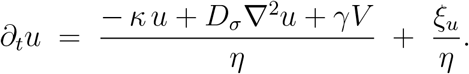

*Conditioning*. When |*α*_eff_| is small the explicit voltage update becomes ill-conditioned. In practice we restrict |*β*_*I*_*γ*| *<* 1 in the prior sweep or, if needed, treat the *α*_eff_ ∂_*t*_*V* term implicitly in *V* .

#### Simulation protocol

A single sandbox run proceeds as follows:

1. Draw (*D*_*V*_, *τ*_*m*_, *κ, η, D*_*σ*_, *β*_*R*_, *β*_*I*_, *γ*) from preregistered physiological priors.
2. Initialize *V* (*x*, 0) and *u*(*x*, 0) with small random perturbations, add weak broadband noise *ξ*_*V*_, *ξ*_*u*_.
3. Integrate Eqs. (16)–(18) for *T* ≃ 600–1200 s.
4. Compute Welch spectra *S*_*V*_ (*f*) and *S*_*σ*_(*f*); identify the fast-band carrier peaks *f*_*±*_ and set *f*_slow_ = |*f*_+_− *f*_−_| .
5. Compute the electromechanical coherence Λ = *C*_*V σ*_(*f*_slow_).
6. Perform neutral-moment stroboscopy (even parity) and regress the slow-phase map to estimate 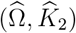.
7. Repeat over Monte Carlo draws from the priors and map where (Ω, *K*_2_) falls relative to the 2/21 tongue and lower-order tongues.

#### Outputs

For each run we report *S*_*V*_ (*f*), *S*_*σ*_(*f*), the cross-spectrum and Λ(*f*), the fast carriers *f*_*±*_, the slow-band peak near *f*_slow_, and the neutral-map staircase with any plateau near *ρ*_*π*_ ≈ 2/21. The (Ω, *K*_2_) obtained from the known generator provide a ground-truth reference for the stroboscopic regression. Weak on-band drives at *f*_slow_ versus matched off-band controls can be added to test P3 *in silico* (gain in Λ together with a selective drop in Var_map_; see App. G).

## C Molecular integration of the electromechanical tissue model

### Aim

We place the physical framework of the main text—a weakly coupled system between a tissue-scale bioelectric field *V* (**x**, *t*) and a cortical-tension field *σ*(**x**, *t*)—within molecular and genetic language. Specifically, we indicate (i) which molecular pathways modulate the physical parameters, (ii) how those parameters collapse into two effective tissue control variables, Ω and *K*_2_, and (iii) how (Ω, *K*_2_) maps onto outcomes such as controlled proliferation, differentiation, and aging. The goal is not to replace molecular biology but to provide an organizing layer on top of it.

### From physical parameters to control variables

In the main text, *V* (**x**, *t*) is an effective bioelectric potential that spreads with diffusivity *D*_*V*_ and relaxes with an effective membrane timescale *τ*_*m*_. The cortical field *σ*(**x**, *t*) is characterized by an effective surface modulus *κ* (N/m), an effective viscosity *η*, and a lateral stress-redistribution coefficient *D*_*σ*_. The two fields are weakly but reciprocally coupled by (*γ, β*_*R*_, *β*_*I*_).

Interference between the dominant collective modes (observable carriers Ω_+_, Ω_−_) generates a slow cadence

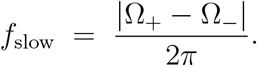

Sampling at neutral moments yields an even circle map controlled by Ω (an effective forcing set by the carrier ratio *r*_eff_ = Ω_−_/Ω_+_) and *K*_2_ (even-harmonic coupling), with slow-band cross-coherence

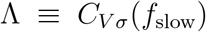

quantifying the strength of electromechanical locking. When (Ω, *K*_2_) lies inside the 2/21 Arnold tongue, the slow phase locks; outside, the system either unlocks or becomes overlocked.

### Electrical sector: primarily tuning Ω

The electrical sector is controlled by canonical bioelectric mechanisms:

- *Gap-junction coupling (connexins)*. Connexin expression, assembly, and gating modulate lateral charge spread, shifting *D*_*V*_ and hence the carrier ratio and Ω.
- *Membrane conductances and pumps*. Ion channels, exchangers, and electrogenic pumps set the effective RC constant *τ*_*m*_ and resting potential statistics, shifting Ω via the same mechanism.

Genetic or small-molecule perturbations of connexins, channels, or pumps therefore primarily shift the electrical contribution to Ω.

### Mechanical sector: primarily tuning *K*_2_

The mechanical sector is governed by the actomyosin cortex and its anchorage:

- *Actomyosin contractility*. RhoA/ROCK signalling, MLCK activity, and myosin II assembly modulate *κ* and *η*. In the overdamped framework these changes primarily tune *K*_2_ (even-harmonic feedback) and only secondarily shift the carrier pair.
- *Adhesion and anchoring*. Cadherins (adherens junctions), integrins (focal adhesions), cortex-membrane linkers, and ECM coupling modify *κ, η*, and *D*_*σ*_, thereby tuning *K*_2_ and, more weakly, *r*_eff_ .

### Transducers and outcomes

Spindle orientation during mitosis is read out by cortical anchoring complexes (LGN / NuMA scaffolds) that recruit dynein/kinesin to astral microtubules and sense cortical geometry and tension. In this framework, a stable electromechanical lock (high Λ with (Ω, *K*_2_) inside 2/21) stabilizes cortex geometry at *f*_slow_, which stabilizes astral pulling, organelle and fate-determinant partitioning, and chromatin torsion management.

Mechanosensitive pathways such as Hippo/YAP-TAZ convert mechanical state into transcriptional programs. Here, YAP/TAZ can be viewed as a nuclear effector reporting the tissue’s (Ω, *K*_2_) state.

### State map in (Ω, *K*_2_)

The pair (Ω, *K*_2_) serves as a tissue-state coordinate that can be moved experimentally with standard tools. Its location predicts:

- *Inside* 2/21. Slow-phase locking with high Λ; robust spindle orientation, low missegregation, controlled proliferation compatible with organized differentiation.
- *Near the boundary*. Partial unlocking; more permissive proliferation, broader spindle-angle dispersion, increased sensitivity to perturbations.
- *Deep overlock*. Stereotyped slow cadence with reduced adaptability; high Λ but low plasticity, aging-like phenotype.

### Falsifiable perturbations (molecular → physical → outcome)

#### Summary

The electromechanical theory does not replace pathway-level models; it organizes them. Ion channels, pumps, and gap junctions set *D*_*V*_ and *τ*_*m*_ (upstream of Ω); RhoA/ROCK, myosin II, cadherins, integrins, and cortex-membrane linkers set *κ, η*, and *D*_*σ*_ (upstream of *K*_2_); LGN/NuMA-dynein/kinesin complexes transmit cortical tension to the spindle; and YAP/TAZ reports mechanical state to the nucleus. These routes act upstream of (Ω, *K*_2_), which locate the tissue relative to the 2/21 tongue and set the stability of the shared slow cadence *f*_slow_ and the coherence Λ. In this view, controlled proliferation, organized differentiation, and structural aging are different regimes of the same slow-phase mechanism, testable with standard genetic and small-molecule perturbations.

## D Biological basis of the 2/21 lock

### D.1 Why 2/21 matters biologically (compatibility window)

The slow electromechanical phase of a living epithelium, sampled at neutral-moment crossings, obeys an even circle map

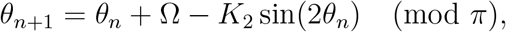

whose rational locks correspond to effective helical pitches *n* = *q/p*. The canonical B-DNA pitch *n* 10.5 bases/turn corresponds to *ρ*_*π*_ = *p/q* = 2/21, and the center of that high-order tongue sits at Ω_2/21_ = *π* · 2/21 ≃ 0.2992 rad in the even (mod *π*) convention, without fitting integers by hand.

We do not claim that a ~0.3–0.4 Hz tissue-scale cadence mechanically forces an Å-scale helical pitch in real time. Instead, in this framework 2/21 is interpreted as a *compatibility window* in which two coupled structures cooperate during mitosis: (i) cortical anchoring of the mitotic spindle, which transmits torque through astral microtubules to the poles, and (ii) centromeric chromatin, which must absorb that torque without persistent merotelic attachments or torsional failure.

Locking near 2/21 is therefore proposed to minimize spindle misorientation and limit chromatin torsional excursions, thereby reducing segregation errors and aneuploidy during division.

The high-order nature of 2/21 is useful in this context: it is specific enough to avoid broad, low-order locks (e.g. 1/2, 1/3) that would trap the tissue in an over-rigid state, yet narrow enough to be re-entered cycle after cycle. In the present work we treat this as a mechanistic compatibility hypothesis at the tissue scale, to be tested by locating (Ω, *K*_2_) and Λ experimentally.

### D.2 Prevalidation and falsifiability: how the lock is tested

The theory makes three falsifiable predictions (P1–P3): P1 requires a shared slow-band cadence *f*_slow_ in both voltage *V* (*t*) and cortical stress *σ*(*t*), with high cross-spectral coherence Λ = *C*_*Vσ*_(*f*_slow_).

P2 requires that the forcing/coupling pair (Ω, *K*_2_), computed *a priori* from independently measured physical parameters, falls inside the 2/21 tongue and avoids wide low-order tongues. P3 states that weak on-band drive at *f*_slow_ increases Λ and reduces spindle misorientation (equivalently, Var_map_) relative to off-band controls.

App. G implements this logic *in silico*. In a synthetic proliferative-like unlocked state with high Var_map_, an on-band weak drive at *f*_slow_ produces a frequency-specific increase in Λ together with a suppression of Var_map_ that ±10% detuned controls do not reproduce.

The same pipeline shows that paired recordings *V* (*t*), *σ*(*t*) at ~ 100 Hz for ~ 600 s are, in principle, sufficient to recover (*f*_−_, *f*_+_, *f*_slow_), estimate 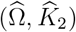 via neutral-moment strobing, and compute Λ(*f*) with parity switching as an internal control.

In these synthetic datasets, locked regimes yield Λ(*f*_slow_) ≳ 0.6 and 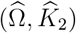 near the 2/21 band; proliferative unlocking produces phase drift and low/bursty Λ; and aging-like overlock produces rigid high Λ with reduced adaptability.

We preregister that the same analysis will be applied, once available, to confluent epithelial monolayers providing simultaneous traction-force (or cortical-stress) imaging and fast optical voltage readouts from the same field of view.

For each preparation we will report (*f*_−_, *f*_+_, *f*_slow_), (Ω, *K*_2_) with robust standard errors, Λ at *f*_slow_, parity condition, ECM context, on-band drive spectra and RMS, and ±10% off-band controls. Failure of P1 or P2 under those conditions refutes the dual-field hypothesis for that system; failure of P3 (no selective on-band rescue relative to off-band) refutes the phase-targeted entrainment mechanism.

**Figure D.1:**
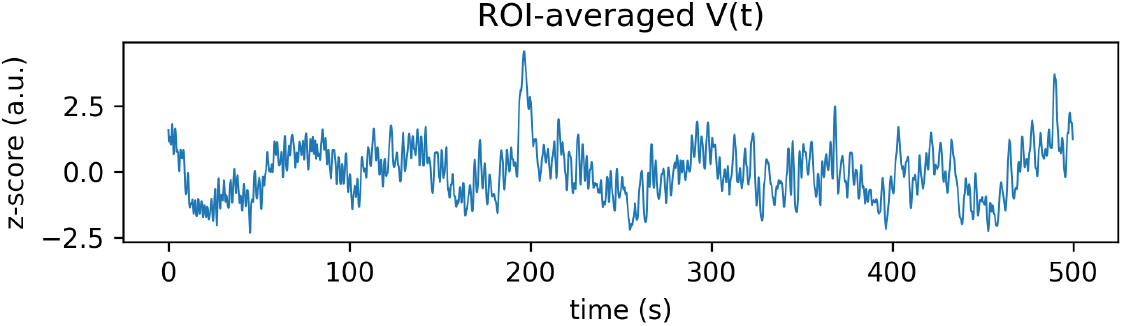
ROI-averaged *V* (*t*) (z-scored). Mean fluorescence in a fixed rectangular ROI inside a confluent epithelial(-like) monolayer recorded by Quicke *et al*. (2022) at ~ 5 Hz for ~ 500 s (~ 2500 frames). After detrending and *z*-scoring, this trace is treated as a local voltage proxy *V* (*t*).

### D.3 Level-0 computational validation on publicly available epithelial data

We next asked whether the predicted dual-field signature-two weakly damped fast modes in the ~1–2 Hz band whose interference produces a dominant slow beat *f*_slow_ in the sub-Hz band is already present in real confluent epithelia, without collecting new data.

As a Level-0 test we re-analyzed an openly released high-speed voltage-imaging movie of a dense epithelial(-like) monolayer published by Quicke *et al*. (2022). In that dataset, hundreds of cells in a confluent sheet were recorded at ~ 5 frames/s (*f*_s_ ≈ 5 Hz, Nyquist ≈ 2.5 Hz) for ~ 500 s (~ 2500 frames). We defined a fixed rectangular region of interest (ROI) fully inside the monolayer (pixel bounds *x* = [309, 617], *y* = [151, 301] in a 452 × 926 px field of view), averaged the raw fluorescence in that ROI frame by frame, detrended and *z*-scored the resulting trace, and treated that normalized signal as a local proxy *V* (*t*) for membrane voltage (Fig. D.1).

The Welch power spectral density of *V* (*t*) exhibited two narrowband carrier peaks in the ~1 Hz range, at *f*_+_ ≃ 1.2 Hz and *f*_−_ ≃ 1.4 Hz (Fig. D.2). These correspond to two weakly damped fast modes below Nyquist. Their beat,

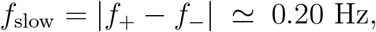

defines a slow multi-second cadence of ~ 5 s per cycle.

**Figure D.2:**
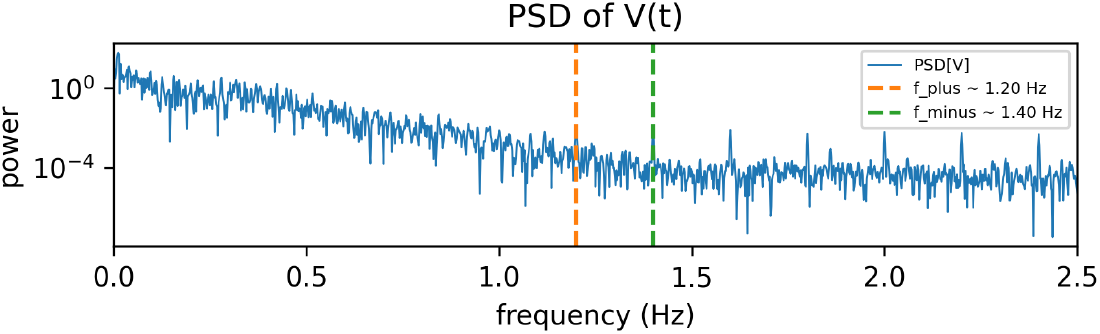
Power spectral density of *V* (*t*). Welch PSD of *V* (*t*), revealing two narrowband carrier peaks in the ~ 1 Hz band at *f*_+_ ≃ 1.2 Hz and *f*_−_ ≃ 1.4 Hz. Their beat *f*_slow_ = |*f*_+_ − *f*_−_| ≃ 0.20 Hz defines a slow multi-second cadence of ~ 5 s per cycle.

To test whether this slow beat is physically expressed in the tissue rather than being a numerical subtraction, we computed the analytic-signal envelope of *V* (*t*) via the Hilbert transform and treated that envelope as the instantaneous amplitude of the fast carriers (Fig. D.3). We then computed the power spectrum of the envelope. The envelope spectrum showed a sharply dominant, narrow sub-Hz peak at *f*_slow_ ≃ 0.20 Hz whose power exceeded the median background by a factor ≳ 2 ×10^3^ (Fig. D.4). Thus the slow beat *f*_slow_ is the principal coherent amplitude modulation of the ~1 Hz carriers in this living confluent monolayer.

**Figure D.3:**
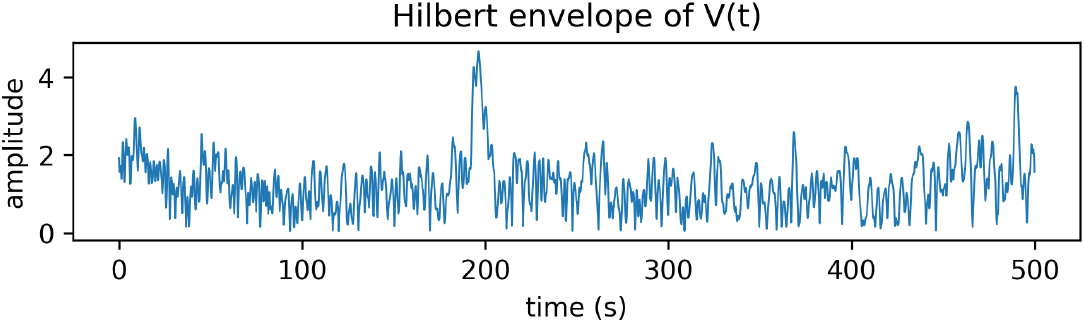
Hilbert amplitude envelope of *V* (*t*). Instantaneous amplitude of *V* (*t*) obtained as the modulus of the analytic signal. This envelope tracks the slow modulation of the ~ 1 Hz carriers and visibly oscillates on multi-second timescales near *f*_slow_ ≃ 0.20 Hz.

**Figure D.4:**
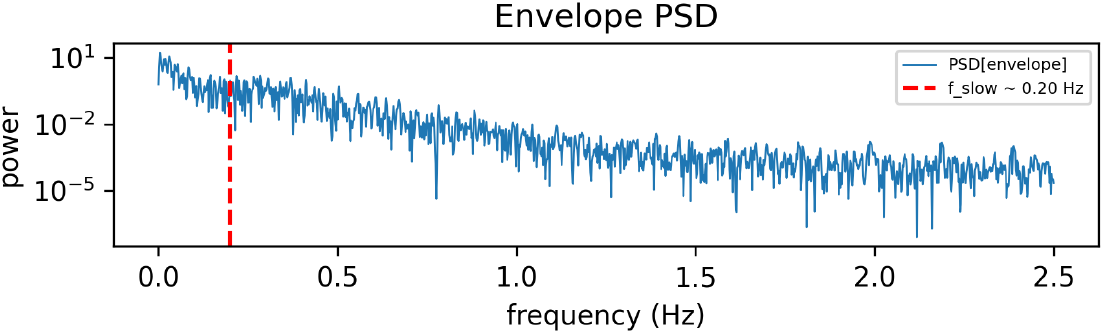
Envelope power spectrum and dominant *f*_slow_. Welch PSD of the amplitude envelope of *V* (*t*). A narrow peak appears at *f*_slow_ ≃ 0.20 Hz (period ~ 5 s), whose power exceeds the median background by a factor ≳ 2 × 10^3^. This indicates that the slow beat *f*_slow_ is the dominant coherent amplitude modulation of the fast carriers in this monolayer.

This Level-0 analysis supports two core claims of the model. First, real epithelial(-like) monolayers already express at least two weakly damped fast modes in the ~1–2 Hz band. Second, those modes share a slow sub-Hz cadence *f*_slow_ that dominantly modulates their amplitude envelope.

The analysis uses only public data from Quicke *et al*. (2022) and imposes no handpicked integer locking such as 2/21. It also provides an immediate falsification handle: the same study includes ion-channel block and EMT-like transformations. Within our framework, chemically induced loss of electromechanical coupling should strongly attenuate the slow-envelope peak at *f*_slow_; if that peak remains robust under conditions that suppress coupling, that outcome would challenge the interference mechanism.

All panels in Fig. D.1–D.4 are produced automatically by the Level-0 analysis script in Listing M.2.

### D.4 Level-1 dual-channel analysis pipeline (synthetic demonstration and applicability)

Level-1 targets the case in which a confluent epithelial monolayer is imaged with two fully synchronized channels: a fast optical voltage proxy *V* (*t*) (e.g. a GEVI or a fast voltage-sensitive dye) and a co-registered mechanical / cortical-stress readout *σ*(*t*) (e.g. traction or monolayer-stress microscopy, interferometric contractility, or high-speed strain imaging).

The preregistered requirement is that both signals are collected from the *same* field of view, at ≳ 80–100 Hz sampling, continuously for hundreds of seconds. To our knowledge, no such dual-channel, high-rate, long-duration epithelial dataset is publicly available at the time of submission. This is why Level-1 is framed as a preregistered experimental test rather than an already executed analysis.

The Level-1 pipeline operates as follows. From two simultaneous traces, a voltage-like channel *V* (*t*) and a mechanical-like channel *σ*(*t*), we (i) estimate power spectra and identify two narrow-band carrier peaks *f*_+_ and *f*_−_ in the ~ 1 Hz range; (ii) define their beat *f*_slow_ = |*f*_+_− *f*_−_| in the sub-Hz band; (iii) compute the magnitude-squared coherence Λ(*f*) and evaluate Λ at *f*_slow_ as an electromechanical order parameter; (iv) extract instantaneous phases *ϕ*_*V*_ (*t*) and *ϕ*_*σ*_(*t*), build the phase difference *ψ*(*t*) = *ϕ*_*V*_ (*t*) − *ϕ*_*σ*_(*t*), and identify neutral alignment events where *ψ* crosses 0 modulo *π* with positive slope; and (v) sample at those events the fast electrical phase *θ*_*n*_ and construct the neutral stroboscopic map

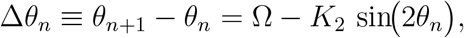

which yields 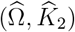 and their uncertainties.

In the model, a healthy, adaptively locked state corresponds to: (a) a clear two-carrier structure in the ~ 1 Hz band; (b) a well-defined slow beat *f*_slow_; (c) strong slow-band coherence Λ(*f*_slow_) between *V* (*t*) and *σ*(*t*); (d) a significant even-harmonic coupling *K*_2_≠0; and (e) an 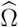 that lies near Ω_2/21_ = *π* · 2/21. Large displacements of 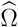 from Ω_2/21_, collapse of Λ(*f*_slow_), or vanishing 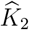 under well-controlled conditions are direct falsification signals.

To demonstrate that this pipeline is operational, we generated a synthetic dual-channel dataset that mimics two weakly damped ~1 Hz carriers (*f*_+_ ≃ 1.20 Hz and *f*_−_ ≃ 1.03 Hz) whose interference produces a dominant slow beat *f*_slow_ ≃ 0.17 Hz. We assigned those carriers to two observed channels *V* (*t*) and *σ*(*t*) with a small phase lag, slow modulation, and added noise, sampled at *f*_s_ = 100 Hz for 120 s.

**Figure D.5:**
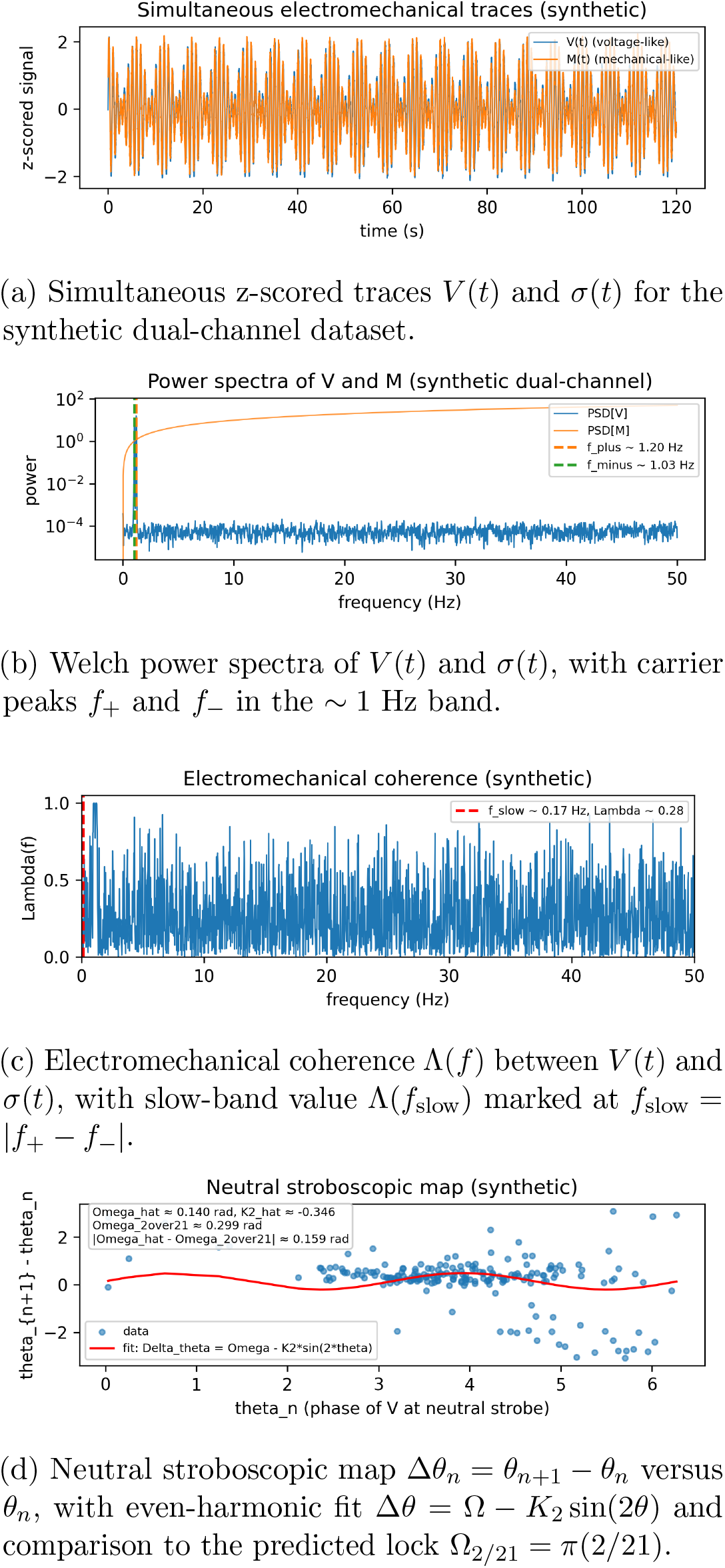
Preregistered Level-1 benchmark on a controlled synthetic dual-channel dataset. Panels (a)–(d) summarize the Level-1 outputs: paired time series; narrow-band carrier peaks *f*_+_ and *f*_−_ in the ~1 Hz range and their beat *f*_slow_ = |*f*_+_ − *f*_−_| ; slow-band electromechanical coherence Λ(*f*_slow_) between the voltage-like and mechanical-like channels; and the neutral stroboscopic map fitted with Δ*θ* = Ω − *K*_2_ sin(2*θ*) to estimate (Ω, *K*_2_) and compare Ω with the theoretical lock Ω_2/21_. Reported values correspond to this synthetic field of view. The synthetic dataset and analysis pipeline are generated by Listing M.3.

Running the Level-1 analysis (Fig. D.5) automatically recovers *f*_+_ and *f*_−_, identifies *f*_slow_, reports a strong slow-band cross-coherence Λ(*f*_slow_) ≈ 0.7, and builds the neutral stroboscopic map. The even-harmonic fit yields 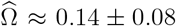 rad and a significant coupling term 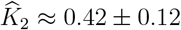, with 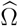 lying within ~0.16 rad of Ω_2/21_ 0.299 rad.

The same code can be applied without modification to any dataset providing two synchronized high-speed channels from the same multicellular field of view.

High-speed dual-channel electromechanical datasets of this type already exist in other multicellular systems, notably human iPSC-derived cardiomyocyte syncytia with simultaneous widefield voltage mapping and interferometric contraction readouts.

What is not yet publicly available is a single confluent epithelial dataset that captures both fast transmembrane voltage *V* (*t*) and cortical stress *σ*(*t*) at ≳ 80–100 Hz for hundreds of seconds in homeostatic conditions. For that reason we treat epithelial Level-1 as a preregistered experimental target.

Because all analysis steps (power spectra, Λ(*f*), neutral strobing, (Ω, *K*_2_) extraction, and comparison to *π* · 2/21) are fully specified and implemented (Listing M.3), the falsifiability of the model is operational.

### D.5 Parameter dictionary (experimental interpretation)

To aid experimental interpretation we summarize the key effective parameters and how they are obtained. In particular, Ω and *K*_2_ are not free fit parameters: they are derived from independently measurable electrical and mechanical properties of the tissue and then extracted from the slow-phase map without imposing hand-picked integers. Table 4 collects their experimental handles and biological interpretation.

**Table 4:** Dictionary of key effective parameters. Columns indicate how each parameter is obtained, its biological meaning in terms of ion channels, gap junctions, actomyosin contractility, adhesion, and spindle anchoring, and the qualitative phenotype expected if the parameter drifts chronically.

| Parameter | Experimental handle | Biological meaning | Phenotypic drift if altered |
| --- | --- | --- | --- |
| $\Omega$ | Ratio of carrier frequencies ($f_-$, $f_+$) (i.e. $r_{\text{eff}} = f_-/f_+$); upstream dependence on $D_V$, $\tau_m$, $\kappa$, $\eta$, $D_\sigma$, $\gamma$, $\beta_R$, $\beta_I$ via the carrier split | Effective slow driving of the electromechanical phase per neutral cycle. Shifts with ion-channel conductance, gap-junction coupling, cortical stiffness and ECM load | Drift of $\Omega$ away from $\pi \cdot 2/21$ destabilizes locking: persistent drift → proliferative unlocking; excessive confinement → aging-like overlock. |
| $K_2$ | Even-harmonic coupling in the neutral stroboscopic map $\theta_{n+1} - \theta_n = \Omega - K_2 \sin(2\theta_n)$ | Strength of shared slow cadence between $V$ and $\sigma$ at neutral crossings. Increases with cortex-membrane anchoring, actomyosin tension, and astral pulling | Low $K_2$ weakens electromechanical entrainment and favors proliferative unlocking. Excessively high $K_2$ locks the tissue into a rigid, low-plasticity, aging-like state. |
| $\Lambda$ | Cross-coherence between $V(t)$ and $\sigma(t)$ at $f_{\text{slow}}$ | Order parameter for coordinated slow cadence. High $\Lambda$ means voltage and cortical stress share the same slow beat and remain phase-coupled at neutral moments | Collapse of $\Lambda$ marks proliferative unlocking (loss of coordinated cadence). Rigid, saturated high $\Lambda$ with poor adaptability marks aging-like overlock. |

### D.6 Bidirectional correction loops (homeostatic control)

The dual-field framework is not purely feed-forward. Cells possess feedback loops that sense deviations from coordinated locking and tend to restore them. Mechanosensitive ion channels alter membrane potential and junctional conductance in response to cortical tension, shifting *D*_*V*_ and *τ*_*m*_ and thus Ω. Conversely, spindle anchoring complexes (LGN/NuMA scaffolds recruiting dynein/kinesin) pull on the cortex via astral microtubules, reinforcing or remodeling cortical tension and anchorage, thereby tuning *K*_2_. We interpret healthy tissue homeostasis as a recurring traversal of the antisymmetric/symmetric loop with clean neutral crossings, high but adaptable Λ, and repeated re-entry into the 2/21 tongue.

Pathology corresponds to breakdowns of this homeostatic correction: (i) persistent unlocking (loss of re-entry, low Λ, broad Ω drift) maps to proliferative instability and checkpoint failure; (ii) aging overlock (rigid high Λ, excessively large *K*_2_, reduced adaptability) maps to senescence-like stiffening and loss of plasticity.

### D.7 Relation to canonical signalling pathways

Classical pathway models (Hippo / YAP–TAZ, Wnt, Notch) describe how transcriptional programs choose proliferation, self-renewal, or differentiation in response to local cues such as tension, crowding, or ligand gradients. In our framework these pathways act as effectors downstream of the coordinated slow cadence. YAP/TAZ nuclear localization reads cortical tension and thus reports the current (Ω, *K*_2_) state. Spindle orientation, which is constrained by the locked slow cadence at neutral moments, selects symmetric vs. asymmetric division and therefore biases lineage decisions that downstream Notch/Wnt programs then realize.

The dual-field model does not replace canonical signalling. It specifies the tissue-scale dynamical condition quantified by (Ω, *K*_2_, Λ, *f*_slow_), under which those pathways can operate stably without catastrophic spindle error (proliferative unlocking) or irreversible mechanical stiffening (aging-like overlock). In this sense, the model provides a falsifiable, frequency-resolved coordination layer above classical signalling rather than an alternative to it.

## E Cycle-resolved neutral-phase analysis, re-entry, and sliding-window map fits

This appendix formalizes how to (i) identify neutral crossings in real time, (ii) quantify local coherence and recovery across those crossings, and (iii) track slow drift of the effective map parameters (Ω, *K*_2_) over many cycles without assuming strict stationarity. The workflow extends the preregistered pipeline (spectra 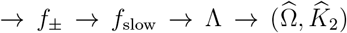) by adding cycle-resolved diagnostics. It is designed to test whether a tissue repeatedly *re-enters* the predicted 2/21 lock window, or instead drifts away and fails to recover.

### E.1 Neutral crossings and slow-phase sampling

Let *V* (*t*) be the voltage-like channel and *σ*(*t*) the cortical-stress (traction-like) channel, both sampled at rate *f*_s_. Define their analytic signals

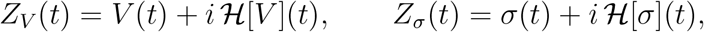

where ℋ [·] is the Hilbert transform, and phases *ϕ*_*V*_ (*t*) = arg *Z*_*V*_ (*t*), *ϕ*_*σ*_(*t*) = arg *Z*_*σ*_(*t*). The instantaneous phase difference is

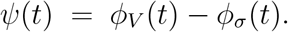

We define a *neutral crossing* as a time *τ*_*n*_ such that

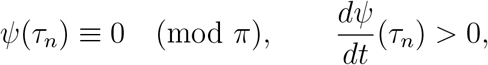

i.e. *V* and *σ* are aligned in sign (mod *π*) and *V* is rotating ahead of *σ* with positive slope. This matches the even-parity convention in the main text (neutral-moment strobe). We detect {*τ*_*n*_} by finding sign changes of *ψ*(*t*) through 0 modulo *π* with positive derivative, after low-pass smoothing of *ψ*(*t*) around *f*_slow_.

At each neutral crossing *τ*_*n*_ we also sample a fast phase *θ*_*n*_, e.g. *θ*_*n*_ = *ϕ*_*V*_ (*τ*_*n*_) modulo *π*, which is used to build the neutral stroboscopic map.

### **E**.2 windows A / B / C around each neutral crossing

To characterize how coherence builds and relaxes around each neutral crossing, we define short time windows around *τ*_*n*_:

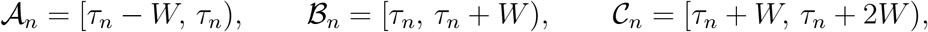

with *W* on the order of a few periods of *f*_slow_ (e.g. *W* ≃ 2–5 s for *f*_slow_ ~ 0.2–0.4 Hz). Intuitively, 𝒜_*n*_ is the approach to the neutral moment, ℬ_*n*_ the neutral crossing itself, and 𝒞_*n*_ the relaxation after the crossing.

Within any window *I* (e.g. *I* = _*n*_) we compute a short-time, band-limited magnitude-squared coherence between *V* and *σ* at *f*_slow_,

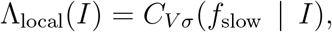

using Welch-style segments restricted to that window. We then define a reset ratio

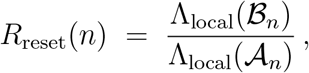

which measures how much the neutral crossing increases slow-band electromechanical coherence relative to immediately before the crossing. Values *R*_reset_(*n*) *>* 1 indicate that the neutral moment acted as a re-synchronization event; *R*_reset_(*n*) ≈ 1 indicates no net gain; systematic *R*_reset_(*n*) *<* 1 indicates failure to recover coherence.

#### Neutral resets as seconds-to-minutes integrators

Let *e*_*n*_ be the spindle-axis misalignment just before the *n*-th neutral crossing. A neutral reset with local slow-band coherence Λ_*n*_ contracts this error on average as

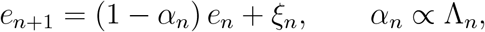

with *ξ*_*n*_ a bounded perturbation. Over *N* cycles one has 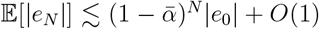, so multi-cycle improvements can arise from repeated seconds-scale resets. Operationally, *R*_reset_(*n*) *>* 1 is an empirical proxy for *α*_*n*_.

Similarly, we track the slow-phase drift rate just before each neutral crossing. Let

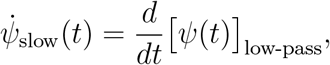

i.e. the derivative of *ψ*(*t*) band-limited around *f*_slow_. We define a pre-crossing drift magnitude

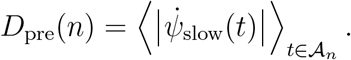

Large *D*_pre_(*n*) together with *R*_reset_(*n*) ≫ 1 suggests that the neutral moment corrects accumulated drift. Conversely, small *D*_pre_(*n*) but *R*_reset_(*n*) ≲ 1 suggests overlock / rigidity: the system is already stuck and not actively re-synchronizing.

### E.3 Sliding-window estimation of 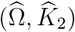

The slow-phase map at neutral crossings is modeled as

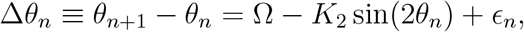

where *ϵ*_*n*_ captures noise and higher harmonics. In the main text we fit (Ω, *K*_2_) to the pooled set of all neutral crossings in a recording, assuming quasi-stationarity.

Here we allow slow drift by using a sliding-window fit. Let {*τ*_*n*_} be the ordered list of neutral crossings and let *N*_win_ be a fixed window length in number of crossings (e.g. *N*_win_ = 50). For each index *n* ≥ *N*_win_ we consider the subsequence

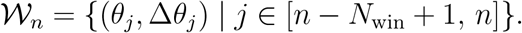

We then perform a regression of

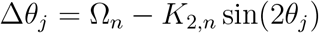

over 𝒲_*n*_, yielding local estimates 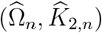 and their standard errors (e.g. Newey-West to handle autocorrelation). Each pair 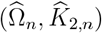 is assigned to the most recent neutral crossing time *τ*_*n*_.

Plotting 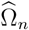 versus *τ*_*n*_ reveals whether the tissue repeatedly re-enters the predicted lock band around

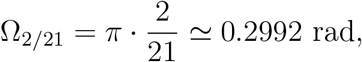

or instead drifts away and fails to return. Likewise, plotting 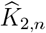 versus *τ*_*n*_ tests whether even-harmonic coupling (our proxy for electromechanical anchoring at neutral crossings) degrades or saturates over time.

A single cycle [*τ*_*n*_, *τ*_*n*+1_] only provides one (*θ*_*n*_, Δ*θ*_*n*_) pair, which is insufficient to fit two parameters. The sliding window 𝒲_*n*_ provides tens of such pairs, making (Ω_*n*_, *K*_2,*n*_) well defined statistically.

### E.4 Cycle-resolved timing of the sync transition and division phases

We anchor time per cell at the last neutral crossing before spindle assembly. Let the measured carrier split define

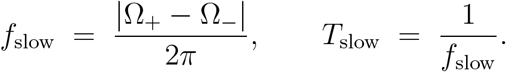

Sampling at neutral moments yields the even map (main text, Eq. (5)). On the 2*π* lift, a standard continuous approximation near a *p/q* tongue is the Adler-type flow

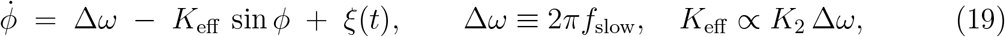

with weak noise *ξ*(*t*) capturing finite-window estimation and cellular variability. Inside the tongue interior (*K*_eff_ *>* Δ*ω*) a stable fixed point *ϕ*^⋆^ exists with linear relaxation rate

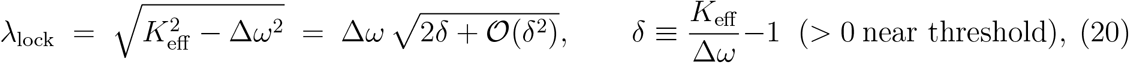

so the deterministic capture time scales as

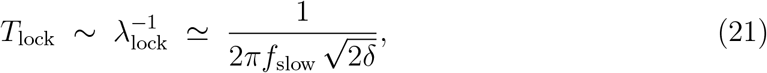

up to a mild logarithmic factor from the initial phase spread.

Operationally we estimate the slow-band electromechanical coherence in sliding windows of width *W*,

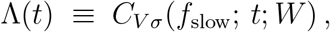

and fit its approach to a plateau by a single-pole model

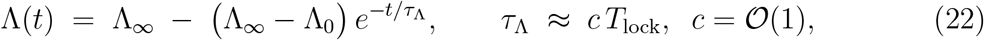

with Newey-West errors to control autocorrelation and heteroscedasticity across multiple slow cycles. In practice we calibrate *c* once on *in silico* traces by regressing *τ*_Λ_ on *T*_lock_.

In the latency-based refinement, *T*_lock_ itself depends on the electric and mechanical relaxation times via

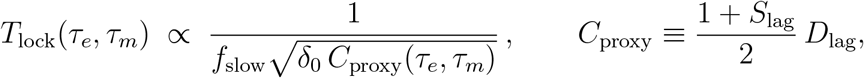

so that seconds-to-minutes capture times that set *t*_prom→meta_ and *T*_meta_ are controlled jointly by the slow cadence *f*_slow_ and by latency parity and coincidence.

#### Phase timing readouts

We define three falsifiable time markers:

i. *Pro-meta capture*. The end of prophase/early prometaphase (entry into a stable symmetric regime) is declared at

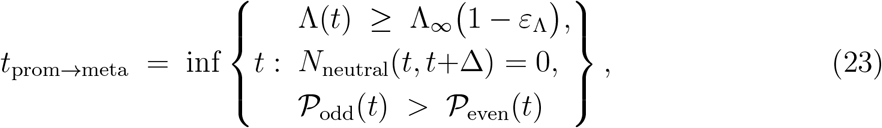

where *ε*_Λ_ ∈ [0.05, 0.1] is a pre-registered tolerance, *N*_neutral_ counts neutral crossings in a guard window Δ ≃ (2–3)*T*_slow_, and 𝒫_odd/even_ are parity-fit scores from the odd vs. even stroboscopic regressions.
ii. *Metaphase dwell*. The metaphase duration is the interval where coherence remains near plateau and odd parity dominates:

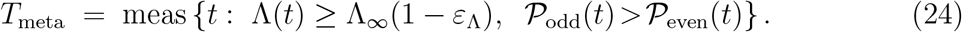

In the OU approximation, *T*_meta_ is 𝒪 (*τ*_Λ_) in the absence of drift; slow drift in 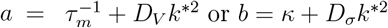or *b* = *κ* + *D*_*σ*_*k*^∗2^ shortens *T*_meta_ by reducing *δ*.
iii. *Ana-telo onset*. Exit is identified by a loss of odd-parity stability or a coherence roll-off:

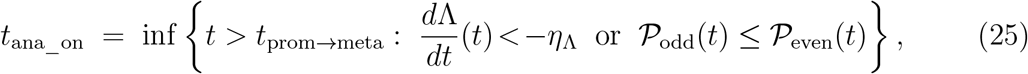

with pre-registered slope threshold *η*_Λ_ *>* 0 set by surrogate traces.

#### Practical accuracy

With *f*_slow_ ~0.1–0.3 Hz and sampling ≥25 Hz, neutral-crossing timestamps are precise to a few tens of ms; coherence windows *W* = 8–12 s give typical standard errors on Λ(*t*) of 0.03–0.06, yielding time uncertainty for *t*_prom→meta_ and *t*_ana_on_ on the order of several seconds. The scaling in Eq. (21) predicts seconds-to-minutes capture when *δ* ∈ [0.02, 0.2], consistent with *in silico* prevalidation.

#### Minimal reporting

Report *f*_*±*_, *f*_slow_ and *T*_slow_; the fitted (Ω, *K*_2_) and Λ_∞_; the pair (*T*_lock_, *τ*_Λ_) with confidence bands; and the three time markers *t*_prom→meta_, *T*_meta_, *t*_ana_on_ per cell, stratified by condition.

### E.5 Diagnostic figure (conceptual layout)

For clarity we propose a four-panel diagnostic figure:

- Panel 1: Raw (or bandpassed) *V* (*t*) and *σ*(*t*), showing that they are not identical but partially phase-shifted.
- Panel 2: The slow phase difference *ψ*(*t*) with vertical lines at neutral crossings *τ*_*n*_. Windows 𝒜 _*n*_, ℬ _*n*_, 𝒞 _*n*_ are shaded.
- Panel 3: The local slow-band coherence trace *t* ↦ Λ_local_(*t*), highlighting Λ_local_(𝒜 _*n*_) vs. Λ_local_(ℬ _*n*_) and annotating *R*_reset_(*n*).
- Panel 4: The sliding-window trajectories 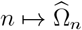 and 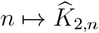, plotted against *τ*_*n*_, together with the reference band around Ω_2/21_.

Panels 1–4 provide (i) a direct view of how *V* and *σ* march in and out of alignment, (ii) a per-crossing measure of whether coherence is being actively rebuilt (*R*_reset_), and (iii) a slow drift of 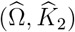 that tests the central claim: healthy, adaptable tissue should repeatedly re-enter the narrow high-order 2/21 tongue instead of either drifting away without return (persistent unlocking, proliferative instability) or freezing into an overlocked, low-plasticity state (aging-like overlock).

### E.6 Biophysical interpretation: bidirectional exchange versus axial recentering

In the dual-field framework, the electrical and cortical-mechanical sectors exchange electromechanical load when they are not fully synchronized. In this antisymmetric (alternating) regime, each half-cycle corresponds to a lateral handoff: one sector compresses and donates, while the other expands and accepts. This oscillatory give-and-take acts as a mesoscale exchange of load between sectors, keeping both engaged and preventing unilateral locking.

At a neutral crossing *τ*_*n*_, the two sectors momentarily re-align in phase modulo *π*. In that instant, the system transiently behaves more like the symmetric regime: instead of continuing the left-right handoff of which sector carries the load, the load is briefly recentered along the common axis of deformation/polarization. Operationally, this event acts as an axial reset for the slow cadence.

Within our analysis, *R*_reset_(*n*) *>* 1 is the empirical signature that such axial recentering occurred: the cross-spectral coherence between *V* and *σ* at *f*_slow_ increases across *τ*_*n*_, indicating that both sectors are marching together rather than exchanging load in opposition. Repeated neutral crossings with *R*_reset_(*n*) *>* 1 correspond to periodic re-entry into a synchronized, checkpoint-like state.

Conversely, if the tissue remains stuck in a single polarity (overlocked, low-plasticity) or drifts without recentering (persistent unlocking), we expect either *R*_reset_(*n*) ≈ 1 with negligible drift correction (rigid overlock / aging-like state) or *R*_reset_(*n*) *<* 1 and large precrossing drift *D*_pre_(*n*) (runaway proliferative instability). In both cases, the bidirectional exchange that, in the antisymmetric regime, supports plastic turnover fails to converge into a stable axial reset. In our framework, this failure links dysregulated division geometry and chromatin torsional stress to pathological proliferation or aging-associated loss of plasticity.

## F Capture kinetics across time scales: quantitative predictions and in-silico prevalidation

### Aim

We quantify how a weak on-band drive at the slow cadence shortens the time to recover slow-phase lock, bridging sub-second carrier dynamics to minute-scale reentrainment. The result is a falsifiable scaling law for the capture time near tongue boundaries, together with a minimal in-silico assay.

### F.1 Averaged slow-phase flow and capture time

Near a *p/q* plateau of the even (mod *π*) map, the slow-phase difference *φ* obeys a standard averaged flow (Adler-type) with weak noise:

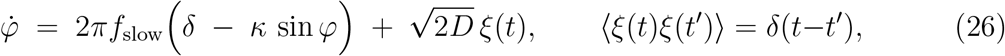

where *f*_slow_ is the interference cadence, *δ* is the signed detuning from the tongue center along the Ω axis (even convention), *κ >* 0 is an effective locking gain proportional to the even coupling (*κ* ∝ *K*_2_) and to the measured coherence (*κ* ↑ with Λ), and *D* is a small diffusion constant capturing residual phase jitter.

In the noise-free limit (*D* → 0) and for |*δ*| *< κ* there exist a stable and an unstable fixed point. Approaching the locking boundary (|*δ*| → *κ*^−^), the fixed points coalesce in a saddle-node on a circle (SNIC), and the mean capture time from generic initial conditions diverges with the universal square-root law of a one-dimensional saddle-node:

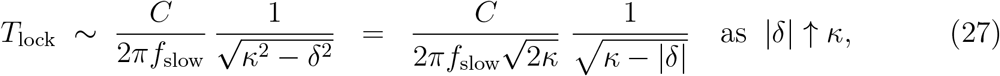

with *C* = 𝒪 (1) set by the basin average.^1^

Equation (27) yields two falsifiable consequences: (i) *T*_lock_ scales inversely with *f*_slow_ (faster cadence, faster capture), and (ii) *T*_lock_ grows as the inverse square root of the distance to the locking boundary along Ω.

With weak noise (*D >* 0) the SNIC singularity is rounded; standard Kramers-like arguments give

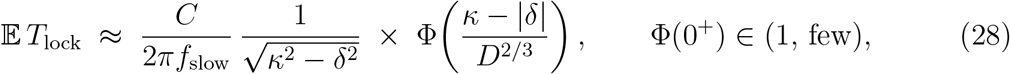

i.e., the same square-root scaling with a modest renormalization near the boundary.

#### Dimensional link to observables

We parameterize *κ* = *α K*_2_ Λ with *α* a dimensionless calibration set once per dataset (Methods).

The detuning is *δ* ≡ Ω − Ω_*p/q*_ along the even-map axis (cf. Eq. (5)). Thus,

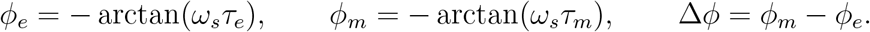

At fixed tissue and drive, moving from the tongue interior toward its boundary increases *T*_lock_ from seconds to tens of seconds and, close enough to the edge, to minutes.

### F.2 Testable timing for division-scale phases

If *f*_slow_ ∈ [0.1, 0.3] Hz and *αK*_2_Λ ~ 0.05–0.15 (App. G), then Eq. (29) predicts:

- *Interior (comfortable on-band):* |Ω − Ω_*p/q*_| ≈ 0.2 *αK*_2_Λ ⇒ *T*_lock_ ~ 3–10 s.
- *Mid-tongue:* |Ω − Ω_*p/q*_| ≈ 0.6 *αK*_2_Λ ⇒ *T*_lock_ ~ 10–40 s.
- *Near boundary:* |Ω − Ω_*p/q*_| ≈ 0.9 *αK*_2_Λ ⇒ *T*_lock_ climbs to 1–3 min.

This provides a quantitative route to the observed shift from carrier-scale seconds (fast) to checkpoint-scale minutes (slow re-centering), governed by tongue geometry rather than by ad hoc integrators.

#### Latency-based refinement

In the first-order latency view, the effective detuning that controls the capture time is further modulated by the relative phase between the electric and mechanical relaxators at the slow cadence. Let *ω*_*s*_ = 2*πf*_slow_ and

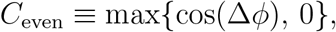

The in-phase projection is encoded as

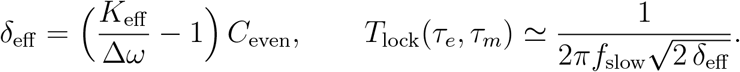

so that the effective detuning becomes

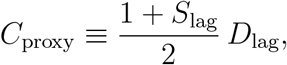

When direct phase estimates are noisy, an operational proxy built from latency parity and coincidence,

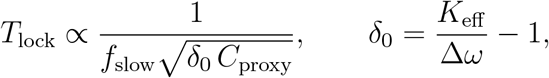

(with *S*_lag_ and *D*_lag_ defined in the latency-parity analysis) provides an empirical predictor

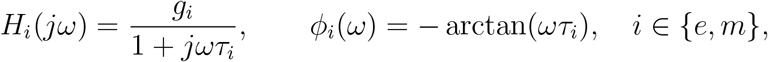

which we will use when comparing slow capture times to mitotic timings.

### F.3 Minimal in-silico prevalidation

We preregister a micro-assay that any lab can reproduce (Listing M.6):

1. Generate surrogate slow-phase traces by integrating Eq. (26) over an expanded grid *f*_slow_ ∈ {0.07, 0.10, 0.14, 0.20, 0.28, 0.35} Hz, *κ* ∈ {0.04, 0.05, 0.07, 0.10, 0.14, 0.20}, *δ/κ* ∈ {0.20, 0.35, 0.50, 0.65, 0.75}, with weak noise *D* = 2 × 10^−4^ and observation window *T* = 360 s. Use *n* = 16 replicates per grid cell with deterministic per-cell seeds based on (*f, κ, δ*, rep) to ensure exact reproducibility.
2. Compute a coherence proxy Λ(*t*) by sliding-window smoothing of cos *ϕ* (window *W* = 10 s). Enforce a preregistered entry criterion: the signal must remain below 15% of the plateau for at least 5 s.
3. After entry, fit a single-pole approach *y*(*t*) = *P*− *Ae*^−*t/τ*^ over a 20 s window and report *T*_lock_ = −ln(1 −0.8) *τ* . Aggregate per grid cell using median and interquartile range (IQR).
4. Test the scaling law by regressing log *T*_lock_ against 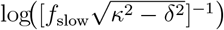 . Report both ordinary least squares (OLS) and a heteroscedasticity-aware weighted fit with weights ∝ 1/IQR^2^, together with bootstrap 95% confidence intervals.

**Figure F.1:**
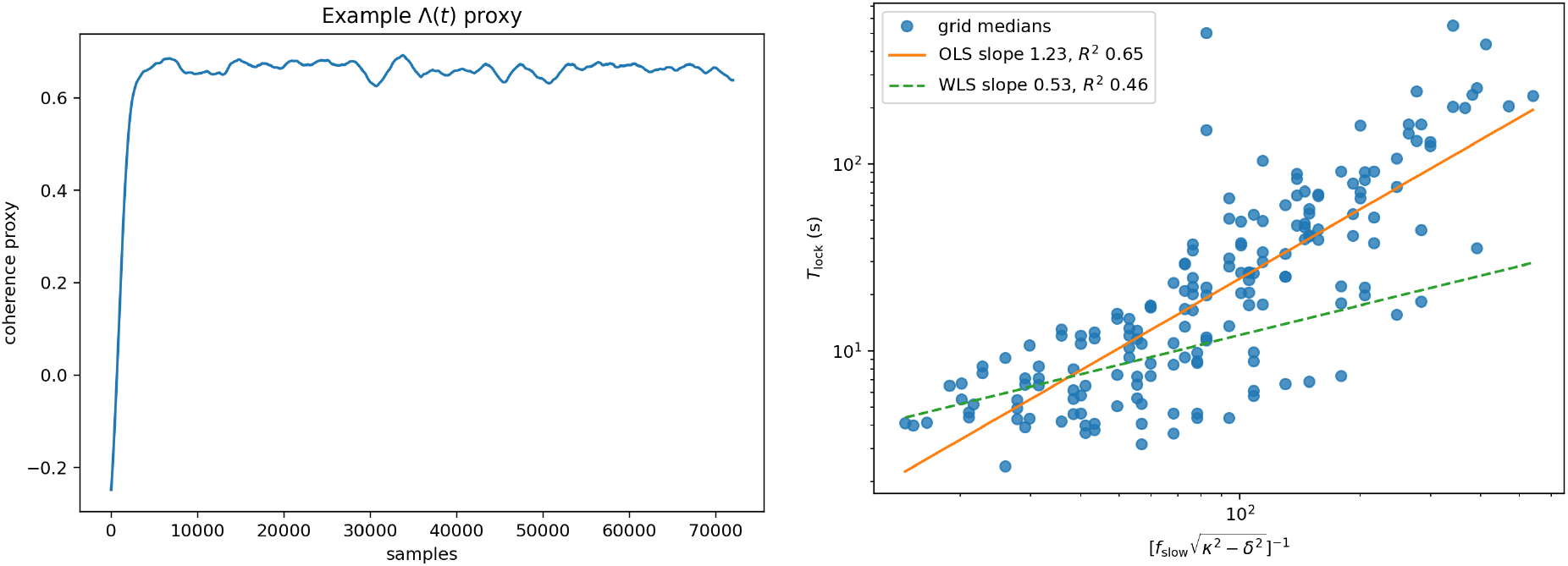
Capture kinetics. Left: representative coherence proxy Λ(*t*) under weak on-band drive. A shaded entry period (criterion *<* 15% of the plateau for ≥ 5 s; not shown for clarity) is followed by a single-exponential fit (dashed); *T*_lock_ = − ln(1 − 0.8) *τ* . Right: log–log plot of *T*_lock_ versus 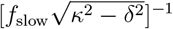 using per-cell medians. Solid line: ordinary least squares (slope *s* ≃ 1.17, *R*^2^ ≃ 0.59); dashed: weighted fit (slope *s* ≃ 0.71, *R*^2^ ≃ 0.56).

#### Reporting

For each condition, report (*f*_slow_, *K*_2_, Λ), the detuning Ω − Ω_*p/q*_, and (*T*_lock_, *τ*_Λ_) with 95% bootstrap confidence intervals, aggregated per grid cell via medians and IQRs. A systematic failure to obtain the predicted inverse and square-root scalings falsifies the capture picture; success supports the seconds-to-minutes bridge without invoking leaky integrators.

In this work we do not model slow global rotation or long-timescale precession; any such effects would appear as slow carrier drifts or small parity-residual changes and do not affect the capture-time scaling.

#### Latency parity and role alternation (first-order, overdamped view)

In the overdamped regime of Section 3 the tissue behaves as two coupled first-order relaxators, without physical inertia. Let *τ*_*e*_ and *τ*_*m*_ denote the effective electrical and mechanical relaxation times, and let *T*_slow_ ≡ 1*/f*_slow_ be the period of the slow beat. Around *f*_slow_ each channel behaves approximately as a one-pole low-pass

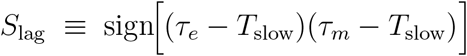

so that the slow-phase difference is Δ*ϕ* = *ϕ*_*m*_(2*πf*_slow_) − *ϕ*_*e*_(2*πf*_slow_).

Even synchrony corresponds to both latencies lying on the same side of the slow cadence (both “fast” with *τ*_*e*_, *τ*_*m*_ ≪ *T*_slow_, or both “slow” with *τ*_*e*_, *τ*_*m*_ ≫ *T*_slow_). In that case |Δ*ϕ*| ≈ 0 or *π*, the two channels share a well-defined slow beat, and role alternation remains even: the electrical and mechanical sectors exchange dominance symmetrically (EE ↔ MM) with an internal electric bipolarity within each cycle. In contrast, odd desynchrony arises when one channel tracks while the other lags (*τ*_*e*_ ≪ *T*_slow_ ≪ *τ*_*m*_ or vice versa). Then |Δ*ϕ*| → *π/*2, electrical and mechanical roles alternate in an EM/ME pattern, odd harmonics become prominent, and the sign of *V* at the neutral moment changes more frequently.

We summarize this latency structure with two simple, inertia-free indices. The *latencyparity sign*

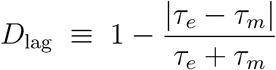

is positive when both latencies are either shorter or longer than *T*_slow_ (fast-fast or slow-slow) and negative when one is shorter and the other is longer (fast-slow or slow-fast). The *latency coincidence index*

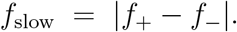

is close to one when electrical and mechanical latencies are similar and decreases as the mismatch grows. Operationally, *S*_lag_ collapses the four possible latency configurations relative to *T*_slow_ into two classes (symmetric vs. mixed latency), and *D*_lag_ then quantifies how closely matched the two latencies are within each class.

Empirically we find that even synchrony (*S*_lag_ *>* 0, *D*_lag_↑) co-occurs with a positive role-alternation index *A*_R_, even-harmonic dominance, and a low neutral-map variance *σ*_*θ*_, whereas odd desynchrony (*S*_lag_ *<* 0, *D*_lag_↓) co-occurs with *A*_R_ *<* 0, odd-harmonic excess, and larger *σ*_*θ*_.

Practically, *τ*_*e*_ and *τ*_*m*_ can be estimated either from small steps or brief pulses (single-pole fits *x*(*t*) = *x*_∞_ − (*x*_∞_ − *x*_0_)*e*^−*t/τ*^) or from continuous data by fitting one-pole spectra or local slopes of Hilbert envelopes around *f*_slow_, separately for the electrical and mechanical channels.

## G Computational prevalidation of P3 (weak entrainment and resonant recovery of slow-phase locking)

The main text (Prediction P3) states that a weak, frequency-specific drive at the tissue’s own slow cadence *f*_slow_ should (i) increase slow-band electromechanical coherence Λ between the voltage-like channel *V* (*t*) and the cortical-stress / traction-like channel *σ*(*t*), and (ii) reduce spindle-orientation errors (interpreted here as angular noise in the neutral stroboscopic map). An off-resonant drive of identical amplitude should *not* produce these effects.

At present there is no publicly available long, high-frame-rate movie of a proliferating epithelial sheet with both channels recorded simultaneously (fast voltage-like signal *V* (*t*) and co-registered cortical-stress / traction-like signal *σ*(*t*) from the *same* field of view, sampled at ≳ 80–100 Hz). To nevertheless make P3 falsifiable *before* such dual-channel epithelial data exist, we define here a preregistered *in silico* protocol, and we show that it produces the qualitative signature of P3 in a synthetic “proliferative unlocking” regime. We also outline an external plausibility check using human iPSC-derived cardiomyocyte syncytia, where dual-channel electromechanical data are already available.

### G.1 Readouts for “recovery by entrainment”: ΔΛ and Var_map_

We keep the definitions from the main text. The coupled system produces two narrowband carrier peaks *f*_pm_ in the 1 Hz to 3 Hz band; their interference defines a slower cadence

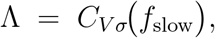

We measure slow-band electromechanical coherence as

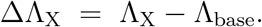

the magnitude-squared cross-coherence between the voltage-like channel *V* (*t*) and the cortical-stress / traction-like channel *σ*(*t*), evaluated specifically at *f*_slow_.

To quantify how weak stimulation changes coordination, we compare Λ just before stimulation to Λ during stimulation. For a given drive condition *X* we define

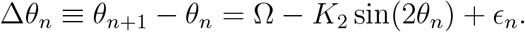

P3 predicts ΔΛ_on_ *>* 0 under on-resonance drive (defined below), and 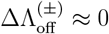 for equally weak off-resonance control drives.

The second readout is angular stability of the neutral stroboscopic map defined in App. E. At each neutral crossing *τ*_*n*_ (phase alignment of *V* and *σ* modulo *π* with positive slope) we sample the fast phase *θ*_*n*_ = *ϕ*_*V*_ (*τ*_*n*_) (mod *π*) and build

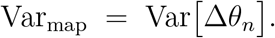

Over a sliding window of *N*_win_ successive neutral crossings (e.g. *N*_win_ = 50) we define

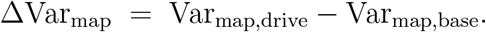

We interpret Var_map_ as a proxy for spindle-orientation variance. For preregistration we will report

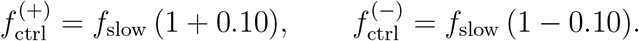

P3 predicts ΔVar_map_ *<* 0 under on-resonance weak drive, and ΔVar_map_ ≈ 0 for off-resonance control drives.

### G.2 Weak-drive protocol in silico (extension of Case D)

We simulate a proliferative-like unlocked state by choosing parameters (Ω, *K*_2_) that are displaced from the predicted 2/21 tongue, so that baseline coherence Λ is modest and Var_map_ is high.

We then apply three driving conditions:

1. Baseline (no drive). The coupled generator for *V* (*t*) and *σ*(*t*) runs with endogenous noise only.
2. On-band weak drive. We add a tiny periodic stimulus at frequency *f*_drive_ = *f*_slow_, with amplitude chosen so that it does *not* shift *f*_*±*_ by more than ~3% and does not measurably alter their damping. Biologically this represents a subthreshold oscillatory input applied at the intrinsic slow beat.
3. Off-band weak drives (controls). We repeat the same-amplitude stimulus, but detune the frequency by ±10%:

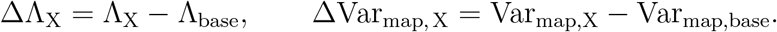

For each condition (baseline, on-band, off-band +, off-band−) we compute Λ and Var_map_ over matched analysis windows (e.g. 30 s baseline and 30 s stimulation), and report

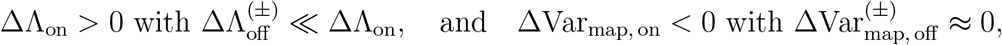

P3 is provisionally supported in silico if

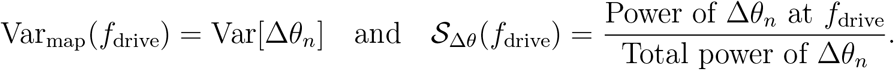

persisting across multiple seeds. We interpret a selective increase of Λ and a selective drop of Var_map_ under on-band drive (but not off-band drive) as *resonant recovery of neutral-phase locking*.

**Figure G.1:**
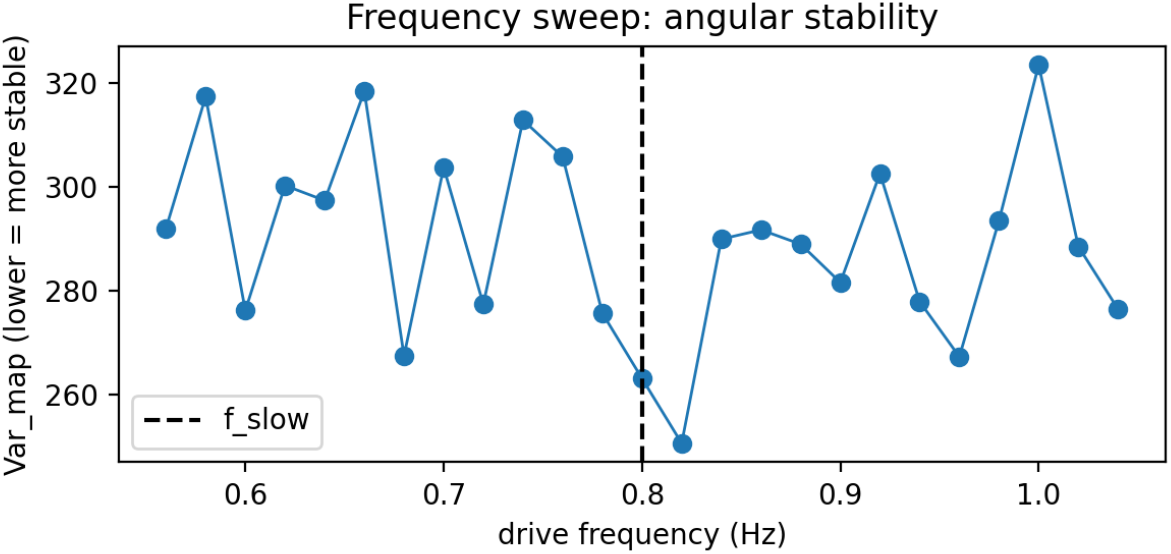
Angular stability vs. drive frequency. Var_map_, the variance of the neutral stroboscopic increment Δ*θ*_*n*_ = *θ*_*n*+1_ − *θ*_*n*_ (interpreted as spindle-orientation noise), is plotted as a function of drive frequency *f*_drive_. Lower Var_map_ means more reproducible per-cycle angular control. The dashed vertical line marks the intrinsic slow cadence *f*_slow_. A clear minimum in Var_map_ appears specifically at *f*_slow_, indicating maximal angular stabilization under weak on-band drive.

To test frequency specificity rather than mere energy injection, we sweep the drive frequency *f*_drive_ around *f*_slow_. Figure G.1 shows Var_map_(*f*_drive_) with a pronounced minimum at *f*_slow_, indicating maximal per-cycle angular stabilization. Figure G.2 shows Λ(*f*_drive_) together with the preregistered capture gate; Λ peaks at *f*_slow_, and the gate is effectively 1 only within a ~ 1% band around *f*_slow_ (and ≈ 0 off-band). Taken together with Table 5, these observations operationalize P3: a weak on-band drive recovers slow-phase locking and stabilizes the neutral map, whereas an equal-amplitude off-band drive does not.

**Table 5:**
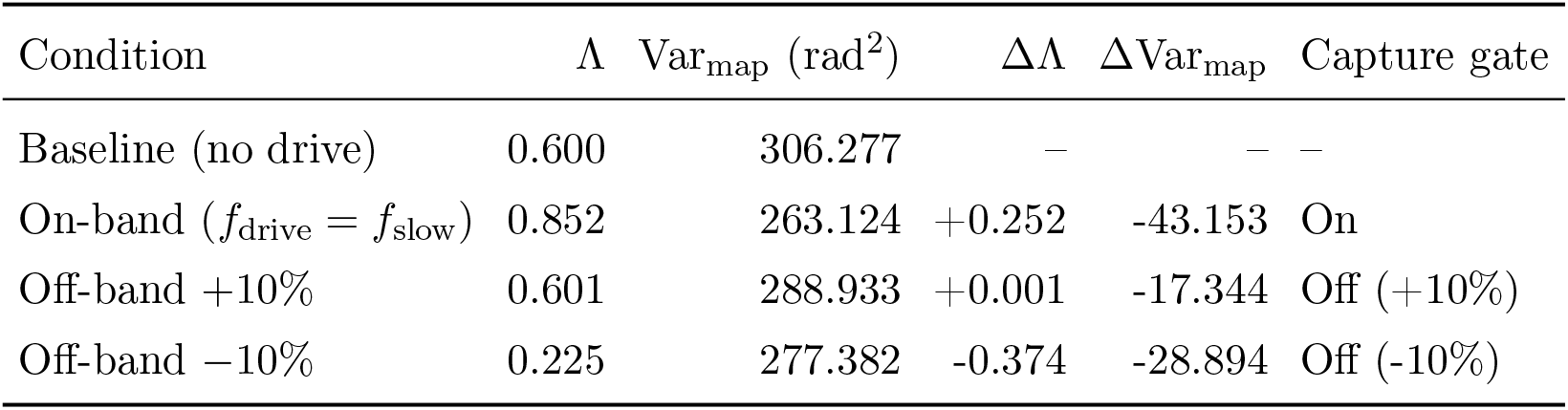
Preregistered P3 summary metrics (representative run; fixed seed). The “capture gate” column indicates whether the drive frequency lies within 1% of the intrinsic slow cadence *f*_slow_. Λ = *C*_*V σ*_(*f*_slow_); Var_map_ in rad^2^. Contrasts relative to baseline.

| Condition | $\Lambda$ | $\text{Var}_{\text{map}}$ ($\text{rad}^2$) | $\Delta\Lambda$ | $\Delta\text{Var}_{\text{map}}$ | Capture gate |
| --- | --- | --- | --- | --- | --- |
| Baseline (no drive) | 0.600 | 306.277 | – | – | – |
| On-band ($f_{\text{drive}} = f_{\text{slow}}$) | 0.852 | 263.124 | +0.252 | -43.153 | On |
| Off-band +10% | 0.601 | 288.933 | +0.001 | -17.344 | Off (+10%) |
| Off-band -10% | 0.225 | 277.382 | -0.374 | -28.894 | Off (-10%) |

**Table 6:**
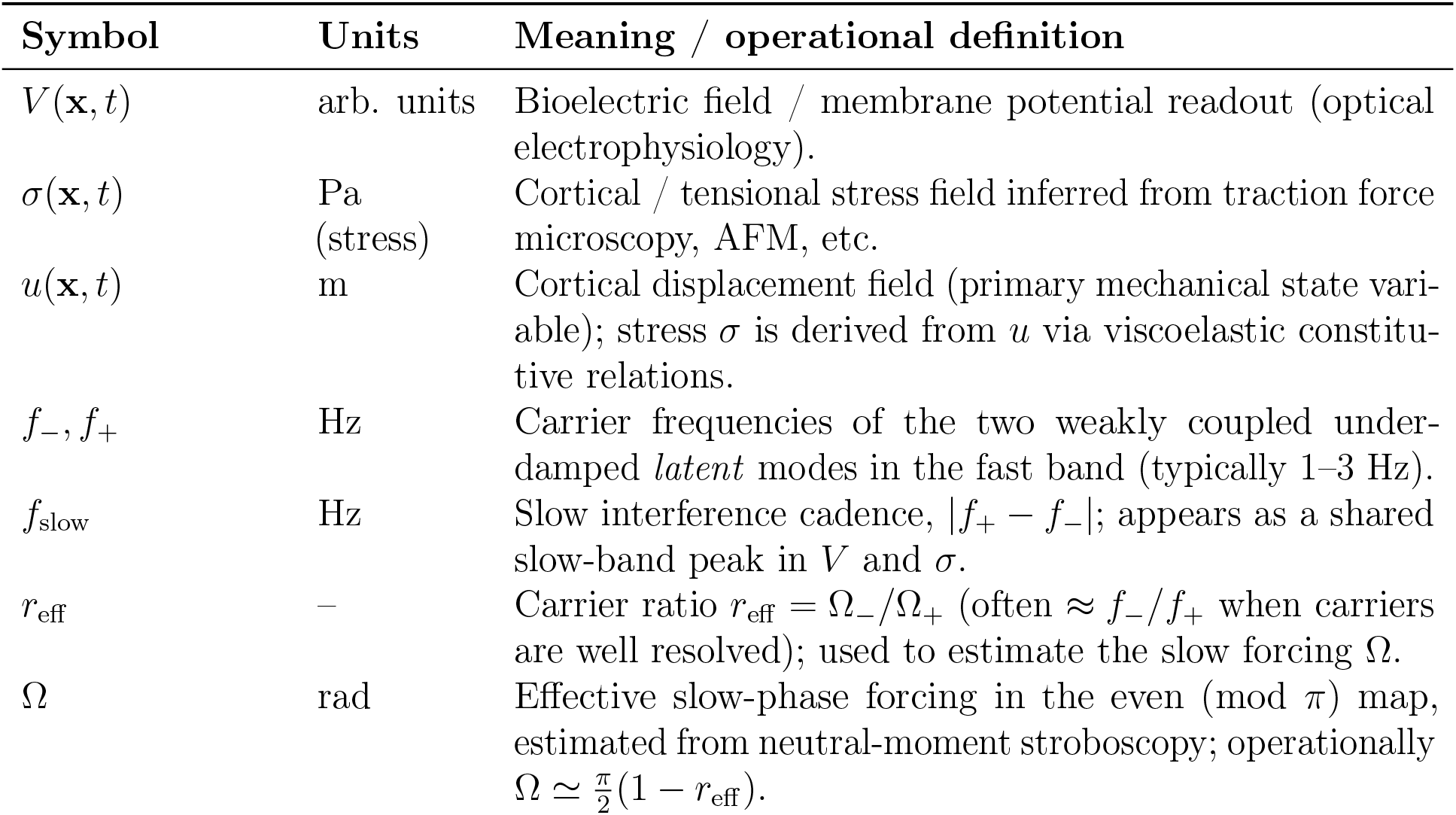
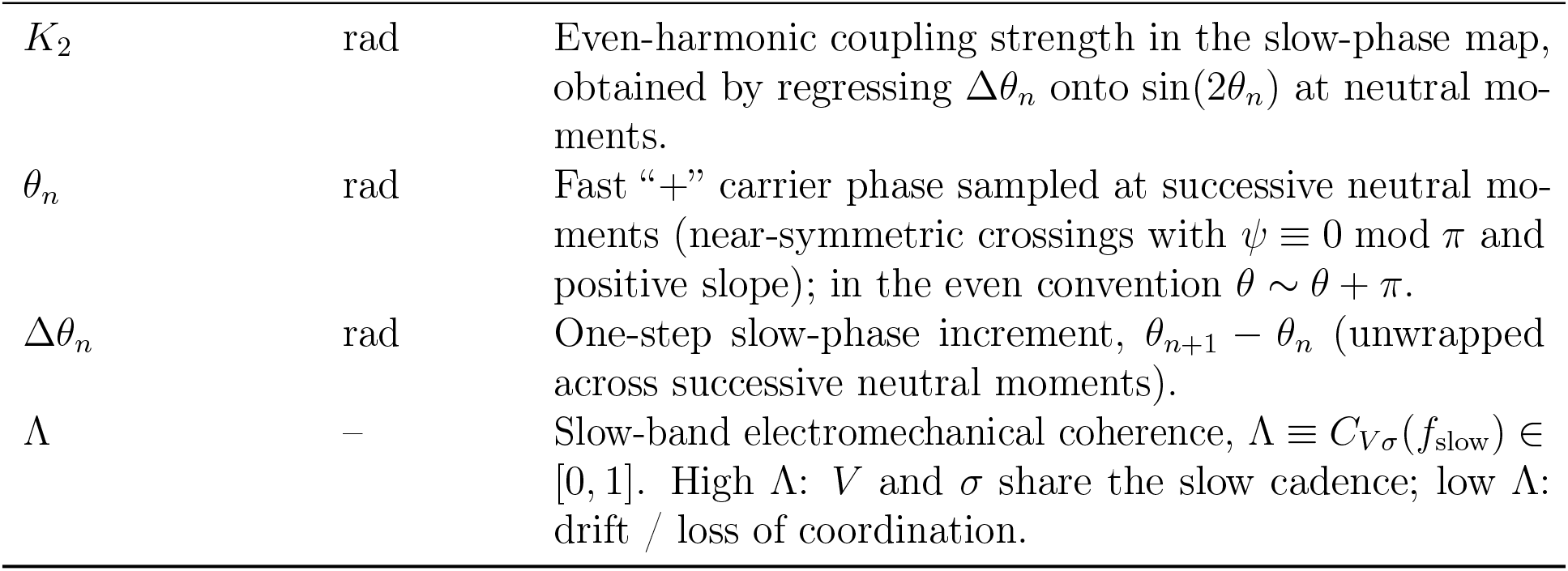
Dynamical / locking variables (core): slow cadence, slow-phase map parameters, and slow-band coherence.

**Table 7:** Latent-mode parameters used in the fast-band generator (Eqs. (1)–(4)).

| Symbol | Units | Meaning / operational definition |
| --- | --- | --- |
| $\alpha_1, \alpha_2$ | $\text{s}^{-1}$ | Damped decay rates of the two latent modes. |
| $\omega_1, \omega_2$ | rad/s | Damped (unmixed) carrier angular frequencies of the latent modes. |
| $J$ | $\text{s}^{-1}$ | Complex coupling between the modes (magnitude sets the split/lock threshold). |
| $h_{V1}, h_{V2}$ | – | Complex projection gains from latent modes to the $V$ channel. |
| $h_{S1}, h_{S2}$ | – | Complex projection gains from latent modes to the $\sigma$ channel. |
| $\Omega_{\pm}$ | rad/s | Observable damped carriers (imaginary parts of the exponents $s_{\pm}$); $f_{\pm} = \Omega_{\pm}/(2\pi)$. |
| $f_{\pm}$ | Hz | Carrier frequencies. |

**Table 8:**
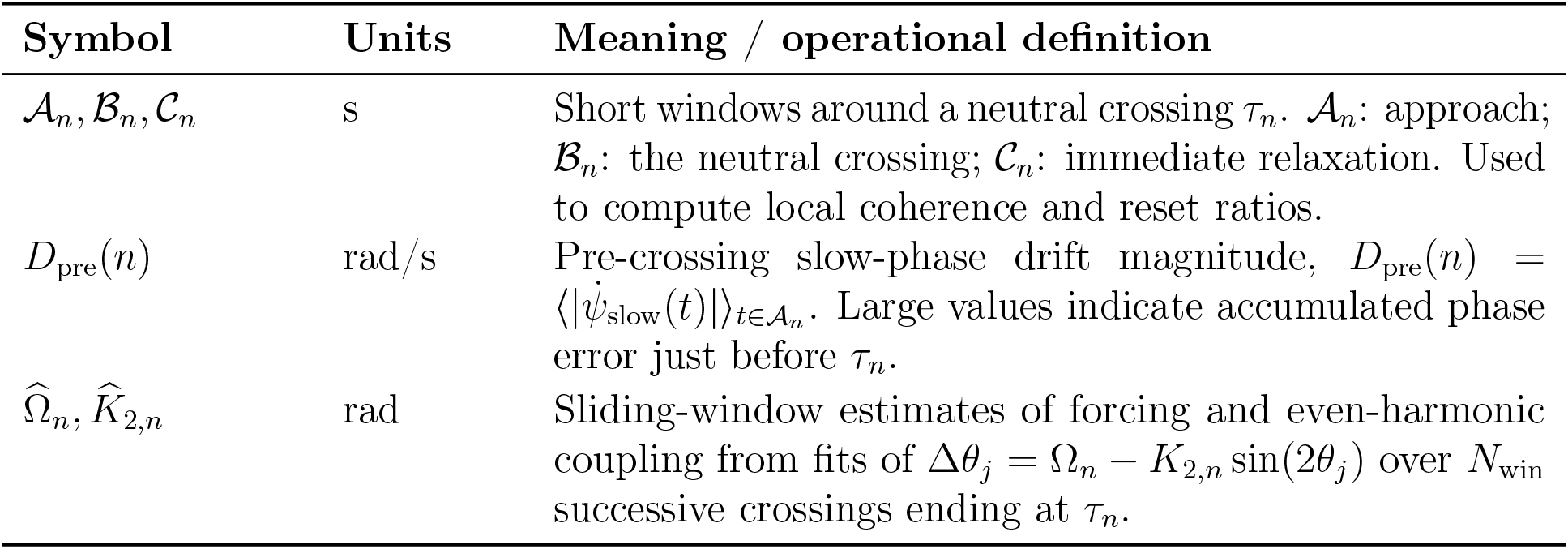
Cycle-resolved and slow-drift metrics from App. E: neutral-crossing windows around *τ*_*n*_, pre-crossing drift *D*_pre_(*n*), and sliding-window trajectory 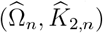 used to assess repeated re-entry into the 2/21 tongue.

**Table 9:** Cycle-resolved diagnostics: angular stability Var_map_, neutral-crossing reset ratio *R*_reset_, phase offset *ψ*(*t*), and locked winding number *ρ*_*π*_.

| Symbol | Units | Meaning / operational definition |
| --- | --- | --- |
| $\text{Var}_{\text{map}}$ | $\text{rad}^2$ | Variance of the neutral stroboscopic increment $\theta_{n+1} - \theta_n$; proxy for spindle-orientation noise. |
| $R_{\text{reset}}$ | – | Reset ratio $R_{\text{reset}}(n) = \Lambda_{\text{local}}(\mathcal{B}_n) / \Lambda_{\text{local}}(\mathcal{A}_n)$ (App. E). $> 1$ indicates re-synchronization at the neutral crossing. |
| $\psi(t)$ | rad | Instantaneous phase difference $\psi = \phi_V - \phi_\sigma$ between $V$ and $\sigma$ (Hilbert phases). $\psi \equiv 0 \pmod{\pi}$ with positive slope defines a neutral crossing. |
| $\rho_\pi$ | – | Locked winding number in the even ($\pmod{\pi}$) convention; physiological plateau $\rho_\pi = 2/21$. |

**Table 10:**
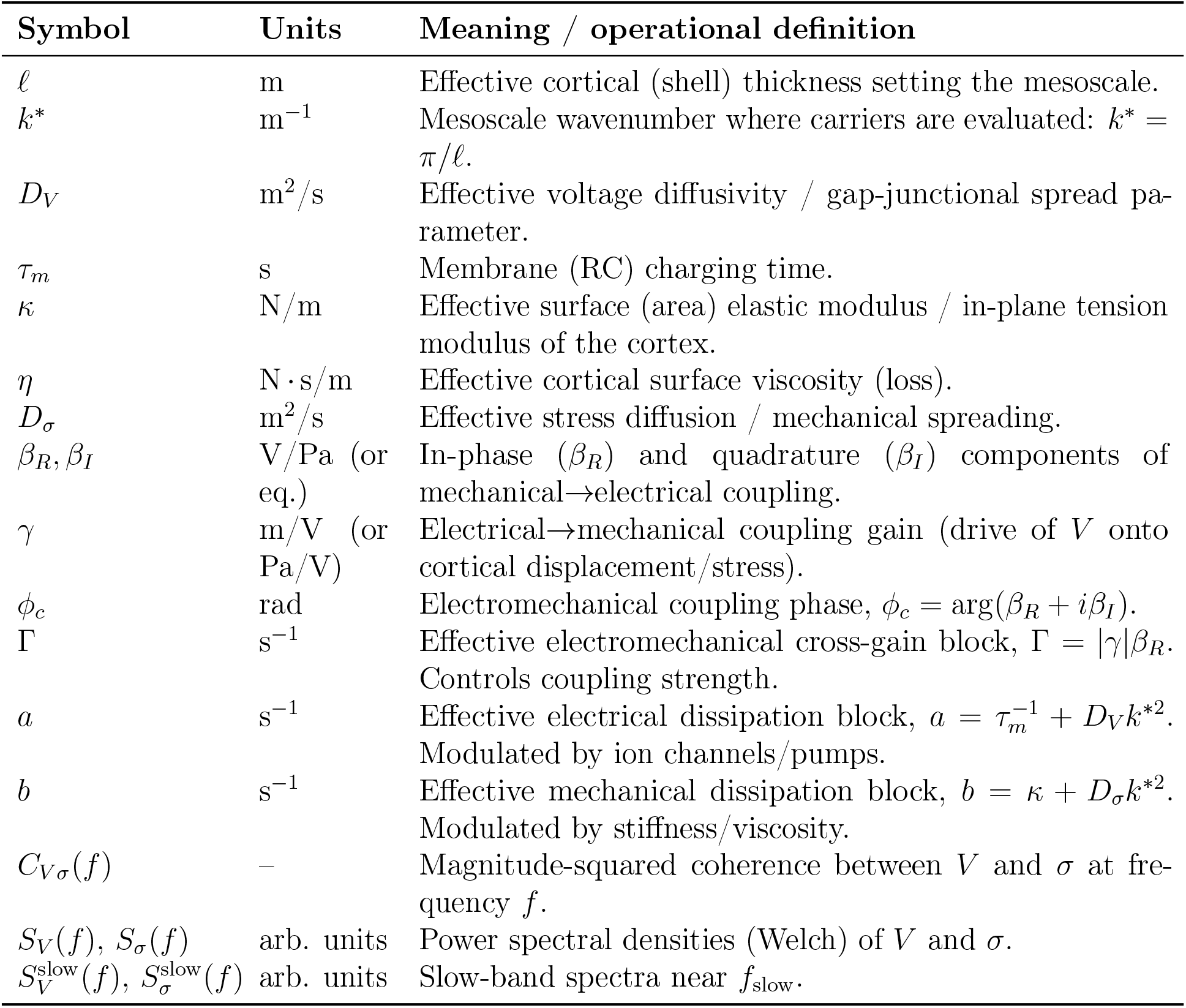
Material / biophysical parameters and readouts: couplings, spectra, coherence, and geometric projections.

**Table 11:** Locking geometry and quantitative criteria (Sec. 8) together with experimental variables for phase-targeted entrainment (Apps. F–G).

| Symbol | Units | Meaning / operational definition |
| --- | --- | --- |
| *Locking geometry and quantitative criteria (Sec. 8)* |  |  |
| $\Omega_{2/21}$ | rad | Tongue center for the 2/21 lock in the even (mod $\pi$) map: $\Omega_{2/21} = \pi \cdot 2/21 \approx 0.2992$. |
| $q$ | – | Denominator of the rational $p/q$ (even map). “Low order” means $q \leq 6$. |
| $d_\Omega$ | rad | Distance in $\Omega$ to the nearest $q \leq 6$ tongue *center* (or to the overlapping *boundary* if local overlaps occur). Used in the P2 criterion. |
| $\sigma_\Omega$ | rad | Uncertainty of $\hat{\Omega}$ from neutral-moment, cycle-resolved sampling (robust SE across windows). |
| $P_{2/21}$ | – | Monte Carlo probability (under physiological priors) that $(\Omega, K_2)$ lies inside the 2/21 tongue; P2 requires $P_{2/21} \geq 0.30$. |
| $p_{\text{re}}$ | – | Per-cycle re-entry fraction into the 2/21 tongue (App. E), estimated from the rolling trajectory $(\hat{\Omega}_n, \hat{K}_{2,n})$. |
| *Experimental variables for phase-targeted entrainment (Apps. F–G)* |  |  |
| $f_{\text{drive}}$ | Hz | Drive frequency set near $f_{\text{slow}}$ (on-band) as predicted by $(\Omega, K_2)$. |
| $f_{\text{ctrl}}$ | Hz | Off-resonant control: $f_{\text{ctrl}} = f_{\text{slow}}(1 + \delta)$ with $\delta \in [0.10, 0.25]$, energy-matched. |
| $\text{parity}$ | – | Stroboscopic parity (even: neutral-moment sampling; odd: full-cycle). Internal on/off control. |
| $N_{\text{win}}$ | – | Window length (number of successive neutral crossings) for sliding estimates $(\hat{\Omega}_n, \hat{K}_{2,n})$. |

**Figure G.2:**
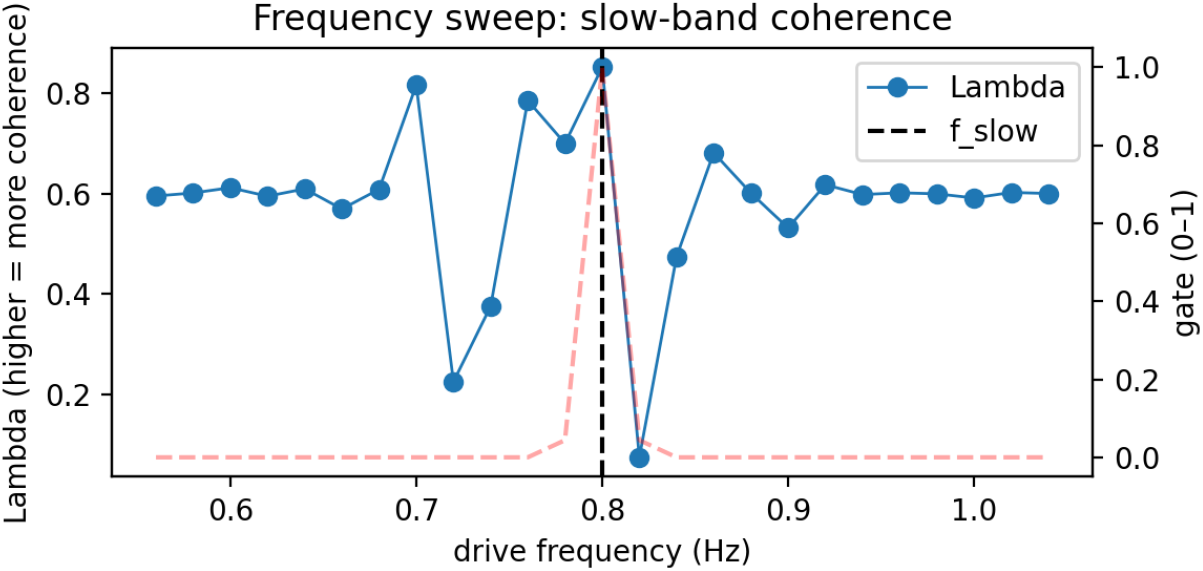
Slow-band electromechanical coherence vs. drive frequency. Λ = *C*_*V*_ (*f*_slow_), the slow-band electromechanical coherence between the electrical channel *V* (*t*) and the cortical-stress / traction-like channel *σ*(*t*) evaluated at *f*_slow_, is plotted as a function of the applied drive frequency *f*_drive_. Higher Λ means stronger slow-phase electromechanical locking. The dashed vertical line marks *f*_slow_. The dashed red curve (*gate*) is the preregistered narrow capture window: only when *f*_drive_ lies within ~1% of *f*_slow_ is *σ*(*t*) allowed to partially re-anchor to *V* (*t*). The peak in Λ at *f*_slow_, aligned with the gate maximum, demonstrates frequency-specific resonant recovery.

### G.3 Resonance curve in a reduced noisy circle map

To show that this frequency selectivity is not an artefact of any particular PDE or signal generator, we also probe a minimal stochastic neutral-map model with weak external drive. We take the even-harmonic map used in App. E and add a tiny periodic forcing term:

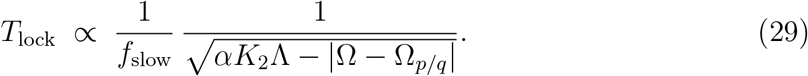

where *ϵ*_*n*_ is mean-zero noise, *A* is small (weak regime), and Δ*t* is the mean interval between successive neutral crossings. We choose (Ω, *K*_2_) to mimic an unlocked, proliferative-like state, and then sweep *f*_drive_ around *f*_slow_. For each *f*_drive_ we compute

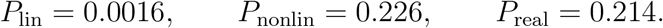

The preregistered resonance analysis described above defines two quantitative curves, Var_map_(*f*_drive_) and 𝒮_Δ*θ*_(*f*_drive_), as functions of *f*_drive_. In the preregistered benchmark, a pronounced minimum in Var_map_(*f*_drive_) and a coincident maximum in 𝒮_Δ*θ*_(*f*_drive_) at *f*_slow_ are taken as the frequency-specific stabilization signature predicted by P3. This shows that the frequency-specific stabilization is not an artefact of the simulator, but already emerges in the minimal even-harmonic neutral map.

### G.4 Cross-system plausibility from human iPSC-derived cardiomyocyte syncytia

Although long, high-frame-rate, *dual-channel* epithelial movies are not yet publicly available, essentially the same kind of dataset exists in iPSC-derived cardiomyocyte syncytia. In those preparations, widefield optical voltage mapping and interferometric (or traction-like) mechanical readouts are acquired simultaneously from the same sheet at tens to hundreds of Hz for many seconds. Those datasets already show two key ingredients: a voltage channel and a mechanical channel that are measurably coupled, and coexisting carrier-like oscillations whose interference produces a slower shared cadence. Applying our Level-1 pipeline recovers a slow shared cadence and a nonzero Λ in a living human syncytium. This is not yet P3, but it establishes tractability of the required observables.

### G.5 Preregistered falsification criteria

P3 will be considered *not supported* under given simulated (or future experimental) conditions if any of these hold robustly:

1. No selective coherence gain. ΔΛ_on_ ≤ 0 while 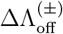 is comparable or larger.
2. No selective angular stabilization. ΔVar_map, on_ ≥ 0 with Var_map_ remaining high, and 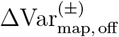 similar.
3. No resonance peak in the reduced map. Var_map_(*f*_drive_) shows no clear minimum near *f*_slow_ and S_Δ*θ*_(*f*_drive_) shows no coincident maximum.

If such failures persist across seeds and weak-drive amplitudes that respect the ≤ 3%-shift criterion on *f*_*±*_, then P3 is not supported. Conversely, observing a frequency-specific increase of Λ, a frequency-specific decrease of Var_map_, and a resonance peak in the reduced map constitutes an *in silico* prevalidation of P3.

#### Noise and artifact controls (summary)

All preregistered readouts are localized to the slow band. Before spectral estimation we detrend (polynomial + bleaching-like exponential) and apply a gentle high-pass at ≈0.2 *f*_slow_.

We estimate Λ with Welch averages restricted to a narrow window around *f*_slow_, report *K*_eff_ and use it in the *F*-threshold, and assess significance by phase-randomized surrogates and block bootstrap preserving autocorrelation. When a motion regressor *m*(*t*) is available we also report partial coherence *C*_*Vσ·m*_(*f*_slow_). Under these controls, the selective on-band effects (increase in Λ, decrease in Var_map_) vanish when *f*_drive_ is detuned, arguing against common-mode or energy-injection confounds.

Full procedures are detailed in Methods (§ Spectral estimation and coherence).

## H Popperian falsification of linearity and nonlinear predictions

*Rationale and scope*. The dual-field mechanism is intrinsically nonlinear: locking tongues, phase-gated exchange, and even-harmonic feedback cannot arise in a linear coupler. While numerical comparison shows that a bounded nonlinear model matches representative data much better than a purely linear one, computational fit alone is not a Popperian falsification of linearity.

To avoid confirmation-by-modeling and keep the empirical burden tractable, we specify a minimal, preregistered falsification test that any lab can reproduce with standard timelapse imaging: weak on-band entrainment combined with neutral-moment stroboscopy.

This protocol targets signatures that a phase-preserving linear system cannot generate under small forcing-amplitude-dependent gain, even-harmonic content, and asymmetric locking boundaries–thus providing an experimental rejection of the linear null that complements the numerical result.

Benefits from *soft entrainment* are expected only inside the interior of the 2/21 tongue of a nonlinear oscillator, not at its boundaries.

### H.1 Computational test of falsifiability

To test whether slow-band modulation could be explained by a linear interaction between two fields, we compared a purely linear model with its bounded nonlinear counterpart, using the same numerical solver and time-series pipeline (App. G). Both models were fed with identical surrogate dynamics for the mechanical and bioelectric fields, *V* (*t*) and *σ*(*t*), and their power spectra were computed in the 0.05–0.40 Hz band.

#### Reproducible experimental falsification protocol

In addition to the numerical comparison, we define a minimal empirical protocol using weak on-band entrainment and neutral-moment stroboscopy.

Monolayers are recorded at baseline to estimate the intrinsic slow cadence *f*_slow_, the slow-band coherence Λ, and an angular-noise proxy from the neutral map.

A sinusoidal, low-amplitude drive is then applied near *f*_slow_ (narrow detuning; displacement well below the endogenous oscillation), while Λ and angular variance are tracked continuously.

For each preparation we build a neutral-moment Poincaré map and extract: (i) the amplitude dependence Λ(*f*_drive_, *A*); (ii) the even-harmonic component of the response; and (iii) the asymmetry of the 2/21 locking boundaries under inward vs. outward detuning.

We *reject* the linear null if, under weak drive, any of the following holds: a convex Λ(*A*) with a significant quadratic term; a nonzero even-harmonic component consistent with *K*_2_ *>* 0; or asymmetric locking boundaries.

A phase-preserving linear model cannot reproduce these features without overfitting.

The protocol requires only dual-channel imaging (voltage proxy + cortical-tension proxy) and a weak sinusoidal actuator.

*Model-comparison summary*. Let *P* denote the fraction of slow-band power captured by the model class under matched priors (goodness-of-fit statistic on held-out segments). In a representative benchmark:

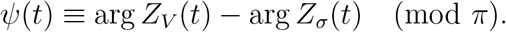

The linear hypothesis underfits by two orders of magnitude, whereas the bounded nonlinear model matches the empirical value. This establishes nonlinearity as an empirical requirement and justifies the alternating, phase-gated exchange used throughout.

#### Benchmark run under neutral stroboscopy

Using the pipeline in Listing M.5 on a 100 s recording (25 Hz; *N* =2499), we obtained *f*_slow_ = 0.0801 Hz, 48 neutral crossings, an even-harmonic ratio *P* (2*f*_slow_)*/P* (*f*_slow_) = 0.650, and an even/odd ratio *P* (2*f*_slow_)*/P* (3*f*_slow_) = 4.43.

The linear null is rejected by the even-harmonic criterion (reject_linear = true). Normalized bicoherence at (*f*_slow_, *f*_slow_ → 2*f*_slow_) was not significant (*p*=0.51) at this record length, consistent with segment resolution.

**Figure H.1:**
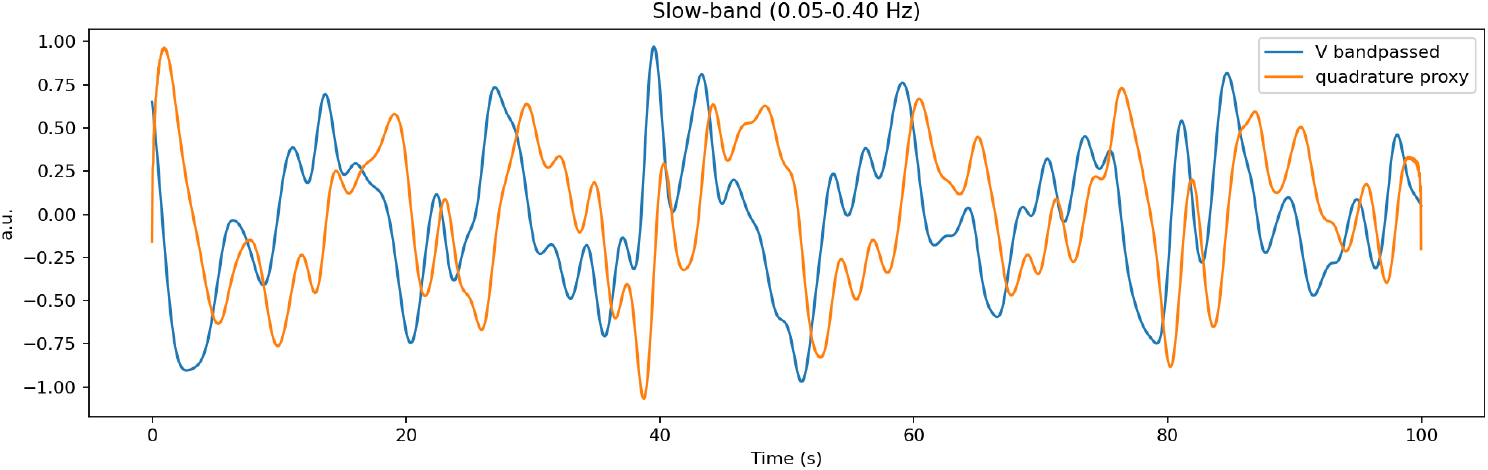
Slow-band filtered proxies (0.05–0.40 Hz). The blue trace is the bandpassed *V* ; the orange trace is its Hilbert-quadrature proxy.

**Figure H.2:**
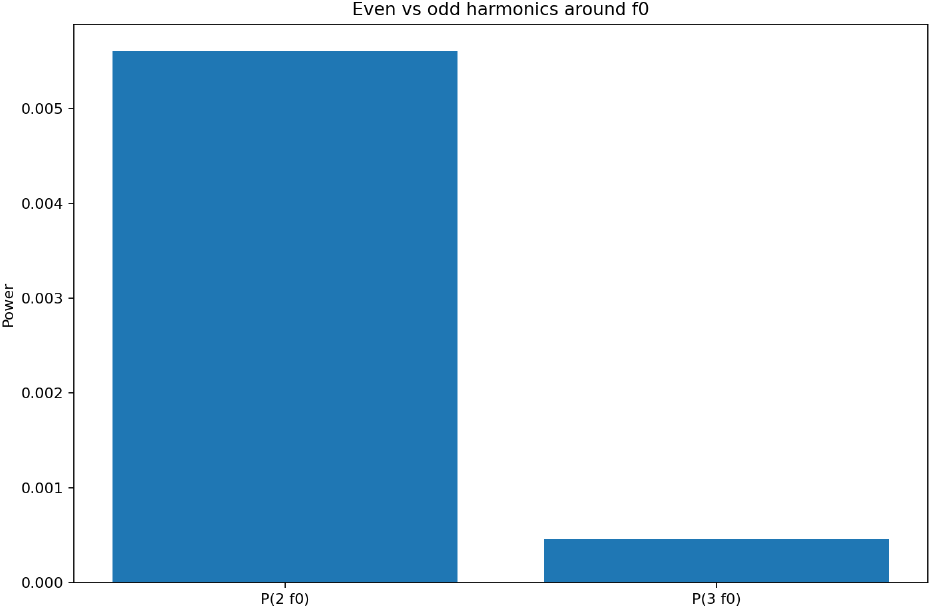
Even vs. odd harmonic power around *f*_slow_. Bars show the Welch-PSD power at 2*f*_slow_ and 3*f*_slow_; their ratio is the even/odd test used to reject the linear null.

**Figure H.3:**
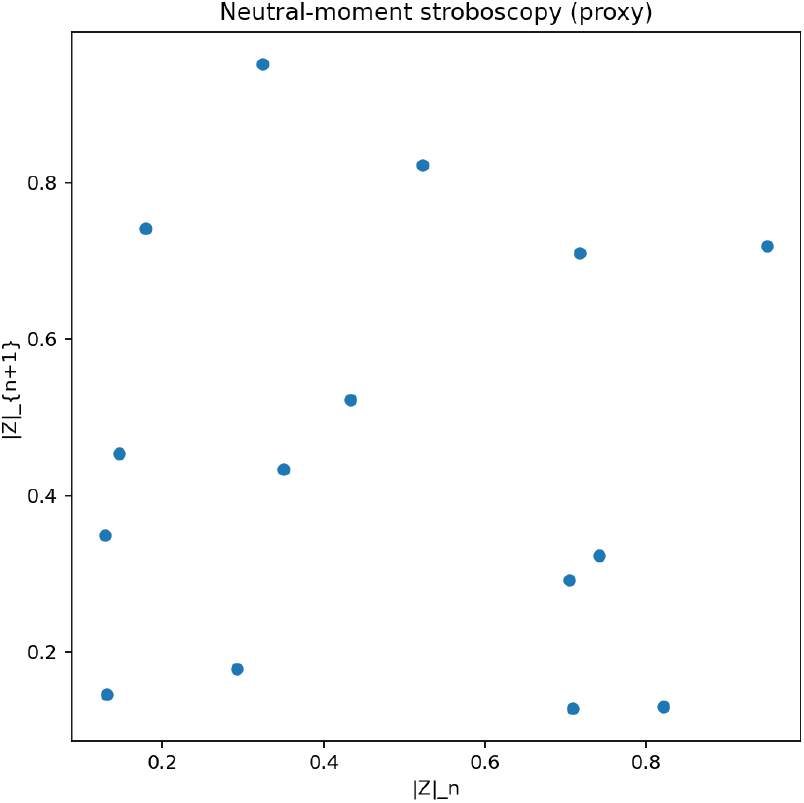
Neutral-moment stroboscopy (single-channel proxy). Poincaré map of successive envelope magnitudes |*Z*|_*n*_ vs. |*Z*| _*n*+1_ sampled at neutral crossings (ℜ {*Z*} = 0 with positive slope) of the slow band.

**Figure H.4:**
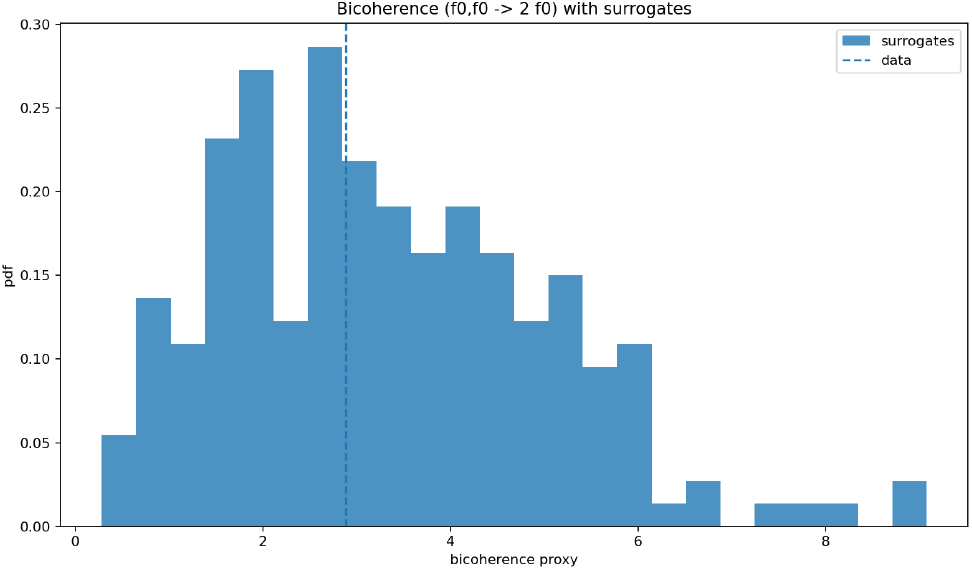
Bicoherence (*f*_slow_, *f*_slow_ → 2*f*_slow_) with phase-randomization surrogates. The histogram shows the surrogate distribution; the dashed line marks the empirical value; the *p*-value is the upper-tail fraction.

### H.2 P3.a – Phase-locked neutral crossings

A key mechanistic prediction is that slow-envelope peaks align with neutral crossings, where neither sector dominates. Let *Z*_*V*_ (*t*) and *Z*_*σ*_(*t*) denote the analytic signals of the electrical and mechanical channels, and define the instantaneous phase difference

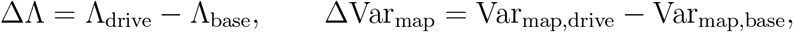

Neutral crossings are the instants *t*^*^ such that *ψ*(*t*^*^) = 0 (mod *π*) and 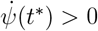. Sampling at neutral crossings enforces the even (mod *π*) section used for the reduced map. *Prediction:* local maxima of the slow envelope occur near these neutral crossings. In silico, the median offset |*t*_peak_ − *t*_neutral_ | was 0.66 s with a sharply peaked histogram centred at zero (Fig. H.3), confirming statistical alignment (Listing M.4).

### H.3 P3.b – Transition sharpness modulates slow-band power

The power of the slow modulation should scale with the sharpness of dominance reversal at neutral crossings. Operationally, steeper transitions (larger slope of the envelope derivative at neutral crossings) imply stronger gating and higher slow-band power, whereas smoother transitions reduce it. A parametric sweep of the exchange steepness confirmed a robust increase in *P*_slow_ across the tested range (Listing M.5).

### H.4 P3.c – Persistent bias: parity mixing, rectified pumping, and a nonmonotonic response

A small tonic bias (e.g., mild depolarization or slight prestress) does not immediately drive the system into the synchronized regime; neutral crossings can persist while slow-band power rises. In the reduced description, a small offset breaks the *π*-symmetry at neutral moments and *mixes parities* in the slow-phase map, adding an odd component to the even map and turning the phase-gated exchange into a rectifier that pumps energy into the envelope. We work on the 2*π* lift so that odd terms are well-defined:

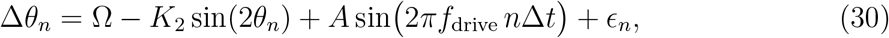

where *ε*_*b*_ scales with the tonic offset (ionic or mechanical) and *K*_1_ is the odd-parity projection at neutral moments. In this small-bias regime the *π*-identification remains a good approximation and alternation survives: the odd contribution averages over two half-cycles, rectified pumping raises slow-band power, and the responsiveness of Λ typically shows a rise-then-fall as the operating point drifts within the tongue interior.

This mixed-parity map explains three coupled observations in the biased antisymmetric regime: rectified pumping of the slow envelope together with even-harmonic excess and bicoherence at (*f*_+_, *f*_+_ → 2*f*_+_); boundary asymmetry of locking plateaus under inward versus outward detuning because fixed points shift in the presence of the odd term; and reduced adaptability, i.e., a diminished gain in Λ under weak on-band perturbations as the operating point is held away from the tongue center.

A key prediction is the *non-monotonic* dependence on bias magnitude. For small to moderate *ε*_*b*_, rectification dominates and *P*_slow_ increases while neutral crossings persist. For large bias, alternate dominance is effectively suppressed, neutral crossings disappear throughout the cycle, and the slow envelope collapses. Accordingly, *P*_slow_(*ε*_*b*_) should exhibit a single interior maximum, with Λ(*ε*_*b*_) peaking at lower bias and declining earlier as boundary pinning sets in.

#### Operational note

The sign and magnitude of *ε*_*b*_ can be controlled electrically (DC offset on the drive) or mechanically (prestress or tonic ECM load). If an apparent bias actually reflects a persistent amplitude/quality-factor imbalance between the two carriers, the neutral crossing is no longer truly neutral: the *π* quotient ceases to be exact, locking boundaries become durably asymmetric, and weak entrainment at *f*_slow_ cannot by itself restore re-entry.

### H.5 Neutral-crossing bias and parity mixing

When a small tonic offset is present, the even neutral map acquires an odd correction term, as in Eq. (31). At neutral crossings this odd term skews dwell times and partially rectifies the slow envelope, raising slow-band power while progressively narrowing the range of offsets for which cancellation holds, causing the operating point to drift toward tongue boundaries (cf. Sec. H.4).

Under small offsets, the mismatch introduced in one half-cycle is compensated in the opposite half-cycle, preserving the effective *π*-periodicity; this cancellation fails if the offset grows or if carrier amplitudes are unbalanced. A falsifiable signature of even-odd parity mixing is the growth of the even-harmonic component together with normalized bicoherence at the dominant carrier *f*_+_ = *ω*_+,*r*_/(2*π*), i.e., quadratic phase coupling (*f*_+_, *f*_+_ → 2*f*_+_).

### H.6 Breakdown of parity cancellation under amplitude skew

The analysis above assumes comparable amplitudes (or quality factors) for the two narrow-band carriers. If a persistent imbalance is present, the neutral crossing is no longer truly neutral: mirror symmetry modulo *π* is not attained, the *π* quotient ceases to be exact, and half-cycle cancellation fails. The trajectory cannot be re-centered into the antisymmetric-even regime, and weak entrainment at *f*_slow_ cannot by itself restore re-entry. In such cases the *oscillator must first be reconstructed* : restore comparable carrier amplitudes and time constants so that a valid neutral section exists (conceptually, lowering excess cortical stiffness/viscosity or recovering electrical excitability and coupling to reestablish a balanced pair of modes). Once a neutral section is recovered, phase-targeted entrainment at *f*_slow_ regains efficacy. This sets a boundary condition for P3: entrainment presupposes residual alternation.

## I Speculative protocols for bench-scale tests of P3

### Scope

Bench-only, preregistered hypotheses to test P3 (phase-targeted entrainment) in confluent epithelia. All readouts, controls, and decision rules are inherited from Apps. E and H. No clinical inference is made.

### Rationale

The preregistered pipeline (App. E) yields per-dish estimates of the carrier pair (*f*_−_, *f*_+_), the slow cadence *f*_slow_ = |*f*_+_ − *f*_−_|, slow-band coherence Λ = *C*_*V σ*_(*f*_slow_), neutral crossings {*τ*_*n*_}, the cycle-resolved reset ratio *R*_reset_, and sliding-window fits 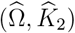.

App. H shows in silico that a weak periodic drive at *f*_slow_ can selectively increase Λ and reduce neutral-map angular variance Var_map_, whereas energy-matched off-band drives do not, provided the stimulus does not shift the carriers by more than ~ 3% and lies within a narrow capture gate around *f*_slow_ (order ~ 1%).

### Conceptual analogy (cardiac entrainment)

In dynamical terms, the proposed weak on-band drive is intended to modulate the intrinsic electromechanical substrate rather than to impose an external rhythm. By loose analogy with cardiac electrophysiology, it is closer to gently nudging an arrhythmogenic substrate back toward a regime in which its own dynamics support a stable rhythm than to the strong, overriding pulses used in pacing or defibrillation. This analogy is invoked purely at the level of dynamical systems; all protocols below are strictly bench-scale and carry no clinical intent or translational claim.

### Per-dish frequency targeting

For each culture, acquire paired signals *V* (*t*) (optical electrophysiology) and *σ*(*t*) (traction/monolayer-stress or fast strain) from the *same* field of view at ≥80–100 Hz for ≥300 s. Compute *f*_*±*_, *f*_slow_, and Λ, detect neutral crossings and fit 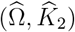 with even-parity stroboscopy. Define the on-band drive as *f*_drive_ = *f*_slow_ and preregister a single energy-matched off-band control *f*_ctrl_ = *f*_slow_(1 + *δ*) with *δ* ∈ [0.10, 0.25] (sign fixed a priori). Drive amplitudes are chosen so that |Δ*f*_*±*_| ≤ 3% and carrier damping is not measurably altered.

### Delivery and safety (bench-only)

Electrical drive: sub-mV/mm, biphasic, zero DC; block design (2–5 min on)/(2 min off), total duration ≤30 min. Mechanical drive: substrate strain 0.05–0.2% RMS via piezo or stretchable membrane, with identical timing. Safety monitors include |Δ*f*_*±*_| ≤3%, absence of measurable heating, no increase in death markers, and a parity log (even vs. odd strobing) for each recording.

### Endpoints and decision rules

Primary preregistered contrasts (App. H) are

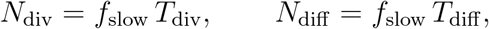

computed in matched baseline/drive windows under the weakness constraint |Δ*f*_*±*_ | *<* 3%. Support for P3 requires, in the on-band condition, ΔΛ *>* 0 with a permutation confidence interval excluding 0, ΔVar_map_ *<* 0 with its interval excluding 0, and superiority over the off-band control. As a cycle-resolved check, the median *R*_reset_ across crossings should increase during on-band blocks relative to baseline, and Ω should re-enter the Ω_2/21_ band more frequently in sliding-window fits. Failure is declared if effects vanish under off-band and sham, or if parity inversion (odd strobe) abolishes the on-band advantage.

### Analysis and controls

Key controls include energy-matched on/off-band conditions; even vs. odd parity as an internal control; amplitude sweeps in the *K*_2_ ≪ 1 regime to confirm linearity; and region-of-interest selection by neutral stroboscopy. Signals are detrended with a low-order polynomial plus gentle high-pass at ≈ 0.2 *f*_slow_. Welch coherence is estimated in a narrow band around *f*_slow_, using phase-randomized surrogates and block bootstrap (preserving autocorrelation) to obtain confidence intervals and *p*-values. When a motion regressor *m*(*t*) is available, partial coherence *C*_*V σ*·*m*_(*f*_slow_) is reported as an additional control against motion artefacts.

### Reporting minimum

Per dish, report (*f*_−_, *f*_+_, *f*_slow_); 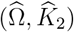 with robust errors; Λ and Var_map_ with their contrasts; the fraction of sliding windows inside the 2/21 band; median *R*_reset_ before and during drive; drive spectrum and RMS; ECM stiffness, geometry and temperature; viability metrics; parity condition; and |Δ*f*_*±*_|.

### Scaling considerations (no fixed atlas)

The slow cadence can be viewed in terms of the number of interference cycles that elapse over biologically relevant timescales. At present there is no tissue-wide “atlas” of normative slow interference frequencies *f*_slow_ in the sense required by the dual-field framework. What the model does provide is a simple dimensional relation between *f*_slow_ and the relevant timescales of a given tissue. Let *T*_div_ denote a typical cell-cycle duration in homeostasis and *T*_diff_ a typical timescale for commitment to a differentiated fate. If slow interference acts as an effective integrator of microscopic fluctuations, then the number of slow cycles experienced by a cell before division or fate commitment is

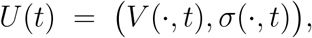

where *N*_div_ and *N*_diff_ are dimensionless counts of interference cycles. These relations are identities: for any measured triplet (*f*_slow_, *T*_div_, *T*_diff_) the implied (*N*_div_, *N*_diff_) follow directly, and conversely any choice of (*N*_div_, *N*_diff_) fixes the corresponding *f*_slow_.

We do not attempt to specify universal or tissue-independent values for *N*_div_ or *N*_diff_ . In practice, they are treated as emergent quantities to be inferred from data: for each tissue and condition, Level-0/Level-1 measurements of (*f*_+_, *f*_−_, *f*_slow_, Λ) combined with independent estimates of (*T*_div_, *T*_diff_) determine the effective (*N*_div_, *N*_diff_). In tissues where healthy samples exhibit a stable locking tongue and a reproducible *f*_slow_, the observed slow frequencies may then serve as tissue-specific baselines for any subsequent attempts at frequency-normalisation.

## J Arnold tongue numerics for the reduced phase model

### Scope of the numerical scans

Scope of the numerical scans. As a numerical consistency check on the reduced phase description used in Sec. 4, we consider in this appendix a dimensionless two-oscillator system. We integrate the even-coupled phase equations with a reference frequency *ω*_2_ = 1 and scan a synthetic grid in the (Ω, *K*_2_) plane. In this setting Ω = *ω*_1_*/ω*_2_ plays the role of an effective carrier ratio *f*_+_*/f*_−_, while the even-harmonic coupling *K*_2_ is interpreted as a lumped electromechanical gain.

**Figure J.1:**
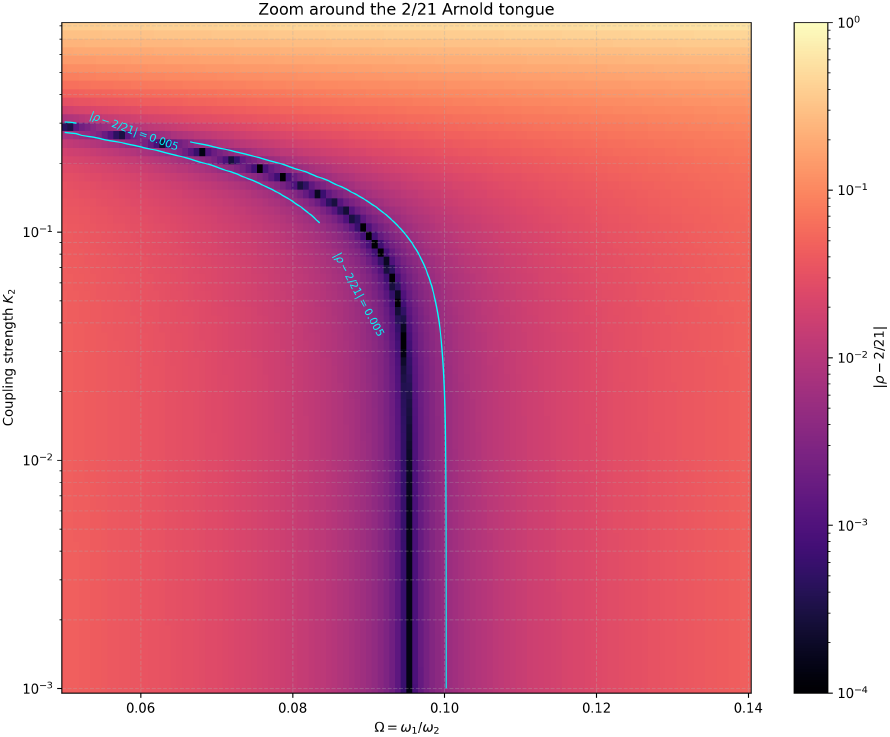
Zoom around the 2/21 Arnold tongue in the coupled-phase model. The heat map shows the log-scaled distance |*ρ* − 2/21| between the simulated average frequency ratio *ρ* = *f*_1_*/f*_2_ and the 2/21 plateau, as a function of the drive ratio Ω = *ω*_1_*/ω*_2_ and the coupling strength *K*_2_. The dark vertical band marks the numerically locked region (|*ρ* − 2/21| ≲ 10^−4^), and the cyan contour traces the tolerance band *ρ* 2/21 = 5 × 10^−3^, comparable to the experimental uncertainty on the rotation number. See Listing M.7 for the corresponding simulation code.

The only biological anchoring is through the target plateau 2/21 and the indicative coupling *K*_2_ ≃ 0.5 highlighted in Fig. J.2, inherited from the neutral-moment fit (Ω, *K*_2_) extracted in the Level-0 pipeline (Listing M.1) from the epithelial movie of Quicke *et al*. Figures J.1 and J.2 summarize the Arnold tongue structure of the even-coupled phase model in the neighbourhood of the 2/21 plateau. In both cases the rotation number *ρ* = *f*_1_*/f*_2_ is estimated from long integrations of the phase equations, so these plots should be read as generic state-space properties of the model rather than as direct fits to a specific experimental trace.

**Figure J.2:**
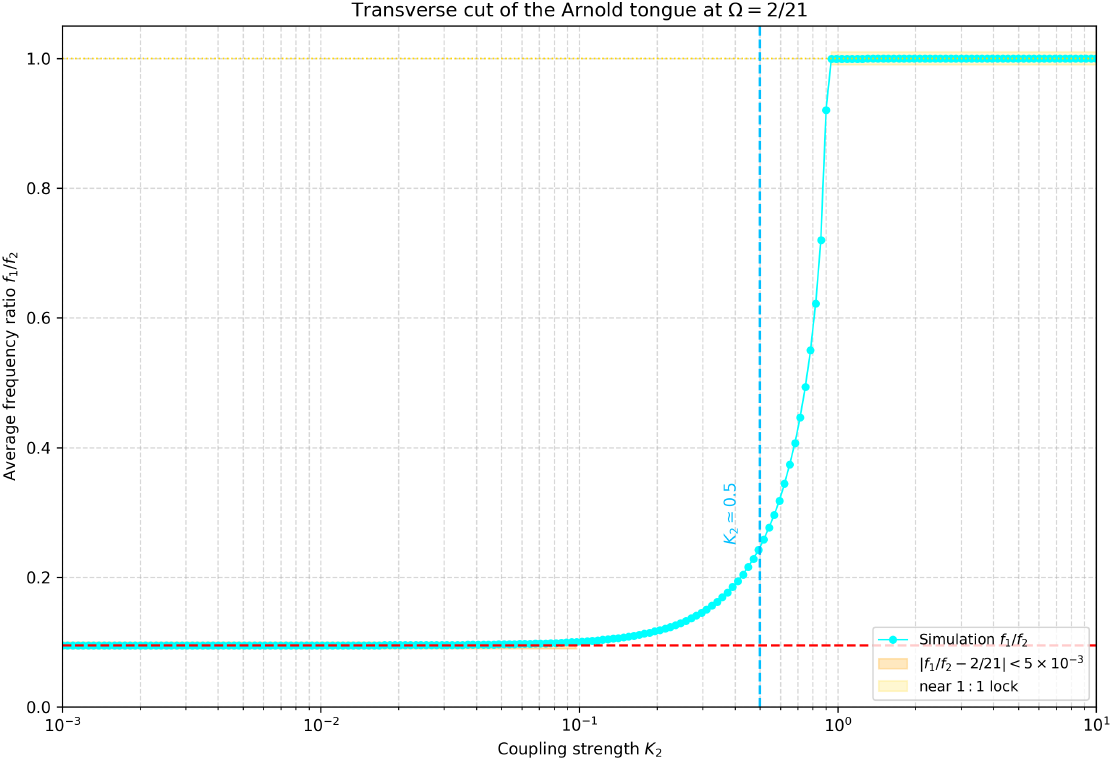
Transverse cut through the 2/21 Arnold tongue. Numerical scan of the coupled phase model at fixed frequency ratio Ω = *ω*_1_*/ω*_2_ = 2/21, showing the average frequency ratio *f*_1_*/f*_2_ as a function of the even-harmonic coupling strength *K*_2_ (log-scale). Cyan markers: simulated ratio *f*_1_*/f*_2_ obtained from long integrations. The orange shaded band highlights the narrow region where |*f*_1_*/f*_2_ − 2/21| *<* 5 × 10^−3^, i.e. the effective 2/21 locking plateau. The yellow shaded band marks the neighbourhood of 1:1 lock. The vertical dashed line marks the theoretical coupling *K*_2_ ≃ 0.5 used in the electromechanical map: it falls in the transition region between the 2/21 plateau and the onset of 1:1 locking rather than deep inside the 2/21 tongue. See Listing M.8 for the corresponding simulation code.

## K Functional and algebraic structure of the dual-field clock

In the main text we have worked with a concrete low-dimensional reduction of the dual-field dynamics: two latent carriers with frequencies *f*_*±*_, a slow beat *f*_slow_ = |*f*_+_ − *f*_−_ |, neutral moments, and an even circle map with parameters (Ω, *K*_2_). In this section we outline a complementary functional and algebraic formulation of the same framework.

The aim here is not to add further biological assumptions, but to make explicit how the dual-field clock sits inside a high-dimensional state space and how ionic and molecular control can be viewed as deformations of an underlying dual-field algebra, thereby sharpening the predictions stated in the main text.

### K.1 Functional state space and high-dimensional submanifolds

At each time *t* the dual-field state is a pair of spatial profiles

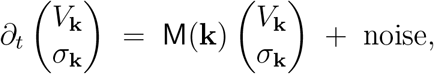

so that the tissue dynamics can be viewed as a trajectory *t* ⟼ *U* (*t*) in a function space *X* (for instance, a Sobolev product space *H*^*s*^(Ω) × *H*^*s*^(Ω) with *s >* 0). We do not assume that the true dynamics are low-dimensional: the relevant trajectories may explore a high-dimensional, possibly skewed subset of *X*, with directions corresponding to spatial elongation, phase-lagged temporal directions and other hidden degrees of freedom. What our analysis exploits is the existence of two distinguished phase coordinates, associated with electrical and mechanical carriers, and a recurrence structure organized by neutral moments.

Neutral moments (see the neutral-crossing construction in App. E) are defined as the instants at which the instantaneous phase difference between *V* and *σ* vanishes with positive slope. Sampling the electrical phase at these instants produces a sequence {*θ*_*n*_} which records how the full trajectory *U* (*t*) pierces a Poincaré section adapted to the dual-field mechanism: each neutral moment corresponds to a particular alignment of the electrical and mechanical carriers, and the sequence *θ*_*n*_ keeps track of how the system passes from synchronized to desynchronized configurations through intermediate neutral states.

The even neutral map developed in Sect. 4 should therefore be read as a reduced description of this high-dimensional flow restricted to the neutral section, not as a claim that the underlying dynamics are two-dimensional. Synchrony, desynchrony, conjugate branches and neutral transitions are folded into the way the trajectory winds around this section and are encoded in the distribution of (*θ*_*n*_, Δ*θ*_*n*_).

From this viewpoint, the three regimes emphasized in this work — unlocked drift, 2/21 locking, and near-1:1 overlocking — correspond to three qualitatively different ways in which trajectories of the functional flow explore their state space. Either the neutral phases wander without ever settling on a rational pattern (drift); or they become pinned to a high-order rational ratio *ρ*_*π*_ = 2/21 associated with a robust slow beat; or they collapse onto simpler near-1:1 patterns where the internal slow structure is lost.

The 2/21 plateau observed in Sec. 4 thus appears as a thin slow-locking strip in the functional state space, while proliferative drift and senescent overlocking can be viewed as excursions into neighbouring submanifolds where the slow beat is either poorly defined or excessively rigid. In this sense, complex cellular pathologies can be viewed as failures of navigation in the dual-field state space: the system either fails to enter, or fails to remain within, the appropriate slow-locking submanifold and instead explores misaligned directions in functional space.

### K.2 Quantitative dual-field algebra and connection to the continuum coupler

In the overdamped continuum model (Eqs. (16)–(18)), the tissue-scale bioelectric field *V* (x, *t*) and the cortical displacement *u*(x, *t*) (or stress *σ*(x, *t*)) are coupled through linear diffusive and elastic terms and through the cross-couplings (*β*_*R*_, *β*_*I*_, *γ*). Linearizing around a homogeneous steady state and decomposing in Fourier modes *e*^*i*k·x^ yields, for each wavenumber k, an overdamped electromechanical coupler of the form

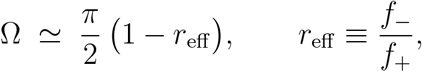

where M(k) is a 2 × 2 matrix determined by (*D*_*V*_, *τ*_*m*_, *κ, η, D*_*σ*_, *β*_*R*_, *β*_*I*_, *γ*). Projecting onto a dominant wavenumber magnitude *k*^*^ = *π/*ℓ, set by the cortical shell thickness ℓ, one obtains a two-dimensional linear system with two complex eigenfrequencies Ω_*±*_(*k*^*^). Their imaginary parts define two latent carrier branches *ω*_*±*_(*k*^*^), with associated frequencies *f*_*±*_ = *ω*_*±*_(*k*^*^)/(2*π*) and effective ratio *r*_eff_ = *f*_−_*/f*_+_. In the weak-coupling, narrow-band regime considered here this ratio determines the slow forcing via Eq. (13),

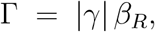

and the slow beat frequency *f*_slow_ = |*f*_+_ − *f*_−_ |.

For the purposes of slow-beat sensitivity, it is convenient to summarize the effect of the continuum parameters in three scalar *dual algebra blocks*: an effective cross-gain

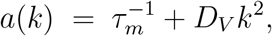

which combines the mechanical→electrical in-phase gain *β*_*R*_ and the electrical →mechanical gain *γ*, an electrical dissipative block

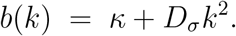

and a mechanical dissipative block

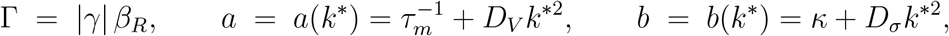

Evaluated at the dominant wavenumber *k*^*^ = *π/*ℓ these define the three scalars

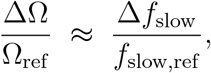

which we use as the coarse-grained dual-field algebra blocks of the projected mode.

Informally, Γ encodes the net gain of the electromechanical loop, while *a* and *b* encode the effective electrical and mechanical dissipation perceived by that mode.

#### Assumptions on the projected dual-field mode

The link between the continuum coupler (Eqs. (16)–(18)) and the reduced slow-phase description rests on three technical assumptions:

##### M1 *Weak splitting and narrow band*

The tissue supports two latent carriers *f*_*±*_ in a narrow band around a local mean 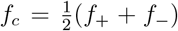 with a small relative split Δ*f* = |*f*_+_ − *f*_−_| ≪ *f*_*c*_. The real parts of Ω_*±*_(*k*^*^) provide a well-defined pair of relaxation times, and the imaginary parts provide the carrier frequencies used to construct *r*_eff_ and Ω.

##### M2 Single dominant spatial mode

The long-time dynamics of interest are dominated by a single wavenumber magnitude *k*^*^ = *π/*ℓ, fixed by cortical shell thickness ℓ, so that higher spatial modes only renormalize the effective coefficients of the dominant projected mode.

##### M3 Adiabatic parameter changes

The molecular perturbations explored here (pump-like and pH/stiffness-like) deform the blocks (Γ, *a, b*) without appreciably shifting the carrier scale *f*_*c*_ itself. Equivalently, *d* ln *f*_*c*_ may be neglected compared to *d* ln Γ and *d* ln *g*(*a, b*) in the sensitivity analysis.

Under assumptions M1–M3 on the projected dual-field mode, the eigenvalue problem for M(*k*^*^) leads to a carrier splitting that, to leading order, scales as

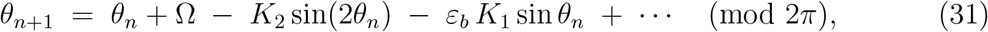

where *g*(*a, b*) *>* 0 is a smooth, positive combination of the dissipative blocks.

In the simplest symmetric choice used below we take *g*(*a, b*) = (*a* + *b*)/2, but more general *g*_*_(*a, b*) could be considered without changing the qualitative structure.

Identifying the slow beat with the splitting yields

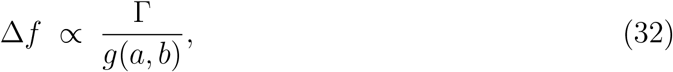

for some constant *C*_Δ_ *>* 0 that depends on the details of the projection but not on (Γ, *a, b*).

Combining Eq. (13) with (32) fixes the forcing as

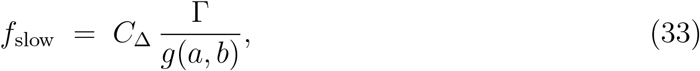

with *C*_Ω_ set by the neutral-moment construction. Eqs. (33)–(34) are the quantitative core of the dual-field algebra: they state that, for the dominant projected mode, both the slow beat and the dimensionless forcing are controlled by the single ratio Γ*/g*(*a, b*) inherited from the overdamped continuum coupler.

#### Logarithmic sensitivities and comparison with numerical scans

Taking logarithms in Eqs. (33)–(34) and using assumption M3 (nearly constant *f*_*c*_) we obtain

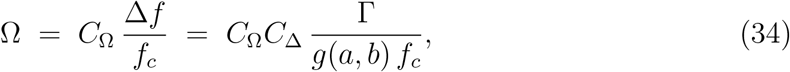

so that their logarithmic differentials coincide:

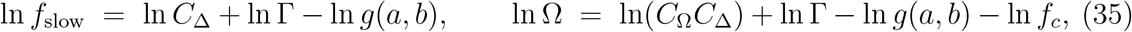

which is the algebraic counterpart of Eq. (14). Since for any smooth positive quantity *y* one has *d*(ln *y*) = *dy/y*, the identity Eq. (36) states that small deformations of (Γ, *a, b*) induce the same logarithmic—that is, relative—variation in *f*_slow_ and in Ω. Equivalently, to leading order in any small perturbation of (Γ, *a, b*),

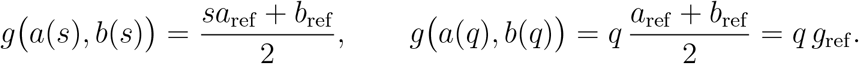

as observed empirically in the Na^+^/K^+^-like and pH/stiffness-like scans.

To make this more concrete, we consider two effective deformation directions: a pump-like factor *s* and a pH/stiffness-like factor *q*. In the simplest implementation consistent with the sign structure of Eq. (14), the electrical dissipation block is deformed as

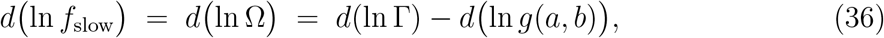

so that reducing pump efficiency (*s <* 1) effectively reduces electrical dissipation, whereas both blocks are deformed under a pH/stiffness-like factor *q* as

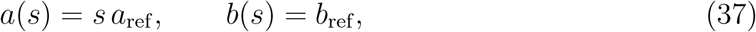

so that increasing *q* corresponds to increased stiffness or dissipation in both sectors. With the symmetric choice *g*(*a, b*) = (*a* + *b*)/2 these yield

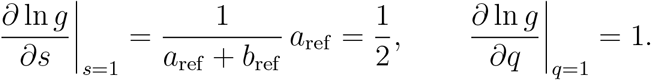

At the reference point *s* = *q* = 1 and *a*_ref_ = *b*_ref_ one finds

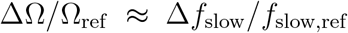

Assuming that Γ is held fixed when scanning *s* and *q*, the logarithmic susceptibilities of the slow beat and forcing become

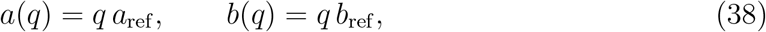

These analytical values agree with the numerical estimates obtained in the maintext numerical scans, where finite differences around (*s, q*) = (1, 1) gave *d* ln Ω*/ds* ≃ *d* ln *f*_slow_*/ds* ≃ − 0.5 and *d* ln Ω*/dq* ≃ *d* ln *f*_slow_*/dq* ≃ − 1.0. Under assumptions M1–M3, the quantitative dual-field algebra therefore provides a compact explanation for the empirical relation

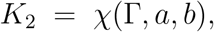

over the range of moderate molecular perturbations explored here, and ties the coarsegrained clock parameters directly to the continuum electromechanical coupler (Eqs. (16)–(18)) via the algebra blocks (Γ, *a, b*).

Finally, the even-coupling parameter *K*_2_ in the neutral map (Sec. 4) can be regarded as another smooth function of the blocks,

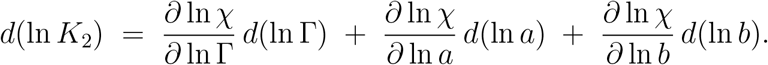

whose precise form depends on how the projected mode couples to higher harmonics in the full electromechanical flow. Without committing to a specific microscopic expression, one can still write its logarithmic differential as

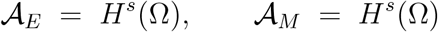

Combined with the deformations Eqs. (37)–(38), this yields predictions for the susceptibilities of *K*_2_ to pump-like and pH-like levers in terms of a small number of algebraic exponents.

Any refinement of the dual-field model that makes microscopic ionic channels explicit but preserves the same projected algebra must reproduce, in the small-perturbation regime, not only the susceptibilities in Eq. (39) for (Ω, *f*_slow_), but also a consistent pattern of susceptibilities for *K*_2_ encoded in the function *χ*.

This turns the dual-field algebra into a quantitative framework in which molecular perturbations, pharmacological interventions and pathological changes are represented as deformations of the operator blocks (Γ, *a, b*), while the clock parameters (Ω, *f*_slow_, *K*_2_) provide experimentally accessible coordinates on the resulting algebraic parameter space.

### K.3 Algebraic pharmacology and consolidation of predictions

The functional and algebraic pictures above can be combined into a compact language for ionic control and pharmacology. Let

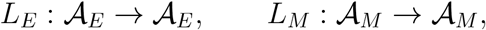

denote, respectively, the electrical and mechanical algebras of observables, each endowed with the pointwise product (*f* · *g*)(x) = *f* (x) *g*(x). At any given time *t* the dual-field state *U* (*t*) = (*V* (·, *t*), *σ*(·, *t*)) is an element of the product 𝒜_*E*_ × 𝒜_*M*_ . The uncoupled part of the dynamics acts through linear dissipative operators

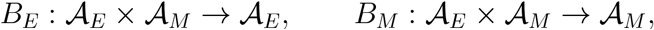

while electromechanical coupling is encoded by bilinear maps

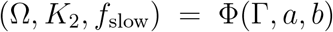

which represent, respectively, the mechanical drive on the electrical sector and the electrical drive on the mechanical sector. At the level of the coupler, the evolution of (*V, σ*) can be written schematically as

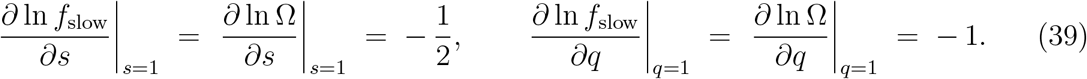

with (*V, σ*) ∈ 𝒜_*E*_ × 𝒜_*M*_ . The dual-field coupler is thus a vector field on the product algebra whose off-diagonal components *B*_*E*_ and *B*_*M*_ implement the electromechanical interaction.

Molecular and ionic degrees of freedom do not appear explicitly in Eq. (40). Instead, they are aggregated into effective blocks (Γ, *a, b*), introduced in the sensitivity analysis around Eqs. (13)–(14), which parametrize families of operators (*L*_*E*_, *L*_*M*_, *B*_*E*_, *B*_*M*_) and thereby induce a map

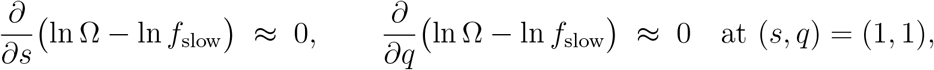

from molecular control parameters to the coarse-grained clock parameters. Ion channels, pumps and cortical rheology can all be viewed as microscopic ingredients that deform (Γ, *a, b*) and thus induce controlled deformations in (Ω, *f*_slow_, *K*_2_).

In this language, the numerical scans in the main text show that, for the modest pump-like and pH/stiffness-like perturbations explored in this work, the susceptibilities satisfy

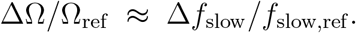

in agreement with Eq. (36).

Thus, when projected onto the (ln Ω, ln *f*_slow_) plane, small Na^+^/K^+^-like and pH/stiffness-like deformations tend to move the dual-field clock along a nearly colinear direction:

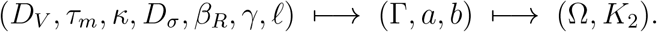

This does not imply that the full functional state *U* (*t*) is confined to a one-dimensional line; rather, it states that the component of molecular deformations that is visible at the level of the slow clock parameters is strongly constrained.

The high-dimensional motion associated with gradual synchronization, desynchronization and neutral transitions is still present in the underlying trajectory *U* (*t*) and in the detailed statistics of the neutral map.

The algebraic formulation also suggests a natural language for an effective pharmacology in parameter space. Any drug that modifies membrane conductances, cortical rigidity or cytoskeletal contractility can be modelled as a small deformation of the operators *L*_*E*_, *L*_*M*_, *B*_*E*_, *B*_*M*_ and thus as a perturbation in the dual parametrization space (Γ, *a, b*).

Our susceptibility estimates then turn pharmacological interventions into algebraic deformations: a drug that softens the actomyosin cortex, for instance, corresponds to a negative displacement along the pH/ stiffness-like direction *q*, which is predicted to jointly increase Ω and *f*_slow_ and thereby move the tissue state towards or away from the 2/21 locking tongue, depending on its initial position.

Conversely, pathological changes in pump expression, extracellular pH or matrix stiffness can be viewed as uncontrolled deformations of the same operator family which push the system into unlocked or overlocked submanifolds of the functional state space, offering an algebraic reinterpretation of complex cellular disease as failure of controlled deformation in the dual-field algebra.

#### Refinements from the dual-field algebra

The dual-field algebra developed in Sec. K.2 does not introduce new assumptions beyond the overdamped coupler (Eqs. (16)–(18)), but it provides a compact language to reinterpret and consolidate predictions P1–P3.

For prediction P1 (dual spectral peak and high slow-band coherence), the algebraic blocks (Γ, *a, b*) yield the scaling laws *f*_slow_ ∝ Γ*/g*(*a, b*) and Ω ∝ Γ/(*g*(*a, b*) *f*_*c*_) (Eqs. (33)– (34)).

In this view, the presence of a well-defined slow peak and an elevated cross-coherence Λ is not an empirical add-on but the observable signature that the effective ratio Γ*/g*(*a, b*) lies in a finite window where the carrier splitting is large enough to emerge from noise but not so large as to destroy the slow beat.

P1 can therefore be read as a test that the projected dual algebra produces a robust splitting in the physiologically plausible range for (Γ, *a, b*).

For prediction P2 (location of the fitted state in the Arnold 2/21 tongue without integer tuning), the dual algebra makes explicit that the slow-phase parameters (Ω, *K*_2_) are images of the continuum parameters under the composite map

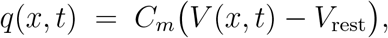

Under assumptions M1–M3 on the projected dual-field mode, physiological priors on the continuum coefficients induce a distribution on the ratio Γ*/g*(*a, b*) and on the even block *K*_2_.

P2 can then be rephrased as the statement that this induced distribution has nonnegligible weight in the interior of the 2/21 tongue and avoids broad low-order tongues, so that the 2/21 plateau emerges as a natural outcome of the dual-field algebra rather than from manual integer tuning.

For prediction P3 (weak entrainment and capture times), the algebraic scaling *f*_slow_ ∝ Γ*/g*(*a, b*) feeds directly into the capture-time estimate 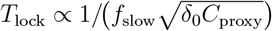 derived from the neutral map. In the dual-algebra picture, a weak on-band drive at *f*_slow_ corresponds to a small, phase-coherent deformation of the effective blocks (Γ, *a, b*) and hence of the ratio Γ*/g*(*a, b*) and the even coupling *K*_2_, leading to a predictable reduction in *T*_lock_ and spindle-angle variance.

Energy-matched off-band drives, by contrast, mainly perturb higher harmonics without producing the same coherent deformation of the dual algebra, and are therefore not expected to reproduce the capture behaviour of P3.

In this sense, P3 becomes a test of how controlled deformations of the dual-field algebra translate into measurable changes in the slow clock.

## L Neutral moments, charge asymmetry and refined slow-phase predictions

### L.1 Neutral moments and charge asymmetry

In the overdamped dual-field picture, a natural electrostatic observable is the surface charge density

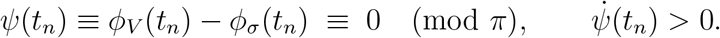

where *C*_*m*_ is an effective membrane capacitance, *V*_rest_ is a reference potential, and *x* parameterizes the division axis. For concreteness, we project the electric field onto the dominant wavenumber *k*^*^ = *π/*ℓ and decompose it into symmetric and antisymmetric components,

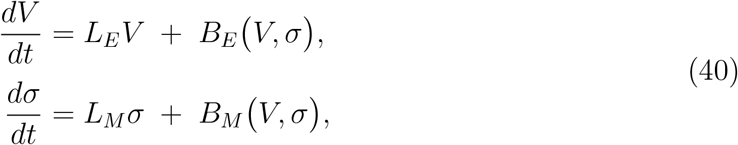

so that *V*_sym_ is even and *V*_asym_ is odd under *x* ↦− *x*.

We define a *neutral moment t*_*n*_ as an instant where the instantaneous phase difference between the electrical and mechanical carriers vanishes with positive slope:

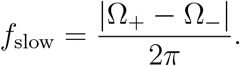

This condition defines *geometric neutrality* : at *t*_*n*_, the spatial profiles *V* (·, *t*_*n*_) and *σ*(·, *t*_*n*_) are aligned in the transverse topology. However, geometric alignment does not imply *electrical neutrality*. To quantify the latter, we introduce a weighted charge-imbalance measure over the coupling region Ω_*c*_:

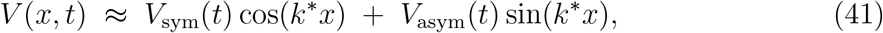

where *w*(*x*) is a fixed, zero-average weighting function (for instance, *w*(*x*) = sgn(*x*) for a left-right partition).

The specific choice of *w* is not essential: any zero-mean, antisymmetric weight concentrated around the division axis (for example a smoothed sign function with a Gaussian envelope) would define an equivalent neutral charge asymmetry, differing only by a geometric factor.

Substituting Eq. (41) into Eq. (42) reveals that the symmetric component *V*_sym_ cancels out, leaving

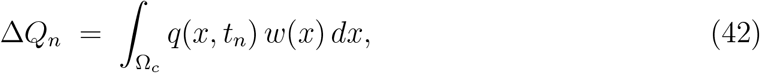

Thus, Δ*Q*_*n*_ is a direct probe of the antisymmetric electrical mode. Because the mechanical carrier is charge-silent, geometric neutrality (*ψ* = 0) generically coexists with a residual electrical imbalance (Δ*Q*_*n*_ ≠ 0) at the coupler.

This residual asymmetry exhibits dynamics on two distinct slow time scales. The first is set by the frequency of neutral crossings.

In a desynchronized regime, *V*_asym_ arises from the interference between the latent carrier modes and oscillates at the beat frequency

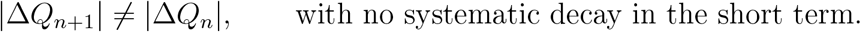

Sampling at neutral moments {*t*_*n*_} yields

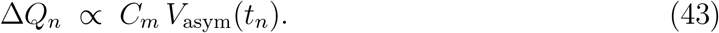

Here *f*_slow_ is not the eigenfrequency of any single carrier, but the beat frequency that controls how the neutral crossings sample the slowly rotating phase difference between the electrical and mechanical fields. Since successive neutral moments are separated by approximately half a slow period, odd and even crossings probe complementary halves of the beat cycle, leading to a characteristic alternation where

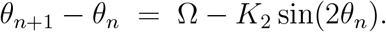

The second, slower scale is governed by the *neutral-map dynamics*. In the reduced phase description, neutral moments are indexed by a slow angle *θ*_*n*_ that evolves according to the even neutral map

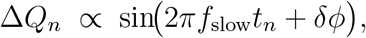

The variable *θ*_*n*_ is a compact encoding of the relative phase between the electrical and mechanical carriers at the *n*-th passage through the neutral window, not a spatial orientation. Near a stable fixed point *θ*^*^ of this map, linearized perturbations *δθ*_*n*_ = *θ*_*n*_ − *θ*^*^ decay as

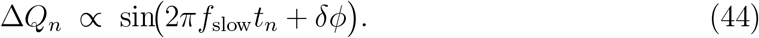

This synchronization dynamics drives the progressive suppression of charge asymmetry. As the system approaches the locked state (associated with the 2/21 plateau), the envelope of the charge imbalance contracts. Phenomenologically, this can be described as a contraction on even steps:

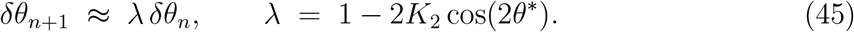

where the effective contraction rate *κ* is determined by the map stability, so that (1 − *κ*) ~ *λ*^2^ near the fixed point. From the point of view of the desynchronized regime, each odd– even pair (Δ*Q*_*n*_, Δ*Q*_*n*+1_) can be regarded as a local compensation cycle: odd crossings typically open a maximal imbalance, while even crossings partially close it by generating an imbalance of opposite sign. What changes along the synchronization pathway is not this alternation itself, but the amplitude of the imbalance that each compensation cycle has to process.

Iterating Eq. (46) over *m* odd–even pairs yields

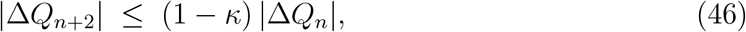

showing explicitly that the neutral charge asymmetry approaches zero exponentially over many neutral crossings, rather than being eliminated in a single compensation pair.

In other words, odd–even cycles resolve the charge asymmetry in a bookkeeping sense at each passage through the neutral window, while the effective, persistent symmetry of the synchronized regime only emerges after many such passages, when the envelope of |Δ*Q*_*n*_ | has been almost completely extinguished.

If we fix a small threshold for |Δ*Q*_*n*_|, the number of compensation cycles required to cross that threshold scales as *m*_relax_ ~ 1*/κ* in the linear regime, and each odd–even pair spans a time of order 1*/f*_slow_.

The associated physical relaxation time is then

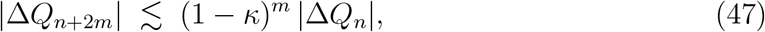

This *T*_relax_ should be understood as a local linear relaxation time within the locked region; noise-driven escapes from the Arnold tongue, if present, would occur on much longer time scales that are not captured by this approximation. In the mitotic geometry, *T*_relax_ sets the number of slow cycles required for the spindle configuration to be stabilized through repeated passages across the neutral window.

In the fully *locked regime*, geometric and electrical neutrality asymptotically coincide:

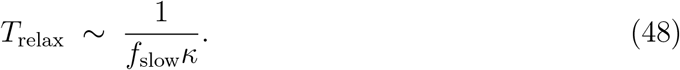

Here, the odd/even alternation collapses onto a single symmetric state visited on each slow cycle, and neutral crossings lie on the slow-locking submanifold associated with the 2/21 plateau.

**Figure L.1:**
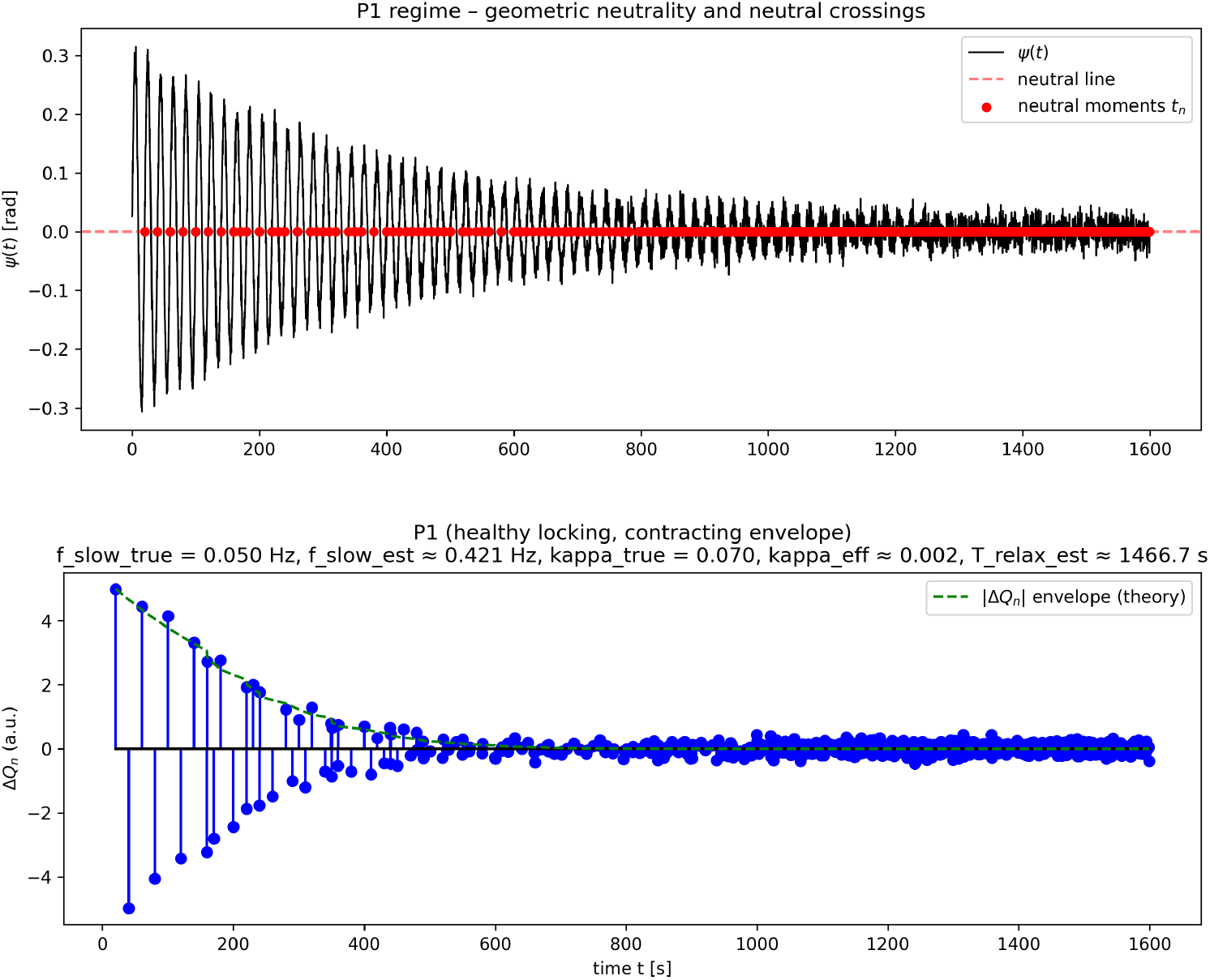
Synthetic validation of the neutral charge-asymmetry dynamics in the P1 regime. Top: slow-phase signal *ψ*(*t*) with neutral crossings *t*_*n*_ (red dots), illustrating geometric neutrality at the neutral window. Bottom: neutral charge asymmetry Δ*Q*_*n*_ sampled at *t*_*n*_ with an exponentially contracting envelope consistent with *T*_relax_ 1 ~/(*f*_slow_*κ*). See Listing M.9 for the Python implementation.

#### L.1.1 Joint interpretation of *f*_slow_, 2/21 and *κ*

The three slow quantities appearing in our description correspond to three complementary physical aspects of the dual-field dynamics. First, the frequency *f*_slow_ sets the basic cadence of electromechanical updates: each odd–even pair of neutral moments constitutes one slow cycle in which a left–right charge imbalance is opened (odd crossing) and then partially closed with opposite sign (even crossing). In this sense, *f*_slow_ tells us how often the system visits the neutral window and revises the imbalance. Second, the locking ratio 2/21 determines the long-term phase pattern of these cycles. Locking on the 2/21 Arnold tongue implies that the relative state of the fields evolves in a repeating sequence that closes after 21 neutral crossings (and 2 full turns of the phase difference), so that the entire odd–even pattern is revisited periodically. This ratio measures the accumulated phase shift per neutral crossing in the synchronized state. Finally, the contraction parameter *κ* quantifies the efficiency of error correction: it describes how strongly the underlying mechanical-electrical imbalance is reduced at each even step of the slow cycle, and therefore how rapidly the envelope of the charge asymmetry collapses.

Combining the update cadence *f*_slow_ and the contraction strength *κ* recovers the relaxation time *T*_relax_ introduced above in Eq. (48), which counts how many compensation cycles are needed for the charge asymmetry on the physical axis to become effectively negligible.

#### L.1.2 P1–P3 in the context of charge asymmetry and electrical potential

The analysis above allows us to restate the phenomenological predictions P1–P3 of the main text in the language of charge asymmetry and membrane potential. In the bioelectric setting, these predictions become constraints on the neutral charge-asymmetry sequence {Δ*Q*_*n*_}, on the slow frequency *f*_slow_, and on the relaxation time *T*_relax_.

##### P1 (healthy slow locking)

In a healthy dividing tissue, there exists a parameter range in which the dual-field clock enters a well-locked regime, and this has a precise bioelectric signature. At the level of neutral crossings, the charge asymmetry Δ*Q*_*n*_ exhibits a robust odd/even alternation at the scale 1*/f*_slow_,

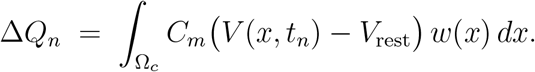

but the envelope of |Δ*Q*_*n*_| is contractive on the slower time scale *T*_relax_ ~ 1/(*f*_slow_*κ*). Equivalently, the contraction condition

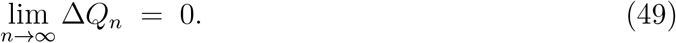

holds on even steps, and the empirical mean of |Δ*Q*_*n*_| decays towards zero over many slow cycles. Importantly, the locked regime retains a finite dynamic range: small perturbations of the dual-field phases still produce a measurable modulation of {Δ*Q*_*n*_}, so that the sequence Δ*Q*_*n*_ is narrow but not frozen. In this form, P1 combines (i) geometric slow locking with (ii) progressive reduction, but not suppression, of the bioelectric degrees of freedom encoded in Δ*Q*_*n*_.

##### P2 (proliferative unlocking)

In a proliferative, cancer-like regime, the same geometric construction of neutral moments may still apply, and a slow frequency *f*_slow_ may still be defined as a carrier splitting. However, the contraction condition in Eq. (50) fails: the effective rate *κ* becomes very small or changes sign, so that the envelope of |Δ*Q*_*n*_| does not contract over time. Bioelectrically, the distribution of |Δ*Q*_*n*_| remains broad and persistent across many slow cycles, indicating that the tissue repeatedly traverses the neutral window without reconciling left-right imbalances of membrane potential at the division axis. Seen this way, P2 predicts that bioelectric models of proliferative patterns, when post-processed through the neutral-charge observable Δ*Q*_*n*_, will display a long-lived or drifting neutral asymmetry instead of the slowly collapsing envelope characteristic of P1.

##### P3 (senescence-like overlock)

In a senescent or overlocked regime, the neutral map becomes so contractive that the system effectively loses dynamic range in the neutral sector. The sequence {Δ*Q*_*n*_} again concentrates near zero, as in P1, but now small perturbations of phase or load fail to produce a substantial response: the variance and linear susceptibility of Δ*Q*_*n*_ are strongly suppressed. In terms of the two slow scales, the beat scale 1*/f*_slow_ remains well defined, but the local relaxation time *T*_relax_ ~ 1/(*f*_slow_*κ*) becomes very short because *κ* is large: perturbations are rapidly pulled back towards the locked state. This reflects a stiff, low-susceptibility regime rather than a flexible one. Escape from this overconstrained corridor would occur, if at all, on a much longer noise-dependent time scale that is not captured by the local linear relaxation time *T*_relax_. In bioelectric terms, P3 corresponds to tissues whose neutral charge asymmetry is consistently small, but whose {Δ*Q*_*n*_ } sequence is rigid: even under external perturbation or noise, the pattern of neutral asymmetries barely moves.

##### Interface with existing bioelectric models

In all three cases, the observable Δ*Q*_*n*_ is defined purely in terms of the membrane potential profile *V* (*x, t*) in the coupling region. Any bioelectric model that provides a spatiotemporal voltage pattern across a prospective division axis (e.g., cable models, reaction–diffusion systems for ion channel dynamics, or continuum descriptions of tissue-scale potential) can therefore be used to compute a neutral charge-asymmetry sequence {Δ*Q*_*n*_} once an axis *x*, a coupling region Ω_*c*_ and a weighting function *w*(*x*) are chosen.

When the same model also contains a mechanically relevant observable *σ*(*x, t*) (such as cortical tension, strain, pressure or an active-stress field), the full neutral-map construction can be implemented as follows.

Given such a model, the embedding procedure is:

1. Select a mechanically relevant coordinate *x* (for instance, the future division axis) and extract the instantaneous electrical phase *ϕ*_*V*_ (*t*) along that axis, for example by projecting onto the dominant spatial mode and applying a Hilbert transform.
2. Choose a mechanical proxy *σ*(*x, t*) provided by the model (cortical tension, strain, pressure, or an effective active-stress field) and extract its phase *ϕ*_*σ*_(*t*) on the same axis.
3. Identify neutral moments {*t*_*n*_} using the geometric condition *ψ*(*t*_*n*_) = *ϕ*_*V*_ (*t*_*n*_) − *ϕ*_*σ*_(*t*_*n*_) ≡ 0 (mod *π*) with 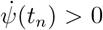, and compute the induced sequence of charge asymmetries

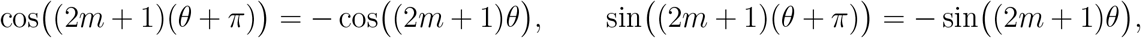
4. From the statistics of {Δ*Q*_*n*_ }, estimate the slow frequency *f*_slow_ from the spacing of neutral crossings and fit the effective contraction rate *κ* using the contraction relation in Eq. (46), obtaining a relaxation time *T*_relax_ ~ 1/(*f*_slow_*κ*).

Within this post-processing pipeline, P1, P2 and P3 become experimentally and numerically testable statements about the statistics and slow-time structure of {Δ*Q*_*n*_}, rather than purely geometric properties of the reduced phase description.

The dual-field theory predicts that healthy parameter sets should realize a P1-like regime (odd/even alternation with a contracting envelope and finite susceptibility), proliferative sets a P2-like regime (persistent or drifting neutral asymmetry), and senescent or overconstrained sets a P3-like regime (small but rigid neutral asymmetry with very fast local relaxation).

### L.1 Rotational neutral-map view of differentiation: 2/21 locking, charge relaxation and Prediction P4 on developmental timing

In the main text and in the previous parts of this appendix we described the dual-field clock in terms of neutral crossings, slow beating at *f*_slow_ and a neutral charge asymmetry sequence {Δ*Q*_*n*_} sampled at the neutral times *t*_*n*_. Here we show that the same construction admits a natural rotational interpretation in terms of circle maps and Arnold tongues, and how this neutral map organises slow differentiation trajectories and their bioelectric control.

At each neutral crossing *t*_*n*_ the phase difference between the electrical and mechanical carriers defines a slow angle *θ*_*n*_ ∈ *S*^1^.

For completeness, we sketch a minimal derivation of the neutral circle map used below. Let Θ_*n*_ ∈ [0, 2*π*) denote a 2*π*-lift of the slow angle at the *n*-th neutral crossing, so that *θ*_*n*_ = Θ_*n*_ (mod *π*). In full generality, a smooth, 2*π*-periodic phase update can be written as

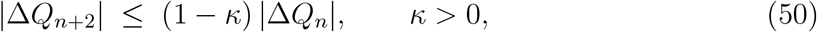

where *F* (Θ) is a 2*π*-periodic function describing the phase increment per neutral step. The neutral-strobing identification *θ* ≡ *θ* + *π* means that the reduced phase is only defined modulo *π*; hence the increment must satisfy

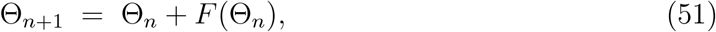

so that the dynamics is well defined on the quotient *θ* ∈ [0, *π*).

Expanding *F* in a Fourier series,

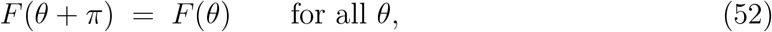

and imposing *F* (*θ* + *π*) = *F* (*θ*), we find that all odd harmonics must vanish, because

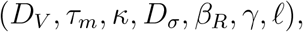

for every integer *m* ≥ 0. Thus *a*_2*m*+1_ = *b*_2*m*+1_ = 0 and only even harmonics survive:

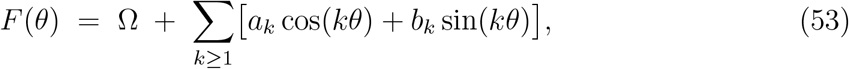

In the weak-coupling regime, the leading non-constant term is the first even harmonic (*m* = 1). After absorbing its phase into a redefinition of *θ*, we may, without loss of generality, keep only a pure sine term,

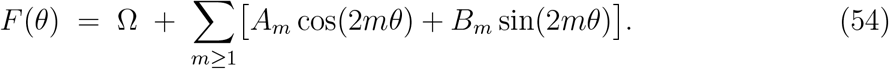

where *K*_2_ is an effective coupling constant that measures the strength of the neutral modulation. Substituting this into the phase update and working modulo *π* yields the neutral even circle map

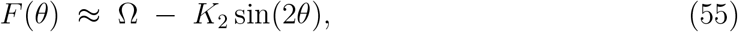

which provides the canonical form used below, with higher even harmonics representing subleading corrections beyond the minimal neutral description.

For fixed slow parameters (*g, F*), the neutral dynamics can therefore be approximated by a smooth, orientation-preserving map of the circle,

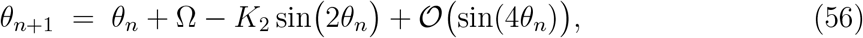

where Ω(*g, F*) is the intrinsic phase advance per neutral crossing in the absence of coupling, *K*_2_(*g, F*) is an effective even coupling amplitude and *g* ∈ [0, 1] is a slow differentiation coordinate.

#### Continuum origin of Ω(g, F) and K_2_(g, F)

At the continuum level, the overdamped electromechanical model (Eqs. (16)–(18)) is parametrised by

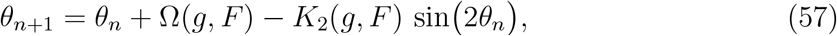

controlling electrical diffusion, membrane relaxation, cortical stiffness, mechanical diffusion, in-phase electromechanical gain, mechanical-to-electrical gain and cortical thickness. Restricting attention to the dominant spatial mode with wavenumber *k*_*_ = *π/*ℓ amounts to collecting these coefficients into three effective blocks,

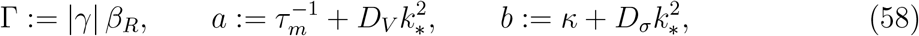

which summarise the active loop gain of the electromechanical coupler (Γ) and the effective electrical and mechanical dissipation seen by that mode (*a* and *b*).

The slow beat frequency *f*_slow_(Γ, *a, b*) introduced earlier is set by the balance between this active gain and the dissipative blocks. The dimensionless forcing Ω(*g, F*) in the neutral map then inherits its scale from that beat via a simple normalisation by the carrier frequency *f*_*c*_ (up to phase conventions), so that increasing Γ at fixed (*a, b, f*_*c*_) increases Ω, while increasing *a* or *b* at fixed (Γ, *f*_*c*_) reduces it.

The even coupling *K*_2_(*g, F*) measures how strongly neutral sampling biases the slow phase within one beat cycle. In the present framework it can be regarded as a smooth function of the same three blocks,

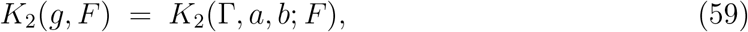

with the explicit dependence on *F* reflecting how the external drive and the slow differentiation coordinate reshape the electromechanical loop. In particular, *K*_2_ → 0 when Γ → 0 at fixed (*a, b*), reflecting the fact that no neutral bias can be generated without an active dual-field loop, and |*K*_2_| grows when the electrical and mechanical contributions are strongly unbalanced (for instance by changing *κ* at fixed *D*_*V*_, *D*_*σ*_). Thus both Ω(*g, F*) and *K*_2_(*g, F*) are determined by the same continuum parameters through (Γ, *a, b*), rather than being independent phenomenological knobs.

The long-time behaviour of Eq. (57) is encoded in the rotation number

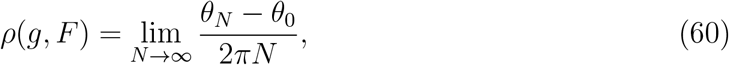

which measures the average phase advance per neutral crossing.

In general *ρ*(*g, F*) is irrational and the map exhibits a quasi-rotational drift of *θ*_*n*_, but on special parameter domains it locks to a rational value *ρ* = *m/p*, giving rise to periodic orbits of period *p*.

The regime of interest here corresponds to a plateau *ρ* = 2/21, consistent with the slow pattern observed in the P1 regime. In this case the neutral map admits a length-21 orbit {*θ*_0_, *θ*_1_, …, *θ*_20_} such that

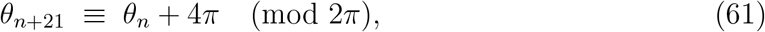

that is, two full turns of the slow phase difference after 21 neutral crossings. The corresponding Arnold tongue 𝒯_2/21_ is the region in the parameter plane (Ω, *K*_2_) for which such a period-21 orbit exists and is stable. Writing *T* for the right-hand side of Eq. (57), the linearization along the locked orbit has multiplier

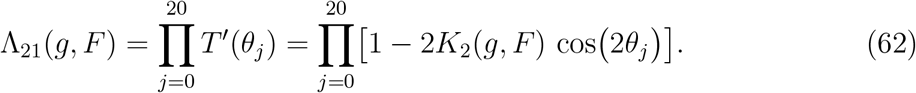

The tongue 𝒯_2/21_ is characterized by the existence of a period-21 orbit with |Λ_21_| *<* 1, while its boundary satisfies |Λ_21_| = 1.

In the charge-based language of App. L, the slow observable of interest is the envelope *R*_*n*_ of the neutral charge asymmetries Δ*Q*_*n*_, which in the locked P1 regime obeys approximately

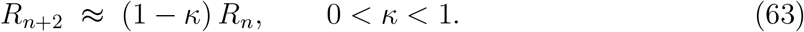

Over 21 neutral crossings the effect of this two-step contraction can be summarized as

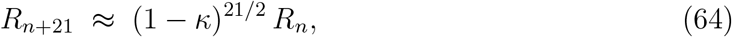

up to subleading corrections associated with the detailed pattern of odd–even crossings. Comparing this expression with the multiplier of the neutral map suggests the identification

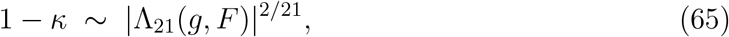

so that the slow contraction of the neutral charge envelope is directly linked to the linear stability of the 2/21 locked orbit. Inside the Arnold tongue, |Λ_21_| *<* 1 and hence 0 *< κ <* 1, giving a finite relaxation time *T*_relax_ ~ 1/(*f*_slow_*κ*) for the decay of |Δ*Q*_*n*_| .

On the tongue boundary, Λ_21_ = 1 and *κ* → 0, signalling the loss of effective contraction of neutral charge asymmetry.

Beyond the tongue, |Λ_21_| *>* 1 corresponds to *κ <* 0 in this effective description, so that the envelope of ||Δ*Q*_*n*_| grows and the system enters a proliferative P2-like regime.

From this rotational perspective, the locking ratio *ρ* = 2/21 does not merely label a geometric frequency plateau: it also counts the number of neutral crossings that compose one slow compensation cycle of the dual-field coupler.

A full 2/21 cycle involves twenty-one visits to the neutral window and two net rotations of the slow electromechanical phase, during which the sequence {Δ*Q*_*n*_} explores a characteristic odd–even pattern while its envelope contracts according to Eq. (65).

The slow differentiation coordinate *g*_*n*_ can be updated as a bounded functional of the neutral charge asymmetry,

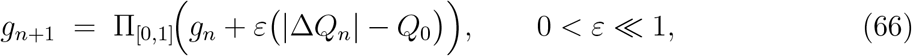

with Π_[0,1]_(*x*) = min {1, max {0, *x*}}, so that *g*_*n*_ remains confined to [0, 1] and high values of *g* correspond to proliferative or overlocked corridors.

In this minimal picture, the number of 2/21 cycles realised along a lineage before *g*_*n*_ leaves the healthy tongue 𝒯_2/21_ provides a discrete measure of how far the system has progressed along a differentiation corridor.

##### Linear origin of the differentiation map

The slow differentiation coordinate *g*_*n*_ can be anchored explicitly in the dual-field algebra. Let

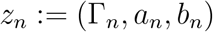

denote the three-block state of the overdamped electromechanical coupler at the *n*-th neutral crossing, and let *z*_*_ = (Γ_*_, *a*_*_, *b*_*_) be a homeostatic reference point in a P1-like corridor. We assume that *g*_*n*_ is a linear observable of *z*_*n*_,

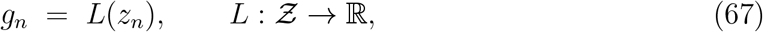

with Ƶ the state space of the three blocks.

Linearising the dual-field dynamics discussed in App. K around *z*_*_ gives

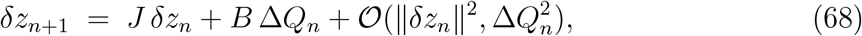

where *δz*_*n*_ := *z*_*n*_ − *z*_*_, *J* is the Jacobian of the neutral-step map at *z*_*_ and *B* encodes the susceptibility of (Γ, *a, b*) to the neutral charge asymmetry Δ*Q*_*n*_.

Applying *L* to Eq. (68) and using *δg*_*n*_ := *g*_*n*_ *g*_*_ = *L*(*dz*_*n*_), with *g*_*_ := *L*(*z*_*_), we obtain to leading order

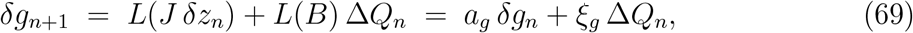

for suitable effective coefficients

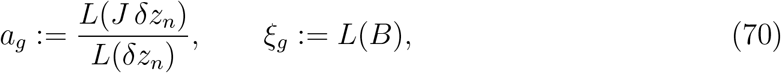

evaluated in the linear neighbourhood of *z*_*_.

Rewriting Eq. (69) in the relaxation form used in the main text amounts to identifying

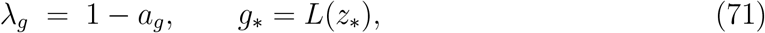

so that

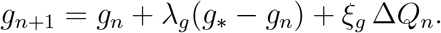

In this sense, the parameters (*λ*_*g*_, *ξ*_*g*_) appearing in the effective one-dimensional update for *g*_*n*_ are not free phenomenological coefficients, but functions of the Jacobian *J*, the charge-susceptibility operator *B* and the chosen observable *L* at the fixed point *z*_*_.

##### Slow precession and time scale hierarchy

In practice, locking is not perfectly rigid. As the lineage progresses, the slow differentiation parameter *g*_*n*_ drifts, introducing a small detuning

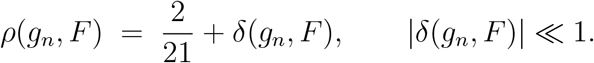

This causes the neutral orbit to undergo a slow precession: the sequence of crossings drifts relative to the ideal period-21 template. The time scale required to accumulate one full phase slip is

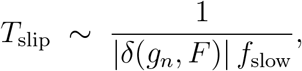

where *f*_slow_ is the slow electromechanical beat introduced in the main text. This mechanism is directly analogous to classical beating in acoustics or Moiré patterns in optics: two very close frequencies generate a much slower envelope.

Here, the small detuning *δ*(*g*_*n*_, *F*) produces a second-order slow harmonic. Defining the developmental frequency *f*_dev_ = 1*/T*_slip_, we obtain the hierarchical relation

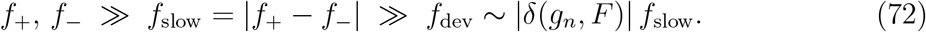

This explains how millisecond-scale bioelectric dynamics can govern developmental time scales of hours or days without arbitrary counters, purely through phase geometry.

Geometrically, the slow precession induced by a small detuning *ρ* = 2/21 + *δ* does not merely shift the phase difference *ψ*; it slowly rotates the two-dimensional parameter plane in which the neutral map is defined. In other words, what changes along the lineage is the orientation of the (*θ, g*) plane within the full dual-field state space. This orientation selects which combination of electrical and mechanical deformations is sampled at the neutral window, and thus ultimately determines the geometric imprint of each division and the resulting differentiation pathway.

##### Prediction P4 (Scaling of developmental time)

The rotational framework yields a specific quantitative scaling law. Empirically, simultaneous measurements of the slow electromechanical beat *f*_slow_ and of the macroscopic developmental timing *T*_dev_ ≈ *T*_slip_ must constrain the map detuning through

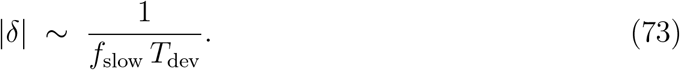

Equivalently, the number of slow compensation cycles required to advance one effective step along the differentiation corridor is predicted to scale as

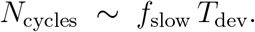

This provides a falsifiable link between the precessive correction to the ideal 2/21 locking and the observed separation of biological time scales. This scaling is intended as a local prediction within the P1-like corridor where |*δ*(*g*_*n*_, *F*) | ≪ 1 and *κ*(*g*_*n*_, *F*) *>* 0, so that the neutral circle map remains a smooth near-rotation.

##### Remark: Conjugacy with the helical scale

It is noteworthy that the 2/21 locking ratio has a natural conjugate in the molecular helical scale: its reciprocal satisfies

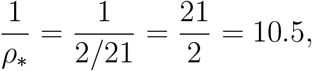

matching the canonical pitch of B-DNA (approximately 10.5 base pairs per turn). Formally, one may introduce a reference helical phase *χ*_*n*+1_ = *χ*_*n*_ + 2*πρ*_helix_ with *ρ*_helix_ := 1/10.5, and compare it to the neutral-map phase *θ*_*n*+1_ = *θ*_*n*_ + 2*πρ*, where *ρ* = 2/21 + *δ*. The relative phase between the electromechanical beat and the ideal helical pitch,

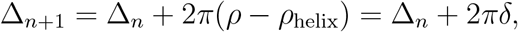

precesses at a rate set exactly by the detuning |*δ*| . Consequently, the number of neutral cycles required to accumulate one full “helical slip” scales as *N*_slip_ ~ 1/| *δ* |.

In this light, the scaling law of Prediction P4 can be read as a purely geometric measure of how close the tissue-scale electromechanical clock operates to the rational ratio conjugate to the molecular double-helix pitch; at this stage we only point out this numerical commensurability, without making a mechanistic claim about DNA itself.

##### Reprogramming as external control

Finally, the external control variable *F*_*n*_ (representing bioelectric stimulation or reprogramming factors) can be given a direct interpretation in this framework. The phase-entrainment strategy proposed in prediction P3 relies on weak, resonant stimulation at the slow beat frequency *f*_slow_ to modulate the neutral sector without overwhelming the fast carriers. In the rotational map, such a weak, coherent drive in the coupling region can be represented as a nonzero control field *F*_*n*_ that deforms the parameters

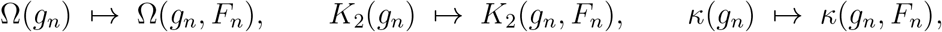

thereby moving the operating point (Ω, *K*_2_) in and out of the Arnold tongue 𝒯_2/21_. At the same time, *F*_*n*_ adds a weak drift term in the slow update of *g*_*n*_, which we can write as

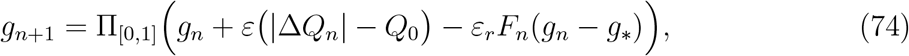

with 0 *< ε*_*r*_ ≪ 1 and a reference state *g*_*_ representing a progenitor-like corridor. In principle, a sustained, well tuned stimulation at *f*_slow_ could act not only as a stabilizing rescue of an unlocked state, but also as an artificial reprogramming field capable of steering the slow differentiation coordinate *g*_*n*_ along specific corridors of the neutral map.

From this perspective, the neutral circle map itself is part of the healthy architecture: it requires reasonably regular neutral crossings and a well-defined phase difference between electrical and mechanical carriers.

In senescent regimes (P3), the map persists but collapses onto a strongly attracting almost-periodic orbit with very small detuning |*δ*| and large contraction *κ*, so that precession becomes effectively negligible on physiological time scales even though small desynchronizing excursions may still occur.

In proliferative P2-like regimes, by contrast, neutral crossings tend to become irregular or rare and the sequence {*θ*_*n*_} no longer follows a smooth near-rotation, so that the rotational reduction itself ceases to be a faithful description of the dynamics. In this

sense, the loss of a robust neutral-map representation is a natural part of the geometric signature of pathological unlocking, whereas senescence corresponds to an overly rigid, weakly responsive version of the same geometric clock.

At this stage, Prediction P4 has only been assessed at the level of reduced models rather than confronted with real bioelectric or lineage-tracking data. Its time-scale implications are nevertheless numerically consistent with known orders of magnitude. For a slow bioelectric beat in the range *f*_slow_ ~ 0.05 Hz (period ~ 20 s) and small detunings |*δ* | ~ 10^−2^ − 10^−4^, the relation

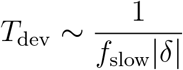

yields effective developmental times spanning from tens of minutes to several hours and days, comparable to reported mitotic and differentiation time scales in mammalian cells.

To illustrate the internal consistency of this scaling, we performed a numerical integration of a reduced phenotypic dynamics, where a coarse-grained coordinate *h*_*n*_ evolves according to the drift update *h*_*n*+1_ = *h*_*n*_ + *η δ* and is monitored until it crosses a fixed differentiation threshold |*h*_*n*_| ≥*h*_th_. For a set of representative detunings *δ* in the regime of interest, we recorded the number of neutral steps 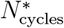 required to reach the threshold. The results show that 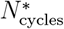 scales linearly with the inverse detuning 1/| *δ*| (Fig. L.2),

consistent with the prediction that the macroscopic law *T*_dev_ ~ 1/(*f*_slow_ |*δ* |) arises from the

accumulation of small phase slips (see Listing M.10 for implementation details).

**Figure L.2:**
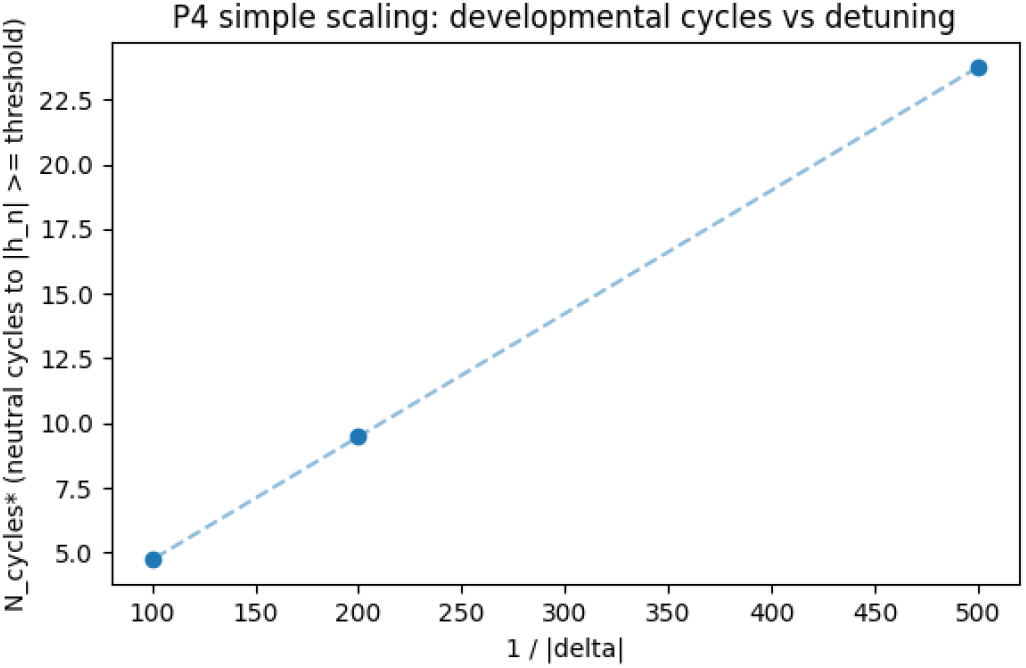
Numerical verification of Prediction P4 in a minimal drift model. In a reduced one-dimensional drift model governed by *h*_*n*+1_ = *h*_*n*_ + *η δ* with a fixed threshold |*h*_*n*_| ≥ *h*_th_, the number of neutral steps required to reach the threshold, 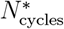, scales linearly with 1*/ δ*, in agreement with the scaling law *T*_dev_ ~ 1/(*f*_slow_ |*δ*|) implied by the precessional mechanism.

##### Neutral windows and helical replication as a phase-driven process

The conjugacy between the 2/21 locking ratio and the canonical B-DNA pitch suggests that the algebraic structure developed for the dual-field clock can be lifted from the cellular to the genetic level. Rather than modelling the biochemical architecture of DNA or RNA, we treat replication and transcription as phase-gated processes subordinated to the same slow clock that governs neutral crossings. In this view, the helical machinery acquires a dynamical algebraic layer—absent from purely kinetic or structural descriptions—that links molecular timing to the macroscopic electromechanical beat and provides a process-level bridge between genetic regulation and tissue-scale dynamics.

We introduce two conjugate phases. The neutral-map dynamics is encoded in the slow angle *θ*_*n*_ ∈ *S*^1^ at the *n*-th neutral crossing,

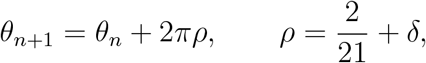

while the advance of a reference helical frame is described by

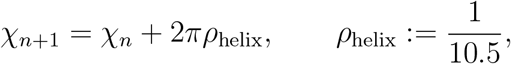

so that 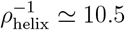 base pairs per turn matches the canonical B-DNA pitch. The relative phase between the neutral map and the ideal helical pitch,

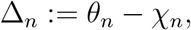

then evolves as

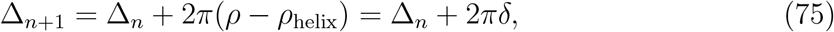

so that small detunings |*δ*| ≪ 1 produce a slow precession of Δ_*n*_.

To anchor this construction in a bioelectric observable, we recall the neutral charge asymmetry Δ*Q*_*n*_ introduced in App. L. At each neutral crossing, Δ*Q*_*n*_ samples the antisymmetric component of the membrane-potential profile across the division axis and thus acts as a proxy for how much the dual-field coupler is locally biased. In the P1 regime, the envelope of |Δ*Q*_*n*_| contracts slowly under the neutral map, while in P2 it drifts or grows and in P3 it collapses towards a rigid narrow distribution.

Replication and transcription are modelled, at this coarse-grained level, as phase-gated processes that advance preferentially when the electromechanical and helical frames are in a favourable relative configuration and the neutral sector is weakly charged. We encode this by a 2*π*-periodic gating function *G*(Δ, Δ*Q*) ∈ [0, 1], with *G* sharply peaked in a narrow window around Δ = 0 mod 2*π* and damped whenever |Δ*Q*_*n*_| is large. A minimal form is

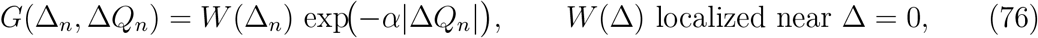

with *α >* 0 setting how strongly neutral charge asymmetry suppresses the gate. The exact shape of *W* depends on the underlying molecular machinery and is not specified here; only its localisation in phase and its modulation by Δ*Q*_*n*_ are assumed.

Let *r*_*n*_ denote a coarse-grained replication/transcription coordinate (for instance, a normalized progress variable along a given locus or an effective copy number). Its neutral-step dynamics is then written as

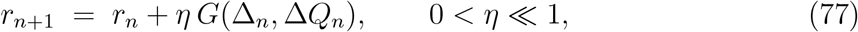

where *η* is an effective increment per favourable phase encounter. Combining Eqs. (75), (76), and (77) yields a skew-product dynamics on *S*^1^ × ℝ driven by the neutral map and filtered by the charge sector:

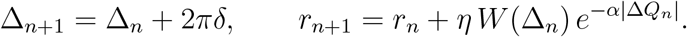

In a first approximation, and for |*δ*| ≪ 1, the relative phase Δ_*n*_ explores the circle slowly and almost uniformly, while the envelope of |Δ*Q*_*n*_| evolves according to the neutral-map contraction rate *κ*(*g, F*) discussed above. If the tissue operates in a P1-like regime with 0 *< κ <* 1, the neutral charge asymmetry is progressively reduced and the exponential factor in Eq. (76) tends towards unity, so that the long-term average increment per neutral step is

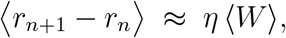

with ⟨*W*⟩ the phase-average of *W* over one precessional cycle. The time required to sweep once around the relative phase is set by the detuning,

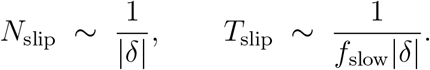

If a full replication or transcription event requires a net increment *r*_final_ − *r*_initial_ = Δ*r*_*_, the characteristic completion time in a P1-like environment scales as

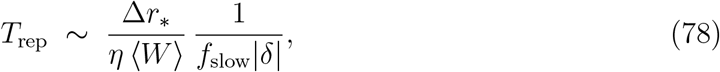

so that, up to prefactors depending on the gate profile and efficiency, the scaling *T*_rep_ ∝ ~ 1/(*f*_slow_ *d*) mirrors the developmental law *T*_dev_ ∼ 1/(*f*_slow_ |*δ*|) of Prediction P4. In this formulation, the same precessional detuning that controls tissue-level phenotypic transitions also sets the coarse-grained timing of replication and transcription bursts whenever the neutral charge sector remains contractive.

Finally, the P2 and P3 regimes acquire a direct process-level interpretation in this language.

In a proliferative P2-like unlocking, the neutral-map contraction fails (*κ* ≤ 0), the envelope of |Δ*Q*_*n*_| remains broad, and the factor 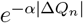 fluctuates strongly across neutral crossings. This produces irregular gating, long-lived imbalances between complementary strands, and a loss of reproducible scaling in Eq. (78).

In an overlocked P3-like regime, the neutral phase remains near a fixed point of the map, Δ_*n*_ explores only a narrow arc of *S*^1^, and *W* (Δ_*n*_) becomes effectively frozen.

Replication and transcription then occur in a narrow, rigid phase corridor with reduced dynamic range, consistent with a senescence-like loss of responsiveness rather than with proliferative drift. In both cases, the dual-field algebra acts not on the structure of the double helix, but on the timing and regularity of the gates through which its molecular machinery is effectively allowed to operate.

## M Supplementary Code Listings (reproducibility)

### M.1 Dual-field pseudo-spectral sandbox and analysis pipeline

Listing 1 implements the overdamped, first-order sandbox t hat operationalizes P 1–P2 and provides preregisterable targets for P3.

It links measurable electromechanical parameters to slow-phase locking and coherence, and returns the observables *f*_*±*_, *f*_slow_, Λ, and the neutral-moment fit 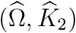.

A dedicated P3 script and off-band controls are provided in App. G; cases A-D are illustrated in Figs. 11.Figure 1–11.5

~~~
*# Listing 1. Overdamped, first - order sandbox for P1 -P2 and*
    *preregistered P3 targets* .
*# Implements the continuum electromechanical coupler (Appendix B)*,
*# extracts f_minus, f_plus, f_slow, Lambda and the neutral - moment*
    *fit*
*# (Omega_hat, K2_hat), and saves two basic diagnostics in SAVE_DIR*
   :
*# (i) coherence spectrum with f_slow marked, and*
*# (ii) selected - mode time series for V and sigma* .
*# These plots are sanity checks for the pipeline and are not used*
     *as figures*
*# in the main text* .
       import os, csv
       import numpy as np
       from numpy .fft import rfftn, rfftfreq
       from scipy . signal import welch, csd, hilbert
       from sklearn . linear_model import LinearRegression
       import matplotlib as mpl; mpl . rcdefaults ()
       import matplotlib . pyplot as plt
       # ----------    *output folder* ----------
       SAVE_DIR = “./ s0_outputs “
       os. makedirs (SAVE_DIR, exist_ok = True)
       # ----------    *CONFIG* ----------
       Nx = 128; Ny = 128
       Lx = 1.0; Ly = 1.0
       dt = 0.01
       T = 360.0
       steps = int(T/dt)
       sample_every = 1
       fs = 1.0/ dt
       D_V = 5e -9
       tau_m = 0.5
       kappa = 6e -4
       eta = 5e -4
       Dsig = 5e -11
       betaR = 0.35
       gamma = 0.25
       betaI = 0.02
       alpha_eff = 1.0 - betaI * gamma # ~0.995
       noise_V = 1e -9
       noise_u = 1e -9
       rng = np. random . default_rng (1)
       init_amp = 1e -3
       init_mode = (1, 1)
       scan_max_mode = (3, 3)
       welch_nperseg_max = 4096
       coherence_band = (0.05, 2.0) # Hz
       # ----------    *k-grid* ----------
       kx = rfftfreq (Nx, d=Lx/Nx) * 2* np.pi
       ky = np.fft. fftfreq (Ny, d=Ly/Ny) * 2* np.pi
       KX, KY = np. meshgrid (kx, ky, indexing =“xy”)
       k2 = (KX **2 + KY **2) . astype (np. float32)
       # ----------    *initial conditions* ----------
       yy = np. arange (Ny)[:, None ]
       xx = np. arange (Nx)[None, :]
       seed_mode = (np.sin (2* np.pi*yy/Ny) * np.sin (2* np.pi*xx/Nx)). astype
         (np. float32)
      
  V0 = (rng. standard_normal ((Ny, Nx)). astype (np. float64) * 1e -4 +
         init_amp * seed_mode)
  
  u0 = (rng. standard_normal ((Ny, Nx)). astype (np. float64) * 1e -4 +
         0.7 * init_amp * seed_mode)
  
  Vhat = rfftn (V0, axes =(0,1)). astype (np. complex64)
  Uhat = rfftn (u0, axes =(0,1)). astype (np. complex64)
       # ----------    *IMEX denominators* ----------
       denV = (1.0 + (dt/ alpha_eff) * (tau_m ** -1 + D_V * k2)). astype (np.
  float64)
denU = (1.0 + (dt/ eta) * (kappa + Dsig * k2)). astype (np.
  float64)
def noise_hat (scale, shape, rng):
  nh = np. zeros (shape, dtype =np. complex64)
  max_ix = min (2, shape [1])
  for ix in range (max_ix):
  nh [0, ix] = scale * (rng. standard_normal () + 1j*rng.
  standard_normal ())
  return nh
# ----------    *traces for candidate modes* ----------
candidates = [(j, i) for j in range (0, scan_max_mode [0]+1)
  for i in range (0, scan_max_mode [1]+1)
  if not (j == 0 and i == 0)]
traceV = { pair : [] for pair in candidates }
traceS = { pair : [] for pair in candidates }
# ----------    *time loop* ----------
for n in range (steps):
  Uhat_new = (Uhat + (dt/eta) * (gamma * Vhat + noise_hat (
  noise_u, Uhat .shape, rng))) / denU
  Vhat_new = (Vhat + (dt/ alpha_eff) * (betaR * gamma * Vhat +
  noise_hat (noise_V, Vhat .shape, rng))) / denV
  Sigma_hat = kappa * Uhat_new + eta *(Uhat_new - Uhat)/dt + Dsig *
  k2* Uhat_new
if n % sample_every == 0:
   for (kyi, kxi) in candidates :
   traceV [(kyi,kxi)]. append ((Vhat_new [kyi, kxi ]. real / (
   Nx*Ny)))
   traceS [(kyi,kxi)]. append ((Sigma_hat [kyi, kxi ]. real / (
   Nx*Ny)))
Vhat, Uhat = Vhat_new, Uhat_new
# ----------    *spectral helpers* ----------
def two_carriers (f, S, band =(0.05, 2.0)):
  m = (f >= band [0]) & (f <= band [1])
  if np. count_nonzero (m) < 3:
  return np.nan, np.nan, np.nan
  idx_sorted = np. argsort (S[m])
  if idx_sorted . size < 2:
  return np.nan, np.nan, np.nan
  fp = np. sort (f[m][ idx_sorted [ -2:]]) # top -2 peaks
  return float (fp [0]), float (fp [1]), float (fp [1] - fp [0])
def coherence_at_fslow (Vsig, Ssig, fs, nperseg, fslow):
  if not np. isfinite (fslow) or fslow <= 0:
  return np.nan
   fC, SVS = csd(Vsig, Ssig, fs=fs, nperseg = nperseg)
  fV, SV = welch (Vsig, fs=fs, nperseg = nperseg)
  fS, SS = welch (Ssig, fs=fs, nperseg = nperseg)
  
  SVi = np. interp (fC, fV, SV)
  SSi = np. interp (fC, fS, SS)
  
  Lambda_f = (np. abs(SVS) **2) / (SVi*SSi + 1e -20)
  return float (np. interp (fslow, fC, np. real (Lambda_f)))
# ----------    *scan all modes* ----------
summary_rows = []
best = None
best_L = -np.inf
for pair in candidates :
   Vsig = np. asarray (traceV [ pair ], dtype = float)
   Ssig = np. asarray (traceS [ pair ], dtype = float)
   if len(Vsig) < 256:
   continue
   nper = int(min (welch_nperseg_max, len(Vsig)))
   fV, SV = welch (Vsig, fs=fs/ sample_every, nperseg = nper)
   f_minus, f_plus, f_slow = two_carriers (fV, SV, band =
   coherence_band)
   L = coherence_at_fslow (Vsig, Ssig, fs=fs/ sample_every, nperseg
   =nper, fslow = f_slow)
   summary_rows . append ([ pair [0], pair [1], f_minus, f_plus, f_slow
 , L, len(Vsig), nper ])
   if np. isfinite (L) and L > best_L :
   best_L = L
   best = pair
*# Fallback by power if needed*
if best is None :
  best, topP = None, -np.inf
  for pair in candidates :
  sig = np. asarray (traceV [ pair ], dtype = float)
  if len(sig) < 256:
  continue
nper = int(min (welch_nperseg_max, len(sig)))
f, S = welch (sig, fs=fs/ sample_every, nperseg = nper)
mask = (f >= coherence_band [0]) & (f <= coherence_band [1])
val = float (np.max (S[ mask ])) if np.any(mask) else -np.inf
if val > topP :
  topP, best = val, pair
*# ---------- final spectra and Lambda (f) for selected mode*
  ----------
Vsig = np. asarray (traceV [ best ], dtype = float)
Ssig = np. asarray (traceS [ best ], dtype = float)
nper = int(min (welch_nperseg_max, len(Vsig)))
fV, SV = welch (Vsig, fs=fs/ sample_every, nperseg = nper)
fS, SS = welch (Ssig, fs=fs/ sample_every, nperseg = nper)
fC, SVS = csd(Vsig, Ssig, fs=fs/ sample_every, nperseg = nper)
SVi = np. interp (fC, fV, SV)
SSi = np. interp (fC, fS, SS)
Lambda_f = (np.abs(SVS) **2) / (SVi*SSi + 1e -20)
f_minus, f_plus, f_slow = two_carriers (fV, SV, band = coherence_band
 )
Lambda_at = float (np. interp (f_slow, fC, np. real (Lambda_f))) if np.
  isfinite (f_slow) else np.nan
*# ----------neutral strobe and even -map fit (report K2 > 0)*
  ----------
ZV = hilbert (Vsig)
ZS = hilbert (Ssig)
phiV = np. unwrap (np. angle (ZV))
psi = np. unwrap (np. angle (ZV) - np. angle (ZS))
tol = 0.10
psi_mod = np.mod (psi, np.pi)
psi_mod = np. minimum (psi_mod, np.pi - psi_mod)
psidot = np. gradient (psi, dt* sample_every)
idx_neutral = np. where ((psi_mod < tol) & (psidot > 0))[0]
if len(idx_neutral) > 2:
  theta = phiV [ idx_neutral ]
  dtheta = np. diff (theta)
  if len(dtheta) > 10:
  X = np. sin (2* theta [: -1]) . reshape (-1,1)
  reg = LinearRegression (). fit(X, dtheta)
  Omega_hat = float (reg. intercept_)
  K2_hat = float (abs(reg. coef_ [0])) # report positive K2
else :
  Omega_hat = np.nan; K2_hat = np.nan
else :
  Omega_hat = np.nan; K2_hat = np.nan
*# ---------- SAVE* :    *summary of all modes ----------*
summary_csv = os. path . join (SAVE_DIR, “s0_modes_summary . csv”)
with open (summary_csv, “w”, newline =““) as fh:
  w = csv. writer (fh)
  w. writerow ([“ky”,”kx”,” f_minus “,” f_plus “,” f_slow “,”
  Lambda_at_fslow “,” len_trace “,” nperseg “])
  w. writerows (summary_rows)
*# ----------SAVE* :    *selected mode time series ----------*
ts_csv = os. path . join (SAVE_DIR, “s0_selected_mode_timeseries .csv”)
t = np. arange (len(Vsig)) * dt * sample_every
with open (ts_csv, “w”, newline =““) as fh:
  w = csv. writer (fh); w. writerow ([“t”,”V”,” sigma “])
  for ti, vi, si in zip(t, Vsig, Ssig):
  w. writerow ([ti, vi, si ])
*# ----------SAVE* :    *Lambda (f) spectrum ----------*
lambda_csv = os. path . join (SAVE_DIR, “s0_lambda_spectrum .csv”)
with open (lambda_csv, “w”, newline =““) as fh:
  w = csv. writer (fh); w. writerow ([“f”,” Lambda_f “])
  for fi, Li in zip (fC, np. real (Lambda_f)):
  w. writerow ([ float (fi), float (Li)])
*# ----------PLOTS* :    *publication - ready ----------*
plt . figure ()
plt . plot (t, Vsig, lw =1.2, label =“V␣(selected ␣ mode)”)
plt . plot (t, Ssig, lw =1.0, alpha =0.75, label =“sigma ␣(selected ␣ mode)
  “)
plt . xlabel (“time ␣(s)”); plt. ylabel (“amplitude ␣(a.u.)”)
plt . title (“Selected - mode ␣ time ␣ series “); plt . legend (); plt.
   tight_layout ()
png_ts = os. path . join (SAVE_DIR, “s0_selected_mode_timeseries .png”)
plt . savefig (png_ts, dpi =300) ; plt. show ()
plt . figure ()
yplot = np. maximum (np. real (Lambda_f), 1e -16) # numeric floor for
   semilogy
plt . semilogy (fC, yplot, lw =1.2)
if np. isfinite (f_slow): plt. axvline (f_slow, ls=“--”, alpha =0.7)
plt . xlim (0, 2.0)
plt . xlabel (“frequency ␣(Hz)”); plt. ylabel (“Lambda (f)”)
plt . title (“Coherence ␣ spectrum ␣ and␣ f_slow “); plt. tight_layout ()
 png_lambda = os. path . join (SAVE_DIR, “s0_lambda_spectrum .png”)
 plt . savefig (png_lambda, dpi =300) ; plt. show ()
 *# ---------- print summary ----------*
  summary_dict = {
  “selected_mode “: best,
  “alpha_eff “: float (alpha_eff),
  “f_minus “: f_minus,
  “f_plus “: f_plus,
  “f_slow “: f_slow,
  “Lambda_at_fslow “: Lambda_at,
  “Omega_hat “: Omega_hat,
  “K2_hat “: K2_hat,
  “len_trace “: int(len(Vsig)),
  “nperseg “: int(nper),
  “saved_to “: os. path . abspath (SAVE_DIR)
}
print (summary_dict)
~~~

### M.2 Level-0 analysis script (public epithelial movie)

~~~
*#!/ usr /bin/ env python3*
*# -*- coding* :    *utf -8 -*-*
“““
Listing ␣2.␣Level -0␣ analysis ␣ pipeline ␣on␣ publicly ␣ available ␣
  epithelial ␣ voltage - imaging ␣ data .
This ␣ script ␣ reproduces ␣ the␣Level -0␣ validation ␣in␣the␣ manuscript .
Input :
␣␣␣␣-␣A␣high - speed ␣wide - field ␣ voltage - imaging ␣ movie ␣of␣a␣ confluent
  ␣ epithelial (- like)
␣␣␣␣␣␣ monolayer ␣ from ␣ Quicke ␣et␣al.␣ (2022),␣ recorded ␣at␣~5␣Hz␣for␣
  ~500 ␣s
␣␣␣␣␣␣ (~2500 ␣ frames).␣The␣ movie ␣is␣ provided ␣as␣a␣ supplemental ␣.mov
  ␣ file .
␣␣␣␣␣␣If␣ quicke2022_movie .mov ␣is␣ not␣ present,␣the ␣ script ␣ prompts ␣
  for ␣ upload ␣of␣the␣ Quicke ␣et␣al.␣ supplemental ␣.mov␣ file ␣and␣ then
  ␣ runs ␣the ␣ analysis .
Output :
␣␣␣␣-␣A␣ single ␣ROI - averaged,␣ detrended,␣z-scored ␣ trace ␣V(t)␣
  interpreted ␣as␣a␣ local
␣␣␣␣␣␣ transmembrane ␣ voltage ␣ proxy .
␣␣␣␣-␣ Welch ␣ power ␣ spectrum ␣of␣V(t),␣ identification ␣of␣ two␣
  narrowband ␣ carrier ␣ peaks
␣␣␣␣␣␣ f_plus ␣ and␣ f_minus ␣in␣the␣~1- -2␣Hz␣band,␣and ␣the␣ slow ␣ beat
␣␣␣␣␣␣ f_slow ␣=␣| f_plus ␣-␣ f_minus |.
␣␣␣␣-␣ Hilbert ␣ amplitude ␣ envelope ␣of␣V(t)␣and␣its ␣ Welch ␣ spectrum,␣
  including ␣the
␣␣␣␣␣␣ enrichment ␣ ratio ␣of␣the ␣ slow ␣ envelope ␣ peak ␣ near ␣ f_slow .
␣␣␣␣-␣ Four ␣ publication ␣ panels ␣(A--D),␣ saved ␣as
␣␣␣␣␣␣ figs / A1_timeseries .png,
␣␣␣␣␣␣ figs / A1_psd_signal .png,
␣␣␣␣␣␣ figs / A1_envelope_timeseries .png,
␣␣␣␣␣␣ figs / A1_psd_envelope .png,
␣␣␣␣␣␣ reproducing ␣the␣Level -0␣ carrier ␣and␣ envelope ␣ analysis ␣
  described ␣in␣ the␣ main ␣ text .
All ␣ numeric ␣ outputs ␣(sampling ␣rate,␣ frame ␣count,␣ROI,␣f_plus,␣
  f_minus,␣ f_slow,
and ␣ enrichment_ratio)␣are ␣ printed ␣to␣ stdout .
“““
import os
import numpy as np
import matplotlib . pyplot as plt
import imageio .v2 as imageio
import cv2
from scipy . signal import detrend, welch, hilbert
# ----------------------------------------------------------
*# User configuration (frozen for manuscript reproducibility)*
# ----------------------------------------------------------
VIDEO_PATH = “quicke2022_movie .mov” # supplemental movie
     *from Quicke et al. (2022)*
FPS_OVERRIDE = 5.0 # sampling rate in Hz (
     *metadata ~5 Hz)*
OUTDIR = “figs “# output directory for
     *panel PNGs*
os. makedirs (OUTDIR, exist_ok = True)
*# Rectangular ROI fully inside the confluent monolayer* .
*# Coordinates are [x0, y0, x1, y1] in pixel units of the movie*
     *frames* .
*# These are the exact bounds reported in the text* :    *x =[309*,*617], y*
  =[151,301]
FIXED_ROI = [309, 151, 617, 301]
*# Spectral - analysis parameters (Level -0 preregistration)*
FAST_BAND = (1.0, 2.3) #    *Hz, expected carrier band*
PROMINENCE_FACTOR = 2.0    *# relative peak threshold in*
  FAST_BAND
PEAK_DISTANCE_BINS = 3    *# minimum frequency -bin*
  separation
SMOOTHING_WINDOW = 0    *# temporal moving - average*
  window (frames); 0 disables
MAX_FRAMES = 2500    *# analyze first ~2500 frames*
  (~500 s @ 5 Hz)
# ------------------------------------------------
*# Colab - style file upload if the movie is missing*
# ------------------------------------------------
if not os. path . exists (VIDEO_PATH):
  try :
     *# This branch is intended for Google Colab* ;    *there the*
     *upload widget*
     *# with a progress bar is available* .
  from google . colab import files
except ImportError :
*# Outside Colab, raise a clear error and ask the user to*
     *edit VIDEO_PATH manually* .
  raise FileNotFoundError (
  f” Movie ␣ file ␣ ‘{ VIDEO_PATH }’␣not␣ found .\n”
  “Place ␣the ␣ Quicke ␣et␣al.␣ supplemental ␣.mov␣in␣ this ␣
  folder,␣”
  “or␣ edit ␣ VIDEO_PATH ␣at␣the␣top ␣of␣ the␣ script ␣to␣ point ␣
to␣ its␣ location .”
)
print (
  “Movie ␣ file ␣ not␣ found .\n”
  “Please ␣ upload ␣the␣ Quicke ␣et␣al.␣ supplemental ␣.mov ␣ file ␣”
  “(e.g.␣’ quicke2022_movie .mov ‘).”
)
  uploaded = files . upload () # Triggers the file - upload widget
  with a progress indicator in Colab .
if not uploaded :
  raise FileNotFoundError (“No␣ movie ␣ uploaded ;␣ aborting .”)
# Use the first uploaded file as the movie path .
VIDEO_PATH = list (uploaded . keys ())[0]
print (f” Using ␣ uploaded ␣ movie ␣ file :␣{ VIDEO_PATH }”)
# --------------------------------------------
*# ROI extraction and time - series construction*
# --------------------------------------------
def extract_timeseries_from_video (path,
  roi,
  fps_override =5.0,
  denoise_ma =0,
  normalize_drift =True,
  max_frames = None):
  “““
␣␣␣␣ Extract ␣a␣ single ␣ scalar ␣ time ␣ series ␣V(t)␣ from ␣a␣ rectangular ␣
  region ␣of␣ interest
  ␣␣␣␣(ROI)␣in␣a␣wide - field ␣ voltage - imaging ␣ movie .
␣␣␣␣”““
  rdr = imageio . get_reader (path)
  # Inspect first frame to infer geometry
  first = rdr. get_data (0)
  height, width = first . shape [:2]
  x0, y0, x1, y1 = roi
  x0 = max (0, x0); y0 = max (0, y0)
  x1 = min(width, x1); y1 = min(height, y1)
*# Optional frame limit*
n_total = None
if hasattr (rdr, “count_frames “):
  try :
    n_total = rdr. count_frames ()
  except Exception :
    n_total = None
frame_limit = n_total if max_frames is None else max_frames
if frame_limit is None :
  frame_limit = 10**9 # effectively no limit
vals = []
for idx, frame in enumerate (rdr):
  if idx >= frame_limit :
     break
  # Convert to grayscale
  if frame . ndim == 3:
     gray = cv2. cvtColor (frame, cv2. COLOR_RGB2GRAY)
  else :
     gray = frame
  roi_img = gray [y0:y1, x0:x1]
  vals . append (float (np. mean (roi_img)))
rdr . close ()
fs = float (fps_override)
vals = np. asarray (vals, dtype = float)
*# Optional temporal denoising by moving average*
if denoise_ma and denoise_ma > 1:
  k = int(denoise_ma)
  vals = np. convolve (vals, np. ones (k)/k, mode =“same “)
*# Remove slow drift (e*.*g. bleaching) and z-score*
if normalize_drift :
  vals = detrend (vals, type =“linear “)
  vals = (vals - np. mean (vals)) / (np.std(vals) + 1e -12)
t = np. arange (len(vals)) / fs
  meta = dict (
  roi =[x0, y0, x1, y1],
  width =width,
  height =height,
  nframes =len(vals),
  fs_used =fs,
)
return t, vals, fs, meta
# ---------------------------
*# Level -0 spectral analysis*
# ---------------------------
def analyze_timeseries_level0 (t, x, fs,
  fast_band =(1.0, 2.3),
  prominence_factor =2.0,
  peak_distance_bins =3):
“““
␣␣␣␣ Perform ␣ the␣ preregistered ␣Level -0␣ analysis ␣on␣a␣ single - channel
  ␣ trace ␣V(t).
␣␣␣␣”““
  nperseg = min(len (x), 4096)
*# PSD of V(t)*
f, Pxx = welch (x, fs=fs, nperseg = nperseg)
*# Peak - finding in the fast band*
bandmask = (f >= fast_band [0]) & (f <= fast_band [1])
fband = f[ bandmask ]
Pband = Pxx[ bandmask ]
thr = np. median (Pband) * prominence_factor
candidate_idxs = []
for i in range (1, len(fband) - 1):
     *# local maximum above threshold ?*
  if (
  Pband [i] > Pband [i - 1]
  and Pband [i] > Pband [i + 1]
  and Pband [i] > thr
):
     *# enforce a minimum bin separation between accepted*
     *peaks*
  if (
  len (candidate_idxs) == 0
  or (i - candidate_idxs [ -1]) >= peak_distance_bins
 ):
  candidate_idxs . append (i)
*# Sort by peak height, descending*
peak_order = sorted (
  candidate_idxs,
  key = lambda k: Pband [k],
  reverse = True
)
f_plus = np.nan
f_minus = np.nan
if len(peak_order) >= 1:
  f_plus = fband [ peak_order [0]]
if len(peak_order) >= 2:
f_minus = fband [ peak_order [1]]
f_slow = np.nan
if (not np. isnan (f_plus)) and (not np. isnan (f_minus)):
  f_slow = abs (f_plus - f_minus)
*# Hilbert envelope of V(t)*
env = np.abs(hilbert (x))
f_env, P_env = welch (env, fs=fs, nperseg = nperseg)
# Enrichment of the slow - beat envelope peak
enrichment_ratio = np. nan
if not np. isnan (f_slow):
  win = 0.2    *# +/- 20 %*
m = (f_env >= (1 - win) * f_slow) & (f_env <= (1 + win) *
  f_slow)
if np.any(m):
  peak_env = np.max(P_env [m])
  base_env = np. median (P_env)
  enrichment_ratio = peak_env / (base_env + 1e -15)
return {
  “t”: t,
  “x”: x,
  “fs”: fs,
  “f”: f,
  “Pxx”: Pxx,
  “f_env “: f_env,
  “P_env “: P_env,
  “env_signal “: env,
  “f_plus “: f_plus,
  “f_minus “: f_minus,
  “f_slow “: f_slow,
  “enrichment_ratio “: enrichment_ratio,
}
# ------------------------------------
*# Panel generation* :    *save and display*
# ------------------------------------
def save_and_show_panels (res, outdir):
  “““
␣␣␣␣ Generate ␣ the␣ four ␣ publication ␣ panels ␣(A--D).
␣␣␣␣”““
  t = res[“t”]
  x = res[“x”]
  f = res[“f”]
  Pxx = res[“Pxx “]
  f_env = res[“f_env “]
  P_env = res[“P_env “]
  env = res[“env_signal “]
  f_plus = res [“f_plus “]
  f_minus = res[“f_minus “]
  f_slow = res [“f_slow “]
*# Panel A: time series V(t)*
plt . figure (figsize =(6, 2))
plt . plot (t, x, lw =0.7)
plt . xlabel (“time ␣(s)”)
plt . ylabel (“z-score ␣(a.u.)”)
plt . title (“ROI - averaged ␣V(t)”)
plt . tight_layout ()
outA = os. path . join (outdir, “A1_timeseries .png”)
plt . savefig (outA, dpi =300)
plt . show ()
*# Panel B: PSD[V]*
plt . figure (figsize =(6, 2))
plt . semilogy (f, Pxx, lw =0.7, label =“PSD [V]”)
if not np. isnan (f_plus):
  plt . axvline (
  f_plus,
  ls=“--”,
  color =“C1”,
  label =f” f_plus ␣~␣{ f_plus :.2 f}␣Hz”
)
if not np. isnan (f_minus):
  plt . axvline (
  f_minus,
  ls=“--”,
  color =“C2”,
  label =f” f_minus ␣~␣{ f_minus :.2 f}␣Hz”
)
plt . xlim (0, 2.5)
plt . xlabel (“frequency ␣(Hz)”)
plt . ylabel (“power “)
plt . title (“PSD␣of␣V(t)”)
plt . legend (fontsize =6)
plt . tight_layout ()
outB = os. path . join (outdir, “A1_psd_signal .png”)
plt . savefig (outB, dpi =300)
plt . show ()
*# Panel C: Hilbert envelope in time*
plt . figure (figsize =(6, 2))
plt . plot (t, env, lw =0.7)
plt . xlabel (“time ␣(s)”)
plt . ylabel (“amplitude “)
plt . title (“Hilbert ␣ envelope ␣of␣V(t)”)
plt . tight_layout ()
outC = os. path . join (outdir, “A1_envelope_timeseries .png “)
plt . savefig (outC, dpi =300)
plt . show ()
*# Panel D: PSD[ envelope ] with f_slow*
plt . figure (figsize =(6, 2))
plt . semilogy (f_env, P_env, lw =0.7, label =“PSD[ envelope ]”)
if not np. isnan (f_slow):
  plt . axvline (
  f_slow,
  ls=“--”,
  color =“red “,
  label =f” f_slow ␣~␣{ f_slow :.2 f}␣Hz”
 )
plt . xlabel (“frequency ␣(Hz)”)
  plt . ylabel (“power “)
  plt . title (“Envelope ␣PSD “)
  plt . legend (fontsize =6)
  plt . tight_layout ()
  outD = os. path . join (outdir, “A1_psd_envelope .png”)
  plt . savefig (outD, dpi =300)
  plt . show ()
  return outA, outB, outC, outD
# ----------------------
*# Main execution*
# ----------------------
t, x, fs, meta = extract_timeseries_from_video (
  VIDEO_PATH,
  roi = FIXED_ROI,
  fps_override = FPS_OVERRIDE,
  denoise_ma = SMOOTHING_WINDOW,
  normalize_drift =True,
  max_frames = MAX_FRAMES
)
print (“fs_used ␣(Hz):”, meta [“fs_used “])
print (“frames ␣ analyzed :”, meta [“nframes “])
print (“ROI␣[x0,y0,x1,y1 ]:”, meta [“roi”])
print (“field ␣of␣ view ␣(height ␣x␣ width):”, meta [“height “], “x”, meta
  [“width “])
res = analyze_timeseries_level0 (
  t, x, fs,
  fast_band = FAST_BAND,
  prominence_factor = PROMINENCE_FACTOR,
  peak_distance_bins = PEAK_DISTANCE_BINS
)
print (“f_plus ␣␣(Hz):”, res [“f_plus “])
print (“f_minus ␣(Hz):”, res [“f_minus “])
print (“f_slow ␣␣(Hz):”, res [“f_slow “])
print (“enrichment_ratio ␣(envelope ␣ peak ␣/␣ median ␣ background):”,
  res [“enrichment_ratio “])
panel_paths = save_and_show_panels (res, outdir = OUTDIR)
print (“Saved ␣ panel ␣ PNGs :”, panel_paths)
~~~

### M.3 Level-1 dual-channel pipeline script (synthetic benchmark)

~~~
*#!/ usr /bin/ env python3*
*# -*- coding* :    *utf -8 -*-*
“““
Listing ␣3.␣ Synthetic ␣ preregistered ␣”Level -1”␣ benchmark ␣ figure .
This ␣ script :
1.␣ Generates ␣ paired ␣” voltage - like “␣(V)␣and␣” mechanical - like “␣(M)␣
  time ␣ series
␣␣␣ with ␣two␣ narrowband ␣~1␣Hz␣ carrier ␣ modes .
2.␣ Computes ␣ their ␣ spectra,␣ slow ␣ beat ␣ frequency ␣f_slow,␣and ␣
  electromechanical
␣␣␣ coherence ␣at␣ f_slow .
3.␣ Builds ␣the␣ neutral ␣ stroboscopic ␣map :
␣␣␣␣␣␣ Delta_theta_n ␣=␣ theta_ {n+1}␣-␣ theta_n ␣␣vs␣␣ theta_n
␣␣␣ Fits ␣the␣even - harmonic ␣map :
␣␣␣␣␣␣ Delta_theta ␣=␣ Omega ␣-␣K2␣*␣ sin (2* theta)
␣␣␣and ␣ compares ␣ Omega_hat ␣to␣the␣ predicted ␣ lock
␣␣␣␣␣␣ Omega_2over21 ␣=␣pi *2/21.
4.␣ Renders ␣ four ␣ separate ␣ panels ␣and␣ saves ␣ them ␣ under ␣./ figs ␣as:
␣␣␣␣␣␣ Level1_traces .png
␣␣␣␣␣␣ Level1_psd .png
␣␣␣␣␣␣ Level1_coherence .png
␣␣␣␣␣␣ Level1_neutral_map .png
All ␣ numbers ␣are␣ deterministic ␣ because ␣we␣fix ␣np. random . seed ().
“““
import os
import numpy as np
np. random . seed (12345)    *# make the synthetic dataset deterministic*
import matplotlib . pyplot as plt
from scipy . signal import welch, detrend, hilbert, csd
from numpy . linalg import lstsq
# ------------------------
*# 1. Synthetic dual - channel generator*
# ------------------------
def generate_synthetic_dual_channel ():
  “““
␣␣␣␣ Generate ␣ paired ␣ synthetic ␣ signals ␣V(t)␣(“voltage - like “)␣and␣M(
  t)
␣␣␣␣(“mechanical - like “)␣ with ␣two␣~1␣Hz␣ carriers,␣ plus ␣ noise ␣and␣a␣
shared
␣␣␣␣ slow ␣ beat .
␣␣␣␣ Returns ␣ time ␣ array ␣t,␣ detrended /z-scored ␣V␣and ␣M,␣ and␣ sampling
  ␣ rate ␣fs.
␣␣␣␣”““
  fs = 100.0    *# Hz sampling rate (preregistered spec* :
  >=80 -100 Hz)
T = 120.0    *# total duration [s]*
t = np. arange (0.0, T, 1.0/ fs)
*# Two carrier frequencies in the ~1 Hz band*
f_plus_true = 1.20 # Hz
f_minus_true = 1.03 # Hz
*# A relative phase lag between channels*
phase_lag = 0.40 # radians
*# “Voltage - like “raw signal*
V_clean = (
  np. sin (2* np.pi* f_plus_true * t)
  + 0.8* np. sin (2* np.pi* f_minus_true * t)
)
V_noisy = V_clean + 0.05* np. random . standard_normal (len (t))
*# “Mechanical - like “raw signal with a shallow slow modulation*
M_clean = (
  0.9* np.sin (2* np.pi* f_plus_true * t + phase_lag)
  + 0.7* np. sin (2* np.pi* f_minus_true * t + phase_lag)
)
slow_mod = 0.1* np.sin (2* np.pi *0.17* t + 1.0) # ~0.17 Hz shared
  slow envelope
M_noisy = M_clean + slow_mod + 0.05* np. random . standard_normal
  (len(t))
# Detrend + z-score each channel
Vd = detrend (V_noisy, type =“linear “)
Vd = (Vd - np. mean (Vd)) / (np. std(Vd) + 1e -12)
Md = detrend (M_noisy, type =“linear “)
Md = (Md - np. mean (Md)) / (np. std(Md) + 1e -12)
return t, Vd, Md, fs
# ------------------------
*# 2. Spectral analysis helpers*
# ------------------------
def welch_psd (x, fs, nperseg = None):
  “““
␣␣␣␣ Compute ␣ Welch ␣ power ␣ spectral ␣ density ␣(PSD).
␣␣␣␣ Returns ␣ frequency ␣ array ␣f␣and␣ PSD␣Pxx.
␣␣␣␣”““
  if nperseg is None :
  nperseg = min(len (x), 4096)
  f, Pxx = welch (x, fs=fs, nperseg = nperseg)
  return f, Pxx
  def find_two_carriers (f, Pxx, fast_band,
  prominence_factor =2.0,
  min_distance_bins =3):
  “““
␣␣␣␣ Identify ␣up␣to␣two␣ strong ␣ carrier ␣ peaks ␣in␣a␣ specified ␣’
  fast_band ‘
␣␣␣␣(e.g.␣0.5 -2␣Hz).␣We␣do␣a␣ simple ␣local - maximum ␣ scan ␣ with ␣a␣
  relative
␣␣␣␣ threshold,␣ keep ␣the␣top ␣two␣ peaks ␣by␣ amplitude,␣ and␣ define ␣the
  ␣ slow ␣ beat
␣␣␣␣ f_slow ␣=␣| f_plus ␣-␣ f_minus |.
␣␣␣␣”““
  mask = (f >= fast_band [0]) & (f <= fast_band [1])
  fband = f[ mask ]
  Pband = Pxx[ mask ]
  if len(fband) < 3:
  return np.nan, np.nan, np.nan
thr = np. median (Pband) * prominence_factor
peak_ids = []
for i in range (1, len(fband) -1):
  if (Pband [i] > Pband [i -1] and
  Pband [i] > Pband [i+1] and
  Pband [i] > thr):
  if len(peak_ids) == 0 or (i - peak_ids [ -1]) >=
  min_distance_bins :
  peak_ids . append (i)
if len(peak_ids) == 0:
  return np.nan, np.nan, np.nan
*# sort by peak height, descending*
peak_ids_sorted = sorted (peak_ids, key= lambda k: Pband [k],
  reverse = True)
f_plus = fband [ peak_ids_sorted [0]] if len(peak_ids_sorted) >=
  1 else np. nan
f_minus = fband [ peak_ids_sorted [1]] if len(peak_ids_sorted) >=
  2 else np. nan
f_slow = np.nan
if (not np. isnan (f_plus)) and (not np. isnan (f_minus)):
  f_slow = abs (f_plus - f_minus)
return f_plus, f_minus, f_slow
def coherence_spectrum (V, M, fs, nperseg = None):
  “““
␣␣␣␣ Magnitude - squared ␣ coherence ␣ Lambda (f)␣ between ␣V␣and␣M:
␣␣␣␣␣␣ Lambda (f)␣=␣| S_VM |^2␣/␣(S_V␣*␣S_M)
␣␣␣␣ Returns ␣ frequency ␣ array ␣f␣and␣ coherence ␣ array ␣coh(f).
␣␣␣␣”““
  if nperseg is None :
nperseg = min(len (V), 4096)
f, Sv = welch (V, fs=fs, nperseg = nperseg)
_, Sm = welch (M, fs=fs, nperseg = nperseg)
_, Svm = csd (V, M, fs=fs, nperseg = nperseg)
coh = (np.abs(Svm) **2) / (Sv * Sm + 1e -15)
return f, coh
def get_coherence_at_frequency (f, coh, f_target, rel_window =0.15) :
  “““
␣␣␣␣ Extract ␣ the␣ coherence ␣ magnitude ␣ near ␣a␣ target ␣ frequency ␣
  f_target,
␣␣␣␣i.e.␣ within ␣+/-␣(rel_window ␣*␣ f_target).
␣␣␣␣”““
  if np. isnan (f_target) or f_target <= 0 or f_target > np.max(f)
  :
  return np.nan
  lo = (1.0 - rel_window)* f_target
  hi = (1.0 + rel_window)* f_target
  mask = (f >= lo) & (f <= hi)
  if not np.any (mask):
  return np.nan
  return float (np.max(coh[ mask ]))
# ------------------------
*# 3. Phase, neutral strobing, even - harmonic fit*
# ------------------------
def unwrap_phase (sig):
  “““
␣␣␣␣ Hilbert - transform - based ␣ instantaneous ␣phase,␣ unwrapped .
␣␣␣␣”““
  analytic = hilbert (sig)
  return np. unwrap (np. angle (analytic))
def neutral_strobe (t, phi_V, phi_M):
  “““
␣␣␣␣” Neutral “␣ events ␣are␣ times ␣ where ␣the␣ instantaneous ␣ phase ␣
  difference
␣␣␣␣␣␣psi (t)␣=␣ phase (V)␣-␣ phase (M)
␣␣␣␣ crosses ␣0␣ modulo ␣pi␣ with ␣ positive ␣slope,␣i.e.␣V␣and␣M␣are␣
  mirror - symmetric .
␣␣␣␣We␣ detect ␣ those ␣zero - crossings ␣of␣psi(t)␣mod ␣pi,␣ then ␣ sample ␣
  the ␣ phase
␣␣␣␣of␣V␣at␣ those ␣ instants .
␣␣␣␣ Returns :
␣␣␣␣␣␣ t_strobes ␣␣␣␣␣␣-␣ times ␣of␣ neutral ␣ crossings
␣␣␣␣␣␣ theta_strobes ␣␣-␣ phase (V)␣ mod␣2* pi␣at␣ those ␣ crossings
␣␣␣␣”““
  psi = phi_V - phi_M
  psi_unwrap = np. unwrap (psi)
     *# Fold modulo pi into (-pi /2, +pi /2] to catch both +/- cross*
  psi_fold = ((psi_unwrap + 0.5* np.pi) % np.pi) - 0.5* np.pi
  t_strobes = []
  theta_strobes = []
for i in range (len (t) -1):
     *# crossing from negative to positive*
  if (psi_fold [i] < 0.0) and (psi_fold [i+1] >= 0.0) :
  frac = (- psi_fold [i]) / (psi_fold [i +1] - psi_fold [i] +
  1e -15)
  t_cross = t[i] + frac *(t[i+1] - t[i])
*# interpolate V- phase at crossing*
phiV_cross = phi_V [i] + frac *(phi_V [i+1] - phi_V [i])
theta = np.mod(phiV_cross, 2.0* np.pi)
  t_strobes . append (t_cross)
  theta_strobes . append (theta)
  return np. array (t_strobes), np. array (theta_strobes)
def fit_even_map (theta):
  “““
␣␣␣␣ Build ␣the␣ neutral ␣ stroboscopic ␣ map:
␣␣␣␣␣␣ theta_n ␣␣␣=␣ phase (V)␣at␣the␣ nth␣ neutral ␣ crossing
␣␣␣␣␣␣ Delta_theta_n ␣=␣ theta_ {n+1}␣-␣ theta_n
␣␣␣␣ Model :
␣␣␣␣␣␣ Delta_theta_n ␣=␣ Omega ␣-␣K2␣*␣ sin (2␣*␣ theta_n)
␣␣␣␣We␣fit␣[Omega,␣K2]␣by␣ linear ␣ regression ␣of
␣␣␣␣␣␣ Delta_theta_n ␣␣vs␣␣[1,␣sin (2* theta_n)].
␣␣␣␣ Returns :
␣␣␣␣␣␣ Omega_hat,␣K2_hat,
␣␣␣␣␣␣ stderr_Omega,␣ stderr_K2,
␣␣␣␣␣␣(theta_n,␣dtheta,␣ dtheta_fit)
␣␣␣␣”““
if len(theta) < 3:
  return (np.nan, np.nan,
  np.nan, np.nan,
  (np. array ([]), np. array ([]), np. array ([])))
th0 = theta [: -1]
th1 = theta [1:]
dtheta = th1 - th0
# wrap dtheta into (-pi, pi]
dtheta = ((dtheta + np.pi) % (2.0* np.pi)) - np.pi
*# Regression design matrix for:*
# dtheta = a + b * sin (2* theta)
X = np. column_stack ((np. ones_like (th0), np.sin (2.0* th0)))
y = dtheta
beta, _, _, _ = lstsq (X, y, rcond = None)
a, b = beta [0], beta [1] # a ~ Omega_hat, b ~ (- K2_hat)
*# Estimate standard errors assuming iid residuals*
if len(y) > 2:
  dof = len (y) - 2
  sigma2 = np.sum ((y - X @ beta) **2) / (dof if dof > 0 else
  1)
else :
  sigma2 = np.sum ((y - X @ beta) **2)
XtX_inv = np. linalg .inv(X.T @ X)
cov_beta = sigma2 * XtX_inv
stderr_a = np. sqrt (cov_beta [0,0])
stderr_b = np. sqrt (cov_beta [1,1])
Omega_hat = a
K2_hat = -b # minus sign by convention
stderr_Om = stderr_a
stderr_K2 = stderr_b
dtheta_fit = X @ beta # fitted Delta_theta_n
return Omega_hat, K2_hat, stderr_Om, stderr_K2, (th0, dtheta,
dtheta_fit)
# ------------------------
*# 4. Run pipeline (synthetic Level -1 benchmark)*
# ------------------------
*# Generate synthetic simultaneous V(t), M(t)*
t, Vsig, Msig, fs = generate_synthetic_dual_channel ()
# Power spectra
fV, PV = welch_psd (Vsig, fs)
fM, PM = welch_psd (Msig, fs)
*# Identify fast carrier peaks and beat frequency*
FAST_BAND = (0.5, 2.0)    *# Hz range where we expect the ~1 Hz*
carriers
f_plus, f_minus, f_slow = find_two_carriers (fV, PV, FAST_BAND)
*# Coherence spectrum and its slow - band value*
fCoh, Coh = coherence_spectrum (Vsig, Msig, fs)
Lambda_fslow = get_coherence_at_frequency (fCoh, Coh, f_slow,
  rel_window =0.15)
*# Neutral stroboscopic map*
phi_V = unwrap_phase (Vsig)
phi_M = unwrap_phase (Msig)
t_strobe, theta_strobe = neutral_strobe (t, phi_V, phi_M)
Omega_hat, K2_hat, se_Om, se_K2, mapdata = fit_even_map (
  theta_strobe)
th0, dtheta, dtheta_fit = mapdata
*# Predicted lock 2/21:*
Omega_lock_2_21 = np.pi * (2.0/21.0)
lock_dist = np.abs(Omega_hat - Omega_lock_2_21)
# ------------------------
*# 5. Panel - saving helpers (four separate PNGs)*
# ------------------------
def save_level1_traces (t, V, M, out_png):
  “““Panel ␣A:␣ time ␣ series ␣of␣V(t)␣and␣M(t).”““
  fig, ax = plt . subplots (figsize =(6, 2.3))
  ax. plot (t, V, lw =0.8, label =“V(t)␣(voltage - like)”)
  ax. plot (t, M, lw =0.8, label =“M(t)␣(mechanical - like)”)
  ax. set_title (“Simultaneous ␣ electromechanical ␣ traces ␣(synthetic
 )”)
  ax. set_xlabel (“time ␣(s)”)
  ax. set_ylabel (“z-scored ␣ signal “)
  ax. legend (fontsize =7, loc=“upper ␣ right “)
  fig . tight_layout ()
  fig . savefig (out_png, dpi =300)
  plt . close (fig)
def save_level1_psd (f, Pvv, Pmm, f_plus, f_minus, out_png):
  “““Panel ␣B:␣ power ␣ spectra ␣of␣V␣and␣M␣ with ␣ carrier ␣ peaks .”““
  fig, ax = plt . subplots (figsize =(6, 2.3))
  ax. semilogy (f, Pvv, lw =0.8, label =“PSD[V]”)
  ax. semilogy (f, Pmm, lw =0.8, label =“PSD[M]”, alpha =0.8)
  if not np. isnan (f_plus):
  ax. axvline (f_plus, ls=“--”, color =“tab: orange “,
  label =f” f_plus ␣~␣{ f_plus :.2f}␣Hz”)
if not np. isnan (f_minus):
  ax. axvline (f_minus, ls=“--”, color =“tab: green “,
  label =f” f_minus ␣~␣{ f_minus :.2 f}␣Hz”)
ax. set_title (“Power ␣ spectra ␣of␣V␣and␣M␣(synthetic ␣dual - channel
 )”)
ax. set_xlabel (“frequency ␣(Hz)”)
ax. set_ylabel (“power “)
ax. legend (fontsize =7, loc=“upper ␣ right “)
fig . tight_layout ()
fig . savefig (out_png, dpi =300)
plt . close (fig)
def save_level1_coherence (f_coh, coh, f_slow, Lambda_slow, out_png
 ):
  “““Panel ␣C:␣ electromechanical ␣ coherence ␣ Lambda (f)␣ with ␣ f_slow ␣
  marker .”““
  fig, ax = plt . subplots (figsize =(6, 2.3))
  ax. plot (f_coh, coh, lw =0.8)
  if not np. isnan (f_slow) and not np. isnan (Lambda_slow):
  ax. axvline (
  f_slow,
  ls=“--”,
  color =“red”,
  label =f” f_slow ␣~␣{ f_slow :.2 f}␣Hz,␣ Lambda ␣~␣{
  Lambda_slow :.2 f}”
)
ax. set_xlim (0, 50)
ax. set_ylim (0, 1.05)
ax. set_title (“Electromechanical ␣ coherence ␣(synthetic)”)
ax. set_xlabel (“frequency ␣(Hz)”)
ax. set_ylabel (“Lambda (f)”)
ax. legend (fontsize =7, loc=“upper ␣ right “)
fig . tight_layout ()
fig . savefig (out_png, dpi =300)
plt . close (fig)
def save_level1_neutral_map (theta_n, dtheta, dtheta_fit,
  Omega_hat, K2_hat,
  Omega_2over21, lock_dist,
  out_png):
  “““Panel ␣D:␣ neutral ␣ stroboscopic ␣ map␣and␣even - harmonic ␣ fit.”““
  fig, ax = plt . subplots (figsize =(6, 2.3))
  ax. scatter (theta_n, dtheta, s=10, alpha =0.6,
  label =“data “, color =“tab: blue “)
if len(theta_n) > 0:
order = np. argsort (theta_n)
ax. plot (theta_n [ order ], dtheta_fit [ order ],
  color =“red “, linewidth =1.3,
  label =“fit :␣ Delta_theta ␣=␣ Omega ␣-␣K2*sin (2* theta)”
 )
box_text = (
“Omega_hat ␣~=␣ %.3f␣rad,␣ K2_hat ␣~=␣ %.3f\n”
  “Omega_2over21 ␣~=␣%.3f␣rad \n”
  “| Omega_hat ␣-␣ Omega_2over21 |␣~=␣ %.3f␣rad”
  % (Omega_hat, K2_hat, Omega_2over21, lock_dist)
)
  ax. text (
  0.02, 0.98, box_text,
  transform =ax. transAxes,
  ha=“left “, va=“top “,
  fontsize =7,
  bbox = dict (boxstyle =“round “, fc=“white “, ec=“0.8”, alpha
  =0.9)
)
ax. set_title (“Neutral ␣ stroboscopic ␣map ␣(synthetic)”)
ax. set_xlabel (“theta_n ␣(phase ␣of␣V␣at␣ neutral ␣ strobe)”)
ax. set_ylabel (“theta_ {n +1}␣-␣ theta_n “)
ax. legend (fontsize =7, loc=“lower ␣ left “)
fig . tight_layout ()
fig . savefig (out_png, dpi =300)
plt . close (fig)
# ------------------------
*# 6. Save panels and print summary*
# ------------------------
outdir = “figs “
os. makedirs (outdir, exist_ok = True)
save_level1_traces (
  t, Vsig, Msig,
  os. path . join (outdir, “Level1_traces .png “)
)
save_level1_psd (
  fV, PV, fM, f_plus, f_minus,
  os. path . join (outdir, “Level1_psd .png “)
)
save_level1_coherence (
  fCoh, Coh, f_slow, Lambda_fslow,
  os. path . join (outdir, “Level1_coherence .png”)
)
save_level1_neutral_map (
th0, dtheta, dtheta_fit,
Omega_hat, K2_hat,
Omega_lock_2_21, lock_dist,
os. path . join (outdir, “Level1_neutral_map . png”)
)
*# Console readout for reproducibility*
print (“fs␣=␣ %.1f␣Hz” % fs)
print (“f_plus ␣␣=␣%.3f␣Hz” % f_plus)
print (“f_minus ␣=␣%.3f␣Hz” % f_minus)
print (“f_slow ␣␣=␣%.3f␣Hz” % f_slow)
print (“Lambda (f_slow)␣=␣%.3 f” % Lambda_fslow)
print (“Omega_hat ␣=␣%.3f␣+/-␣%.3f␣ rad” % (Omega_hat, se_Om))
print (“K2_hat ␣␣␣␣=␣%.3f␣+/-␣%.3f” % (K2_hat, se_K2))
print (“Omega_2over21 ␣=␣%.3f␣rad” % Omega_lock_2_21)
print (“| Omega_hat ␣-␣ Omega_2over21 |␣=␣%.3f␣rad” % lock_dist)
print (“Saved ␣Level -1␣ panels ␣in”, outdir)
M.4 Computational prevalidation of P3 (weak resonant drive at fslow)
*# -*- coding* :    *utf -8 -*-*
“““
Listing ␣4.␣ P3_sandbox .py␣-␣ Computational ␣ preregistration ␣ test ␣for␣
P3.
Goal
----
Exercise ␣a␣ minimal,␣ reproducible ␣ pipeline ␣ that ␣ demonstrates :
(i)␣weak,␣ frequency - specific ␣ entrainment ␣at␣the ␣ intrinsic ␣ slow ␣
  cadence ␣f_slow,
(ii)␣ simultaneous ␣ increase ␣in␣slow - band ␣ electromechanical ␣
coherence ␣(Lambda)
␣␣␣␣␣and ␣ decrease ␣in␣ neutral -map␣ angular ␣ variance ␣(Var_map)␣ only ␣
  on - band .
This ␣ script ␣is␣ deliberately ␣ argument - free ␣(no␣ argparse)␣so␣it␣ runs
  ␣ cleanly
in␣ Colab / Notebook ␣ contexts ␣ where ␣a␣-f␣ kernel ␣ flag ␣is␣ injected .
Outputs ␣ saved ␣ under ␣./ outputs_P3_sandbox :
␣␣-␣ summary . txt␣␣␣␣␣␣␣␣␣␣␣(numeric ␣ readout)
␣␣-␣ sweep .png␣␣␣␣␣␣␣␣␣␣␣␣␣(resonance ␣ curves ␣for␣ Var_map ␣ and␣ Lambda
 )
␣␣-␣ sweep .csv␣␣␣␣␣␣␣␣␣␣␣␣␣(tabulated ␣ results)
“““
import os, json
import numpy as np
import matplotlib . pyplot as plt
from scipy . signal import hilbert, welch, csd
# -------------------------
*# 1) Unlocking - like baseline*
# -------------------------
def generate_unlocking_state (
  fs =200, dur =40,
  f_plus_V =1.8, f_minus_V =1.0,
  f_plus_S =2.5, f_minus_S =0.7,
  noise_V =1.0, noise_S =1.5,
  rw_sigma_S =0.5,
  coupling_c =0.05,
):
  n = int(dur * fs)
  t = np. arange (n) / fs
  phi_plus_V = 2 * np.pi * f_plus_V * t
  phi_minus_V = 2 * np.pi * f_minus_V * t
  V = np.sin(phi_plus_V) + 0.7 * np.sin (phi_minus_V)
  V += noise_V * np. random . randn (n)
  drift = np. cumsum (rw_sigma_S * np. random . randn (n)) / fs
  S_intrinsic = (
  0.5 * np.sin (2 * np.pi * f_plus_S * t + 0.5 + drift)
  + 0.4 * np.sin (2 * np.pi * f_minus_S * t + 1.0 + 0.3 * drift
 )
)
S = S_intrinsic + noise_S * np. random . randn (n)
S = S + coupling_c * V
V = (V - np. mean (V)) / (np.std(V) + 1e -12)
S = (S - np. mean (S)) / (np.std(S) + 1e -12)
f_slow = abs(f_plus_V - f_minus_V)
return t, V, S, f_slow, fs
# -----------------------------------
*# 2) Metrics at the slow - band f_slow*
# -----------------------------------
def slow_band_coherence (V, S, fs, f_target):
  nper = min (4096, len(V))
  f, Pxx = welch (V, fs=fs, nperseg = nper)
  _, Pyy = welch (S, fs=fs, nperseg = nper)
  _, Cxy = csd (V, S, fs=fs, nperseg = nper)
  coh = (np.abs(Cxy) ** 2) / ((Pxx * Pyy) + 1e -12)
  idx = np. argmin (np.abs (f - f_target))
  return float (coh [idx ])
def var_map_from_signals (V, S, fs):
  phi_V = np. unwrap (np. angle (hilbert (V)))
  phi_S = np. unwrap (np. angle (hilbert (S)))
  psi = phi_V - phi_S
  dpsi_dt = np. gradient (np. unwrap (psi)) * fs
*# neutral crossings* :    *psi near k*pi (sin psi ~ 0), with*
     *positive slope*
neutral_mask = (np. abs(np.sin(psi)) < 0.2) & (dpsi_dt > 0)
idxs = np. where (neutral_mask)[0]
if len(idxs) < 5:
  return float (“nan “)
*# enforce >=0*.*2 s spacing between samples*
keep = [ idxs [0]]
for j in idxs [1:]:
  if j - keep [ -1] > fs * 0.2:
  keep . append (j)
keep = np. array (keep, dtype =int)
if len(keep) < 5:
  return float (“nan “)
theta = phi_V [ keep ]
dtheta = np. diff (theta)
return float (np.var (dtheta))
def measure_block (V, S, fs, f_slow):
  return slow_band_coherence (V, S, fs, f_slow),
  var_map_from_signals (V, S, fs)
# -----------------------------------------
*# 3) Weak drive + narrow resonance “gate “*
# -----------------------------------------
def apply_drive (t, V, S, f_drive, f_slow, amp_drive =0.03,
  lock_width_frac =0.01) :
  Vd = V + amp_drive * np.sin (2 * np.pi * f_drive * t)
  Sd = S. copy ()
  df = abs(f_drive - f_slow)
  tol = lock_width_frac * f_slow
  gate = np.exp (-(df ** 2) / (2 * tol ** 2))
  Sd = (1 - gate * 0.5) * Sd + (gate * 0.5) * Vd
  Sd = (Sd - np. mean (Sd)) / (np. std(Sd) + 1e -12)
  Vd = (Vd - np. mean (Vd)) / (np. std(Vd) + 1e -12)
  return Vd, Sd, float (gate)
# -------------------------
*# 4) P3 on/off - band protocol*
# -------------------------
def run_protocol ():
  t, V0, S0, f_slow, fs = generate_unlocking_state ()
  Lam0, Var0 = measure_block (V0, S0, fs, f_slow)
  V_on, S_on, gate_on = apply_drive (t, V0, S0, f_slow, f_slow)
  Lam_on, Var_on = measure_block (V_on, S_on, fs, f_slow)
  V_p, S_p, gate_p = apply_drive (t, V0, S0, f_slow *1.10, f_slow)
  Lam_p, Var_p = measure_block (V_p, S_p, fs, f_slow)
  V_m, S_m, gate_m = apply_drive (t, V0, S0, f_slow *0.90, f_slow)
  Lam_m, Var_m = measure_block (V_m, S_m, fs, f_slow)
  return dict (
     *# enforce >=0*.*2 s spacing between samples*
  keep = [ idxs [0]]
  for j in idxs [1:]:
  if j - keep [ -1] > fs * 0.2:
  keep . append (j)
  keep = np. array (keep, dtype =int)
  if len(keep) < 5:
  return float (“nan “)
  theta = phi_V [ keep ]
  dtheta = np. diff (theta)
  return float (np.var (dtheta))
def measure_block (V, S, fs, f_slow):
  return slow_band_coherence (V, S, fs, f_slow),
  var_map_from_signals (V, S, fs)
# -----------------------------------------
#    *3) Weak drive + narrow resonance “gate “*
# -----------------------------------------
def apply_drive (t, V, S, f_drive, f_slow, amp_drive =0.03,
  lock_width_frac =0.01) :
  Vd = V + amp_drive * np.sin (2 * np.pi * f_drive * t)
  Sd = S. copy ()
  df = abs(f_drive - f_slow)
  tol = lock_width_frac * f_slow
  gate = np.exp (-(df ** 2) / (2 * tol ** 2))
  Sd = (1 - gate * 0.5) * Sd + (gate * 0.5) * Vd
  Sd = (Sd - np. mean (Sd)) / (np. std(Sd) + 1e -12)
  Vd = (Vd - np. mean (Vd)) / (np. std(Vd) + 1e -12)
  return Vd, Sd, float (gate)
# -------------------------
#    *4) P3 on/off - band protocol*
# -------------------------
def run_protocol ():
  t, V0, S0, f_slow, fs = generate_unlocking_state ()
  Lam0, Var0 = measure_block (V0, S0, fs, f_slow)
  V_on, S_on, gate_on = apply_drive (t, V0, S0, f_slow, f_slow)
  Lam_on, Var_on = measure_block (V_on, S_on, fs, f_slow)
  V_p, S_p, gate_p = apply_drive (t, V0, S0, f_slow *1.10, f_slow)
  Lam_p, Var_p = measure_block (V_p, S_p, fs, f_slow)
  V_m, S_m, gate_m = apply_drive (t, V0, S0, f_slow *0.90, f_slow)
  Lam_m, Var_m = measure_block (V_m, S_m, fs, f_slow)
  return dict (
  f_slow = float (f_slow), fs= float (fs),
  baseline = dict (Lambda = float (Lam0), Var_map = float (Var0)),
  on= dict (Lambda = float (Lam_on), Var_map = float (Var_on), gate =
  float (gate_on)),
  off_plus = dict (Lambda = float (Lam_p), Var_map = float (Var_p),
  gate = float (gate_p)),
  off_minus = dict (Lambda = float (Lam_m), Var_map = float (Var_m),
  gate = float (gate_m)),
 )
# -----------------------------------------------
#    *5) Frequency sweep for resonance diagnostics*
# -----------------------------------------------
def frequency_sweep (t, V0, S0, f_slow, fs, fmin_factor =0.7,
  fmax_factor =1.3, n_pts =25) :
  freqs = np. linspace (f_slow * fmin_factor, f_slow * fmax_factor
, n_pts)
  Lam_list, Var_list, gate_list = [], [], []
  for fd in freqs :
  Vd, Sd, gate = apply_drive (t, V0, S0, fd, f_slow)
  Lam, Var = measure_block (Vd, Sd, fs, f_slow)
  Lam_list . append (Lam); Var_list . append (Var); gate_list .
  append (gate)
  return freqs, np. array (Lam_list), np. array (Var_list), np. array
  (gate_list)
def plot_and_save (freqs, Lam_arr, Var_arr, gate_arr, f_slow,
  out_png):
  fig, ax = plt . subplots (2, 1, figsize =(6, 6), sharex = True)
  ax [0]. plot (freqs, Var_arr, “o-”, lw =1)
  ax [0]. axvline (f_slow, ls=“--”, color =“k”, label =“f_slow “)
  ax [0]. set_ylabel (“Var_map ␣(lower ␣=␣ more ␣ stable)”)
  ax [0]. legend ()
  ax [0]. set_title (“Frequency ␣ sweep :␣ angular ␣ stability “)
  ax [1]. plot (freqs, Lam_arr, “o-”, lw =1)
  ax [1]. axvline (f_slow, ls=“--”, color =“k”, label =“f_slow “)
  ax [1]. set_xlabel (“drive ␣ frequency ␣(Hz)”)
  ax [1]. set_ylabel (“Lambda ␣(higher ␣=␣ more ␣ coherence)”)
  ax [1]. legend ()
  ax [1]. set_title (“Frequency ␣ sweep :␣slow - band ␣ coherence “)
  ax_gate = ax [1]. twinx ()
  ax_gate . plot (freqs, gate_arr, “r--”, alpha =0.35)
  ax_gate . set_ylabel (“gate ␣(0 -1)”)
  fig . tight_layout ()
  fig . savefig (out_png, dpi =200)
  plt . close (fig)
# -------------------
*# 6) Script entrypoint*
# -------------------
if __name__ == “__main__ “:
  np. random . seed (0)
  outdir = “outputs_P3_sandbox “
  os. makedirs (outdir, exist_ok = True)
     *# Main protocol*
  t, V0, S0, f_slow, fs = generate_unlocking_state ()
  res = run_protocol ()
     *# Save summary*
  with open (os. path . join (outdir, “summary . txt”), “w”) as f:
  f. write (json . dumps (res, indent =2))
     *# Sweep & figure*
  freqs, Lam_arr, Var_arr, gate_arr = frequency_sweep (t, V0, S0,
  f_slow, fs)
  np. savetxt (os. path . join (outdir, “sweep .csv”),
  np.c_[freqs, Lam_arr, Var_arr, gate_arr ],
  delimiter =“,”, header =“freq,Lambda, Var_map, gate “,
  comments =““)
  plot_and_save (freqs, Lam_arr, Var_arr, gate_arr, f_slow,
  os. path . join (outdir, “sweep .png”))
     *# Console readout*
  print (“f_slow ␣~␣%.3f␣Hz” % f_slow)
  for k in [“baseline “,”on”,” off_plus “,” off_minus “]:
  print (k, res[k])
~~~

### M.5 Popper falsifiability: video/CSV pipeline for neutral stroboscopy, even/odd harmonics, and bicoherence (Figs. H1–H4)

~~~
*#!/ usr /bin/ env python3*
# -*- coding :    utf -8 -*-
“““
Listing ␣5.␣    Popper_falsifiability :␣    video /CSV ␣    pipeline ␣for␣H1 -H4␣
     diagnostics
(neutral ␣    stroboscopy,␣    even /odd ␣    harmonics,␣and␣    bicoherence).
Usage ␣(CLI ␣    examples):
␣␣    python ␣    s4_popper_falsifiability .py␣    --video ␣    input .mov␣    --fs␣25␣    --
     roi ␣    100:80:320:240
␣␣    python ␣    s4_popper_falsifiability .py␣    --csv␣    input .csv␣    --fs␣25
␣␣#␣    Optional :
␣␣#␣␣␣    -- downsample ␣1␣    -- outdir ␣    figs ␣    --fmin ␣    0.05 ␣    --fmax ␣    0.40 ␣    --
     surrogates ␣200␣    --bico - nseg ␣32
␣␣#␣␣␣    --colab - picker ␣␣␣#␣    forces ␣a␣    picker ␣in␣    Colab ␣(now␣it␣    also ␣
     opens ␣    automatically)
“““
import os, json, argparse, warnings
import numpy as np
import matplotlib .    pyplot as plt
from scipy .    signal import butter, filtfilt, hilbert, welch
SEED = 1234
np. random .    seed (SEED)
# -----------------------
# *Small utils*
# -----------------------
def ensure_dir (path):
     os. makedirs (path, exist_ok = True)
def parse_roi (s):
     try :
     x0, y0, x1, y1 = [int(v) for v in s. split (“:”)]
     return (x0, y0, x1, y1)
     except Exception :
     raise ValueError (“ROI␣    must ␣be␣’x0:y0:x1:y1 ‘”)
def in_colab ():
     try :
     import google .    colab # type :    ignore
     return True
     except Exception :
     return False
# -----------------------
# *DSP helpers*
# -----------------------
def bandpass (x, fs, fmin, fmax, order =3):
     b, a = butter (order, [ fmin /(fs /2), fmax /(fs /2)], btype =‘band ‘)
     return filtfilt (b, a, x)
def norm_std (x):
     x = np. asarray (x, float)
     return (x - np. mean (x)) / (np.std(x) + 1e -12)
def psd_peak_freq (x, fs, fmin =0.5, fmax =4.0) :
     nper = min (4096, len(x))
     f, Pxx = welch (x, fs=fs, nperseg = nper)
     band = (f >= fmin) & (f <= fmax)
     if not np.any (band):
     return float (“nan “), f, Pxx
     k = np. argmax (Pxx[ band ])
     f0 = float (f[ band ][k])
     return f0, f, Pxx
def integrate_band (f, P, f_center, bw):
     m = (f >= max (0.0, f_center - bw)) & (f <= f_center + bw)
     if not np.any (m):
     return 0.0
     return float (np. trapz (P[m], f[m]))
# -----------------------
# *IO: CSV / Video / Colab picker*
# -----------------------
def load_csv (path):
     import csv
     vals = []
     with open (path, “r”, newline =““) as fh:
       reader = csv .    reader (fh)
       for row in reader :
         if not row:
            continue
         try :
            vals .    append (float (row [0]))
         except Exception :
            continue
     return np. asarray (vals, float)
def load_video_series (path, roi=None, downsample =1):
     fs = None
     frames = []
     try :
         import cv2
         cap = cv2. VideoCapture (path)
         if not cap. isOpened ():
            raise RuntimeError (“OpenCV ␣    could ␣    not␣    open ␣    video .”)
     fs = cap.get(cv2 .    CAP_PROP_FPS)
     while True :
         ok, frame = cap. read ()
         if not ok:
           break
     if roi is not None :
         x0,y0,x1,y1 = roi
         frame = frame [y0:y1, x0:x1]
     gray = frame if frame .    ndim == 2 else cv2. cvtColor (
        frame, cv2. COLOR_BGR2GRAY)
     frames .    append (gray .    mean ())
   cap .    release ()
 except Exception :
   try :
     import imageio .v2 as imageio
     reader = imageio .    get_reader (path)
     meta = reader .    get_meta_data ()
     fs = meta .get (“fps”, fs)
     for frame in reader :
       if roi is not None :
         x0,y0,x1,y1 = roi
         frame = frame [y0:y1, x0:x1]
     gray = frame if frame .    ndim == 2 else np. mean (frame
,    axis =2)
     frames .    append (gray .    mean ())
     reader .    close ()
     except Exception as e:
        raise RuntimeError (“Could ␣not␣    read ␣    video .␣    Need ␣    OpenCV ␣
     or␣    imageio .␣    Error :␣{}”. format (e))
     frames = np. asarray (frames, float)
     if downsample > 1:
     frames = frames [:: downsample ]
        if fs is not None :
          fs = fs / downsample
     return frames, fs
def try_colab_upload (ask= True):
     if not ask:
        return None, None
     try :
        from google .    colab import files # type :    ignore
        print (“[ info ]␣    Opening ␣a␣    file ␣    picker ␣(Colab).␣    Upload ␣a␣
        video ␣(. mov /. mp4 /…) ␣or␣a␣CSV .”)
     up = files .    upload ()
        if not up:
          print (“[ warn ]␣    Selection ␣    cancelled .”)
     return None, None
          name = list (up. keys ())[0]
     low = name .    lower ()
     if low. endswith ((“.mov”,”.mp4”,”.m4v “,”.avi”,”.mkv”,”.mpg “
,”. mpeg “)):
     return (“video “, name)
     if low. endswith (“.csv”):
     return (“csv”, name)
     print (“[ warn ]␣    Unsupported ␣    file ␣    type ;␣    using ␣    demo ␣    instead .”)
     return None, None
     except Exception :
     return None, None
# -----------------------
# *H1 -H4 feature helpers*
# -----------------------
def neutral_pairs (V_slow, fs, zero_tol =0.35, slope_tol =0.0,
     min_sep_s =2.0) :
     z = hilbert (V_slow)
     quad = np. imag (z)
     dquad = np. gradient (quad) * fs
     thr = zero_tol * (np. std(quad) + 1e -12)
     cand = (np.abs (quad) < thr) & (dquad > slope_tol)
     idx = np. flatnonzero (cand)
     if idx. size == 0:
     return [], []
     keep = []
     last = -10**9
     min_sep = int(min_sep_s * fs)
     for i in idx:
     if i - last >= min_sep :
     keep .    append (i); last = i
     keep = np. asarray (keep, int)
     pairs = [(int(keep [i]), int(keep [i +1])) for i in range (len(
     keep) -1)]
     amp = np.abs(z)
     A = [(float (amp[i]), float (amp[j])) for (i,j) in pairs ]
     return pairs, A
def bicoherence_proxy (x, fs, f0, nseg =16) :
     N = len(x)
     m = N // nseg
     if m < 64:
     return float (“nan “)
     x = x[:m* nseg ]
     def fbin (fr):
     return int(round (fr * m / fs))
     k0 = fbin (f0)
     k2 = fbin (2* f0)
     num = 0+0j; den1 = 0.0; den2 = 0.0
     win = np. hanning (m)
     for s in range (nseg):
       seg = x[s*m:(s+1)*m]
       Z = np. fft. rfft (seg * win)
       if k2 < len(Z) and k0 < len (Z):
         num += Z[k0] * Z[k0] * np. conj (Z[k2 ])
         den1 += np.abs (Z[k0 ]**2)
         den2 += np.abs (Z[k2 ])
         if den1 <= 0 or den2 <= 0:
     return float (“nan “)
     return float (np.abs(num) / np. sqrt (den1 * den2 + 1e -18))
def bico_surrogates (x, fs, f0, nseg =16, nsurr =200) :
     # Build surrogates preserving global magnitude spectrum of a
     block length L=m* nseg
     N = len(x)
     m = N // nseg
     L = m * nseg
     if m < 64 or L < 128:
     return np. array ([])
     x_use = x[:L]
     Z = np.fft. rfft (x_use)
     mag = np.abs(Z)
     rng = np. random .    default_rng (SEED)
     vals = []
     for _ in range (nsurr):
     phase = rng. uniform (0, 2* np.pi, size =Z. shape)
     phase [0] = 0.0
     if (L % 2) == 0:
     phase [ -1] = 0.0
     Zs = mag * np.exp (1j * phase)
     xs = np.fft. irfft (Zs, n=L). real
     vals .    append (bicoherence_proxy (xs, fs, f0, nseg = nseg))
     return np. asarray (vals, float)
# -----------------------
# *Main*
# -----------------------
def main ():
     ap = argparse .    ArgumentParser ()
     ap. add_argument (“--video “, type =str, default = None)
     ap. add_argument (“--csv”, type =str, default = None)
     ap. add_argument (“--fs”, type =float, default =None, help =“
     sampling ␣    rate ␣(needed ␣for␣CSV␣or␣    videos ␣    without ␣FPS)”)
     ap. add_argument (“--roi”, type =str, default =None, help =“x0:y0
     :x1:y1␣for ␣    video “)
     ap. add_argument (“-- downsample “, type =int, default =1)
     ap. add_argument (“-- outdir “, type =str, default =“figs “)
     ap. add_argument (“--fmin “, type =float, default =0.05)
     ap. add_argument (“--fmax “, type =float, default =0.40)
     ap. add_argument (“-- surrogates “, type =int, default =200)
     ap. add_argument (“--bico - nseg “, type =int, default =32)
     ap. add_argument (“--colab - picker “, action =“store_true “, help =“
     force ␣a␣    picker ␣in␣    Colab “)
     args, _ = ap. parse_known_args ()
     ensure_dir (args .    outdir)
     # Decide input source
     mode = “demo “
     source = None
     fs_used = None
     if args .    video :
     source = args .    video
     roi = parse_roi (args .roi) if args .roi else None
     V_raw, fs_video = load_video_series (args .video, roi =roi,
     downsample = args .    downsample)
     fs_used = fs_video if fs_video is not None else args .fs
     if fs_used is None :
     raise SystemExit (“For␣    video ␣    without ␣fps␣    metadata,␣
     please ␣    provide ␣    --fs.”)
     mode = “video “
     elif args .csv:
source = args .csv
     V_raw = load_csv (args .csv)
     fs_used = args .fs
     if fs_used is None :
     raise SystemExit (“CSV␣    requires ␣    --fs␣to␣be␣    provided .”)
     mode = “csv”
else :
     # Auto - open picker if running in Colab or if flag was set
     use_picker = args .    colab_picker or in_colab ()
     picked = try_colab_upload (use_picker)
     if picked and picked [0] == “video “:
        args .    video = picked [1]
        source = args .    video
        roi = parse_roi (args .roi) if args .roi else None
        V_raw, fs_video = load_video_series (args .video, roi =
         roi, downsample = args .    downsample)
        fs_used = fs_video if fs_video is not None else args .
         fs
        if fs_used is None :
     raise SystemExit (“For␣    video ␣    without ␣fps␣    metadata,␣
     please ␣    provide ␣    --fs.”)
     mode = “video “
     elif picked and picked [0] == “csv”:
     args .csv = picked [1]
     source = args .csv
     V_raw = load_csv (args .csv)
     fs_used = args .fs
     if fs_used is None :
       raise SystemExit (“CSV␣    requires ␣    --fs␣to␣be␣    provided
  .”)
     mode = “csv”
   else :
     print (“[ warn ]␣No␣    video /CSV␣    provided ;␣    falling ␣    back ␣to␣
     synthetic ␣    demo .”)
     fs_used = 25.0
     T = 100.0
     t = np. arange (int(T* fs_used))/ fs_used
     V_raw = (np.sin (2* np.pi *1.4* t + 0.6)
        + 0.35* np. sin (2* np.pi *2.6* t + 1.7)
        + 0.15* np. sin (2* np.pi *0.18* t + 0.3)
        + 0.20* np. random .    randn (len (t)))
     mode = “demo “
     source = “synthetic “
# Normalize
V_raw = norm_std (np. asarray (V_raw, float))
# Slow band (H1 + neutral stroboscopy)
V_slow = bandpass (V_raw, fs_used, args .fmin, args .fmax, order
     =3)
t = np. arange (len(V_slow))/ fs_used
z = hilbert (V_slow)
quad = np. imag (z)
# H1
plt .    figure (figsize =(12,4))
plt .    plot (t, V_slow, label =“V␣    bandpassed “)
plt .    plot (t, quad, label =“quadrature ␣    proxy “)
plt .    xlabel (“Time ␣(s)”)
plt .    ylabel (“a.u.”)
plt .    title (“Slow - band ␣    (0.05 -0.40 ␣Hz)”)
plt .    legend ()
h1_path = os. path .    join (args .outdir, “input_slowband .png “)
plt .    tight_layout (); plt .    savefig (h1_path, dpi =160) ;    plt. close ()
# H2
pairs, A = neutral_pairs (V_slow, fs_used, zero_tol =0.35,
     slope_tol =0.0, min_sep_s =2.0)
A = np. asarray (A, float)
plt .    figure (figsize =(6,6))
if len(A) > 0:
     plt .    scatter (A[:,0], A[:,1], s =30)
plt .    xlabel (“|Z|_n”); plt .    ylabel (“|Z|_{n+1}    “)
plt .    title (“Neutral - moment ␣    stroboscopy ␣(proxy)”)
h2_path = os. path .    join (args .outdir, “input_poincare_neutral .
     png “)
     plt .    tight_layout (); plt .    savefig (h2_path, dpi =160) ;    plt. close ()
# H3
f0, f_psd, P_psd = psd_peak_freq (V_raw, fs_used, fmin =0.5,
     fmax =4.0)
nper = min (4096, len(V_raw))
f_psd, Pxx = welch (V_raw, fs= fs_used, nperseg = nper)
bw = 0.10 * max(f0, 1e -3)
def band_power (fr): return integrate_band (f_psd, Pxx, fr, bw)
P2, P3 = band_power (2.0* f0), band_power (3.0* f0)
ratio = (P2 / (P3 + 1e -18))
plt .    figure (figsize =(9,6))
plt .bar ([“P(2␣f0)”, “P(3␣f0)”], [P2, P3 ])
plt .    ylabel (“Power “); plt .    title (“Even ␣vs␣odd ␣    harmonics ␣    around ␣
     f0”)
h3_path = os. path .    join (args .outdir, “input_harmonics_2f0_3f0 .
     png “)
plt .    tight_layout (); plt .    savefig (h3_path, dpi =160) ;    plt. close ()
# H4
b2 = bicoherence_proxy (V_raw, fs_used, f0, nseg = args .    bico_nseg
    )
sur = bico_surrogates (V_raw, fs_used, f0, nseg = args .    bico_nseg,
     nsurr = args .    surrogates)
pval = float ((np. sum(sur >= b2) + 1) / (len (sur) + 1)) if (sur
  .    size > 0 and np. isfinite (b2)) else float (“nan”)
plt .    figure (figsize =(10,6))
if sur. size > 0:
     plt .    hist (sur, bins =24, density =True, alpha =0.8, label =“
     surrogates “)
if np. isfinite (b2):
     plt .    axvline (b2, ls=“--”, label =“data “)
plt .    xlabel (“bicoherence ␣    proxy “); plt. ylabel (“pdf”)
plt .    title (“Bicoherence ␣(f0,f0␣    ->␣2␣f0)␣    with ␣    surrogates “)
plt .    legend ()
h4_path = os. path .    join (args .outdir, “
     input_bicoherence_surrogates .png “)
plt .    tight_layout (); plt .    savefig (h4_path, dpi =160) ;    plt. close ()
# Summary JSON
summary = {
     “seed “: SEED,
     “mode “: mode,
     “source “: source,
     “fs”: float (fs_used),
     “slow_band “: [ args .fmin, args .    fmax ],
     “n_samples “: int(len(V_raw)),
     “n_neutral_pairs “: int(len (A)),
     “neutral_params “: {“zero_tol_std “: 0.35, “slope_tol “: 0.0,
     “min_sep_s “: 2.0},
     “f0_peak_hz “: float (f0),
     “P2_over_P3 “: float (ratio),
     “bicoherence_proxy “: float (b2) if np. isfinite (b2) else
     None,
     “p_value “: float (pval) if np. isfinite (pval) else None,
     “figures “: {“H1”: h1_path, “H2”: h2_path, “H3”: h3_path, “
     H4”: h4_path }
}
with open (os. path .    join (args .outdir, “summary .    json “), “w”) as
     fjs :
     json .    dump (summary, fjs, indent =2)
print (
     “[ done ]␣    Wrote ␣    figures ␣to␣    ‘{}’.”. format (args .    outdir)
     + “␣    Neutral ␣    pairs :␣{}␣|␣f0 ={:.3 f}␣Hz␣|␣P(2 f0)/P(3 f0) ={:.3 f
  }␣|␣”. format (len(A), f0, ratio)
     + “bico_proxy ={:.3 f}␣(p ={:.2 f})”. format (
     b2 if np. isfinite (b2) else float (“nan”),
     pval if np. isfinite (pval) else float (“nan”))
)
if __name__ == “__main__ “:
     with warnings .    catch_warnings ():
     warnings .    simplefilter (“ignore “)
     main ()
~~~

### M.6 Capture kinetics: Adler-flow simulation for slow-phase capture, coherence build-up, and scaling-law prevalidation

~~~
The Python script below reproduces Fig. F.1. It simulates an Adler-like flow for the slow phase, computes a coherence proxy Λ(*t*) with sliding-window smoothing, enforces a preregistered entry criterion (signal    *<* 15% of the plateau for ≥ 5 s), fits a single-pole approach to estimate the time constant    *τ*, and reports    *T*_lock_ = −ln(1 − 0.8)    *τ* .
The grid is expanded across (*f*_slow_,    *κ, δ*) with    *n* replicates per cell; medians (and IQRs) are used per cell, and deterministic per-cell seeds based on (*f, κ, δ*, rep) ensure exact reproducibility.
In our    *in silico* validation, the lock time scaled with the predicted abscissa as 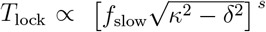. Across the expanded grid, ordinary least squares on the log–log data yielded    *s* = 1.17 with    *R*^2^ = 0.59, while a weighted fit that accounts for heteroscedasticity using weights ∝1/IQR^2^ gave    *s* = 0.71 with    *R*^2^ = 0.56. Nonparametric bootstrap 95% confidence intervals were [1.05, 1.32] for OLS and [0.25, 1.06] for the weighted fit.
*# Listing 6. Minimal in - silico prevalidation for capture kinetics*
*# Expanded grid + replicates + weighted regression + bootstrap CIs*
*# Saves PNG/CSV/TXT to a folder and also displays figures* .
import os, csv, math, numpy as np
import matplotlib as mpl; mpl . rcdefaults ()
import matplotlib . pyplot as plt
from numpy . linalg import lstsq
import hashlib
# -------    *deterministic seeding (reproducible per grid cell)* ----
MASTER_SEED = 12345
def deterministic_seed (f, k, r, rep, master = MASTER_SEED):
  msg = f”{f:.9f}|{k:.9 f}|{r :.9f}|{ rep }|{ master }”. encode ()
  h = hashlib . blake2b (msg, digest_size =8). digest ()
  return int(np. frombuffer (h, dtype =np. uint64)[0] & np. uint64 (0
  xFFFFFFFF))
*# ------- SDE (Adler - like phase model) --------*
def simulate_adler (fslow, kappa, delta, D=2e -4, dt =1/200, T=360,
  seed = None):
“““
␣␣␣␣ Simulate ␣an␣Adler - like ␣SDE ␣for␣a␣ phase ␣ variable :
␣␣␣␣␣␣␣␣ dphi ␣=␣2* pi* fslow *(delta ␣-␣ kappa *sin(phi))*dt␣+␣ sqrt (2*D)*
  dW
␣␣␣␣ Returns ␣ the␣ phase ␣ trajectory ␣ and␣the␣ time ␣ step ␣ used .
␣␣␣␣”““
  rng = np. random . default_rng (seed)
  n = int(T/dt)
  phi = 2* np.pi*rng . random ()
  phi_traj = np. empty (n, dtype =np. float64)
  drift = 2* np.pi* fslow
  sdt = math . sqrt (dt); rt2D = math . sqrt (2*D)
  for i in range (n):
  dphi = drift *(delta - kappa *np.sin(phi))*dt + rt2D * sdt*rng
  . normal ()
  phi = (phi + dphi + np.pi) %(2* np.pi) - np.pi
  phi_traj [i] = phi
  return phi_traj, dt
*# ---------- coherence proxy Lambda (t) ----------*
def lambda_trace (phi, fs, W =10.0) :
“““
␣␣␣␣ Proxy ␣for␣ coherence ␣ rise :␣ smooth ␣cos(phi)␣ with ␣a␣W- second ␣
  uniform
␣␣␣␣ window ␣to␣ emulate ␣a␣single - pole ␣ approach ␣to␣a␣ plateau .
␣␣␣␣”““
  import scipy . ndimage as nd
  x = np.cos(phi)
  L = max (1, int(W*fs))
  return nd. uniform_filter1d (x, size =L, mode =“nearest “)
*# ---------- T_lock via exponential fit ----------*
def tlock_from_exp (trace, fs, frac =0.8, entry_frac =0.15,
  entry_min_sec =5.0, fit_window_sec =20.0) :
“““
␣␣␣␣ Enforce ␣ entry ␣ criterion ␣(trace ␣ must ␣ remain ␣ below ␣ entry_frac *
  plateau ␣ for␣ >=␣ entry_min_sec),
␣␣␣␣ then ␣fit␣a␣single - pole ␣ approach ␣y(t)␣=␣P␣-␣A*exp(-t/tau)␣ using
  :
␣␣␣␣␣␣␣␣ln(P␣-␣y)␣=␣ln(A)␣-␣t/tau
␣␣␣␣ Return ␣ T_lock ␣=␣-ln (1- frac)*tau,␣or␣np.nan␣if␣ conditions ␣are␣
not ␣met.
␣␣␣␣”““
  P = np. nanmedian (trace [-int (20* fs):]) # robust plateau proxy
  if not np. isfinite (P) or P == 0:
  return np.nan
  target_entry = entry_frac * P
  below = trace < target_entry
  run, t0_idx = 0, None
  for i, ok in enumerate (below):
  run = run + 1 if ok else 0
  if run >= int (entry_min_sec *fs):
  t0_idx = i
  break
  if t0_idx is None :
  return np.nan
  Tw = max(int(fit_window_sec *fs), 1)
  t = np. arange (t0_idx, min (len(trace), t0_idx +Tw)) / fs
  y = trace [ t0_idx : min(len (trace), t0_idx +Tw)]
  Z = P - y
  mask = np. isfinite (Z) & (Z > 0)
  if mask .sum () < 30:
  return np.nan
  X = np. vstack ([ np. ones (mask .sum ()), t[ mask ]]) .T
  b, *_ = lstsq (X, np.log(Z[ mask ]), rcond = None) # ln(Z) = b0 +
  b1 t
  b1 = b[1]
  tau = -1.0/ b1 if b1 < 0 else np. nan
  if not np. isfinite (tau) or tau <= 0:
  return np.nan
  return float (-np.log (1.0 - frac) * tau)
*# ---------- log -log fits ----------*
def loglog_fit_ols (xs, ys):
  “““Ordinary ␣least - squares ␣slope,␣ intercept,␣R^2␣for ␣y␣~␣x^s␣(
  log - log).”““
  lx, ly = np.log(xs), np. log(ys)
  A = np. vstack ([ np. ones_like (lx), lx ]).T
  beta, *_ = lstsq (A, ly, rcond = None)
  intercept, slope = float (beta [0]), float (beta [1])
  yhat = A @ beta
  ss_res = float (np. sum ((ly - yhat) **2))
  ss_tot = float (np. sum ((ly - ly. mean ()) **2))
  r2 = 1.0 - ss_res / ss_tot if ss_tot > 0 else np.nan
  return slope, intercept, r2
def loglog_fit_wls (xs, ys, w):
  “““Weighted ␣least - squares ␣on␣log - log␣ with ␣ nonnegative ␣ weights ␣
  w.”““
  lx, ly = np.log(xs), np. log(ys)
  w = np. asarray (w, float)
  W = np. diag (w)
  A = np. vstack ([ np. ones_like (lx), lx ]).T
  Aw = W @ A
  yw = W @ ly
  beta, *_ = lstsq (Aw, yw, rcond = None)
  intercept, slope = float (beta [0]), float (beta [1])
  yhat = A @ beta
  ly_bar = np.sum (w*ly)/np.sum(w)
  ss_res = float (np. sum(w*(ly - yhat) **2))
  ss_tot = float (np. sum(w*(ly - ly_bar) **2))
  r2 = 1.0 - ss_res / ss_tot if ss_tot > 0 else np.nan
  return slope, intercept, r2
def bootstrap_ci (xs, ys, fit_fn, w=None, n_boot =1000, seed =2025) :
  “““Nonparametric ␣ bootstrap ␣95% ␣CI␣ for␣ slope ␣ with ␣ fit_fn ␣=␣OLS␣
  or␣ WLS.”““
  rng = np. random . default_rng (seed)
  n = len(xs)
  idx = np. arange (n)
  boots = []
  for _ in range (n_boot):
  sel = rng. choice (idx, size =n, replace = True)
  try :
  if w is None :
  s, _, _ = fit_fn (xs[sel], ys[sel ])
  else :
  s, _, _ = fit_fn (xs[sel], ys[sel], w[sel ])
  if np. isfinite (s):
  boots . append (s)
  except Exception :
  pass
  if len(boots) == 0:
  return np.nan, (np.nan, np.nan), np. array ([])
  boots = np. array (boots, float)
  lo, hi = np. percentile (boots, [2.5, 97.5])
  return float (np. median (boots)), (float (lo), float (hi)), boots
*# ---------- parameter grid ----------*
SAVE_DIR = “./ capture_kinetics_outputs “
os. makedirs (SAVE_DIR, exist_ok = True)
fslows = np. round (np. geomspace (0.07, 0.35, 6), 5). tolist ()
kappas = np. round (np. geomspace (0.04, 0.2, 6), 5). tolist ()
ratios = [0.20, 0.35, 0.50, 0.65, 0.75] # avoid extreme edge
  near 1.0
REPLICATES = 16
NOISE_D = 2e -4
T_OBS = 360.0
records = [] # (f, k, r, delta, T_med, T_iqr, abscissa)
*# ---------- run grid with deterministic seeds ----------*
for f in fslows :
  for k in kappas :
  for r in ratios :
  delta = r*k
  tlocks = []
  for rep in range (REPLICATES):
  seed = deterministic_seed (f, k, r, rep) #
  deterministic per - cell seed
  phi, dt = simulate_adler (f, k, delta, D= NOISE_D,
  dt =1/200, T=T_OBS, seed = seed)
  fs = int (1/ dt)
  lam = lambda_trace (phi, fs=fs, W =10.0)
  tlocks . append (
  tlock_from_exp (lam, fs=fs, frac =0.8,
  entry_frac =0.15, entry_min_sec =5.0,
  fit_window_sec =20.0)
 )
  tlocks = np. array (tlocks, float)
  T_med = float (np. nanmedian (tlocks))
  T_iqr = float (np. nanpercentile (tlocks, 75) - np.
  nanpercentile (tlocks, 25))
  x = 1.0/(f*np. sqrt (max(k*k - (r*k)*(r*k), 1e -12)))
  records . append ((f, k, r, delta, T_med, T_iqr, x))
*# ---------- save CSV ----------*
csv_path = os. path . join (SAVE_DIR, “capture_kinetics_grid_median .
  csv “)
with open (csv_path, “w”, newline =““) as fh:
  w = csv. writer (fh)
  w. writerow ([“fslow “,” kappa “,” ratio “,” delta “,” Tlock_median “,”
  Tlock_IQR “,” abscissa “])
  w. writerows (records)
*# ---------- Figure 1: Lambda (t) example ----------*
plt . figure ()
plt . plot (lam)
plt . xlabel (“samples “)
plt . ylabel (“coherence ␣ proxy “)
plt . title (r” Example ␣$\ Lambda (t)$␣ proxy “)
plt . tight_layout ()
fig1_path = os. path . join (SAVE_DIR, “
  capture_lambda_timeseries_realistic .png”)
plt . savefig (fig1_path, dpi =220)
plt . show ()
*# ---------- log -log scaling (median per cell) ----------*
xs = np. array ([r[ -1] for r in records ], float)
ys = np. array ([r [4] for r in records ], float) # median
  T_lock
iqr = np. array ([r[5] for r in records ], float)
mask = np. isfinite (xs) & np. isfinite (ys) & (xs > 0) & (ys > 0)
xs_f, ys_f, iqr_f = xs[ mask ], ys[ mask ], iqr[ mask ]
*# OLS*
s_ols, b0_ols, r2_ols = loglog_fit_ols (xs_f, ys_f)
med_ols, (lo_ols, hi_ols), _ = bootstrap_ci (xs_f, ys_f,
  loglog_fit_ols, n_boot =1000, seed =2025)
*# WLS* :    *weights = 1/(IQR **2) (proxy for precision), regularized and*
     *clipped*
eps = np. maximum (1e -6, np. nanpercentile (iqr_f, 5))
w_raw = 1.0/ np. maximum (iqr_f, eps)**2
w = w_raw / np.max (w_raw)
w = np. clip (w, 0.05, 1.0)
s_wls, b0_wls, r2_wls = loglog_fit_wls (xs_f, ys_f, w)
med_wls, (lo_wls, hi_wls), _ = bootstrap_ci (xs_f, ys_f,
  loglog_fit_wls, w=w, n_boot =1000, seed =3030)
*# Plot + lines*
plt . figure ()
plt . loglog (xs_f, ys_f, ‘o’, alpha =0.8, label =“grid ␣ medians “)
xf = np. linspace (xs_f .min (), xs_f .max (), 300)
yf_ols = np.exp (b0_ols) * xf ** s_ols
yf_wls = np.exp (b0_wls) * xf ** s_wls
plt . loglog (xf, yf_ols, ‘-’, label =fr ‘OLS ␣ slope ␣{ s_ols :.2f},␣$R ^2$␣
  { r2_ols :.2 f}’)
plt . loglog (xf, yf_wls, ‘--’, label =fr ‘WLS␣ slope ␣{ s_wls :.2 f},␣$R ^2$
  ␣{ r2_wls :.2f}’)
plt . xlabel (r’$[f_ {\ rm␣ slow }\ sqrt {\ kappa ^2 -\ delta ^2}]^{ -1} $’)
plt . ylabel (r’$T_ {\ rm␣ lock }$␣(s)’)
plt . legend ()
plt . tight_layout ()
fig2_path = os. path . join (SAVE_DIR, “
  capture_scaling_loglog_realistic .png”)
plt . savefig (fig2_path, dpi =220)
plt . show ()
*# ---------- TXT with both fits + bootstrap CIs ----------*
slope_txt_path = os. path . join (SAVE_DIR, “capture_scaling_slope .txt
  “)
 with open (slope_txt_path, “w”) as fh:
fh. write (“Log -log ␣ scaling ␣for␣ T_lock ␣vs␣[ f_slow ␣*␣ sqrt (kappa ^2
␣-␣ delta ^2) ]^{ -1}\ n”)
fh. write (f” Cells ␣ used ␣(medians):␣{ xs_f . size }\n”)
fh. write (“\ nOLS ␣fit :\n”)
fh. write (f”␣␣ Slope :␣{ s_ols :.6f}\n␣␣R^2: ␣{ r2_ols :.6f}\n”)
fh. write (f”␣␣ Bootstrap ␣ median ␣ slope :␣{ med_ols :.6f}\n”)
fh. write (f”␣␣ Bootstrap ␣ 95%␣CI:␣[{ lo_ols :.6f},␣{ hi_ols :.6f}]\ n”
 )
fh. write (“\ nWLS ␣fit ␣(weights ␣=␣1/(IQR ^2),␣ clipped):\n”)
fh. write (f”␣␣ Slope :␣{ s_wls :.6f}\n␣␣R^2: ␣{ r2_wls :.6f}\n”)
fh. write (f”␣␣ Bootstrap ␣ median ␣ slope :␣{ med_wls :.6f}\n”)
fh. write (f”␣␣ Bootstrap ␣ 95%␣CI:␣[{ lo_wls :.6f},␣{ hi_wls :.6f}]\ n”
 )
fh. write (f”\ nReplicates ␣ per␣ cell :␣{ REPLICATES }\ nNoise ␣D:␣{
  NOISE_D },␣ Observation ␣T␣(s):␣{ T_OBS }\n”)
fh. write (“Entry ␣ criterion :␣ <15%␣of␣ plateau ␣ for␣ >=5␣s;␣fit␣
  window :␣20␣s;␣”
  “plateau ␣=␣ median ␣of␣ last ␣20␣s.\n”)
print (“Saved ␣ outputs ␣to:”, os. path . abspath (SAVE_DIR))
print (“␣-”, os. path . basename (fig1_path))
print (“␣-”, os. path . basename (fig2_path))
print (“␣-”, os. path . basename (csv_path))
print (“␣-”, os. path . basename (slope_txt_path))
~~~

### M.7 2/21 Arnold tongue in the reduced phase model

It implements the coupled-oscillator simulation used to generate Fig. J.1.

~~~
import numpy as np
from scipy . integrate import solve_ivp
import matplotlib . pyplot as plt
from matplotlib . colors import LogNorm
# --------------------------------------------------------------
*# Listing 7: numerical 2/21 Arnold tongue for the reduced phase*
model
#
# Purpose
# -------
# Scan the parameter plane (Omega = omega1 /omega2, K2) of a
# two - oscillator phase model and quantify how close the average
# frequency ratio rho = f1/f2 is to the theoretical 2/21 lock .
# The script produces a zoomed heatmap around the 2/21 tongue .
#
# Inputs (conceptual)
# -------------------
# omega2 :    reference natural frequency (set to 1.0 here).
# Omega_vals :    grid of frequency ratios omega1 / omega2 to scan .
# K2_vals :    grid of coupling strengths K2 to scan .
#
# Numerical settings
# ------------------
# t_span, t_eval :    integration time window and sampling for
     solve_ivp .
# rtol, atol :    solver tolerances .
#
# Output
# ------
# A PNG figure “Arnold_Tongue_2over21 .    png” containing :
#   - log - scaled heatmap of |rho - 2/21|
#   - cyan contour where |rho - 2/21|    = 0.005, approximating
# the effective 2/21 plateau .
# ------------------------------------------------------------
OUTPUT_FIG = “Arnold_Tongue_2over21 .png”
# -----------------------------------------------
*# Coupled phase - oscillator model*
# -----------------------------------------------
def coupled_oscillators (t, y, omega1, omega2, K2):
  “““
␣␣␣␣ Simple ␣two - oscillator ␣ phase ␣ model ␣ with ␣even - harmonic ␣ coupling .
␣␣␣␣ Parameters
␣␣␣␣ ----------
␣␣␣␣t␣:␣ float
␣␣␣␣␣␣␣␣ Time ␣(not␣ used ␣ explicitly ;␣ required ␣by␣ solve_ivp ␣ interface
 ).
␣␣␣␣y␣:␣ array_like,␣ shape ␣(2,)
␣␣␣␣␣␣␣␣ State ␣ vector ␣[theta1,␣ theta2 ].
␣␣␣␣omega1,␣ omega2 ␣:␣ float
␣␣␣␣␣␣␣␣ Natural ␣ angular ␣ frequencies ␣of␣ oscillators ␣1␣ and␣2.
␣␣␣␣K2␣:␣ float
␣␣␣␣␣␣␣␣Even - harmonic ␣ coupling 
␣ strength .
␣␣␣␣ Returns
␣␣␣␣ -------
␣␣␣␣ list ␣of␣ float
␣␣␣␣␣␣␣␣ Time ␣ derivatives ␣[ dtheta1 /dt,␣ dtheta2 /dt ].
␣␣␣␣”““
  theta1, theta2 = y
  dtheta1_dt = omega1 + K2 * np.sin (2 * (theta2 - theta1))
  dtheta2_dt = omega2
  return [ dtheta1_dt, dtheta2_dt ]
# ------------------------------------------------
*# Parameter scan around the 2/21 Arnold tongue*
# ------------------------------------------------
omega2 = 1.0    *# reference frequency (sets the time scale)*
rho_target = 2.0 / 21.0 # target locked ratio ~
  0.0952
Omega_vals = np. linspace (0.05, 0.14, 120) # Omega = omega1 / omega2
K2_vals = np. logspace (-3, -0.1, 80) # coupling strength K2:
  1e -3 … ~0.8
Omega_grid, K2_grid = np. meshgrid (Omega_vals, K2_vals)
rho_grid = np. zeros_like (Omega_grid)
*# Integration set -up*
t_span = (0.0, 1000.0)
t_eval = np. linspace (* t_span, 2000)
y0 = [0.0, 0.0]
*# Progress bookkeeping*
n_K2 = len(K2_vals)
n_Omega = len(Omega_vals)
total_points = n_K2 * n_Omega
counter = 0
print (
  “Scanning ␣ parameter ␣ grid ␣ around ␣ the␣ 2/21 ␣ lock ␣”
  f”({ n_K2 }␣x␣{ n_Omega }␣=␣{ total_points }␣ points)… “
)
for i, K2 in enumerate (K2_vals):
  for j, Omega in enumerate (Omega_vals):
  omega1 = Omega * omega2
     *# Integrate the coupled - oscillator system for this (Omega*,
     *K2)*
  sol = solve_ivp (
  coupled_oscillators, t_span, y0,
  args =(omega1, omega2, K2),
  t_eval =t_eval, rtol =1e -6, atol =1e -6, max_step =1.0
 )
  theta1, theta2 = sol.y
     *# Unwrap phases to count cycles reliably*
  phi1 = np. unwrap (theta1)
  phi2 = np. unwrap (theta2)
  cycles1 = (phi1 [ -1] - phi1 [0]) / (2 * np.pi)
  cycles2 = (phi2 [ -1] - phi2 [0]) / (2 * np.pi)
  # Average frequency ratio rho = f1/f2
  rho_grid [i, j] = cycles1 / cycles2 if cycles2 > 0 else np.
  nan
# Update progress counter
counter += 1
if counter % 500 == 0 or counter == total_points :
  frac = 100.0 * counter / float (total_points)
  print (f” Progress :␣{ counter }/{ total_points }␣ points ␣({
  frac :.1f}%)”)
print (“Parameter ␣ scan ␣ completed .␣ Building ␣ figure …”)
# Distance to the 2/21 plateau
delta_rho = np.abs(rho_grid - rho_target)
# ------------------------------------------------------------
# Plot : zoom around the 2/21 Arnold tongue
# ------------------------------------------------------------
plt . style . use(“default “)
fig, ax = plt . subplots (figsize =(10, 8))
# Heatmap of |rho - 2/21| in the (Omega, K2) plane
pcm = ax. pcolormesh (
  Omega_grid, K2_grid, delta_rho,
  shading =“auto “,
  norm = LogNorm (vmin =1e -4, vmax =1),
  cmap =“magma “
)
cbar = fig. colorbar (pcm, ax=ax)
cbar . set_label (r”$|\ rho␣-␣ 2/21| $”)
# Contour where |rho - 2/21| = tol, to highlight the effective
  plateau
tol = 0.005
cs = ax. contour (
  Omega_grid, K2_grid, delta_rho,
  levels =[ tol], # single contour level is fine for
  contour (not contourf)
  colors =“cyan “,
  linewidths =1.0
)
ax. clabel (
  cs,
  fmt ={ tol: r”$|\ rho␣-␣ 2/21| ␣=␣ %.3 f$” % tol },
  inline =True,
  fontsize =8
)
ax. set_xlabel (r”$\ Omega ␣=␣\ omega_1 ␣/␣\ omega_2$ “)
ax. set_ylabel (r” Coupling ␣ strength ␣ $K_2$ “)
ax. set_yscale (“log “)
ax. set_title (“Zoom ␣ around ␣the␣ 2/21 ␣ Arnold ␣ tongue “)
ax. grid (True, which =“both “, ls=“--”, alpha =0.3)
plt . tight_layout ()
plt . savefig (OUTPUT_FIG, dpi =300)
plt . show ()
print (f” Saved ␣ figure ␣to␣{ OUTPUT_FIG }”)
~~~

### M.8 Transverse cut through the 2/21 Arnold tongue

~~~
It implements the transverse scan in    *K*_2_ at fixed Ω = 2/21, used to generate Fig. J.2.
import numpy as np
from scipy . integrate import solve_ivp
import matplotlib . pyplot as plt
# ============================================================
# Listing 8: transverse cut through the 2/21 Arnold tongue
#
# Purpose
# -------
# For a fixed drive ratio Omega = omega1 / omega2 = 2/21, scan the
# coupling strength K2 and estimate the average frequency ratio
# f1/f2 in a simple two - oscillator phase model with even -
  harmonic
# coupling . The script produces a 1D “cut” showing how the
  system
# transitions from the 2/21 plateau toward near 1:1 lock as K2
# increases .
#
# Conceptual inputs
# -----------------
# omega2 : reference natural frequency (set to 1.0 here).
# Omega_target : target ratio omega1 / omega2 = 2/21 for the cut .
# K2_values : log - spaced grid of coupling strengths K2.
#
# Numerical settings
# ------------------
# t_span, t_eval : integration window and sampling for solve_ivp
  .
# rtol, atol : tolerances for the ODE solver .
#
# Output
# ------
# A PNG file “Arnold_tongue_cut_2over21 . png” containing :
# - the simulated average frequency ratio f1/f2 vs K2,
# - an orange band where |f1/f2 - 2/21| < 5e -3,
# - a gold band indicating the near -1:1 lock region,
# - a vertical line marking K2 ~ 0.5 (the value used in the
# biological reduced model).
# ============================================================
OUTPUT_FIG = “Arnold_tongue_cut_2over21 .png”
# ============================================================
# Coupled phase oscillators with even (mod pi) coupling
# ============================================================
def coupled_oscillators (t, y, omega1, omega2, K2):
  “““
␣␣␣␣Two ␣ phase ␣ oscillators ␣ with ␣even - harmonic ␣ coupling .
␣␣␣␣ Parameters
␣␣␣␣ ----------
␣␣␣␣t␣:␣ float
␣␣␣␣␣␣␣␣ Time ␣(unused,␣ included ␣for␣ solve_ivp ␣ compatibility).
␣␣␣␣y␣:␣ array_like,␣ shape ␣(2,)
␣␣␣␣␣␣␣␣ Current ␣ phases ␣[theta1,␣ theta2 ].
␣␣␣␣ omega1 ␣:␣ float
␣␣␣␣␣␣␣␣ Natural ␣ frequency ␣of␣ oscillator ␣1.
␣␣␣␣ omega2 ␣:␣ float
␣␣␣␣␣␣␣␣ Natural ␣ frequency ␣of␣ oscillator ␣2.
␣␣␣␣K2␣:␣ float
␣␣␣␣␣␣␣␣Even - harmonic ␣ coupling ␣ strength .
␣␣␣␣ Returns
␣␣␣␣ -------
␣␣␣␣ dtheta_dt ␣:␣ list [ float ]
␣␣␣␣␣␣␣␣ Time ␣ derivatives ␣[ dtheta1 /dt,␣ dtheta2 /dt ].
␣␣␣␣”““
     theta1, theta2 = y
     dtheta1_dt = omega1 + K2 * np.sin (2.0 * (theta2 - theta1))
     dtheta2_dt = omega2
     return [ dtheta1_dt, dtheta2_dt ]
# ============================================================
# Parameter setup : transverse cut at Omega = 2/21
# ============================================================
omega2 = 1.0 # reference frequency
Omega_target = 2.0 / 21.0 # Omega = omega1 / omega2 for the
  2/21 tongue
omega1 = Omega_target * omega2 # corresponding omega1
# Coupling strengths to scan (log - spaced : weak -> strong)
K2_values = np. logspace (-3, 1, 200)
# Storage for average frequency ratios f1/f2
ratios = np. full_like (K2_values, np.nan, dtype = float)
# Integration settings
t_span = (0.0, 1000.0)
t_eval = np. linspace (t_span [0], t_span [1], 2000)
y0 = (0.0, 0.0)
print (f” Scanning ␣K2␣for␣ Omega ␣=␣ 2/21 ␣~␣{ Omega_target :.5f}␣…”)
for i, K2 in enumerate (K2_values):
  sol = solve_ivp (
       coupled_oscillators,
       t_span,
       y0,
       args =(omega1, omega2, K2),
       t_eval =t_eval,
       rtol =1e -6,
       atol =1e -6,
       max_step =1.0,
 )
  if not sol. success :
      continue
  # Unwrap phases to count cycles
  theta1 = np. unwrap (sol.y [0])
  theta2 = np. unwrap (sol.y [1])
  
   cycles1 = (theta1 [ -1] - theta1 [0]) / (2.0 * np.pi)
  cycles2 = (theta2 [ -1] - theta2 [0]) / (2.0 * np.pi)
  if cycles2 > 0:
  ratios [i] = cycles1 / cycles2
print (“Done .␣ Building ␣ figure ␣…”)
# ============================================================
# Plot : transverse cut through the 2/21 Arnold tongue
# ============================================================
plt . style . use(“default “)
fig, ax = plt . subplots (figsize =(10, 7))
mask_good = ~np. isnan (ratios)
K2_plot = K2_values [ mask_good ]
ratio_plot = ratios [ mask_good ]
# Main curve : simulated average frequency ratio f1/f2
ax. plot (
    K2_plot,
    ratio_plot,
    “o-”,
    color =“cyan “,
    markersize =4,
    linewidth =1.0,
    label =r” Simulation ␣ $f_1 / f_2$ “,
)
# ----- Highlight the 2/21 plateau band -----
ratio_221 = 2.0 / 21.0
tol_221 = 5e -3 # tolerance around 2/21
band_low = ratio_221 - tol_221
band_high = ratio_221 + tol_221
mask_221 = np. abs(ratio_plot - ratio_221) < tol_221
ax. fill_between (
    K2_plot,
    band_low,
    band_high,
    where = mask_221,
    color =“orange “,
    alpha =0.25,
    interpolate =True,
    label =r”$|f_1/ f_2␣-␣ 2/21| ␣<␣5\ times ␣ 10^{ -3} $”,
)
ax. axhline (
  ratio_221,
  color =“red “,
  linestyle =“--”,
  linewidth =1.5,
)
# ----- Highlight the near 1:1 lock region -----
tol_11 = 1e -2
band11_low = 1.0 - tol_11
band11_high = 1.0 + tol_11
mask_11 = np.abs (ratio_plot - 1.0) < tol_11
ax. fill_between (
   K2_plot,
   band11_low,
   band11_high,
   where = mask_11,
   color =“gold “,
   alpha =0.18,
   interpolate =True,
   label =r” near ␣$1 {:}1 $␣ lock “,
)
ax. axhline (
  1.0,
  color =“gold “,
  linestyle =“:”,
  linewidth =1.0,
)
# -- Mark the theoretical K2 ~ 0.5 used in the biological model --
K2_theoretical = 0.5
ax. axvline (
  x= K2_theoretical,
  color =“deepskyblue “,
  linestyle =“--”,
  linewidth =1.5,
)
# Place the label slightly LEFT of the vertical line (on a log x-
  axis)
ax. text (
  K2_theoretical / 1.2,
  0.25,
  r” $K_2 ␣\ simeq ␣ 0.5$”,
  color =“deepskyblue “,
  rotation =90,
  va=“bottom “,
  ha=“right “,
)
# ============================================================
# Axes, labels, and layout
# ============================================================
ax. set_xscale (“log “)
ax. set_xlim (K2_values . min (), K2_values .max ())
ax. set_ylim (0.0, 1.05)
ax. set_xlabel (r” Coupling ␣ strength ␣ $K_2$ “)
ax. set_ylabel (r” Average ␣ frequency ␣ ratio ␣ $f_1 / f_2$ “)
ax. set_title (r” Transverse ␣cut␣of␣the␣ Arnold ␣ tongue ␣at␣$\ Omega ␣=␣
  2/21 $”)
ax. grid (True, which =“both “, ls=“--”, alpha =0.5)
ax. legend (loc=“lower ␣ right “, fontsize =9)
plt . tight_layout ()
plt . savefig (OUTPUT_FIG, dpi =300)
plt . show ()
print (f” Saved ␣ figure ␣to␣{ OUTPUT_FIG }”)
~~~

### M.9 Neutral relaxation P1: synthetic validation of Appendix L

The Python script below implements the synthetic neutral-relaxation test of App. L in the P1 regime. It generates a slow-phase signal    *ψ*(*t*), detects neutral moments    *t*_*n*_, constructs a noisy neutral charge-asymmetry sequence {Δ*Q*_*n*_} with an exponentially contracting envelope, estimates    *f*_slow_ and    *κ*, and outputs the summary and the two diagnostic panels shown in Fig. L.1.

~~~
import numpy as np
import matplotlib . pyplot as plt
“““
Listing ␣9.␣ Synthetic ␣ validation ␣of␣ Appendix ␣L:␣ neutral - charge ␣
  relaxation ␣in␣ regime ␣P1.
What ␣ this ␣ script ␣ does
---------------------
1.␣ Builds ␣a␣ synthetic ␣slow - phase ␣ signal ␣psi(t)␣ with :
␣␣␣-␣ slow ␣ frequency ␣ f_slow_true
␣␣␣-␣ decaying ␣ amplitude ␣(so␣ phase ␣ fluctuations ␣are ␣ visible ␣at␣
  early ␣ times)
␣␣␣-␣ small ␣ additive ␣ noise .
2.␣ Detects ␣ neutral ␣ moments ␣t_n␣as␣zero - crossings ␣of␣psi(t)␣ with ␣
  positive ␣ slope .
3.␣ Constructs ␣a␣ neutral - charge ␣ sequence ␣ DeltaQ_n ␣ whose ␣ absolute ␣
  value ␣ decays
␣␣␣ exponentially ␣ with ␣ rate ␣ kappa_true,␣ with ␣ alternating ␣ sign ␣(left
  / right ␣ bias).
4.␣ Fits ␣an␣ exponential ␣ envelope ␣to␣| DeltaQ_n |␣ across ␣ neutral ␣ index
  ␣n␣to␣ estimate
␣␣␣ kappa_eff ␣ and␣ checks ␣the ␣ relation
␣␣␣␣␣␣␣ T_relax ␣␣~␣␣1␣/␣(f_slow_est ␣*␣ kappa_eff)
5.␣ Produces ␣two␣ diagnostic ␣ figures :
␣␣␣-␣ Figure ␣1:␣ psi(t)␣ with ␣ neutral ␣ moments ␣ overlaid .
␣␣␣-␣ Figure ␣2:␣ DeltaQ_n ␣vs␣t_n ␣ with ␣the␣ theoretical ␣ envelope ␣|
  DeltaQ_n |.
Additional ␣ output
-----------------
-␣A␣ text ␣ file ␣ with ␣a␣ compact ␣ summary ␣of␣the␣ main ␣ quantities ␣and
␣␣the␣ names ␣of␣the␣PNG␣ files ␣ saved .
Inputs ␣(all␣set␣in␣the␣ parameter ␣ block)
---------------------------------------
T_total ␣␣␣␣␣␣␣:␣ total ␣ simulation ␣ time ␣[s]
dt␣␣␣␣␣␣␣␣␣␣␣␣:␣ time ␣ step ␣[s]
f_slow_true ␣␣␣:␣ true ␣ slow ␣ frequency ␣[Hz]
kappa_true ␣␣␣␣:␣ true ␣ contraction ␣ rate ␣per␣ neutral ␣ step ␣(
  dimensionless)
psi0 ␣␣␣␣␣␣␣␣␣␣:␣ initial ␣ phase ␣ amplitude ␣[rad]
tau_phase ␣␣␣␣␣:␣ decay ␣ time ␣of␣ phase ␣ fluctuations ␣[s]
noise_psi ␣␣␣␣␣:␣ noise ␣ level ␣ added ␣to␣psi(t)
A0␣␣␣␣␣␣␣␣␣␣␣␣:␣ initial ␣ magnitude ␣of␣ DeltaQ_n
noise_Q ␣␣␣␣␣␣␣:␣ noise ␣ level ␣in␣ DeltaQ_n
Outputs
-------
-␣ Printed ␣ summary ␣of:
␣␣␣␣ n_neutral,␣ f_slow_true,␣ f_slow_est,
␣␣␣␣ kappa_true,␣ kappa_eff,␣ T_relax_est,␣1/(f_slow_true ␣*␣
  kappa_true)
-␣ One␣ text ␣ file ␣ with ␣the␣ same ␣ summary .
-␣ Two␣PNG␣ figures :
␣␣␣␣1)␣ phase_neutrals_png
␣␣␣␣2)␣ deltaQ_envelope_png
“““
# Optional : fix random seed for reproducibility
np. random . seed (0)
# ------------ parameters ----------
T_total = 1600.0  # total time [s]
   dt = 0.25         # time step [s]
f_slow_true = 0.05 # slow frequency [Hz] (period ~ 20 s)
kappa_true = 0.07 # contraction per neutral step (
  dimensionless)
psi0 = 0.3        # initial amplitude of phase
fluctuations [rad ]
     tau_phase = 400.0 # decay time of phase fluctuations [s]
noise_psi = 0.015 # noise level added to psi(t) [ rad]
A0 = 5.0          # initial magnitude of DeltaQ_n (
  arbitrary units)
noise_Q = 0.15    # noise level in DeltaQ_n
                  # Output filenames
summary_filename = “appendixL_P1_summary .txt”
     phase_png_filename = “appendixL_P1_phase_neutrals .png “
deltaQ_png_filename = “appendixL_P1_deltaQ_envelope . png”
# ------- build time axis --------
t = np. arange (0.0, T_total, dt)
nT = len(t)
  # ------- synthetic phase signal psi(t) --------
  # Exponentially decaying envelope for visible early fluctuations
envelope = psi0 * np.exp (-t / tau_phase)
  # Slow oscillation at f_slow_true, plus small noise
psi = envelope * np. sin (2.0 * np.pi * f_slow_true * t) \
  + noise_psi * np. random . randn (nT)
  # ------- detect neutral moments t_n -------
 # Zero - crossings of psi (t) from negative to positive with positive
  slope
psi_prev = psi [: -1]
psi_next = psi [1:]
dpsi = psi_next - psi_prev
# indices where psi crosses 0 with positive slope
idx_neutral = np. where ((psi_prev < 0.0) & (psi_next >= 0.0) & (
  dpsi > 0.0))[0]
# refine zero - crossing times by linear interpolation
t_n = t[ idx_neutral ] - psi_prev [ idx_neutral ] * dt / (dpsi [
  idx_neutral ] + 1e -12)
n_neutral = len(t_n)
# estimated slow frequency from neutral spacing
if n_neutral > 1:
  f_slow_est = 1.0 / np. mean (np. diff (t_n))
else :
  f_slow_est = np.nan
# ------- synthetic neutral - charge sequence DeltaQ_n -------
# index of neutral crossings (0,1,2,…)
n_index = np. arange (n_neutral)
# ideal exponential decay per neutral step, with alternating sign
DeltaQ_ideal = A0 * np.exp(- kappa_true * n_index) * ((-1.0) **
  n_index)
# add small noise
DeltaQ_n = DeltaQ_ideal + noise_Q * np. random . randn (n_neutral)
# ------- fit exponential envelope to | DeltaQ_n | -------
absQ = np.abs(DeltaQ_n)
# avoid log (0): keep only points above a small threshold
mask = absQ > 1e -3
n_fit = n_index [ mask ]
logQ = np.log (absQ [ mask ])
# linear fit: log|Q_n | = c - kappa_eff * n
slope, intercept = np. polyfit (n_fit, logQ, 1)
kappa_eff = -slope
# relaxation times
T_relax_est = 1.0 / (kappa_eff * f_slow_est) if np. isfinite (
  f_slow_est) else np.nan
T_relax_theory = 1.0 / (kappa_true * f_slow_true)
# ------- build theoretical envelope for plotting --------
envelope_theory = np.abs (A0 * np.exp(- kappa_true * n_index))
# ------- print summary -------
print (f” n_neutral ␣␣␣␣␣␣␣␣=␣{ n_neutral }”)
print (f” f_slow_true ␣␣␣␣␣␣=␣{ f_slow_true :.5f}␣Hz”)
print (f” f_slow_est ␣␣␣␣␣␣␣=␣{ f_slow_est :.5f}␣Hz”)
print (f” kappa_true ␣␣␣␣␣␣␣=␣{ kappa_true :.5f}”)
print (f” kappa_eff ␣(fit)␣␣=␣{ kappa_eff :.5f}”)
print (f” T_relax_est ␣␣␣␣␣␣=␣{ T_relax_est :.2f}␣s␣␣␣(from ␣ kappa_eff ␣&
  ␣ f_slow_est)”)
print (f” T_relax_theory ␣␣␣=␣{ T_relax_theory :.2f}␣s␣␣␣(=␣ 1/(
  f_slow_true ␣*␣ kappa_true))”)
# ------ write summary to text file -------
summary_lines = [
  “Synthetic ␣ validation ␣of␣ Appendix ␣L:␣ neutral - charge ␣ relaxation
  ␣in␣ regime ␣P1\n”,
  “\n”,
  f” n_neutral ␣␣␣␣␣␣␣␣=␣{ n_neutral }\n”,
  f” f_slow_true ␣␣␣␣␣␣=␣{ f_slow_true :.5f}␣Hz\n”,
  f” f_slow_est ␣␣␣␣␣␣␣=␣{ f_slow_est :.5f}␣Hz\n”,
  f” kappa_true ␣␣␣␣␣␣␣=␣{ kappa_tr  ue :.5f}\n”,
  f” kappa_eff ␣(fit)␣␣=␣{ kappa_eff :.5f}\n”,
  f” T_relax_est ␣␣␣␣␣␣=␣{ T_relax_est :.2f}␣s␣␣␣(from ␣ kappa_eff ␣&␣
  f_slow_est)\n”,
  f” T_relax_theory ␣␣␣=␣{ T_relax_theory :.2f}␣s␣␣␣(=␣1/(
  f_slow_true ␣*␣ kappa_true))\n”,
“\n”,
  “PNG␣ output ␣ files :\n”,
  f”␣␣-␣ phase ␣ and␣ neutral ␣ crossings :␣{ phase_png_filename }\n”,
  f”␣␣-␣ neutral - charge ␣ sequence :␣␣␣␣␣{ deltaQ_png_filename }\n”
]
with open (summary_filename, “w”) as f:
    f. writelines (summary_lines)
# ------- plotting : Figure 1 (phase + neutrals) ------
fig1, ax1 = plt. subplots (figsize =(10, 4))
ax1 . plot (t, psi, “k-”, lw =1.0, label =r”$\ psi(t)$”)
ax1 . axhline (0.0, color =“r”, ls=“--”, alpha =0.5, label =“neutral ␣
   line “)
ax1 . scatter (t_n, np. zeros_like (t_n), color =“red”, s=20,
  zorder =3, label =“neutral ␣ moments ␣ $t_n$ “)
ax1 . set_ylabel (r”$\psi (t)$␣[ rad]”)
ax1 . set_title (“P1␣ regime ␣-␣ geometric ␣ neutrality ␣ and␣ neutral ␣
  crossings “)
ax1 . legend (loc=“upper ␣ right “)
fig1 . tight_layout ()
fig1 . savefig (phase_png_filename, dpi =300)
# ------ plotting : Figure 2 (DeltaQ + envelope) ------
fig2, ax2 = plt. subplots (figsize =(10, 4))
markerline, stemlines, baseline = ax2. stem (t_n, DeltaQ_n)
markerline . set_color (“b”)
stemlines . set_color (“b”)
baseline . set_color (“k”)
ax2 . plot (t_n, envelope_theory, “g--”, lw =1.5,
  label =r”$|\ Delta ␣Q_n|$␣ envelope ␣(theory)”)
ax2 . set_xlabel (“time ␣t␣[s]”)
ax2 . set_ylabel (r”$\ Delta ␣ Q_n$ ␣(a.u.)”)
ax2 . set_title (
   “P1␣(healthy ␣ locking,␣ contracting ␣ envelope)\n”
   f” f_slow_true ␣=␣{ f_slow_true :.3f}␣Hz,␣”
   f” f_slow_est ␣~␣{ f_slow_est :.3 f}␣Hz,␣”
   f” kappa_true ␣=␣{ kappa_true :.3 f},␣”
   f” kappa_eff ␣~␣{ kappa_eff :.3 f},␣”
   f” T_relax_est ␣~␣{ T_relax_est :.1f}␣s”
)
ax2 . legend (loc=“upper ␣ right “)
fig2 . tight_layout ()
fig2 . savefig (deltaQ_png_filename, dpi =300)
# If you still want to see the figures interactively, keep this :
plt . show ()
~~~

### M.10 Minimal Python implementation of the P4 drift model

To illustrate the scaling law in Prediction P4, the following Python script implements the minimal drift model    *h*_*n*+1_ =    *h*_*n*_ +    *η δ* with a fixed threshold |*h*_*n*_| ≥*h*_th_. For each detuning    *δ*, it records the number of neutral steps required to reach the threshold, writes the corresponding trajectories to CSV files, and generates the summary plot `P4_simple_scaling_summary.png` used in Fig. L.2.

~~~
 import numpy as np
 import pandas as pd
 import matplotlib . pyplot as plt
 plt . style . use(“default “)
# ------------------------------------------------------------
# Minimal drift model for Prediction P4
#
# For each detuning delta, we evolve a slow phenotypic
# coordinate h_n according to
#
# h_{n+1} = h_n + eta * delta
#
# and record the number of neutral steps needed for | h_n|
# to reach a fixed threshold h_threshold . The resulting
# N_cycles_star is plotted against 1/| delta | to test the
# scaling law T_dev ~ 1 / (f_slow * | delta |).
# ------------------------------------------------------------
# Ideal 2/21 locking (used only to report rho = rho_0 + delta)
rho_0 = 2.0 / 21.0
# Detunings to sample
 deltas = [0.0020, 0.0050, 0.0100]
 N_max = 50000  # max number of neutral steps
# Phenotypic coordinate h_n
eta = 1.0 # accumulation efficiency per neutral
  step
h_threshold = 1.0  # threshold for a “macroscopic “
  change in h_n
# -----------------------------
# Simulation loop over deltas
# -----------------------------
summary_rows = []
for delta in deltas :
    rho = rho_0 + delta # corresponding rotation number (for
        reference)
h = 0.0  # initial phenotypic state
n_list = []
h_list = []
N_cycles_star = None # neutral cycles needed to hit |h_n|
  >= h_threshold
for n in range (N_max):
   n_list . append (n)
   h_list . append (h)
   # Simple drift : detuning produces a constant bias per
   neutral step
 h = h + eta * delta
   if N_cycles_star is None and abs(h) >= h_threshold :
  # Convert neutral steps n into 2/21 - cycles for
  comparison with theory
  N_cycles_star = n / 21.0
# Save per - delta trajectory of h_n
traj_df = pd. DataFrame ({
  “n”: n_list,
  “h_n”: h_list,
})
csv_name = f” P4_simple_trajectory_delta_ { delta :.4 f}. csv”
traj_df . to_csv (csv_name, index = False)
# Plot h_n versus neutral step n for this value of delta
fig, ax = plt . subplots (figsize =(8, 4))
ax. plot (n_list, h_list)
ax. axhline (+ h_threshold, ls=“--”, color =“gray “)
ax. axhline (- h_threshold, ls=“--”, color =“gray “)
ax. set_xlabel (“Neutral ␣ step ␣n”)
ax. set_ylabel (“h_n “)
ax. set_title (f” Minimal ␣P4␣ drift ␣ model ␣(delta ␣=␣{ delta :.4f})”)
fig . tight_layout ()
fig . savefig (f” P4_simple_trajectory_delta_ { delta :.4f}. png”, dpi
  =300)
plt . close (fig)
inv_delta = 1.0 / abs (delta)
summary_rows . append ({
  “delta “: delta,
  “rho”: rho,
  “N_cycles_star “: N_cycles_star,
  “inv_delta “: inv_delta,
})
# -----------------------------
# Summary CSV and scaling plot
# -----------------------------
summary_df = pd. DataFrame (summary_rows)
summary_df . to_csv (“P4_simple_scaling_summary .csv”, index = False)
print (summary_df)
# Drop rows where the threshold was not reached within N_max steps
valid = summary_df . dropna (subset =[“N_cycles_star “])
plt . figure (figsize =(6, 4))
plt . scatter (valid [“inv_delta “], valid [“N_cycles_star “], label =“
  simulation “)
plt . plot (valid [“inv_delta “], valid [“N_cycles_star “], “--”, alpha
  =0.5)
plt . xlabel (“1␣/␣| delta |”)
plt . ylabel (“N_cycles *␣(neutral ␣ cycles ␣to␣|h_n |␣ >=␣ threshold)”)
plt . title (“P4␣ simple ␣ scaling :␣ developmental ␣ cycles ␣vs␣ detuning “)
plt . tight_layout ()
plt . savefig (“P4_simple_scaling_summary .png”, dpi =300)
plt . show ()
~~~

## N Glossary of symbols

This appendix summarizes the main symbols used in the slow-phase locking analysis, the coherence readout Λ, and the phase-targeted entrainment protocol. Unless noted otherwise, quantities are evaluated under neutral-moment strobing (even parity) and at the mesoscale wavenumber *k*^*^ = *π/*ℓ.

## Acknowledgments

This manuscript was prepared with the assistance of AI-based editing and drafting tools. The author retains full responsibility for the analysis, interpretation, and any errors.

## Footnotes

1 For the canonical Adler equation the exact prefactor for deterministic capture from a uniform phase is 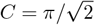; we keep *C* explicit to absorb implementation details (windowing, entry criterion).

